# Viral infection and virophage co-infection induce distinct remodeling of host lipidome

**DOI:** 10.64898/2026.09.28.754671

**Authors:** Guy Schleyer, Max Emil Schön, Christian Hertweck, Matthias G. Fischer

## Abstract

Virophages are small double-stranded DNA viruses that co-infect eukaryotic cells along with giant viruses. They are widespread in aquatic ecosystems and can influence microbial population dynamics by acting as an adaptive antiviral defense system, limiting production of giant virus progeny. Giant viruses extensively reprogram host metabolism to support their replication, yet little is known about how this process is altered by virophages. Characterizing these alterations may reveal the metabolic mechanisms underlying virophage-mediated changes in infection outcomes. Here, we used one of the few established host–giant virus–virophage experimental systems, consisting of the marine heterotrophic flagellate *Cafeteria burkhardae*, its giant virus CroV, and the virophage mavirus, to investigate lipid remodeling during viral infection and virophage co-infection. We found that infection by the giant virus strongly induced triacylglycerols containing furan fatty acids (FuFAs), a class of potent antioxidants that may protect infected cells and newly formed virions from oxidative damage. These lipids were also detected in purified giant virus particles, linking their accumulation to virus production. Co-infection with the virophage reduced the FuFA response and was instead characterized by pronounced accumulation of the betaine lipid diacylglyceryltrimethylhomoserine (DGTS). Most of the induced DGTS species were also present in the giant virus particles, suggesting that their accumulation during co-infection may be associated with altered giant virus production and lipid utilization. Together, these findings show that virophage co-infection reshapes giant virus-induced remodeling of the host lipidome, revealing a potential metabolic basis for their effects on virus production and infection outcomes.

## Introduction

Viruses are major drivers of microbial ecology and biogeochemical cycling in the oceans through their impact on host physiology, cellular metabolism and microbial mortality (1–3). Among them, members of the phylum *Nucleocytoviricota*, commonly referred to as giant viruses, infect a broad diversity of protists and have profound ecological consequences on microbial food webs and nutrient recycling (4–7). Giant viruses possess unusually large genomes encoding an extensive repertoire of metabolic genes. This genetic capacity enables them to actively reprogram host metabolism rather than relying solely on host biosynthetic machinery. Such metabolic rewiring supports viral genome replication, virion assembly, and the production of large numbers of progeny virions (7–9). Some giant viruses are themselves susceptible to infection by virophages, small double-stranded DNA viruses that exploit giant virus replication within cytoplasmic viral factories (10). During co-infection, virophages depend on the transcriptional machinery of the giant virus, often reducing giant virus progeny production while enhancing survival of the host population (10–13). Although the ecological and evolutionary consequences of virophages have received considerable attention, little is known about how co-infection redirects giant virus-induced remodeling of host metabolism.

One of the few experimentally characterized host–giant virus–virophage systems is the marine heterotrophic flagellate *Cafeteria burkhardae* (formerly *C. roenbergensis*) with its lytic giant virus Cafeteria roenbergensis virus (CroV) and its virophage mavirus. Following infection, CroV establishes cytoplasmic viral factories and completes its replication cycle within approximately 24 h, whereas mavirus exploits these viral factories to replicate (Fig. 1A), thereby reducing CroV progeny production during subsequent rounds of infection and increasing host population survival (11, 12, 14). The CroV–mavirus interaction therefore provides a unique opportunity to investigate how co-infection by a virophage impacts infection-associated metabolic reprogramming.

**Figure 1:**
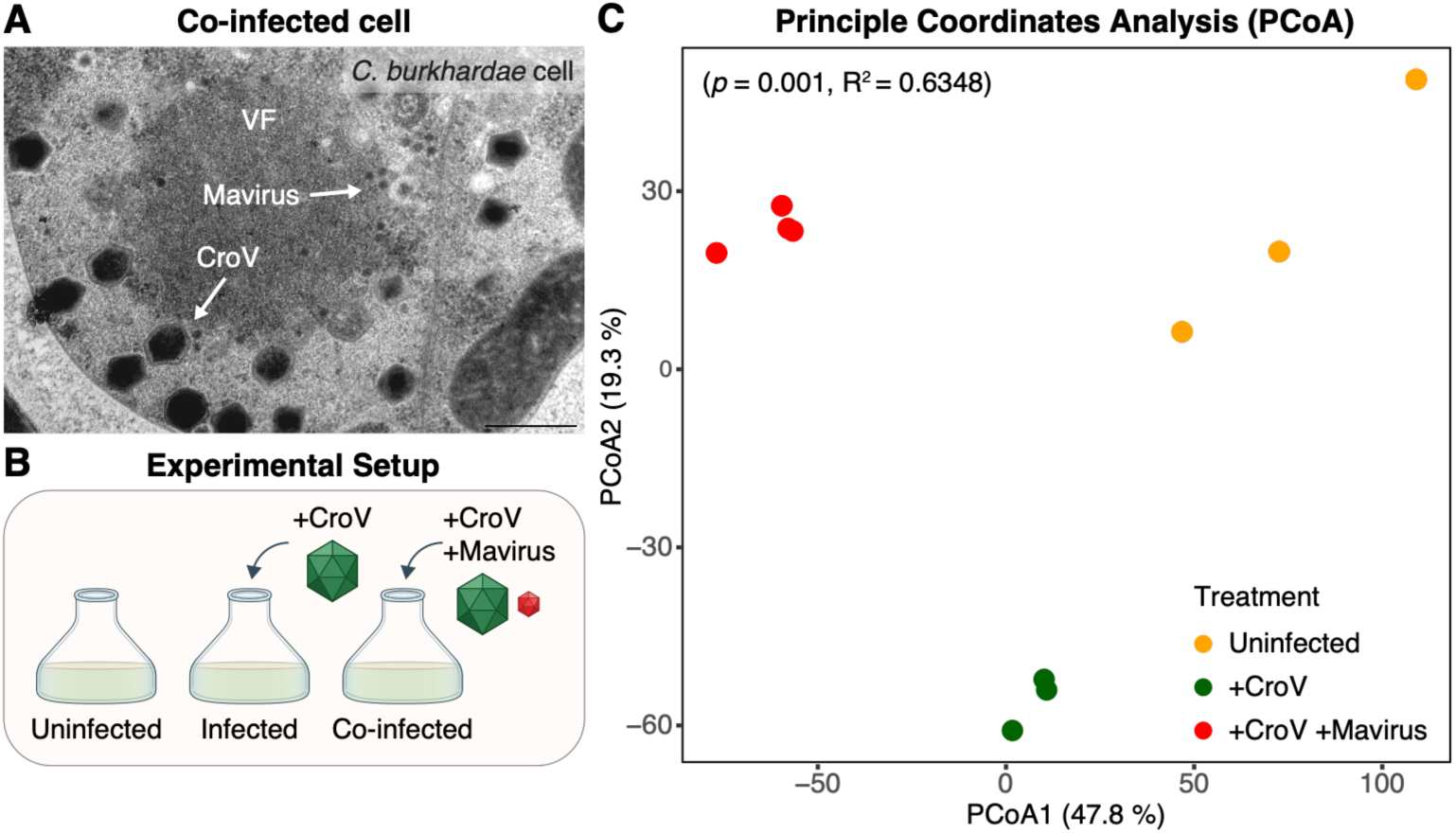
Untargeted LC-MS-based lipidomics analysis of uninfected, infected and co-infected *C. burkhardae*. (A) Transmission electron microscopy of a *C. burkhardae* cell co-infected with CroV and mavirus at 24 hpi. The CroV viral factory (VF) consists of large CroV particles and small mavirus particles. Scale bar = 500 nm. Reproduced and modified from ref (13) with permission from AAAS. (B) Experimental Setup used to analyze the lipid profile of *C. burkhardae* cells following infection with CroV and co-infection with CroV and mavirus. (C) Principal Coordinates Analysis (PCoA) of uninfected *C. burkhardae* cells (‘Uninfected’, orange, *n* = 3), cells infected with CroV (‘+CroV’, green, *n* = 3), and cells co-infected with CroV and mavirus (‘+CroV +Mavirus’, red, *n* = 4) based on untargeted lipidomics (8,821 mass features). Percentage of explained variance is indicated per coordinate.

Because giant virus replication depends on extensive membrane remodeling for viral factory formation and virion assembly, lipid metabolism provides a particularly informative lens on this reprogramming. Notably, while CroV virions contain an internal lipid membrane beneath the protein capsid, mavirus virions lack a lipid membrane (15, 16). Beyond serving as structural components of cellular and viral membranes, lipids also regulate membrane dynamics, signaling and energy storage, making them central to infection and virus release (17, 18). Accordingly, the few eukaryotic host–giant virus systems studied to date have revealed extensive remodeling of the host lipidome, including changes in phospholipids, sphingolipids, betaine lipids and neutral lipids. These lipids are closely linked to infection progression, host susceptibility, virion composition and cell fate (19–27). In some cases, membrane-containing viruses selectively enrich specific host- and virus-derived lipids within their virions rather than simply reflecting the composition of host membranes, indicating active remodeling of cellular lipid metabolism to support virus production (19, 20, 23, 24, 28). Nevertheless, whether and how lipid remodeling is rewired following virophage co-infection remains unknown.

Here, we report the distinct remodeling of the host lipidome following giant virus infection and virophage co-infection. We identify the most differential lipid classes in each infection scenario and study their occurrence in the giant virus virions. Together, our results establish that virophage co-infection redirects infection-associated lipid remodeling and expand viral lipidomics to multipartite host–virus–virophage systems.

## Results

To characterize lipid remodeling during viral infection and co-infection, we analyzed the lipidome of *C. burkhardae* under three conditions: (i) uninfected cells, (ii) cells infected with CroV, and (iii) cells co-infected with CroV and mavirus (Fig. 1B). Based on prior characterization of infection dynamics in this system (14), samples were collected 24 h post infection (hpi), corresponding to an advanced stage of CroV infection prior to complete host lysis. Intact polar lipids (IPLs) were extracted and analyzed using a liquid chromatography-mass spectrometry (LC-MS)-based untargeted lipidomics workflow. An unsupervised analysis using principal coordinates analysis (PCoA, Fig. 1C) was applied to obtain an overview of infection-associated changes, separating the samples according to the treatments. The first PCoA axis (47.8%) accounted for the difference between the three groups, while the second PCoA axis (19.3%) revealed further differences between uninfected and co-infected cells, and cells infected with CroV. Such separation indicates that lipid modulation in cells infected with CroV is significantly different than in those co-infected with both viruses.

To target mass features that were differential between the treatments, *k*-means clustering (*k* = 3, Fig. S1) was applied, followed by comparative statistical analysis (one-way repeated measures ANOVA, *p* < 0.01). This approach yielded three clusters, corresponding to the different treatment, and 1,172 differential mass features (Fig. S2), of which 118 exhibited a fold change ≥100 between at least two treatments. Subsequent feature deconvolution and manual curation grouped these mass features into 101 putative lipid species. Of these, 72 were assigned a putative annotation based on accurate mass, MS/MS fragmentation patterns, and comparison to lipid databases, while the other features remained unclassified (Table S1). Two-dimensional hierarchical clustering was then applied to this subset of 101 putative lipids species, maintaining the separation between treatments (Fig. 2). The putative lipid species were grouped into three clusters: (1) lipids with higher abundance in uninfected cells (19 species); (2) lipids with higher abundance in cells infected with CroV (30 species); and (3) lipids with higher abundance in cells co-infected with CroV and mavirus (52 species).

**Figure 2:**
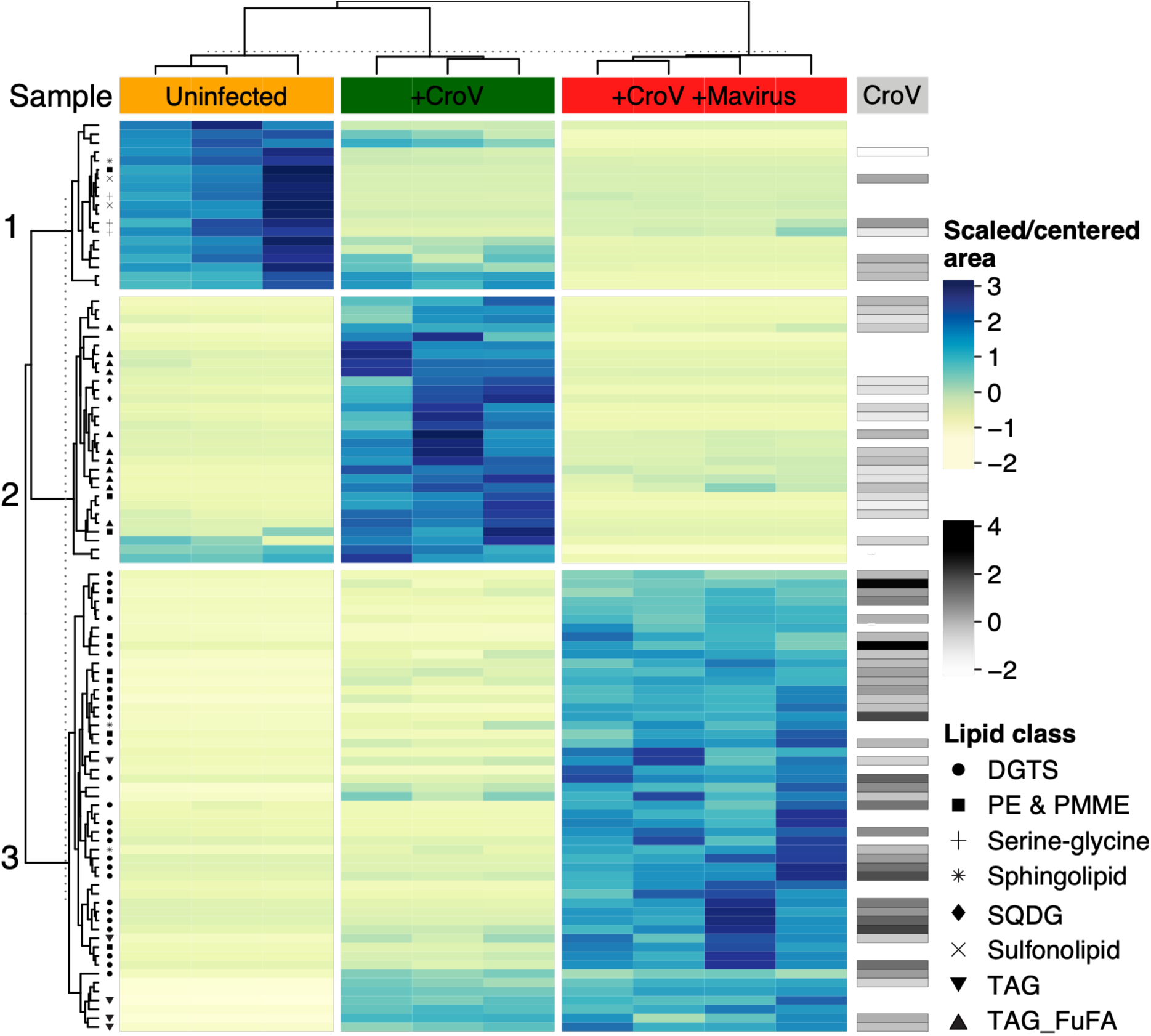
Differential lipid species during infection and co-infection of *C. burkhardae*. Two-dimensional hierarchical clustering of 101 lipid species that show high change in relative abundance (≥100-fold change, Table S1) between uninfected *C. burkhardae* cells (‘Uninfected’, orange, *n* = 3), cells infected with CroV (‘+CroV’, green, *n* = 3), and cells co-infected with CroV and mavirus (‘+CroV +Mavirus’, red, *n* = 4). Clustering was performed on log-transformed and standardized mean peak areas of the adduct ion with the highest peak area. Samples are grouped into three main clusters that separate the different treatments. The lipid species are divided into three clusters (1–3). Lipid classes with two or more annotated species are marked by a symbol. The relative occurrence of the lipids in CroV particles is presented on the right side of the heatmap (gray scale).

In uninfected cells (cluster 1), several phospholipids (phosphatidylcholine (PC) 35:2, phosphatidylethanolamine (PE) 30:0, and phosphatidylinositol (PI) 38:5) and one ceramide (Cer 35:4;O2) were present at higher relative abundance (Fig. S3A and B). Notably, three flavolipins and two sulfonolipids (Rif-1 and Rif-derived ZU-602) were identified in this cluster (Fig. S3C and D). Both flavolipins and sulfonolipids have thus far been described primarily in bacteria of the Flavobacteriaceae family (29–31). Their detection in uninfected *C. burkhardae* cultures could suggest the presence of associated bacteria attached to the cells or uptake through phagotrophic feeding, followed by transient retention within the cells (background signal originating in free-living bacteria was removed prior to the analysis, see Materials and Methods). While we cannot exclude the possibility of an as-yet-unrecognized biosynthetic capacity in *C. burkhardae*, the currently available genomic and biochemical evidence suggests a bacterial origin for these lipid classes.

In cells infected with CroV (cluster 2), two phospholipids (PE 40:8;O and phosphatidylmonomethylethanolamine (PMME) 37:2), one sulfolipid (sulfoquinovosyl diacylglycerol (SQDG) 35:1), one betaine lipid (diacylglyceryl carboxyhydroxymethylcholine (DGCC) 41:4) and one neutral glycerolipid (diacylglycerol (DAG) 40:10) were identified, most of which were not detected in uninfected or co-infected cells (Fig. S4). Notably, more than a third of the identified lipid species were triacylglycerols (TAGs) containing the furan fatty acid (FuFA) 11D5 (Fig. 3A and B, Fig. S5 and Fig. S6, 11 species, of which three contained an additional 9D5 FuFA)(32, 33). These FuFA-containing TAGs were also detected during co-infection, albeit in lower abundance, and most of them were below detection level in uninfected cells (Fig. 3B). Molecular networking analysis using GNPS2 (34) led to the identification of three additional FuFA-containing TAGs and one FuFA-containing DAG, which were clustered together with most of the other FuFA-containing TAGs (7 out of 11 species, Fig. 3C). Similarly, these additional FuFA-containing DAG and TAGs showed higher relative abundance in cells infected with CroV, lower abundance in co-infected cells, and were below detection level in uninfected cells (Fig. 3D). Further targeted search of the characteristic fragment of FuFA 11D5 ([FCO]^+^, mass to charge (*m*/*z*) 333.2794, also characteristic for FuFA 13D3 (33)) using MassQL (35) revealed a group of seven DGTS species containing this FuFA within a larger cluster of DGTS species (Fig. S7A). These FuFA-containing DGTS species were detected in all treatments, with slightly higher relative abundance in cells infected with CroV (Fig. S7B). An additional species containing either both 11D5 and 11D3 or 13D3 and 9D5 FuFAs was identified in cluster 1 (Table S1), having similar relative abundance in uninfected cells and cells infected with CroV while not detected in co-infected cells (Fig. S7B and Fig. S8).

**Figure 3:**
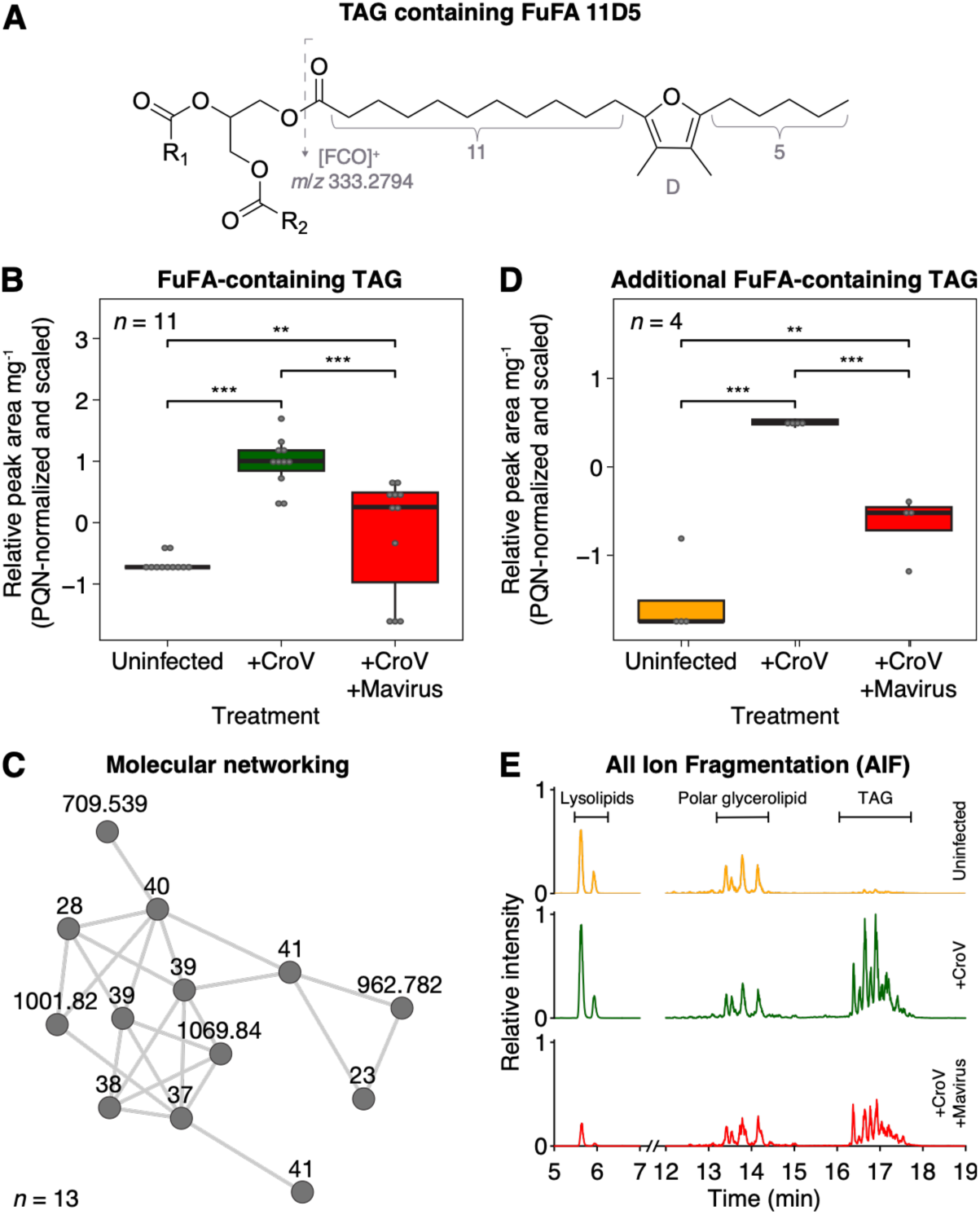
Induction of FuFA-containing TAGs during infection of *C. burkhardae*. (A) Structure of a TAG containing the FuFA 11D5 (11-(3,4-dimethyl-5-pentylfuran-2-yl)-undecanoic acid). R_1_ and R_2_ correspond to the two additional esterified FAs. Numbers 11 and 5 indicate the number of carbon atoms of the carboxyalkyl and the alkyl chain, respectively; ‘D’ indicates a dimethyl-substituted furan moiety (in β,βʹ-positions). [FCO]^+^ represents the characteristic fragment of FuFAs in the MS/MS spectra, with *m*/*z* 333.2794 corresponding to FuFA 11D5 or 13D3 (32, 33). The position of the FuFA in the glycerol backbone does not necessarily reflect the actual order of FAs. (B) Eleven FuFA-containing TAG species were identified through *k-*means clustering and statistical analysis to be highly induced during infection, as well as increased during co-infection (Fig. 2). Relative peak area is presented in uninfected *C. burkhardae* cells (‘Uninfected’, orange), cells infected with CroV (‘+CroV’, green), and cells co-infected with CroV and mavirus (‘+CroV +Mavirus’, red). The indicated *n* denotes the number of lipid species, each presented as the mean relative peak area per mg. Statistical significance was assessed using a linear mixed-effects model, followed by two-sided pairwise comparisons of estimated marginal means with Holm adjustment for multiple comparisons; ** Holm-adjusted *p* < 0.01; *** *p* < 0.001. (C) Three additional FuFA-containing TAG species and one FuFA-containing DAG were identified following network analysis (*m*/*z* values indicated) using a characteristic fragment of FuFA 11D5 or 13D3 ([FCO]^+^, *m*/*z* 333.2794) (33). The network contains seven out of eleven FuFA-containing TAG species identified in the differential analysis (Fig. 2 and Table S1, identification number indicated). (D) Relative peak area of the additional FuFA-containing TAG and DAG species, as in (B). (E) Systemic induction of lysolipids and lipids containing FuFA 11D5 during infection and co-infection. LC-MS extracted ion chromatograms (EICs) of the characteristic fragment of FuFA 11D5 following all ion fragmentation (AIF) are presented for the three treatments, as in (B), *n* = 1. Values were normalized to the sample weight and to the global maximum across the three samples. Comparison of the furan core ion of D5 is presented in Fig. S10.

As FuFA occurred in two different lipid classes, we sought to compare its occurrence across all lipid species by performing an untargeted analysis using the characteristic fragment of FuFA 11D5 (Fig. 3E). The analysis revealed a slight increase in FuFA-containing lysolipids in cells infected with CroV compared to uninfected cells (1.1- to 1.5-fold change) and a lower relative abundance in co-infected cells (0.1- to 0.3-fold change). FuFA-containing polar glycerolipids (including DGTS), on the other hand, had a similar relative abundance in all treatments, while FuFA-containing TAGs were significantly induced both in infected and co-infected cells (36- and 20-fold change respectively), corresponding to the changes observed by comparing the specific lipid species (Fig. 3B and D). Induction was observed also for FuFA 9D5- and 9D3-containing TAGs, albeit to a lesser extent and at lower relative abundance (Fig. S9 and Fig. S10).

In co-infected cells (cluster 3), almost half of the identified lipid species belonged to the class DGTS (24 species, Fig. 4A), with about half also induced in cells infected with CroV, albeit in lower abundances. Interestingly, none of these DGTS species contained FuFA. Additionally, eight phospholipid (five PE, two PMME and one phosphatidylglycerol (PG) species), seven neutral glycerolipids (five TAG and two oxygenated DAG species), two glycosphingolipids, one sulfolipid and one glycine lipid were also present at higher relative abundance in co-infected cells, of which most were also induced to a lesser extent in cells infected with CroV (Fig. 4 and Fig. S11).

**Figure 4:**
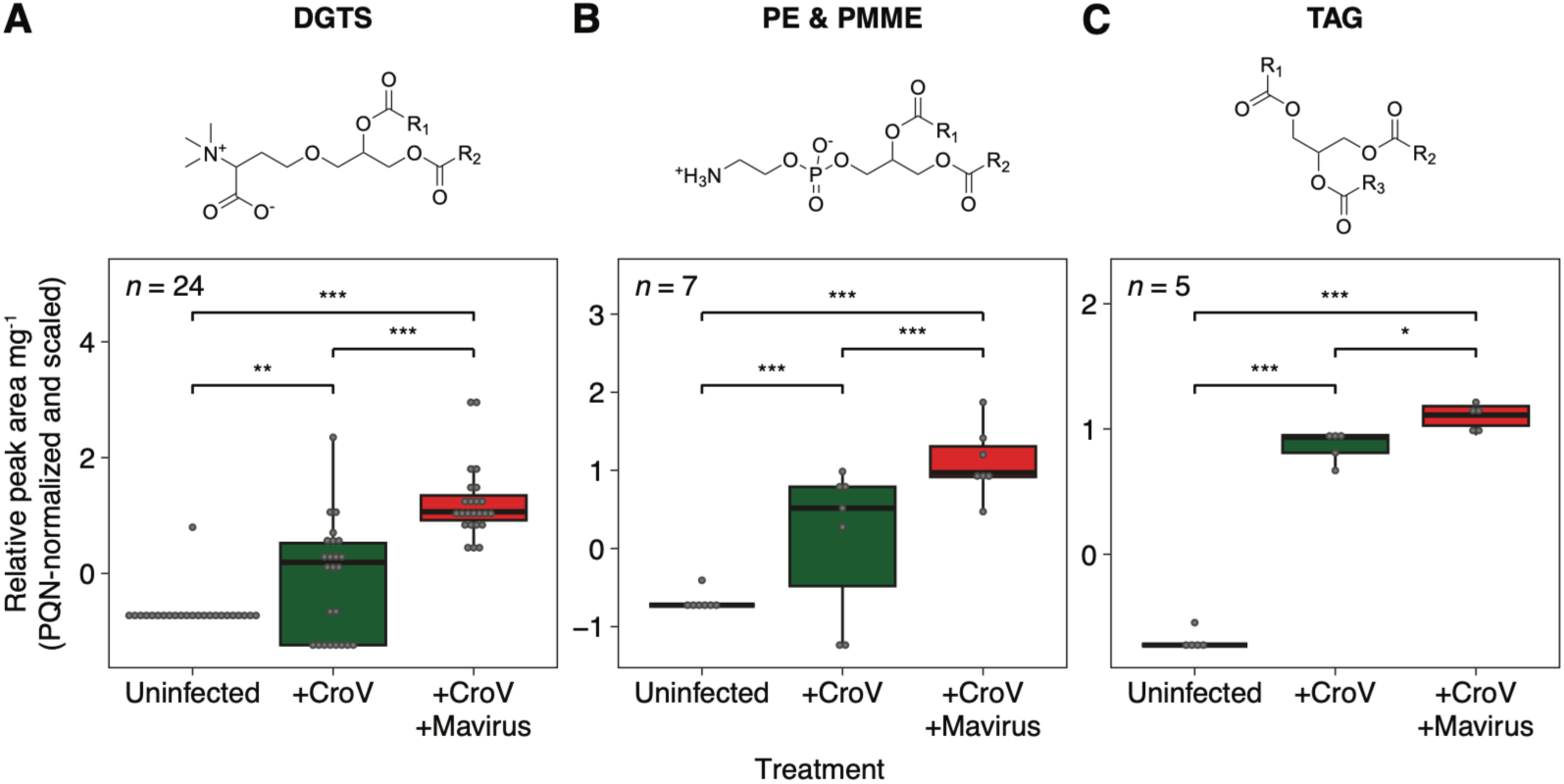
Main lipids species induced during co-infection of *C. burkhardae*. Structure and relative abundance of (A) DGTS, (B) PE & PMME (structure of PE presented), and (C) TAG species in uninfected *C. burkhardae* cells (‘Uninfected’, orange), cells infected with CroV (‘+CroV’, green), and cells co-infected with CroV and mavirus (‘+CroV +Mavirus’, red). The indicated *n* denotes the number of lipid species, each presented as the mean relative peak area per mg. Statistical significance was assessed using a linear mixed-effects model, followed by two-sided pairwise comparisons of estimated marginal means with Holm adjustment for multiple comparisons; * Holm-adjusted *p* < 0.05; ** *p* < 0.01; *** *p* < 0.001. R_1_ and R_2_ and R_3_ correspond to the esterified FAs.

We were further interested in identifying possible genes involved in the biosynthesis of FuFAs and DGTSs both in *C. burkhardae*, CroV and mavirus. Previous studies identified the biosynthetic pathway of FuFA in bacteria (36–38) and soy (39), and a pathway has been suggested for algae (40, 41) (Table S2). We identified 193 genes in the host genome of *C. burkhardae* that are putatively involved in the biosynthesis of sulfonolipids (67), DGTS (38), FuFA (14), PE/PMME (7), and other lipid components (Table S3). While no complete biosynthetic pathway for any of the identified lipids could be reconstructed, four genes showed matches to references from the biosynthetic pathway of both FuFAs and DGTS (KAA0171498.1, KAA0170054.1, KAA0167919.1 and KAA0166380.1; all annotated as methyltransferases, Table S3). Structure predictions of the corresponding protein sequences and subsequent structural homology searches against the PDB confirmed the sequence-based annotation in most cases (Data S1). The majority of the identified genes are involved in ubiquitous processes, such as dehydrogenation or methylation. Interestingly, unlike other systems where genes involved in lipid biosynthesis were identified in the vial genomes (9), no homologous genes were identified in the viral genomes.

Finally, because CroV virions contain an internal lipid membrane (15), we analyzed the lipid composition of purified CroV particles to investigate the occurrence of infection-associated lipids in viral progeny. Most of the lipids that were induced in cells infected with CroV (cluster 2) were also identified in the CroV particles (Fig. 2 and Table S1), including TAG species both with and without FuFAs (11 out of 16 species). Incorporation of infection-derived lipids in viral particles has been observed in diverse systems (19, 23, 24), suggesting that induction of these lipids during infection supports viral stability and infectivity. Notably, CroV particles also contained most of the lipid species that were induced in co-infected cells (cluster 3), including DGTS (21 out of 23 species). Considering that infection with mavirus can hinder production of CroV (11), this might indicate that these lipids, which are induced also during CroV infection, are predominantly incorporated into the membrane of newly produced CroV virions. In co-infected cells, on the other hand, these lipids are not utilized in the CroV membrane and therefore might accumulate at higher concentrations in the co-infected cells (42).

## Discussion

How virophage co-infection modifies the extensive metabolic remodeling commonly induced by giant viruses is largely unknown. Here, we show that CroV infection of the marine heterotrophic protist *C. burkhardae* is associated with a distinctive lipid signature marked by high abundance of FuFA-containing TAGs in both the infected cell and viral progeny. Co-infection with mavirus substantially alters this signature, with a weaker FuFA response and strong enrichment of DGTS species, which are also found in CroV virions. Together with changes in polar glycerolipids, these patterns suggest possible rerouting of acyl flux between membrane lipids and neutral-lipid pools. These findings demonstrate that virophage co-infection substantially reshapes the metabolic consequences of giant virus infection (Fig. 5).

**Figure 5:**
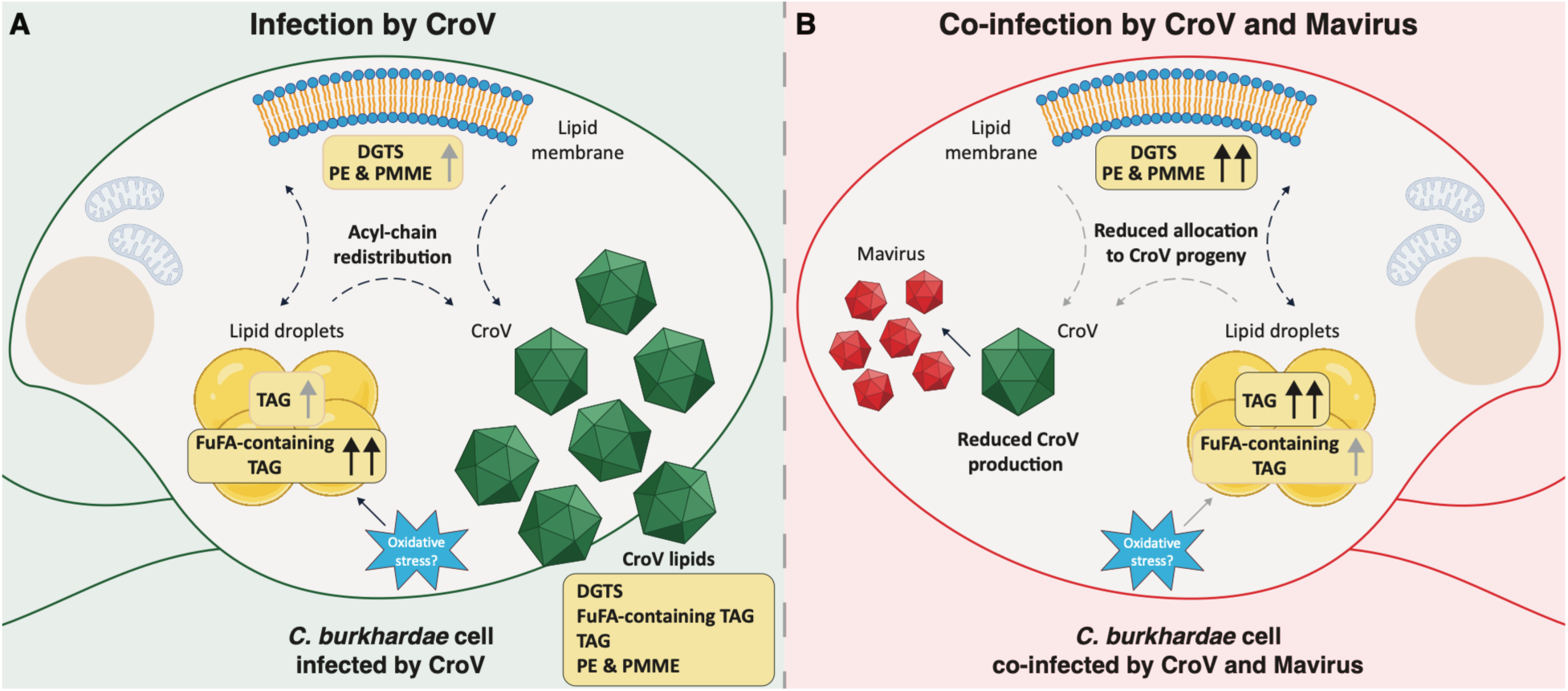
Virophage co-infection reshapes CroV-induced lipid remodeling in *C. burkhardae*. (A) CroV infection is associated with high abundance of FuFA-containing TAGs and increased abundance of DGTS, PE and PMME, together with their presence in CroV particles. The observed lipid changes are consistent with acyl-chain redistribution between membrane lipids, neutral lipid pools and viral progeny. Increased FuFA abundance may be linked to infection-induced oxidative stress. (B) During CroV–mavirus co-infection, CroV production is strongly reduced, the FuFA-containing TAG response is attenuated, and DGTS, PE, PMME and TAGs accumulate intracellularly. Whereas CroV virions contain an internal lipid membrane, mavirus virions lack a lipid membrane. DGTS enrichment during co-infection may therefore partly reflect reduced allocation of lipids to membrane-containing CroV progeny. Dashed arrows indicate proposed lipid redistribution. Several elements of the figure were created with BioRender.com.

### FuFA accumulation is consistent with response to oxidative stress

The preferential accumulation of FuFAs in TAGs during CroV infection (Fig. 3) raises the possibility that these lipids help protect against infection-induced oxidative stress. Such oxidative stress has been documented in other host–giant virus systems (43, 44). FuFAs are potent radical scavengers and can limit lipid peroxidation, thereby contributing to membrane stability and cellular stress tolerance (36, 45–47). Their occurrence in CroV particles raises two non-exclusive possibilities: FuFA moities may protect virion-associated lipids against oxidative damage, whereas the presence of TAGs may reflect a physical association between viral structural components and lipid droplets during assembly, as proposed for another giant virus (24). The weaker abundance of FuFA-containing TAGs in co-infected cells might therefore impact infection outcomes, possibly through antioxidant protection, regulation of redox-sensitive signaling pathways or giant virus assembly.

The biosynthesis of FuFAs remains incompletely understood, with characterized pathways largely restricted to bacteria and emerging evidence from plants and algae (36–41). Our genomic screen identified 14 candidate host genes associated with FuFA biosynthesis, with structural comparisons supporting several annotations. However, most candidates have broadly distributed enzymatic functions, and no complete pathway could be reconstructed. No homologous candidate genes were identified in CroV or mavirus, leaving the enzymatic basis of FuFA production in this system unresolved. The lipidomic data nevertheless indicates that CroV infection promotes accumulation of FuFA predominantly in TAGs, with comparatively small changes in FuFA-containing lysolipids and polar glycerolipids. This pattern could reflect redistribution of existing FuFAs, increased synthesis through an unresolved host pathway, or both.

### DGTS enrichment suggests altered membrane-lipid allocation during co-infection

Whereas the FuFA response was attenuated during co-infection (Fig. 3B, D and E), the accompanying enrichment of DGTS (Fig. 4A) suggests altered allocation of membrane lipids between host cells and viral progeny. One possible explanation is that co-infection changes the balance between DGTS biosynthesis, cellular retention, and incorporation into viral particles. DGTS is a phosphorus-free, zwitterionic membrane lipid associated with stress-induced membrane remodeling, such as phosphate limitation (48–50), and changes in DGTS abundance have also been observed in other host–virus systems (21, 23). Nevertheless, the accompanying increases in PE and PMME (Fig. 4B) suggest broader membrane-lipid remodeling rather than simple replacement of phospholipids by DGTSs. Detection of DGTS in purified CroV particles (Fig. 2 and Table S1) is consistent with recruitment of these lipids during virion formation, as other membrane-containing viruses also acquire selected subsets of host lipids (18, 24, 51). Given that mavirus virions lack a lipid membrane (16), the higher abundance of DGTS in co-infected cells may reflect reduced incorporation into CroV progeny, leaving more DGTS in cellular membranes.

### Lipid accumulation suggests coordinated membrane remodeling and acyl-chain redistribution

The concurrent increases in PE, PMME, and TAG species (Fig. 4B and C) suggest coordinated redistribution of acyl chains between structural phospholipids and neutral-lipid pools during CroV infection and co-infection. PE promotes membrane curvature and fusion (52, 53), and PE-enriched membranes were shown to enhance viral replication complex activity in experimental systems (54, 55), supporting a possible role in generating infection-associated membrane domains. PMME is the first methylated intermediate in the PE-to-PC pathway, and its accumulation is consistent with increased activity or a bottleneck in phospholipid methylation during membrane biogenesis (56). The parallel rise in TAGs could provide temporary storage for excess acyl chains generated during membrane turnover, consistent with the proposed involvement of lipid droplets in viral replication and particle production in other systems (24, 57–59). Dynamic coupling between PE and TAG pools has also been demonstrated in microalgae, with PE serving as a transient carbon sink when neutral-lipid flux is perturbed (60). Together, these patterns support a model in which CroV infection coordinates membrane remodeling with neutral lipid accumulation, while mavirus co-infection alters the balance between cellular lipid retention and allocation to viral progeny.

Overall, our findings show that virophage co-infection redirects the lipid remodeling associated with giant virus infection, shifting a pronounced FuFA-containing TAG signature toward DGTS enrichment (Fig. 5). This highlights the importance of studying changes in the lipidome in multipartite interactions, as the tripartite host–giant virus–virophage system reveals metabolic responses that studies of host–virus pairs alone may not capture. Beyond the infected cell, these changes could alter the biochemical quality of material retained in microbial biomass, transferred to grazers, or released through lysis and viral progeny. Although these ecological consequences remain to be tested, our results identify virophage co-infection as a potential modifier of the composition and fate of organic matter in marine microbial food webs.

## Materials and Methods

### Host and virus strains

The following strains were used in this study: the host *C. burkhardae* strain RCC970-E3, the giant virus CroV strain BV-PW1, and the virophage mavirus strain Spezl.

### Infection experiment and sample collection

*C. burkhardae* cultures (500 mL) were cultivated in f/2 artificial seawater medium in 3 L polycarbonate Fernbach flasks (Corning, NY, USA), as described previously (11). At an initial cell density of 9 × 10^5^ cells per mL, cultures were assigned to one of three treatments: uninfected controls (*n* = 2), infected with CroV (*n* = 4), or co-infected with CroV and mavirus (*n* = 4) (Fig. 1B). CroV was added at a virus-to-host ratio (determined by flow cytometry) of 0.1 and incubated at 22 °C at 50 rpm shaking. For CroV-mavirus co-infections, a suspension of freshly reactivated (from *C. burkhardae* strain E4-10M1 infected with CroV) and 0.22 µm filtered mavirus was added at an approximate virus-to-host ratio of 6, as determined by qPCR (see (11) for details). At 24 hpi, cultures were harvested by centrifugation using a Fiberlite F9 rotor (Thermo Fisher Scientific, MA, USA) for 30 min at 6,774 x g (6,000 rpm) and 20 °C. The resulting cell pellets were resuspended in 8 mL PBS and transferred to clear SW40 ultraclear ultracentrifuge tubes (Beckman Coulter, CA, USA). A layer of 20% (w/v) iodixanol (OptiPrep, Thermo Fisher Scientific, MA, USA), prepared in phosphate-buffered saline (PBS) containing 500 mM NaCl, was carefully placed beneath the cell suspension. Samples were centrifuged in a SW40 rotor (Beckman Coulter, CA, USA) for 20 min at 71,000 x g (20,000 rpm) and 20 °C to separate the host cells from associated bacteria. The resulting visible band of *C. burkhardae* cells was collected by puncturing the side of the centrifuge tube with a syringe and needle and diluted in 14 mL PBS. Cells were washed three times with PBS. Each wash consisted of a centrifugation step of 10 min at 4,000 × g, removal of the supernatant, and resuspension of the pellet in fresh PBS. Following a final centrifugation step, the cells were resuspended in 0.5 mL PBS and lyophilized. Bacterial background controls were collected (*n* = 2) by filtering 500 mL of *C. burkhardae* culture twice through a 3 µm pore-size polycarbonate filter (142 mm diameter, TSTP, Merck Millipore, MA, USA) and checking the filtrate by microscopy for remaining flagellates. The filtrate was then centrifuged in a Fiberlite F9 rotor for 30 min at 6,774 x g (6,000 rpm) and 20 °C to and the bacterial pellet was resuspended in PBS and washed as described above. CroV particles from lysed *C. burkhardae* cultures (*n* = 1) were concentrated by tangential flow filtration using a Vivaflow 200, 0.2 μm PES unit (Thermo Fisher Scientific, MA, USA) to a final volume of 30 mL. The CroV–bacteria concentrate was then purified by density gradient ultracentrifugation as described previously (15) and resuspended in 50 mM Tris-Cl, pH 8.0, 2 mM MgCl_2_. Concentrated cell and virus suspensions were lyophilized using an Alpha 1-2 LDplus freeze-dryer (Christ/Sigma, Osterode, Germany) and stored at –20 °C until further analysis.

### Untargeted lipid profiling using UHPLC-HRMS

Lipid extraction was performed as previously described (61) with slight modifications. Briefly, dry samples were extracted with 1 mL of a pre-cooled (–20 °C) methanol:methyl tert-butyl ether (MTBE) (1:3, *ν*:*ν*) solution containing EquiSPLASH internal standard mixture (100 ng/mL of each species, Avant i Polar Lipids, USA). The samples were shaken for 30 min at 4 °C and sonicated for 30 min in pre-cooled water. The samples were then supplemented with 0.5 mL water:methanol (3:1, *ν*:*ν*) solution, vortexed for 1 min, and centrifuged for 10 min at 16,100 × g and 4 °C. The upper organic phase (0.55 mL) was transferred to 2 mL microcentrifuge tubes and dried under a flow of nitrogen. The polar phase was re-extracted with 0.5 mL of pre-cooled MTBE. The upper organic phase (0.55 mL) was combined with the organic phase from the first extraction and dried under a flow of nitrogen. The samples were stored at –80 °C until LC-MS analysis. Two extraction blanks were collected following the same procedure. One sample of *C. burkhardae* cells infected with CroV was excluded due to low extraction yields.

Data acquisition using ultra-high-performance liquid chromatography-high resolution mass spectrometry (UHPLC-HRMS) was performed as described previously (26), with some modifications. The samples were dissolved in 200 μL acetonitrile:isopropanol (7:3) with 1% 1 M ammonium acetate and 0.1% acetic acid containing internal standards (16:0 PC-d9, Avant i Polar Lipids, USA; and palmitic acid-d4 (C16:0-d4), Santa Cruz Biotechnology, USA), vortexed, sonicated for 10 min and centrifuged for 10 min at 16,100 × g and 10 °C. The supernatants were transferred to 200 μL glass inserts in autosampler vials and directly used for UHPLC-HRMS analysis. A pooled quality control (QC) sample was generated by combining aliquots of 10 μL from all biological samples and was injected at the beginning, middle and end of the run. An aliquot of 2 µL was analyzed using UltiMate 3000 UHPLC (Thermo Fisher Scientific) coupled to a Thermo Fisher Scientific QExactive HF-X Hybrid Quadrupole-Orbitrap equipped with an electrospray ion source. An ACQUITY Premier C8 BEH column (2.1 × 100 mm, 1.7 μm particle; Waters) and pre-column was used for separation with a solvent system of (A) 45% water and 55% acetonitrile:isopropanol (7:3) with 1% (*v*/*v*) 1 M ammonium acetate and 0.1% (*v*/*v*) acetic acid, and (B) acetonitrile:isopropanol (7:3) with 1% (*v*/*v*) 1 M ammonium acetate. The column was maintained at 40 °C and the flow rate of the mobile phase was 0.4 mL per min. The chromatographic gradient was set as follows: 1 min 100% mobile phase A, linear decrease from 100% to 25% mobile phase A over 11 min, from 25% to 0% mobile phase A over 3 min, after which the column was first washed with 100% mobile phase B for 6 min and then returned to initial conditions over 0.5 min and equilibrated for 3.5 min (25 min total run time). Data was acquired in both positive and negative ionization modes (resolution: 60,000 at *m/z* 200) over a mass range *m/z* 150–1,500. One extract of the uninfected culture and one of the bacterial cultures were injected twice to obtain three data files for these treatments.

### Comparative analysis of untargeted lipid profiling data

Before post-processing, the EquiSPLASH, PC-d9 and C16:0-d4 internal standards were assessed to account for *m*/*z* and retention time shifts. UHPLC-HRMS files were converted from the Thermo raw data into .mzML format using msConvert by ProteoWizard (62). Data were processed using MZmine3 (version 3.7.2) for feature extraction and alignment with dataset specific parameters (.xml files with the parameters can be found at [MSV000102757](https://massive.ucsd.edu/ProteoSAFe/dataset.jsp?task=4ec342f6e97e49ebb7386e127d21340c)) (63). Post processing and statistical analysis were performed on the output quantification table from MZmine3 (11,109 mass features, of which 7,192 in position ionization mode and 3,917 in negative ionization mode) using an R script (https://doi.org/10.5281/zenodo.21872557 (64)) based on published workflow (65). Initial PCoA was performed using a Canberra dissimilarity matrix to inspect the data before further background removal and normalization (Fig. S12). For preprocessing, background noise was removed using blank (extraction only) and bacterial background samples with a 30% threshold. Peak areas were further normalized to sample dry weight (Table S4), data was imputed to a low baseline cutoff and then normalized using probabilistic quotient normalization (PQN), scaled and centered. The processed data was used for further statistical analysis: PCoA was performed using a Euclidean dissimilarity matrix to identify statistically significant clusters (Fig. 1C), followed by posthoc PERMANOVA, yielding 8,821 mass features. Differential mass features between uninfected, CroV-infected and co-infected treatments were identified based on *k*-means clustering using the Elbow method (*k* = 3, Fig. S1), followed by one-way repeated measures ANOVA (*p* < 0.01) with false discovery rate (*FDR* ≤ 0.01) correction, yielding 1,172 differential mass features between the clusters, which corresponded to the experimental treatments. The mean peak area of each mass feature was then calculated per cluster, followed by calculation of the fold change between the clusters. A fold change of ≥ 2 (in log scale) between each pair of clusters was selected, yielding 118 differential mass features. Grouping of these mass features into 101 feature groups (Table S1) was performed based on MZmine3 feature deconvolution and manual assignment. These feature groups were then confirmed to have relative peak areas at least two orders-of-magnitude higher than those observed in extraction blanks and bacterial background samples. The peak area of each feature group was then extracted for the CroV samples, after which the resulting data were centered and scaled.

### Putative annotation of lipid species

The putative annotation of the 72 differential lipid species (out of 101 feature groups, Table S1) was performed manually and by using SIRIUS (version 5.6.3)(66) based on accurate mass, MS/MS fragmentation patterns, the Lipid Maps computationally generated database of lipid classes and the Lipid Maps Structure Database (LMSD) (67), and existing literature. Annotation was carried out according to the Metabolomics Standards Initiative, ‘Level 2 – putatively annotated compounds’ (68). MS/MS spectra acquisition was performed using data dependent MS^2^ (ddMS^2^) and targeted MS/MS acquisition modes using UHPLC-HRMS (see above) operated in both ionization modes (resolution: 30,000 at *m/z* 200). ddMS^2^ of the 15 most abundant ions was performed with an isolation window of 1.0 *m/z*, a stepped normalized collision energy (NCE) of 15, 22.5 and 30, and dynamic exclusion of 10 s. Targeted MS/MS analyses were performed with an isolation window of 0.8 *m*/*z* using NCE of 15–35, depending on the lipid species. Identification of FuFA-containing lipids was achieved by comparison to previously reported neutral losses and characteristic fragments: the base peak ion ([FCO]^+^, formally [FuFA-OH]^+^) at *m*/*z* 333.2794 for 11D5 or 13D3 and at *m*/*z* 291.2324 for 9M5 or 13D3, the furan core ion at *m*/*z* 179.1436 for D5 and at *m*/*z* 151.1123 for D3, and the McLafferty rearrangement ion at *m*/*z* 123.0810 (33). Representative MS/MS fragmentation of FuFA-containing TAG and DGTS species can be found in Fig. S5, Fig. S6 and Fig. S8. Identification of glycylserine lipids, glycine lipid and sulfonolipids (RIF-1 and RIF-derived ZU-602) were based on previously published MS/MS fragmentation patterns (29, 31, 69–71). Statistical comparisons at the lipid class level were performed using linear mixed-effects models fitted to log-transformed replicate-level PQN-normalized relative peak areas, with treatment as a fixed effect and lipid species and biological sample as random intercepts. Pairwise treatment contrasts were calculated from estimated marginal means and adjusted within each lipid class using the Holm method. In the corresponding figures, plotted points represent treatment-averaged values for individual lipid species. Statistical comparisons at the individual lipid species level were performed using two-sided Welch’s *t*-tests on PQN-normalized relative peak areas, with pairwise treatment comparisons adjusted within each lipid species using the Holm method.

### Identification of additional FuFA-containing lipid species using molecular networking

Classical molecular networking analysis was performed using GNPS2 (34). Files in .mzml format we processed using the ‘classical_networking_workflow’ (version 2026.03.24) and the parameters specified in Table S5. Targeted search of additional lipid species containing the FuFA 11D5 was performed using the ‘MassQL highlight’ option in the Molecular Networking Browser (35) by querying the characteristic fragment of FuFA 11D5 ([FCO]^+^, *m*/*z* 333.2794, also characteristic for FuFA 13D3 (33)). The molecular network on GNPS2 can be found in the following link: https://gnps2.org/dashboards/networkviewer/?usi=mzdata%3AGNPS2%3ATASK-e61db862ccd443d896be409467fe3874-nf_output%2Fnetworking%2Fnetwork.graphml&usi-mgf=mzdata%3AGNPS2%3ATASK-e61db862ccd443d896be409467fe3874-nf_output%2Fclustering%2Fspecs_ms.mgf#%7B%7D

### Untargeted analysis of FuFA-containing lipids using All Ion Fragmentation (AIF)

AIF was performed using UHPLC-HRMS (see above) operated in both ionization modes (Resolution: 60,000 at *m/z* 200) over a mass range of *m*/*z* 100–1,500 and a stepped NCE of 20, 25 and 30. Extracted ion chromatograms (EICs) were generated for six [FCO]^+^ characteristic fragments (Fig. S9): *m*/*z* 263.2011 (7M5 and 9M3), 277.2168 (7D5 and 9D3), 291.2324 (9M5 and 11M3), 305.2481 (9D5 and 11D3), 319.2637 (11M5 and 13M3), 333.2794 (11D5 and 13D3), and four furan core ions (Fig. S10): 137.0967 (M3), 151.1123 (D3), 165.1280 (M5), 179.1436 (D5) (33). Relative peak area was scaled 0–1 using the maximum peak area value per fragment type ([FCO]^+^ or furan core ion). Fold change was calculated using the peak areas between 5.50–5.80 min and 5.85–6.05 min for lysolipids, 13.20–14.60 min for polar glycerolipids and 16.2–17.8 min for TAGs, normalized to the weight of each sample.

### Bioinformatics analysis of FuFA and DGTS biosynthetic genes

Genomes of the host *C. burkhardae* RCC970, CroV and mavirus were screened for candidate proteins involved in the biosynthesis of the lipid classes detected in this study. Candidate genes were identified from the literature (Table S2) (37, 50, 72–77). Genes were traced to incorporating Pfam families or UniRef Clusters. For Pfam families, pre-build HMM profiles were downloaded, while alignment of UniRef Clusters used MAFFT-E-INS-i v.7.505 (78) and custom HMM profiles for these alignments were built using HMMer v.3.3.2 (79). Both groups of HMM profiles were used to screen all protein sequences from *C. burkhardae*, CroV and Mavirus. The matching protein sequences were grouped into the categories FuFA, DGTS, PE & PMME, TAG, and sulfonolipids based on the matching reference gene clusters/profiles (Table S3). We further included potential lipid biosynthesis-related genes by searching the reference annotations of *C. burkhardae* RCC970 for keywords ‘lipid’, ‘fatty’, ‘triglycer’ and ‘sterol’.

For all candidate proteins involved in lipid biosynthesis, a three-dimensional structure for the monomeric protein was predicted using Alphafold3 v.3.0.1 (80). Each predicted structure was then used as a query to search the PDB database using foldseek v.8.ef4e960 (81) ‘easy-search’. Alignments were visualized with Open-Source PyMOL v.3.1.0 (schrodinger/pymol-open-source 2025 https://github.com/schrodinger/pymol-open-source (22 May 2025, date last accessed)) (Data S1).

### Data, Materials, and Software Availability

All processed data, analysis code, and scripts necessary to reproduce the findings and generate the figures are publicly available in Zenodo (https://doi.org/10.5281/zenodo.21872557)(64). Raw LC-MS files (full MS and ddMS^2^ in .raw and .mzml formats; MS/MS and AIF in .raw format), LC-MS metadata, mzmine3 batch files used for pre-processing and quantification tables following mzmine3 pre-processing are publicly available in MassIVE under the accession number [MSV000102757](https://massive.ucsd.edu/ProteoSAFe/dataset.jsp?task=4ec342f6e97e49ebb7386e127d21340c). The molecular network is available on GNPS2 dashboard: https://gnps2.org/dashboards/networkviewer/?usi=mzdata%3AGNPS2%3ATASK-e61db862ccd443d896be409467fe3874-nf_output%2Fnetworking%2Fnetwork.graphml&usi-mgf=mzdata%3AGNPS2%3ATASK-e61db862ccd443d896be409467fe3874-nf_output%2Fclustering%2Fspecs_ms.mgf#%7B%7D.

## Acknowledgments

This research was supported by an EMBO Postdoctoral Fellowship (grant no. ALTF 192-2021) and an HFSP Postdoctoral Fellowship (grant no. LT0028/2022-L) awarded to G.S.

## Author contributions

G.S. and M.G.F. designed research; G.S. and M.G.F performed experiments; G.S. and M.E.S. analyzed data; G.S. and M.E.S. wrote the manuscript, which was discussed and revised by all co-authors.

