## Supplementary Information for "Viral infection and virophage co-infection induce distinct remodeling of host lipidome"

**This PDF file includes:**

- Figures S1-S12
- Legends for Tables S1-S5
- Legend for Data S1
- SI References

**Other supporting materials for this manuscript include the following:**

- Tables S1-S5 (.xlsx)
- Data S1 (.pdf)

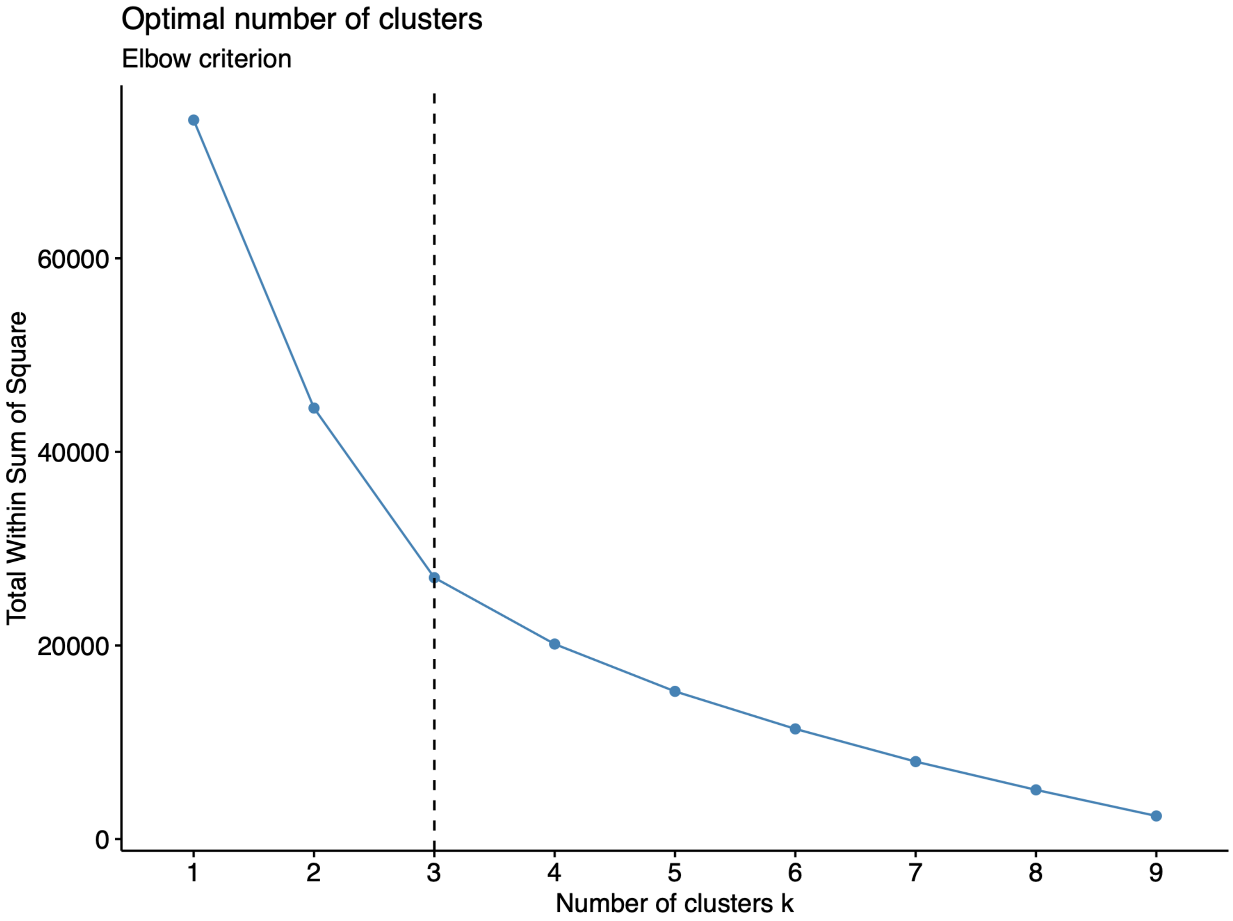


**Figure S1: Elbow method for determining the best number of clusters.** The method was applied on untargeted LC-MS-based lipidomics data using 8,821 mass features derived from cultures of uninfected *C. burkhardae* cells, cells infected with CroV, and cells co-infected with CroV and mavirus. *k* = 3 was chosen as the best number of clusters.

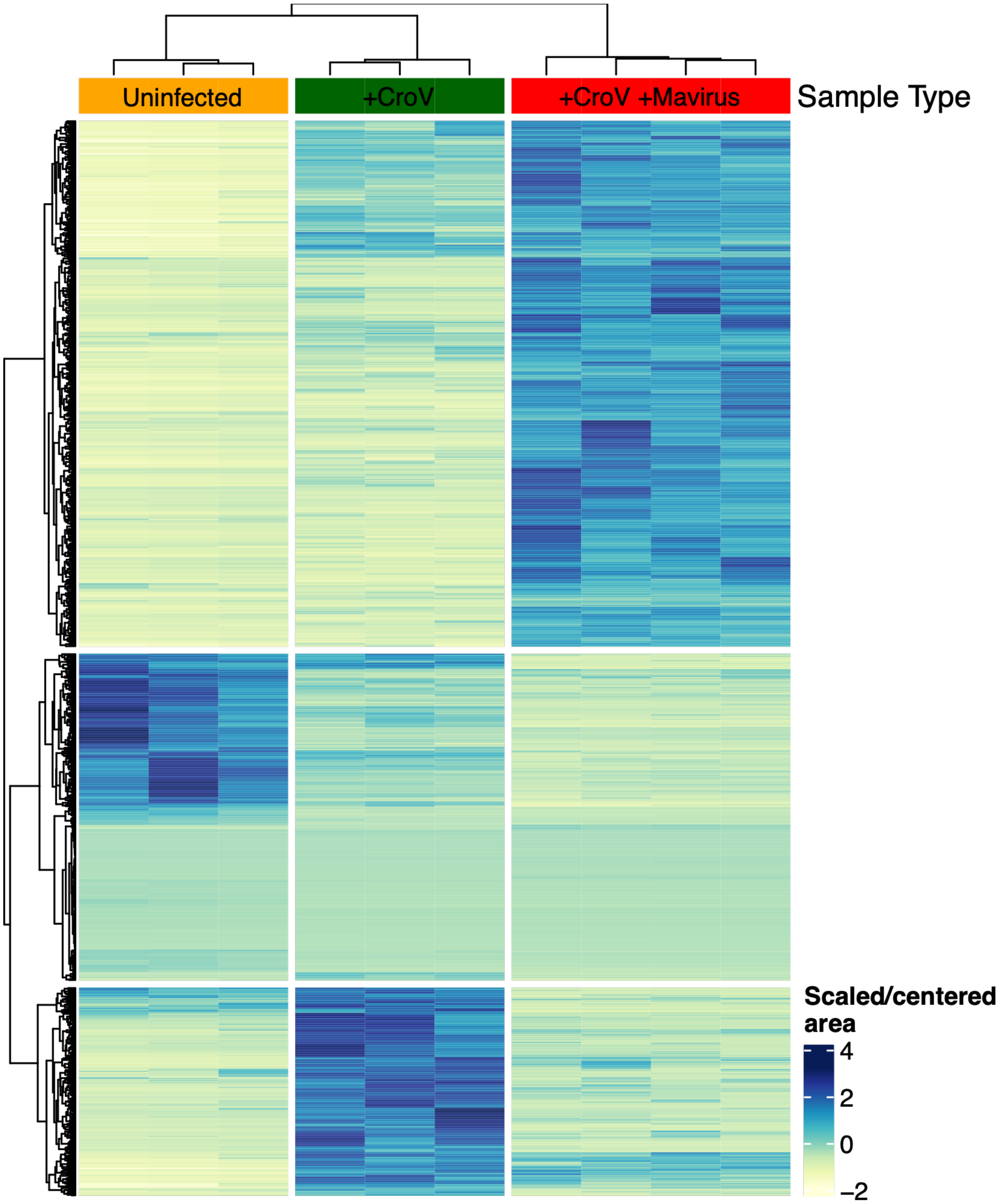


**Figure S2: Differential mass features during infection and co-infection of *C. burkhardae***. Two-dimensional hierarchical clustering of 1,172 statistically significant (one-way repeated measures ANOVA, $p < 0.01$), differential mass features between uninfected *C. burkhardae* cells (‘Uninfected’, $n = 3$), cells infected with CroV (‘+CroV’, $n = 3$), and cells co-infected with CroV and mavirus (‘+CroV +Mavirus’, $n = 4$). Clustering was performed on log-transformed and standardized mean peak areas.

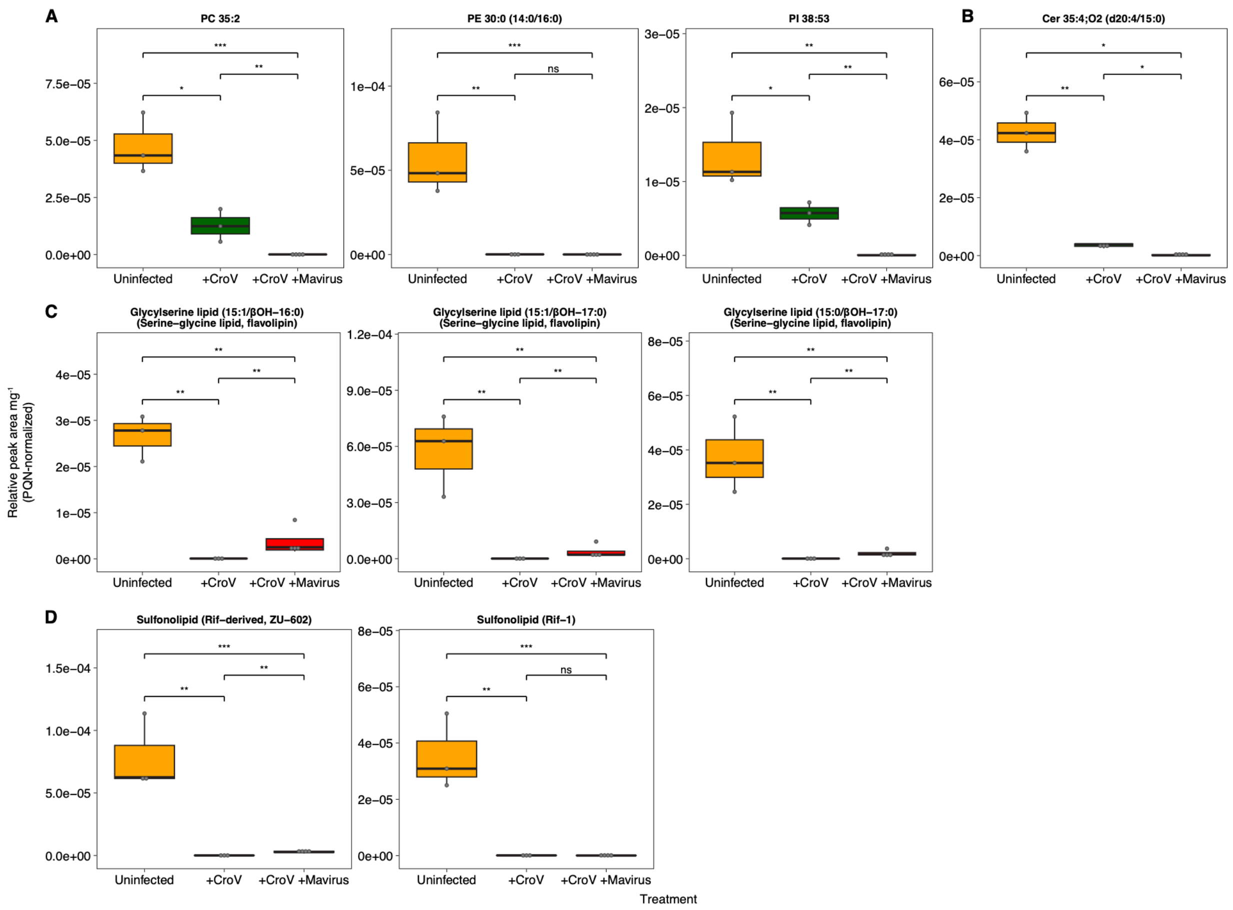


**Figure S3: Main differential lipid species in cluster 1**. Relative peak area of (A) phospholipids, (B) a sphingolipid, (C) glycylserine lipids (flavolipins), and (D) sulfonolipids in uninfected *C. burkhardae* cells ('Uninfected', $n$ = 3, orange), cells infected with CroV ('+CroV', $n$ = 3, green), and cells co-infected with CroV and mavirus ('+CroV +Mavirus', $n$ = 4, red). Statistical comparisons were performed using two-sided Welch's $t$-tests on log-transformed replicate-level PQN-normalized relative peak areas. Pairwise treatment comparisons were adjusted within each lipid species using the Holm method. * Holm-adjusted $p < 0.05$; ** $p < 0.01$; *** $p < 0.001$; ns, not significant.

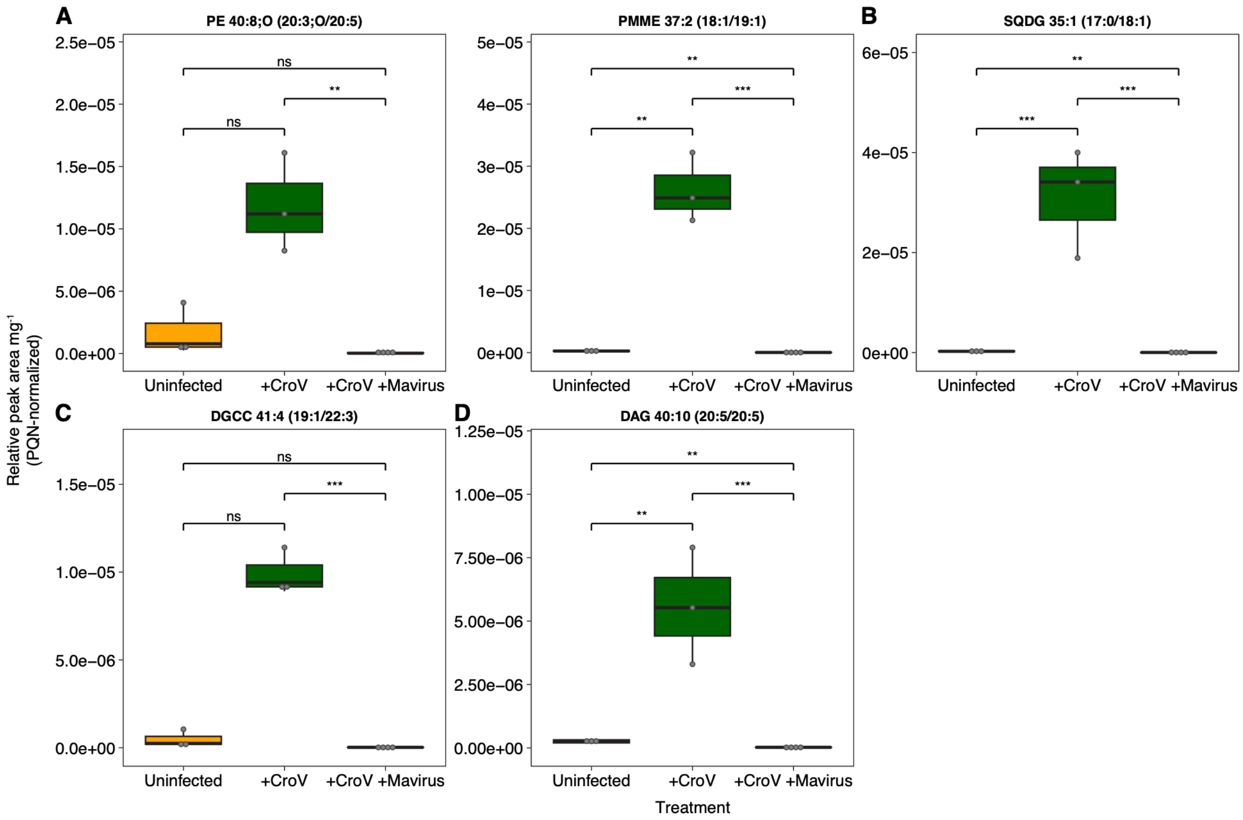


**Figure S4: Additional differential lipid species in cluster 2**. Relative peak area of (A) phospholipids, (B) a sulfolipid, (C) a betaine lipid, and (D) a neutral glycerolipid in uninfected *C. burkhardae* cells ('Uninfected', $n = 3$, orange), cells infected with CroV ('+CroV', $n = 3$, green), and cells co-infected with CroV and mavirus ('+CroV +Mavirus', $n = 4$, red). Statistical comparisons were performed using two-sided Welch's $t$-tests on log-transformed replicate-level PQN-normalized relative peak areas. Pairwise treatment comparisons were adjusted within each lipid species using the Holm method. ** Holm-adjusted $p < 0.01$; *** $p < 0.001$; ns, not significant.

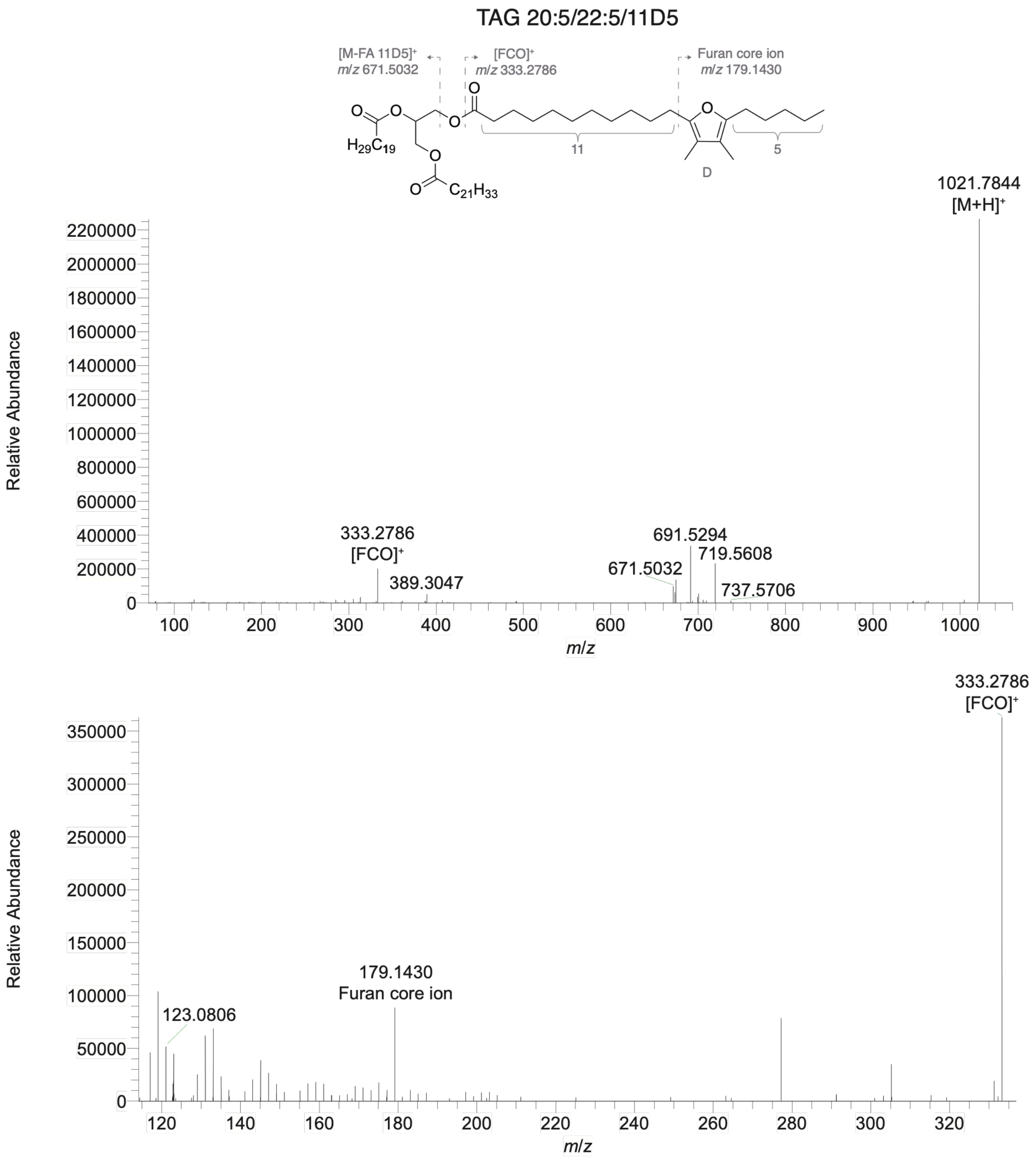


**Figure S5: Identification of the FuFA-TAG 20:5/22:5/11D5**. A putative structure is presented together with the corresponding MS/MS over the full mass range (top) and an expanded view of the region with characteristic fragments for furan fatty acids (FuFAs, bottom). Fragments were detected in positive ionization MS/MS mode using $[M+H]^+$ = 1021.7860 as the precursor ion. Numbers 11 and 5 in the structure indicate the number of carbon atoms of the carboxyalkyl and alkyl chains, respectively; 'D' indicates a dimethyl-substituted furan moiety (in β,β'-positions). $[FCO]^+$ represents the characteristic fragment of FuFAs in the MS/MS spectra, with *m/z* 333.2794 corresponding to FuFA 11D5 or 13D3 (1, 2). The FAs are listed by increasing molacular mass followed by the FuFA in position three of the glycerol backbone, which does not necessarily reflect the actual order. Identificatoin was performed according to the Metabolomics Standards Initiative level 2 annotation (3).

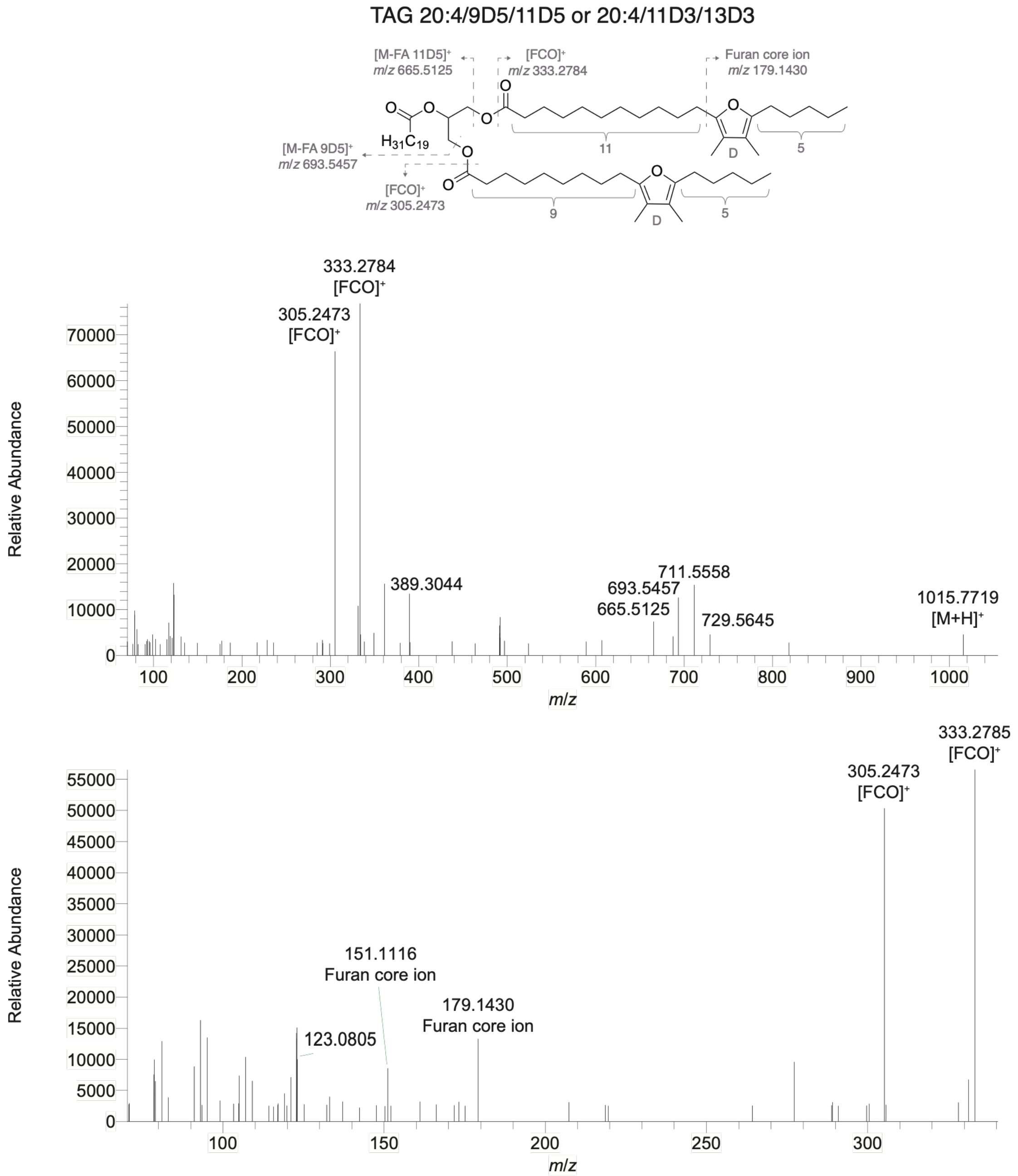


**Figure S6: Identification of the FuFA-TAG 20:4/9D5/11D5 or 20:4/11D3/13D3**. A putative structure is presented for TAG 20:4/9D5/11D5 together with the corresponding MS/MS over the full mass range (top, NCE 25) and an expanded view of the region with characteristic fragments for FuFAs (bottom, NCE 35). Fragments were detected in positive ionization MS/MS mode using $[M+H]^+$ = 1015.7719 as the precursor ion. Numbers 9 or 11 and 5 in the structure indicate the number of carbon atoms of the carboxyalkyl and alkyl chains, respectively; 'D' indicates a dimethyl-substituted furan moiety (in β,β'-positions). $[FCO]^+$ represents the characteristic fragment of FuFAs in the MS/MS spectra, with *m/z* 333.2784 corresponding to FuFA 11D5 or 13D3 and *m/z* 305.2473 corresponding to FuFA 9D5 or 13D3 (1, 2). The FAs are listed by increasing molacular mass, which does not necessarily reflect the actual order. Identificatoin was performed according to the Metabolomics Standard/s Initiative level 2 annotation (3).

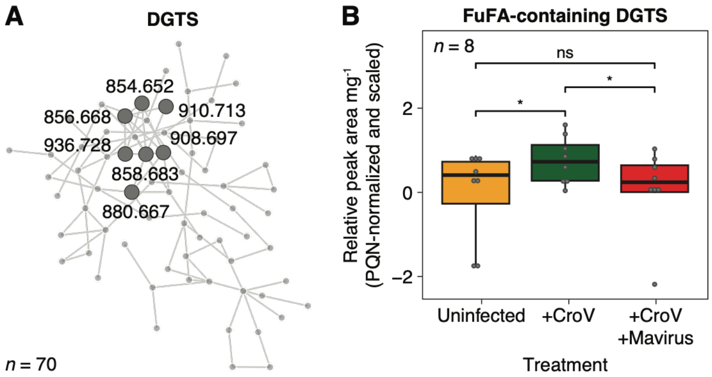


**Figure S7: Identification of FuFA-containing DGTS species**. (A) Seven additional FuFA-containing DGTS species were identified following network analysis (*m*/*z* values indicated) following a MassQL search of the characteristic fragment of FuFA 11D5 ($[FCO]^+$, *m*/*z* 333.2794, also characteristic for FuFA 13D3) (2). The network contained 63 additional DGTS species without a FuFA. (B) Relative peak area of eight FuFA-containing DGTS species (seven from the network analysis and one from cluster 1) in uninfected *C. burkhardae* cells (‘Uninfected’, orange), cells infected with CroV (‘+CroV’, green), and cells co-infected cells with CroV and mavirus (‘+CroV +Mavirus’, red). The indicated *n* denotes the number of lipid species, each presented as the mean relative peak area per mg. Statistical significance was assessed using a linear mixed-effects model, followed by two-sided pairwise comparisons of estimated marginal means with Holm adjustment for multiple comparisons; * Holm-adjusted $p < 0.05$; ns, not significant.

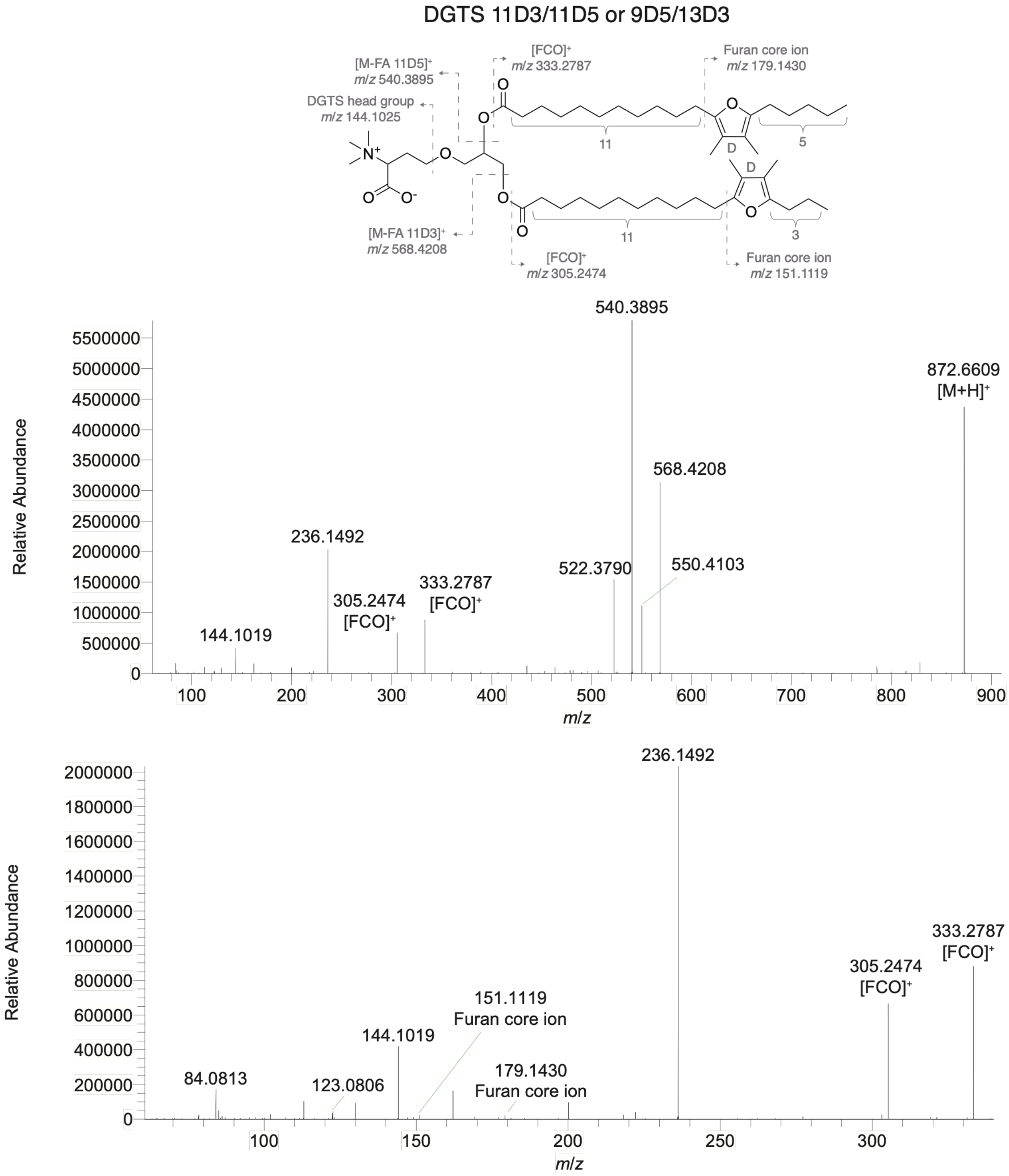


**Figure S8: Identification of the FuFA-DGTS 11D3/11D5 or 9D5/13D3**. A putative structure is presented for DGTS 11D3/13D5 together with the corresponding MS/MS over the full mass range (top) and an expanded view of the region with characteristic fragments for FuFAs and DGTS head group (bottom). Fragments were detected in positive ionization MS/MS mode using $[M+H]^+$ = 872.6609 as the precursor ion. Numbers 11 and 3 or 5 in the structure indicate the number of carbon atoms of the carboxyalkyl and alkyl chains, respectively; 'D' indicates a dimethyl-substituted furan moiety (in β,β'-positions). $[FCO]^+$ represents the characteristic fragment of FuFAs in the MS/MS spectra, with *m/z* 333.2784 corresponding to FuFA 11D5 or 13D3 and *m/z* 305.2473 corresponding to FuFA 9D5 or 13D3 (1, 2). The FuFAs are listed by increasing molacular mass, which does not necessarily reflect the actual order. Identificatoin was prformed according to the Metabolomics Standards Initiative level 2 annotation (3).

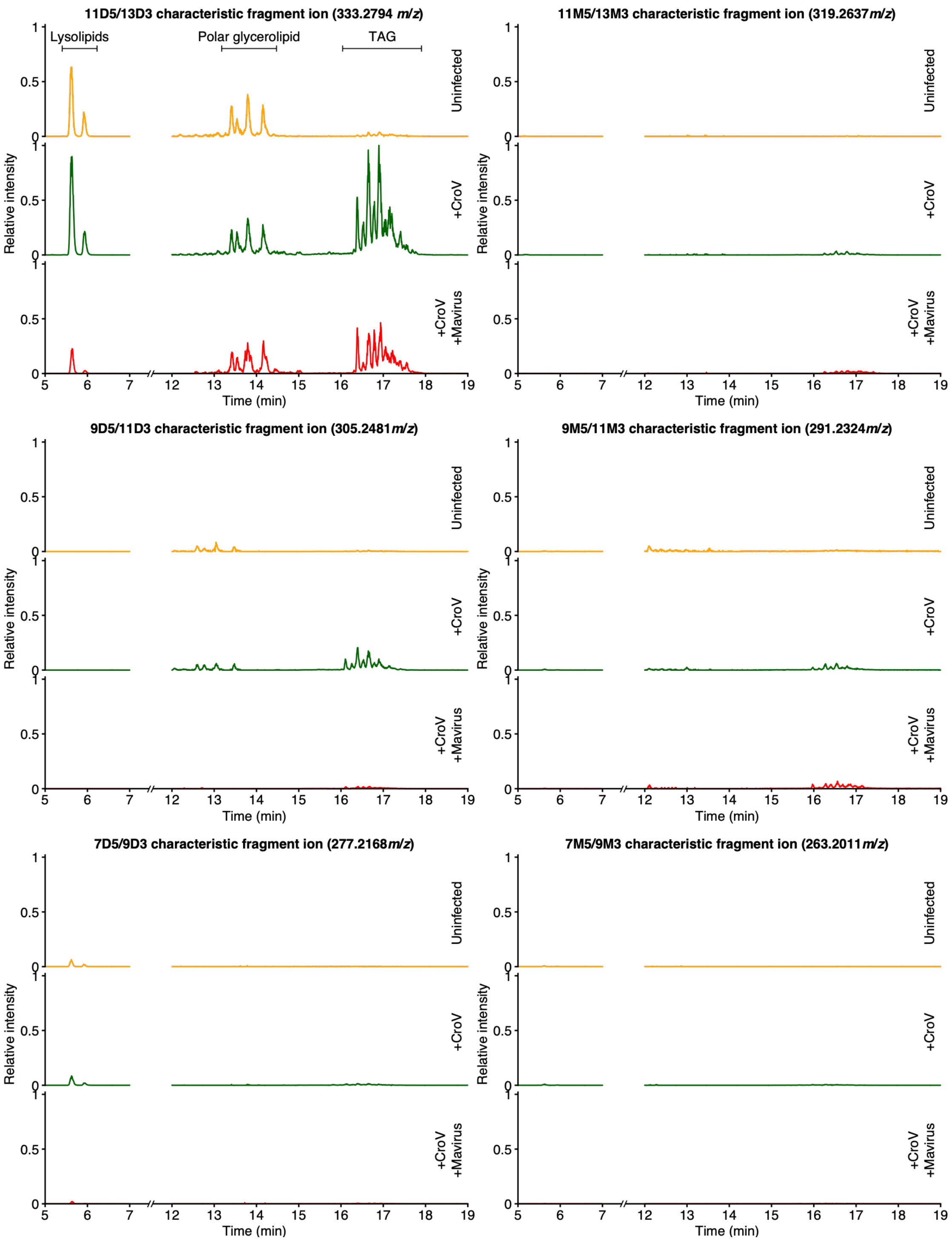


**Figure S9: Relative abundance of various FuFA-containing lipids based on the characteristic fragment ions.** LC-MS extracted ion chromatograms (EICs) of the characteristic fragment ions ($[FCO]^+$, *m/z* values indicated (2)) following all ion fragmentation (AIF) are presented for uninfected *C. burkhardae* cells ('Uninfected', orange), cells infected with CroV ('+CroV', green), and cells co-infected cells with CroV and mavirus ('+CroV +Mavirus', red), *n* = 1. Values were normalized to the sample weight and to the global maximum across the three samples and all characteristic fragment ions.

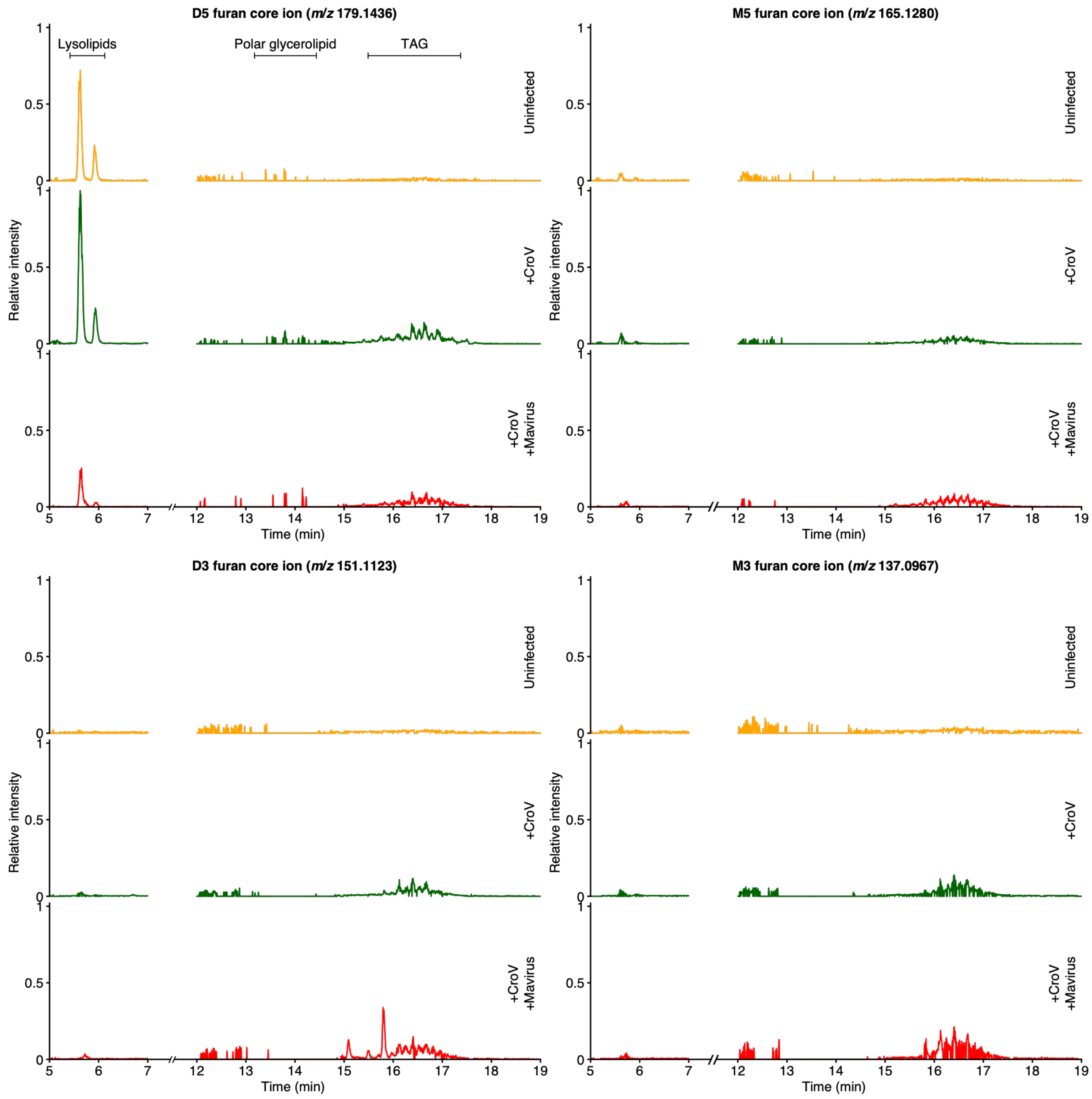


**Figure S10: Relative abundance of various FuFA-containing lipids based on the furan core ions.** LC-MS extracted ion chromatograms (EICs) of the furan core ions (*m/z* values indicated (2)) following all ion fragmentation (AIF) are presented for uninfected *C. burkhardae* cells ('Uninfected', orange), cells infected with CroV ('+CroV', green), and cells co-infected with CroV and mavirus ('+CroV +Mavirus', red), *n* = 1. Values were normalized to the sample weight and to the global maximum across the three samples and all furan core ions.

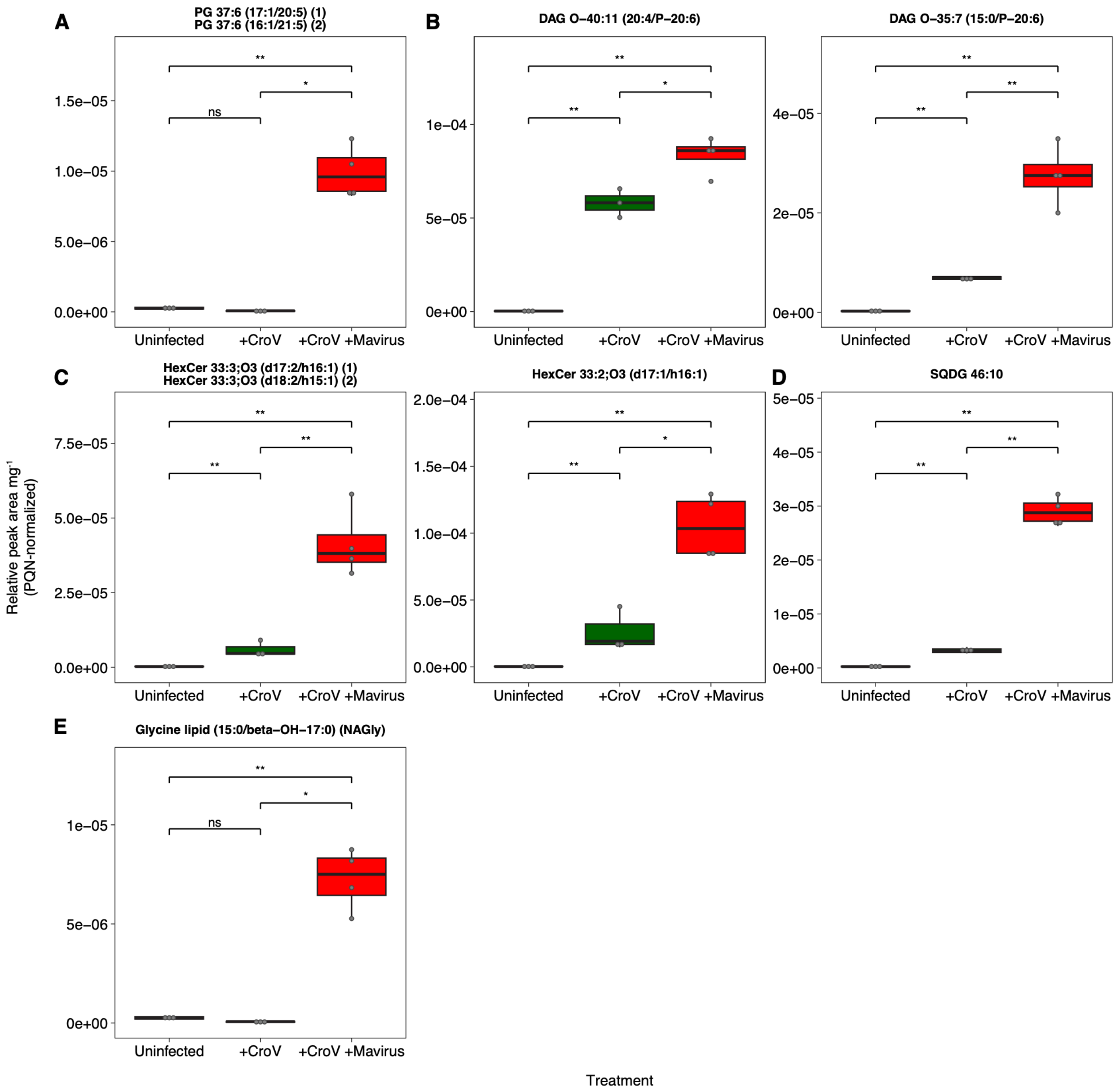


**Figure S11: Additional differential lipid species in cluster 3**. Relative peak area of (A) a phospholipid, (B) neutral glycerolipids, (C) sphingolipids, (D) a sulfolipid, and (E) a glycine lipid in uninfected *C. burkhardae* cells ('Uninfected', *n* = 3, orange), cells infected with CroV ('+CroV', *n* = 3, green), and cells co-infected with CroV and mavirus ('+CroV +Mavirus', *n* = 4, red). Statistical comparisons were performed using two-sided Welch's *t*-tests on log-transformed replicate-level PQN-normalized relative peak areas. Pairwise treatment comparisons were adjusted within each lipid species using the Holm method. * Holm-adjusted $p < 0.05$; ** $p < 0.01$; *** $p < 0.001$; ns, not significant.

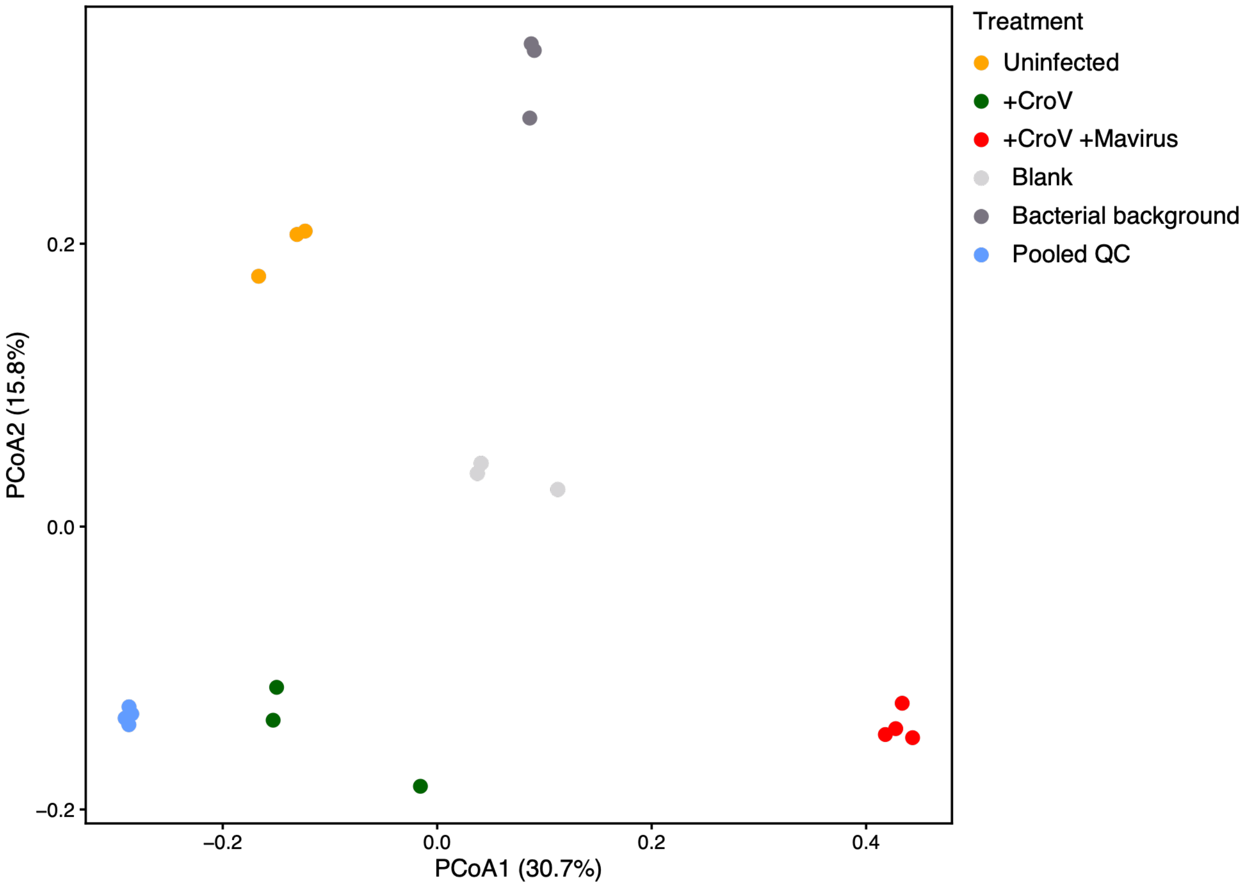


**Figure S12: Principal Coordinates Analysis (PCoA) of all samples before background removal and normalization.** Samples: uninfected *C. burkhardae* cells (‘Uninfected’, orange, *n* = 3), cells infected with CroV (‘+CroV’, green, *n* = 3), and cells co-infected with CroV and mavirus (‘+CroV +Mavirus’, red, *n* = 4) , extraction blank (‘Blank’, light grey, *n* = 3), bacterial background (‘Bacterial background’, dark grey, *n* = 3), and pooled QC (‘Pooled QC’, light blue, *n* = 4) based on untargeted lipidomics (11,109 mass features). Percentage of explained variance is indicated per coordinate.

**Table S1: Differential lipid species during infection and co-infection of *C. burkhardae*.** The table contains information about the 101 putative lipid species that were differential between uninfected cells, cells infected with CroV and cells co-infected with CroV and mavirus (fold change ≥100 between at least two treatments). Heatmap # represents the order of the features in the heatmap (Fig. 2), from top to bottom. Putative annotation was performed manually and by using SIRIUS (version 5.6.3)(4) based on accurate mass, MS/MS fragmentation patterns, the Lipid Maps computationally generated database of lipid classes and the Lipid Maps Structure Database (LMSD) (5), and existing literature. Annotation was carried out according to the Metabolomics Standards Initiative, 'Level 2 – putatively annotated compounds' (3).

**Table S2: Characterized genes and enzymes in the biosynthetic pathways of FuFA, DGTS, PE, PMME and sulfonolipids.** The table contains the accession numbers, function and references.

**Table S3: *C. burkhardae* genes putatively involved in the biosynthesis of FuFA, DGTS, PE/PMME, sulfonolipids and other lipid components.** The table contains the gene locus, accession numbers, annotation, the lipid class.

**Table S4: Dry weight of biological samples used for normalization.** The table contains the sample type, replicate number and dry weight in mg.

**Table S5: Parameters used for molecular networking analysis using GNPS2.** The table contains the parameters, as summarized by GNPS2 following analysis.

**Data S1:** Structure predictions of the corresponding protein sequences and subsequent structural homology searches against the PDB confirmed the sequence-based annotation in most cases.
