## Supplementary material for "Viral infection and virophage co-infection induce distinct remodeling of host lipidome": Data S1

### kaa0145772

- Sequence-based annotation for kaa0145772 (predicted in DGTS from hits to PF11899, PF13649, UniRef50__A0A165VFW6, UniRef50__A0A238JF33, UniRef50__A0A699Z892, UniRef50__A0A9P7X1M9) is hypothetical protein FNF28_07802 [Cafeteria roenbergensis]
- Best hit was 5wp5 chain B: Phosphomethylethanolamine N-methyltransferase 2

| target | prob | fident | alnlen | evalue | theader |
|---|---|---|---|---|---|
| 5wp5-assembly2.cif.gz__B | 1 | 0.127 | 456 | 3.069e-12 | Arabidopsis thaliana phosphoethanolamine N-methyltransferase 2 (AtPMT2) in complex with SAH |
| 5wp4-assembly1.cif.gz__A | 1 | 0.118 | 465 | 3.135e-11 | Arabidopsis thaliana phosphoethanolamine N-methyltransferase 1 (AtPMT1, XIOPTL) in complex with SAH and phosphocholine |
| 4ine-assembly1.cif.gz__B | 1 | 0.129 | 454 | 3.135e-11 | Crystal structure of N-methyl transferase (PMT-2) from Caenorhabditis elegant complexed with S-adenosyl homocysteine and phosphoethanolamine |

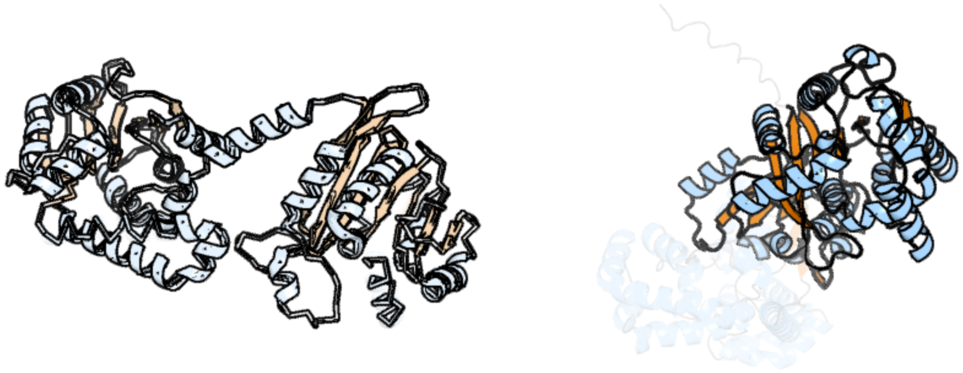

Figure 1: left: reference structure of 5wp5 chain B. right: predicted structure of kaa0145772, unaligned sequences are shown as transparent

### kaa0145785

- Sequence-based annotation for kaa0145785 (predicted in PE_PMME from hits to PF01066) is hypothetical protein FNF28_07795 [Cafeteria roenbergensis]
- No significant structural hit found

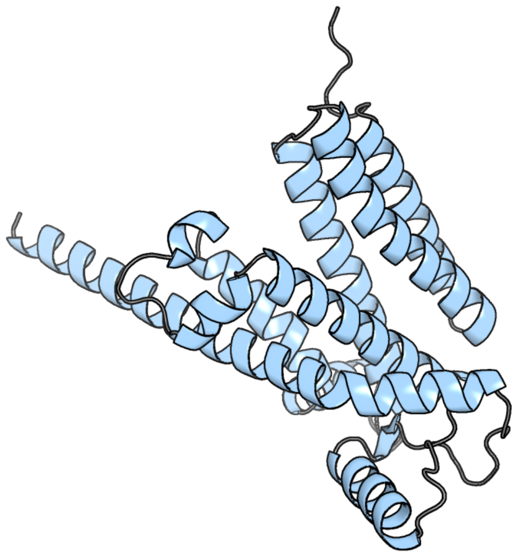

Figure 2: predicted structure of kaa0145785

### kaa0146217

- Sequence-based annotation for kaa0146217 (predicted in Fu-FA from hits to PF01593) is hypothetical protein FNF28_07704 [Cafeteria roenbergensis]
- Best hit was 6wpu chain A: Flavin-containing monooxygenase

| target | prob | fident | alnlen | evalue | theader |
|---|---|---|---|---|---|
| 6wpu-assembly1.cif.gz_A | 1 | 0.249 | 562 | 1.215e-33 | Structure of S-allyl-L-cysteine S-oxygenase from Allium sativum |
| 5nmw-assembly2.cif.gz_D | 1 | 0.234 | 528 | 9.339e-33 | Crystal Structure of the pyrrolizidine alkaloid N-oxygenase from Zonocerus variegatus in complex with FAD |
| 5nmw-assembly1.cif.gz_A-2 | 1 | 0.224 | 535 | 1.742e-32 | Crystal Structure of the pyrrolizidine alkaloid N-oxygenase from Zonocerus variegatus in complex with FAD |

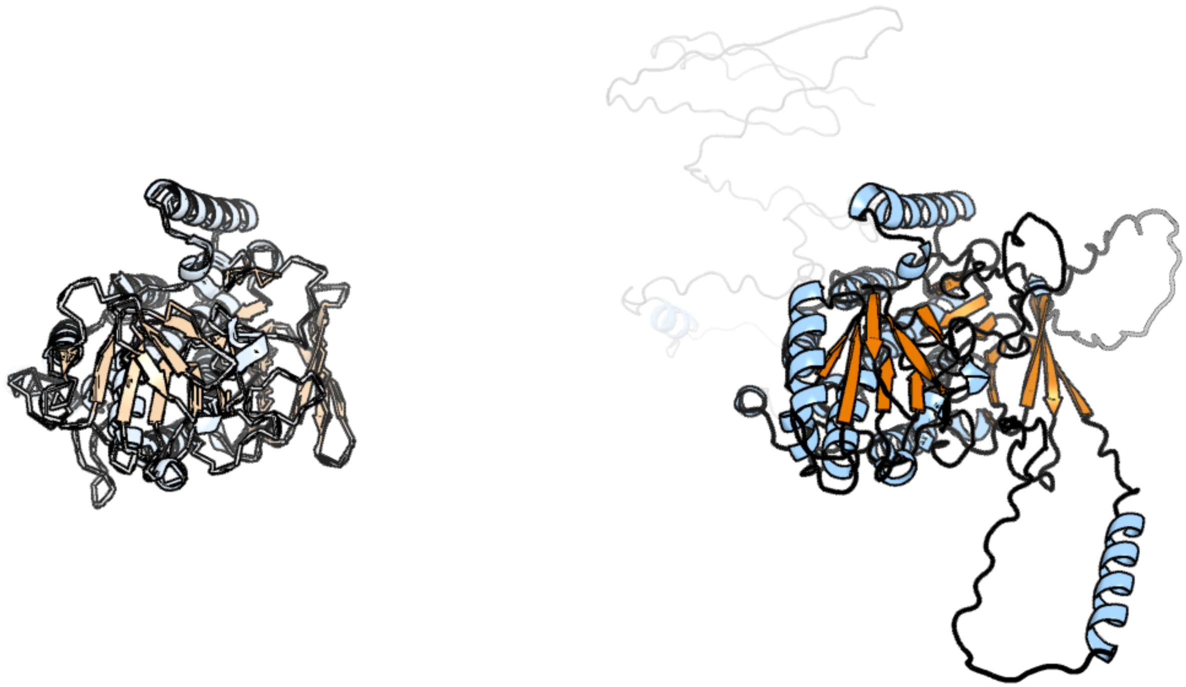

Figure 3: left: reference structure of 6wpu chain A. right: predicted structure of kaa0146217, unaligned sequences are shown as transparent

#### kaa0146863

- Sequence-based annotation for kaa0146863 (predicted in Sulfonolipids from hits to PF00155) is hypothetical protein FNF28_07625 [Cafeteria roenbergensis]
- Best hit was 6t8q chain A: Kynurenine/alpha-aminoadipate aminotransferase, mitochondrial

| target | prob | fident | alnlen | evalue | theader |
|---|---|---|---|---|---|
| 6t8q-assembly1.cif.gz_A | 1 | 0.387 | 462 | 6.466e-43 | HKATII IN COMPLEX WITH LIGAND (2R)-N-benzyl-1-[6-methyl-5-(oxan-4-yl)-7-oxo-6H,7H-[1,3]thiazolo[5,4-d]pyrimidin-2-yl]pyrrolidine-2-carboxamide |
| 4ge4-assembly1.cif.gz_A | 1 | 0.386 | 461 | 6.331e-42 | Kynurenine Aminotransferase II Inhibitors |
| 6d0a-assembly1.cif.gz_A | 1 | 0.387 | 462 | 7.04e-42 | Crystal structure of Kynurenine Aminotransferase-II in apo-form, at 1.47 A resolution |

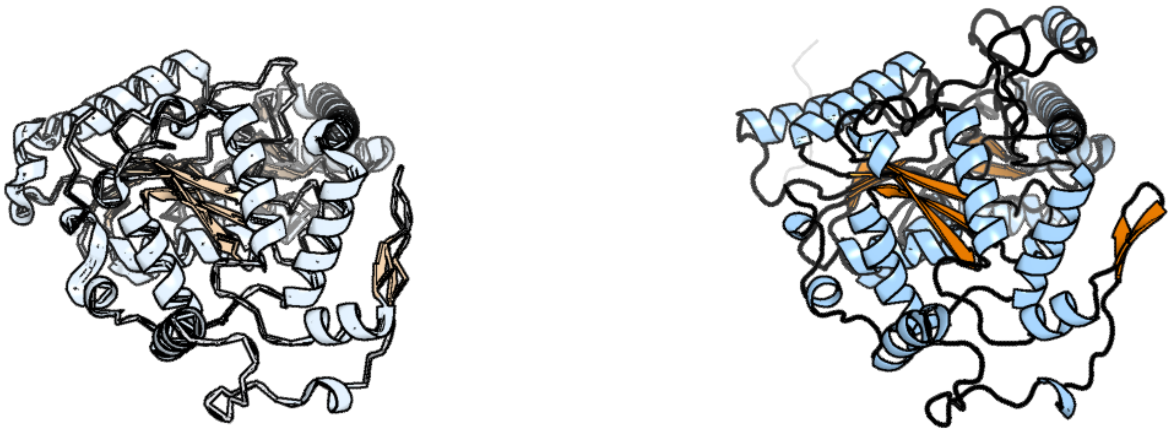

Figure 4: left: reference structure of 6t8q chain A. right: predicted structure of kaa0146863, unaligned sequences are shown as transparent

#### kaa0147976

- Sequence-based annotation for kaa0147976 (predicted in Sulfonolipids from hits to PF00106) is hypothetical protein FNF28_07503 [Cafeteria roenbergensis]
- Best hit was 2gdz chain A: NAD+-dependent 15-hydroxyprostaglandin dehydrogenase

| target | prob | fident | alnlen | evalue | theader |
|---|---|---|---|---|---|
| 2gdz-assembly1.cif.gz_A-2 | 1 | 0.248 | 282 | 2.653e-18 | Crystal structure of 15-hydroxyprostaglandin dehydrogenase type1, complexed with NAD+ |
| 8cwl-assembly1.cif.gz_B | 1 | 0.239 | 284 | 1.363e-16 | Cryo-EM structure of Human 15-PGDH in complex with small molecule SW222746 |
| 3wye-assembly1.cif.gz_A-2 | 1 | 0.251 | 278 | 9.487e-16 | Crystal Structure of chimeric engineered (2S,3S)-butanediol dehydrogenase complexed with NAD+ |

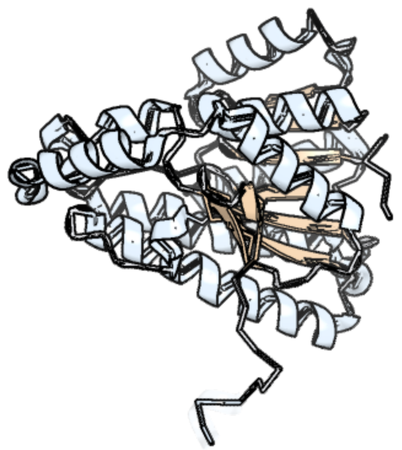

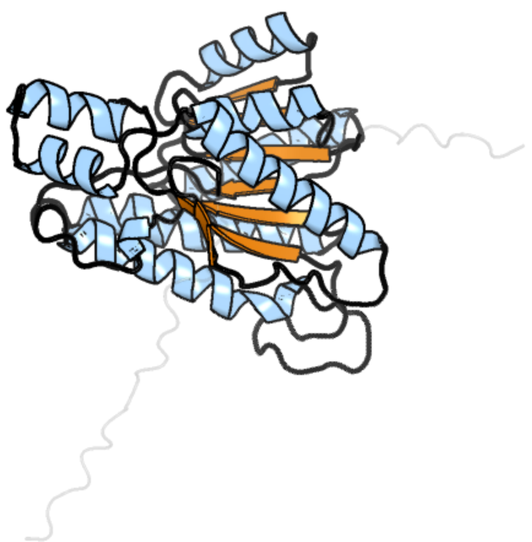

Figure 5: left: reference structure of 2gdz chain A. right: predicted structure of kaa0147976, unaligned sequences are shown as transparent

#### kaa0148283

- Sequence-based annotation for kaa0148283 (predicted in Sulfonolipids from hits to PF00291, UniRef50_A0A2D9B818) is hypothetical protein FNF28_07465 [Cafeteria roenbergensis]
- Best hit was 5jjc chain B: Cysteine synthase

| target | prob | fident | alnlen | evalue | theader |
|---|---|---|---|---|---|
| 5jjc-assembly2.cif.gz_B | 1 | 0.412 | 327 | 3.541e-40 | Crystal Structure of double mutant (Q96A-Y125A) O-Acetyl Serine Sulfhydralase from Brucella abortus |
| 5i7w-assembly2.cif.gz_B | 1 | 0.414 | 335 | 1.538e-39 | Crystal Structure of a Cysteine Synthase from Brucella suis |
| 5jis-assembly1.cif.gz_A | 1 | 0.422 | 327 | 2.32e-39 | The Crystal Structure of O-acetyl serine sulfhydralase from Brucella abortus |

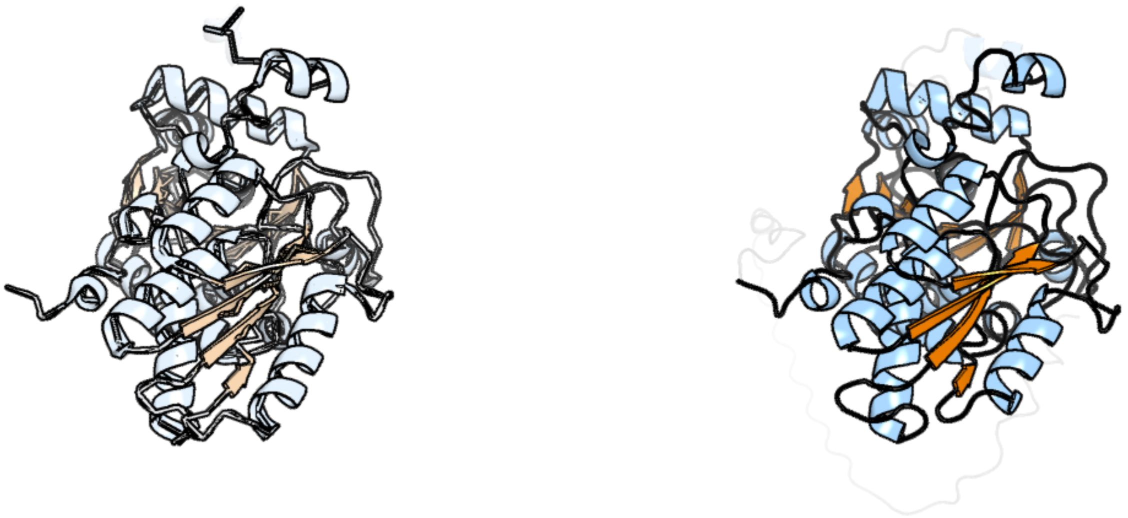

Figure 6: left: reference structure of 5jjc chain B. right: predicted structure of kaa0148283, unaligned sequences are shown as transparent

### kaa0148314

- Sequence-based annotation for kaa0148314 (predicted in DGTS from hits to PF13649) is hypothetical protein FNF28_07464 [Cafeteria roenbergensis]
- Best hit was 3lcc chain A: Putative methyl chloride transferase

| target | prob | fident | alnlen | evalue | theader |
|---|---|---|---|---|---|
| 3lcc-assembly1.cif.gz_A | 1 | 0.294 | 224 | 4.824e-16 | Structure of a SAM-dependent halide methyltransferase from Arabidopsis thaliana |
| 8ajp-assembly1.cif.gz_A | 1 | 0.251 | 227 | 9.51e-14 | Crystal structure of Halogen methyl transferase from Paraburkholderia xenovorans at 1.8 A in complex with SAH |
| 6mro-assembly1.cif.gz_A | 1 | 0.224 | 223 | 7.297e-12 | Crystal structure of methyl transferase from Methanosarcina acetivorans at 1.6 Angstroms resolution, Northeast Structural Genomics Consortium (NESG) Target MvR53. |

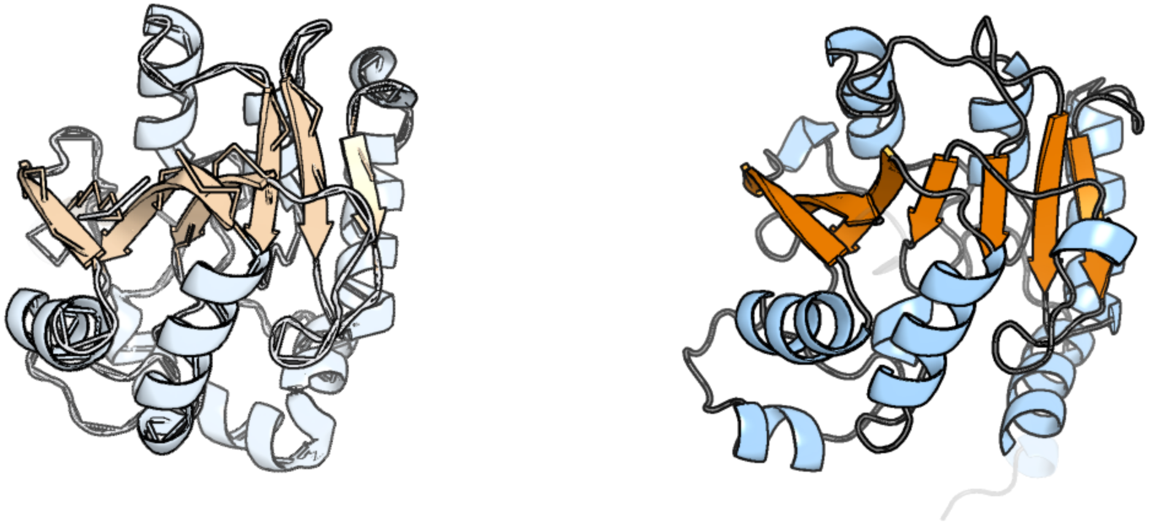

Figure 7: left: reference structure of 3lcc chain A. right: predicted structure of kaa0148314, unaligned sequences are shown as transparent

#### kaa0149273

- Sequence-based annotation for kaa0149273 (predicted in DGTS from hits to PF13649, UniRef50_A0A238JF33) is hypothetical protein FNF28_07376 [Cafeteria roenbergensis]
- Best hit was 5cm2 chain Z: TRNA METHYLTRANSFERASE

| target | prob | fident | alnlen | evalue | theader |
|---|---|---|---|---|---|
| 5cm2-assembly1.cif.gz_Z | 1 | 0.191 | 183 | 1.042e-12 | Structure of Y. lipolytica Trm9-Trm112 complex, a methyltransferase modifying U34 in the anticodon loop of some tRNAs |
| 5ufm-assembly1.cif.gz_A | 1 | 0.178 | 213 | 1.11e-12 | Crystal structure of Burkholderia thailandensis 1,6-didemethyltoxoflavin-N1-methyltransferase with bound 1,6-didemethyltoxoflavin and S-adenosylhomocysteine |
| 5mgz-assembly1.cif.gz_B | 1 | 0.223 | 242 | 1.11e-12 | Streptomyces Spheroides NovO (8-demethylnovbiocic acid methyltransferase) with SAH |

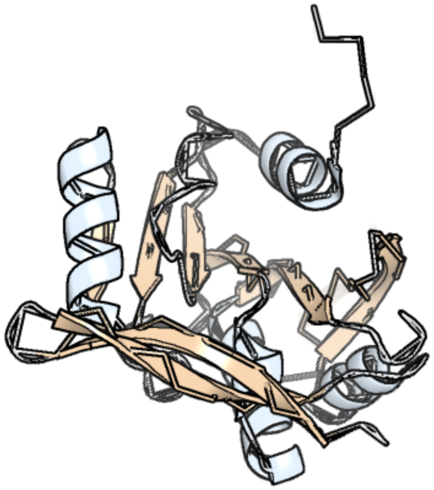

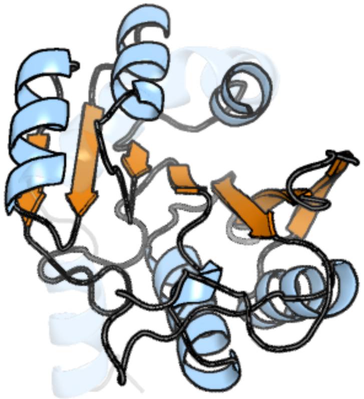

Figure 8: left: reference structure of 5cm2 chain Z. right: predicted structure of kaa0149273, unaligned sequences are shown as transparent

#### kaa0149378

- Sequence-based annotation for kaa0149378 (predicted in Sulfonolipids from hits to PF00106, UniRef50_A0A383U0M1) is hypothetical protein FNF28_07360 [Cafeteria roenbergensis]
- Best hit was 2ilt chain A: Corticosteroid 11-beta-dehydrogenase isozyme 1

| target | prob | fident | alnlen | evalue | theader |
|---|---|---|---|---|---|
| 2ilt-assembly1.cif.gz_A | 1 | 0.355 | 273 | 6.898e-24 | Human 11-beta-Hydroxysteroid Dehydrogenase (HSD1) with NADP and Adamantane Sulfone Inhibitor |
| 2bel-assembly2.cif.gz_C | 1 | 0.362 | 262 | 2.482e-23 | Structure of human 11-beta-hydroxysteroid dehydrogenase in complex with NADP and carbenoxolone |
| 3dwf-assembly3.cif.gz_C | 1 | 0.341 | 275 | 2.482e-23 | Crystal Structure of the Guinea Pig 11beta-Hydroxysteroid Dehydrogenase Type 1 Mutant F278E |

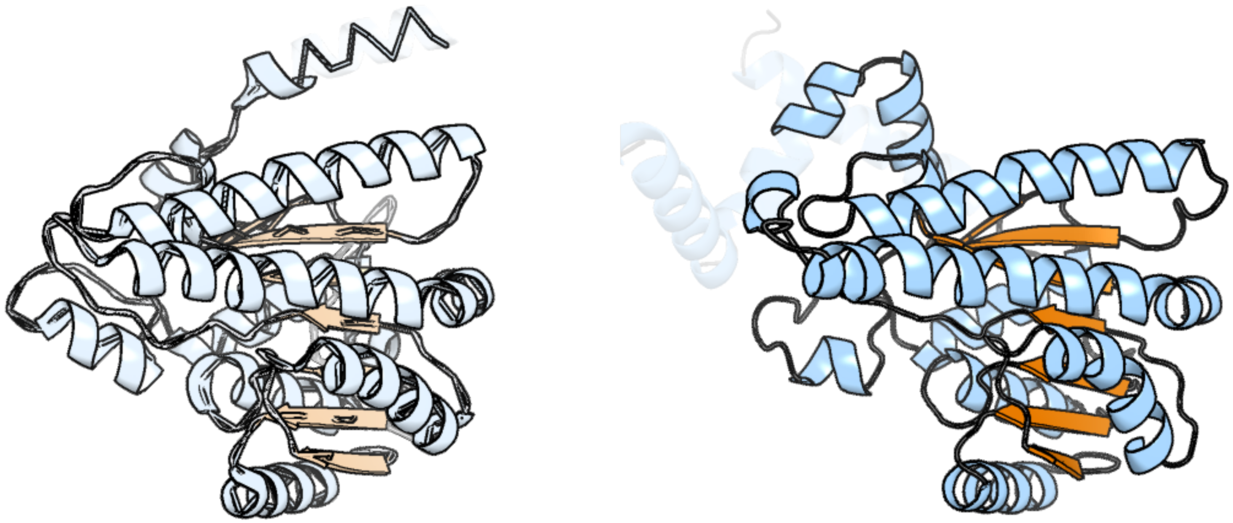

Figure 9: left: reference structure of 2ilt chain A. right: predicted structure of kaa0149378, unaligned sequences are shown as transparent

### kaa0150274

- Sequence-based annotation for kaa0150274 (predicted in DGTS from hits to PF13649) is hypothetical protein FNF28_07275 [Cafeteria roenbergensis]
- Best hit was 2yqz chain B: Hypothetical protein TTHA0223

| target | prob | fident | alnlen | evalue | theader |
|---|---|---|---|---|---|
| 2yqz-assembly2.cif.gz_B | 1 | 0.199 | 276 | 4.406e-15 | Crystal Structure of Hypothetical Methyltransferase TTHA0223 from Thermus thermophilus HB8 complexed with S-adenosylmethionine |
| 5w7k-assembly2.cif.gz_B | 1 | 0.168 | 279 | 2.535e-14 | Crystal structure of OxaG |
| 2yr0-assembly2.cif.gz_B | 1 | 0.183 | 267 | 8.473e-14 | Crystal Structure of Hypothetical Methyltransferase TTHA0223 from Thermus thermophilus HB8 |

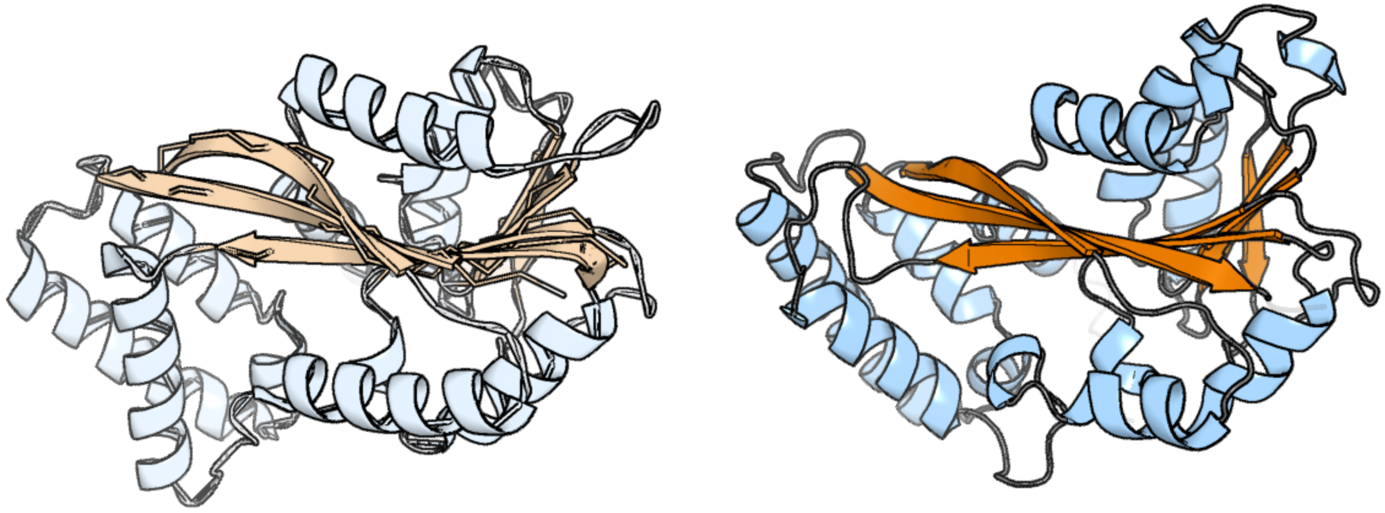

Figure 10: left: reference structure of 2yqz chain B. right: predicted structure of kaa0150274, unaligned sequences are shown as transparent

#### kaa0150947

- Sequence-based annotation for kaa0150947 (predicted in DGTS from hits to PF13649) is hypothetical protein FNF28_07191 [Cafeteria roenbergensis]
- Best hit was 2hnk chain C: SAM-dependent O-methyltransferase

| target | prob | fident | alnlen | evalue | theader |
|---|---|---|---|---|---|
| 2hnk-assembly1.cif.gz_C | 1 | 0.294 | 241 | 1.316e-22 | Crystal structure of SAM-dependent O-methyltransferase from pathogenic bacterium Leptospira interrogans |
| 5log-assembly1.cif.gz_A | 1 | 0.326 | 242 | 2.228e-22 | Crystal Structure of SafC from Myxococcus xanthus bound to SAM |
| 2hnk-assembly2.cif.gz_B-2 | 1 | 0.299 | 244 | 5.361e-22 | Crystal structure of SAM-dependent O-methyltransferase from pathogenic bacterium Leptospira interrogans |

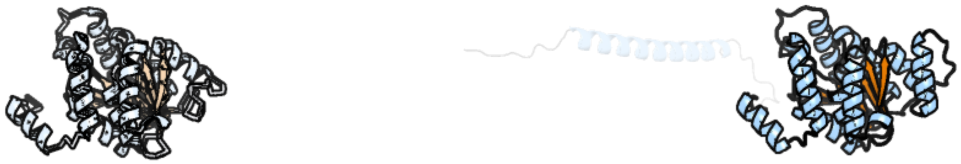

Figure 11: left: reference structure of 2hnk chain C. right: predicted structure of kaa0150947, unaligned sequences are shown as transparent

#### kaa0152887

- Sequence-based annotation for kaa0152887 (predicted in Sulfonolipids from hits to PF00291, UniRef50_A0A2D9B818) is hypothetical protein FNF28_06994 [Cafeteria roenbergensis]
- Best hit was 5i7w chain B: Cysteine synthase A

| target | prob | fident | alnlen | evalue | theader |
|---|---|---|---|---|---|
| 5i7w-assembly2.cif.gz_B | 1 | 0.494 | 340 | 1.108e-42 | Crystal Structure of a Cysteine Synthase from Brucella suis |
| 5jis-assembly1.cif.gz_A | 1 | 0.501 | 327 | 2.121e-42 | The Crystal Structure of O-acetyl serine sulfhydralase from Brucella abortus |
| 5jjc-assembly2.cif.gz_B | 1 | 0.495 | 327 | 1.775e-41 | Crystal Structure of double mutant (Q96A-Y125A) O-Acetyl Serine Sulfhydralase from Brucella abortus |

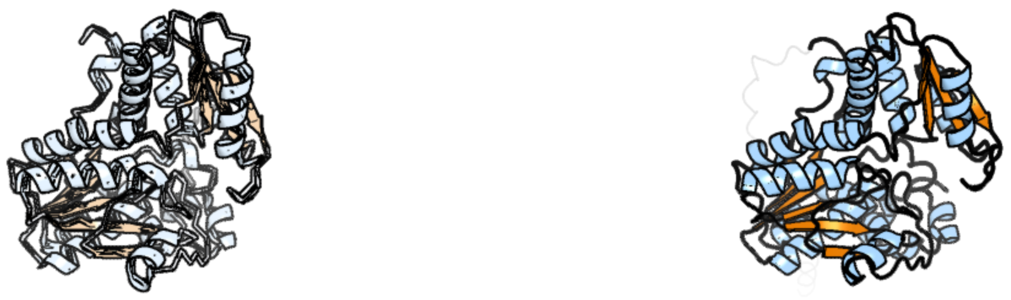

Figure 12: left: reference structure of 5i7w chain B. right: predicted structure of kaa0152887, unaligned sequences are shown as transparent

#### kaa0153987

- Sequence-based annotation for kaa0153987 (predicted in Sulfonolipids from hits to PF00106) is hypothetical protein FNF28_06876 [Cafeteria roenbergensis]
- Best hit was 1ooe chain B: Dihydropteridine reductase

| target | prob | fident | alnlen | evalue | theader |
|---|---|---|---|---|---|
| 1ooe-assembly1.cif.gz_B | 1 | 0.39 | 241 | 4.808e-26 | Structural Genomics of Caenorhabditis elegans : Dihydropteridine reductase |
| 1hdr-assembly1.cif.gz_A-2 | 1 | 0.364 | 239 | 2.892e-25 | THE CRYSTALLOGRAPHIC STRUCTURE OF A HUMAN DIHYDROPTERIDINE REDUCTASE NADH BINARY COMPLEX EXPRESSED IN ESCHERICHIA COLI BY A CDNA CONSTRUCTED FROM ITS RAT HOMOLOGUE |
| 3orf-assembly2.cif.gz_D | 1 | 0.367 | 237 | 5.37e-25 | Crystal Structure of Dihydropteridine Reductase from Dictyostelium discoideum |

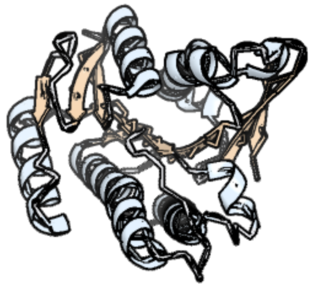

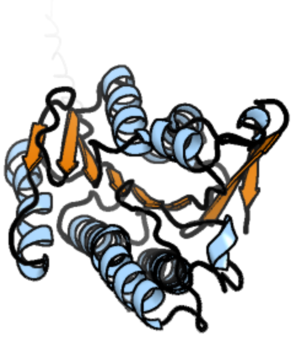

Figure 13: left: reference structure of 1ooe chain B. right: predicted structure of kaa0153987, unaligned sequences are shown as transparent

### kaa0154170

- Sequence-based annotation for kaa0154170 (predicted in DGTS from hits to PF13649) is hypothetical protein FNF28_06855 [Cafeteria roenbergensis]
- Best hit was 5wp5 chain B: Phosphomethylethanolamine N-methyltransferase 2

| target | prob | fident | alnlen | evalue | theader |
|---|---|---|---|---|---|
| 5wp5-assembly2.cif.gz_B | 1 | 0.244 | 254 | 1.108e-16 | Arabidopsis thaliana phosphoethanolamine N-methyltransferase 2 (AtPMT2) in complex with SAH |
| 5wp4-assembly1.cif.gz_A | 1 | 0.246 | 247 | 1.841e-16 | Arabidopsis thaliana phosphoethanolamine N-methyltransferase 1 (AtPMT1, XIOPTL) in complex with SAH and phosphocholine |
| 4kri-assembly1.cif.gz_A | 1 | 0.201 | 253 | 1.011e-14 | Haemonchus contortus Phospholethanolamine N-methyltransferase 2 in complex with phosphomonomethylethanolamine and S-adenosylhomocysteine |

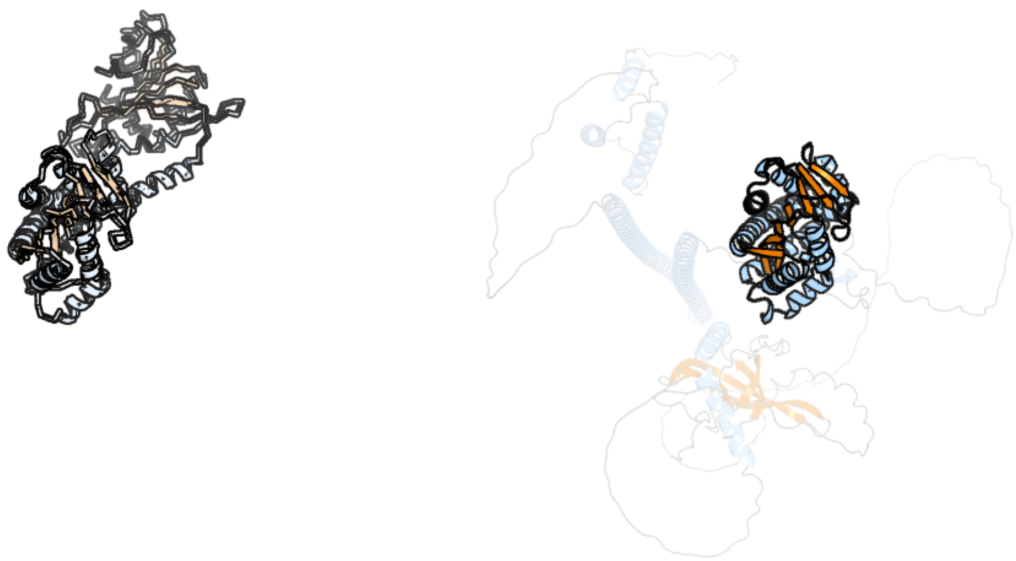

Figure 14: left: reference structure of 5wp5 chain B. right: predicted structure of kaa0154170, unaligned sequences are shown as transparent

#### kaa0155038

- Sequence-based annotation for kaa0155038 (predicted in Fu-FA from hits to PF06966) is hypothetical protein FNF28_06781 [Cafeteria roenbergensis]
- No significant structural hit found

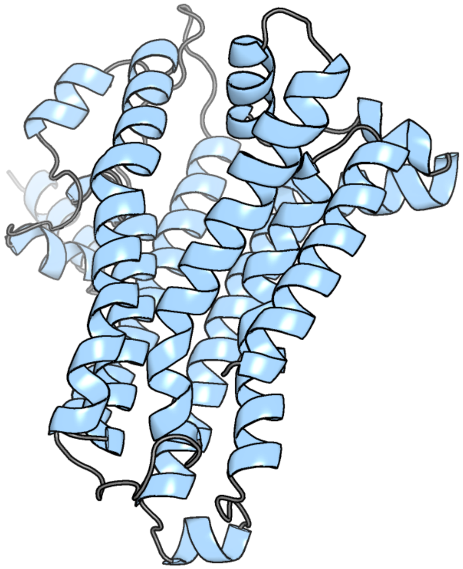

Figure 15: predicted structure of kaa0155038

### kaa0155827

- Sequence-based annotation for kaa0155827 (predicted in Sulfonolipids from hits to PF00155) is hypothetical protein FNF28_06687 [Cafeteria roenbergensis]
- Best hit was 6eei chain A: Tyrosine decarboxylase 1

| target | prob | fident | alnlen | evalue | theader |
|---|---|---|---|---|---|
| 6eei-assembly1.cif.gz_A | 1 | 0.483 | 488 | 1.917e-57 | Crystal structure of Arabidopsis thaliana phenylacetaldehyde synthase in complex with L-phenylalanine |
| 6eei-assembly1.cif.gz_B | 1 | 0.484 | 489 | 2.796e-57 | Crystal structure of Arabidopsis thaliana phenylacetaldehyde synthase in complex with L-phenylalanine |
| 6eem-assembly1.cif.gz_B | 1 | 0.441 | 507 | 7.787e-57 | Crystal structure of Papaver somniferum tyrosine decarboxylase in complex with L-tyrosine |

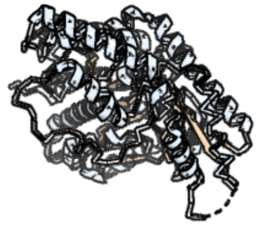

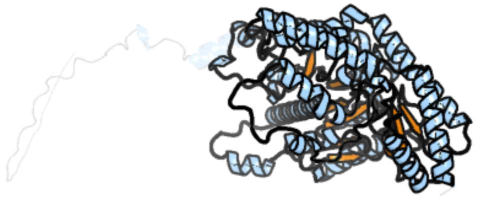

Figure 16: left: reference structure of 6eei chain A. right: predicted structure of kaa0155827, unaligned sequences are shown as transparent

#### kaa0156556

- Sequence-based annotation for kaa0156556 (predicted in Sulfonolipids from hits to PF00155, UniRef90_A0A7Z8YEE0, UniRef90_D4IKV7) is hypothetical protein FNF28_06614 [Cafeteria roenbergensis]
- Best hit was 7x98 chain D: 5-aminolevulinate synthase

| target | prob | fident | alnlen | evalue | theader |
|---|---|---|---|---|---|
| 7x98-assembly2.cif.gz_D | 1 | 0.528 | 422 | 4.121e-53 | 5-Aminolevulinate synthase HemA from Rhodopseudomonas palustris |
| 2bwo-assembly2.cif.gz_D | 1 | 0.502 | 418 | 4.178e-52 | 5-Aminolevulinate Synthase from Rhodobacter capsulatus in complex with succinyl-CoA |
| 5qqr-assembly1.cif.gz_B | 1 | 0.462 | 428 | 5.282e-51 | PanDDA analysis group deposition – Crystal Structure of human ALAS2A in complex with Z1171217421 |

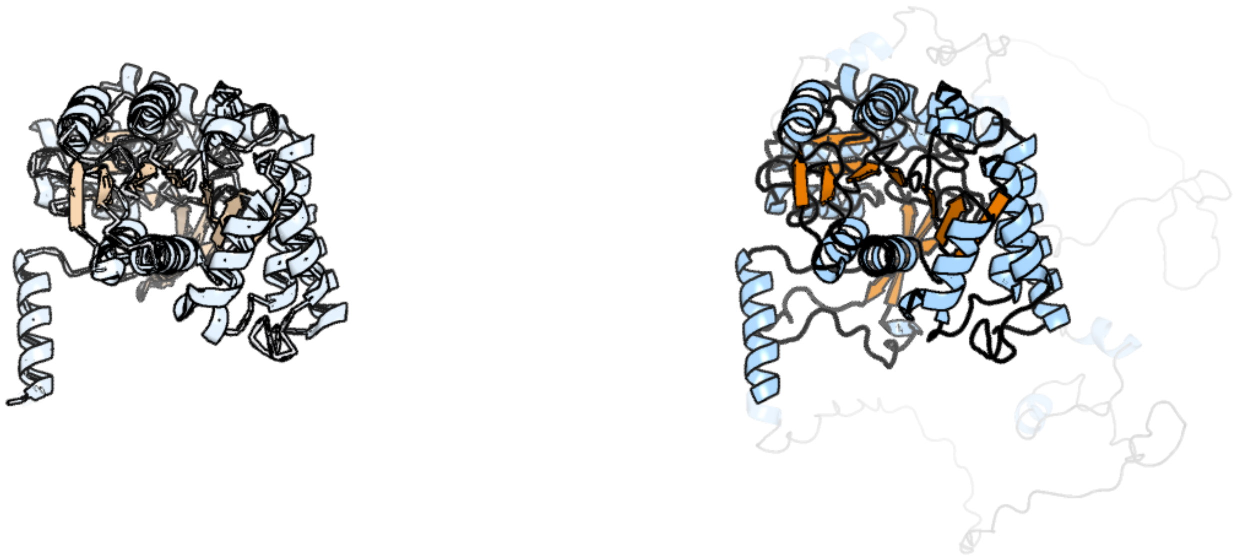

Figure 17: left: reference structure of 7x98 chain D. right: predicted structure of kaa0156556, unaligned sequences are shown as transparent

### kaa0156557

- Sequence-based annotation for kaa0156557 (predicted in DGTS from hits to PF13649) is hypothetical protein FNF28_06615 [Cafeteria roenbergensis]
- Best hit was 1xtp chain A: LMAJ004091AAA

| target | prob | fident | alnlen | evalue | theader |
|---|---|---|---|---|---|
| 1xtp-assembly1.cif.gz_A | 1 | 0.313 | 214 | 1.01e-24 | Structural Analysis of Leishmania major LMAJ004091AAA, a SAM-dependent methyltransferase of the DUF858/Pfam05891 family |
| 6wh8-assembly1.cif.gz_B | 1 | 0.275 | 218 | 9.742e-24 | The structure of NTMT1 in complex with compound BM-30 |
| 2ex4-assembly1.cif.gz_B | 1 | 0.27 | 218 | 2.759e-23 | Crystal Structure of Human methyltransferase AD-003 in complex with S-adenosyl-L-homocysteine |

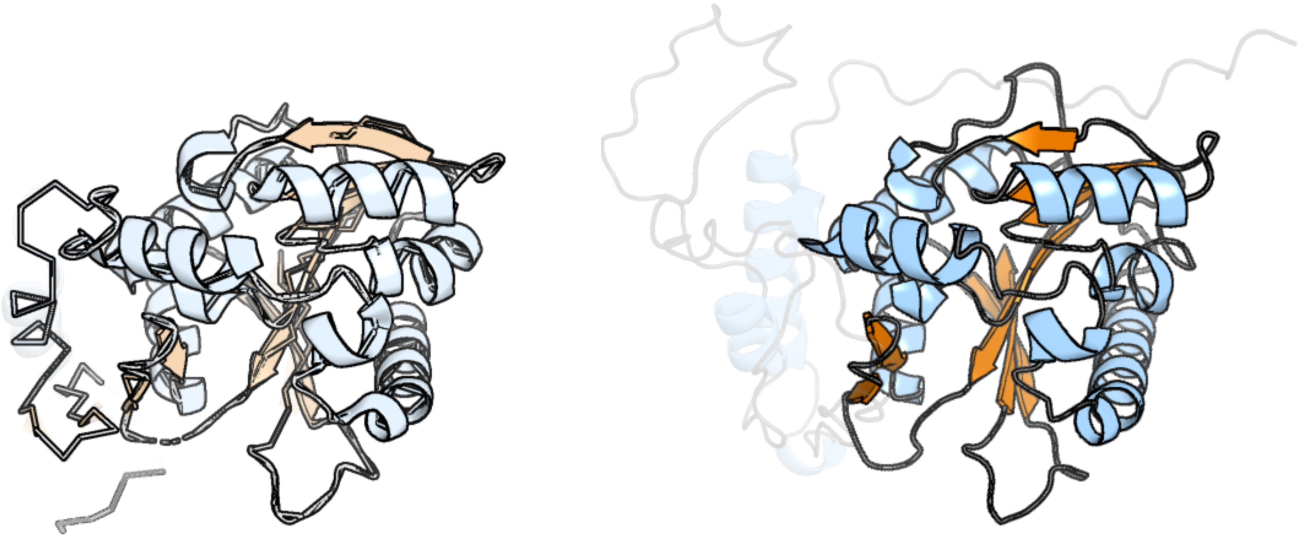

Figure 18: left: reference structure of 1xtp chain A. right: predicted structure of kaa0156557, unaligned sequences are shown as transparent

#### kaa0157937

- Sequence-based annotation for kaa0157937 (predicted in Fu-FA from hits to UniRef50_Q5LTD2) is hypothetical protein FNF28_06444 [Cafeteria roenbergensis]
- Best hit was 5kmp chain A: FAD-dependent pyridine nucleotide-disulfide oxidoreductase

| target | prob | fident | alnlen | evalue | theader |
|---|---|---|---|---|---|
| 5kmp-assembly1.cif.gz_A | 1 | 0.181 | 469 | 9.377e-22 | The structure of G164E variant of type II NADH dehydrogenase from Caldalkalibacillus thermarum |
| 4nwz-assembly1.cif.gz_A | 1 | 0.182 | 466 | 1.111e-21 | Structure of bacterial type II NADH dehydrogenase from Caldalkalibacillus thermarum at 2.5A resolution |
| 5na4-assembly1.cif.gz_A | 1 | 0.182 | 456 | 1.176e-21 | NADH:quinone oxidoreductase (NDH-II) from Staphylococcus aureus - E172S mutant |

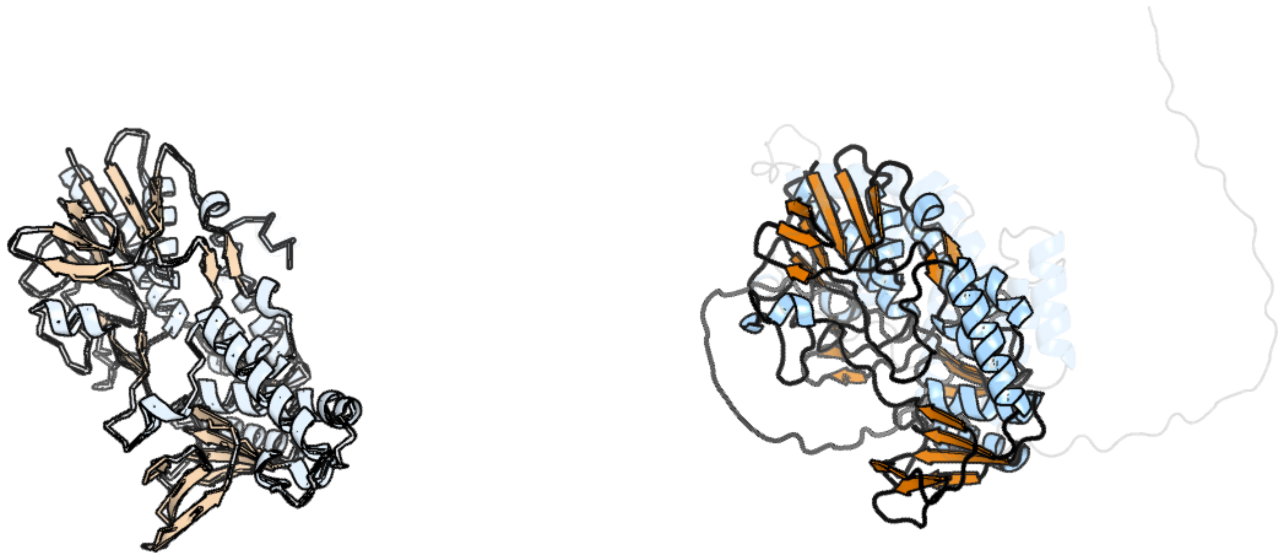

Figure 19: left: reference structure of 5kmp chain A. right: predicted structure of kaa0157937, unaligned sequences are shown as transparent

#### kaa0157993

- Sequence-based annotation for kaa0157993 (predicted in Sulfonolipids from hits to PF00106) is hypothetical protein FNF28_06416 [Cafeteria roenbergensis]
- Best hit was 2wyv chain D: ENOYL-[ACYL CARRIER PROTEIN] REDUCTASE

| target | prob | fident | alnlen | evalue | theader |
|---|---|---|---|---|---|
| 2wyv-assembly1.cif.gz_D | 1 | 0.413 | 232 | 1.541e-19 | High resolution structure of Thermus thermophilus enoyl-acyl carrier protein reductase NAD-form |
| 2yw9-assembly2.cif.gz_G | 1 | 0.409 | 232 | 8.965e-19 | Crystal structure of TT0143 from Thermus thermophilus HB8 |
| 7fcm-assembly1.cif.gz_B | 1 | 0.377 | 236 | 1.429e-18 | Crystal structure of Moraxella catarrhalis enoyl-ACP-reductase (FabI) in complex with NAD and Triclosan |

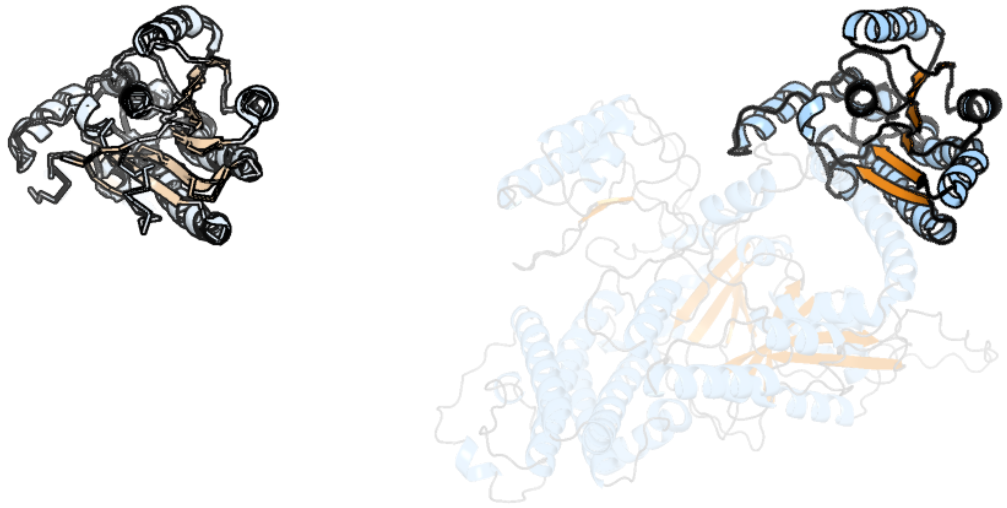

Figure 20: left: reference structure of 2wyv chain D. right: predicted structure of kaa0157993, unaligned sequences are shown as transparent

#### kaa0158001

- Sequence-based annotation for kaa0158001 (predicted in Sulfonolipids from hits to PF00106, UniRef50_A0A383U0M1) is hypothetical protein FNF28_06424 [Cafeteria roenbergensis]
- Best hit was 3rd5 chain A: MYPAA.01249.C

| target | prob | fident | alnlen | evalue | theader |
|---|---|---|---|---|---|
| 3rd5-assembly1.cif.gz_A | 1 | 0.333 | 327 | 3.51e-23 | Crystal structure of a putative uncharacterized protein from Mycobacterium Paratuberculosis |
| 7jk9-assembly1.cif.gz_A | 1 | 0.244 | 344 | 3.271e-19 | Helical filaments of plant light-dependent protochlorophyllide oxidoreductase (LPOR) bound to NADPH, Pchlide, and membrane |
| 6l1h-assembly2.cif.gz_B | 1 | 0.246 | 332 | 4.123e-19 | Crystal structure of light-dependent protochlorophyllide oxidoreductase from Thermosynechococcus elongatus |

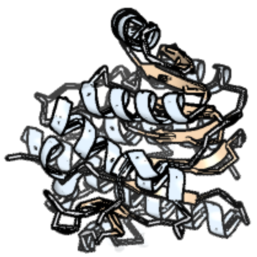

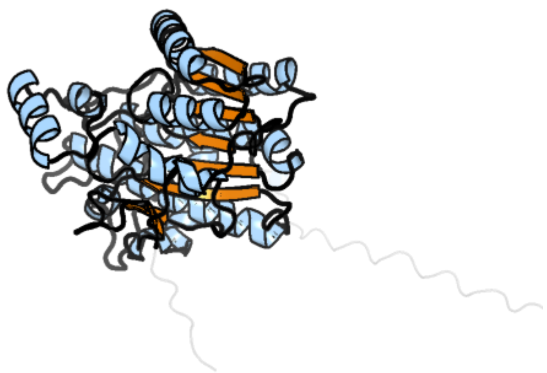

Figure 21: left: reference structure of 3rd5 chain A. right: predicted structure of kaa0158001, unaligned sequences are shown as transparent

### kaa0158005

- Sequence-based annotation for kaa0158005 (predicted in Sulfonolipids from hits to PF00106) is hypothetical protein FNF28_06428 [Cafeteria roenbergensis]
- Best hit was 6l1h chain B: NADPH-protochlorophyllide oxidoreductase

| target | prob | fident | alnlen | evalue | theader |
|---|---|---|---|---|---|
| 6l1h-assembly2.cif.gz_B | 1 | 0.145 | 459 | 1.283e-11 | Crystal structure of light-dependent protochlorophyllide oxidoreductase from Thermosynechococcus elongatus |
| 6l1h-assembly1.cif.gz_A | 1 | 0.152 | 451 | 2.117e-11 | Crystal structure of light-dependent protochlorophyllide oxidoreductase from Thermosynechococcus elongatus |
| 6rnv-assembly1.cif.gz_A | 1 | 0.149 | 448 | 1.853e-10 | The crystal structure of Thermosynechococcus elongatus protochlorophyllide oxidoreductase (POR) |

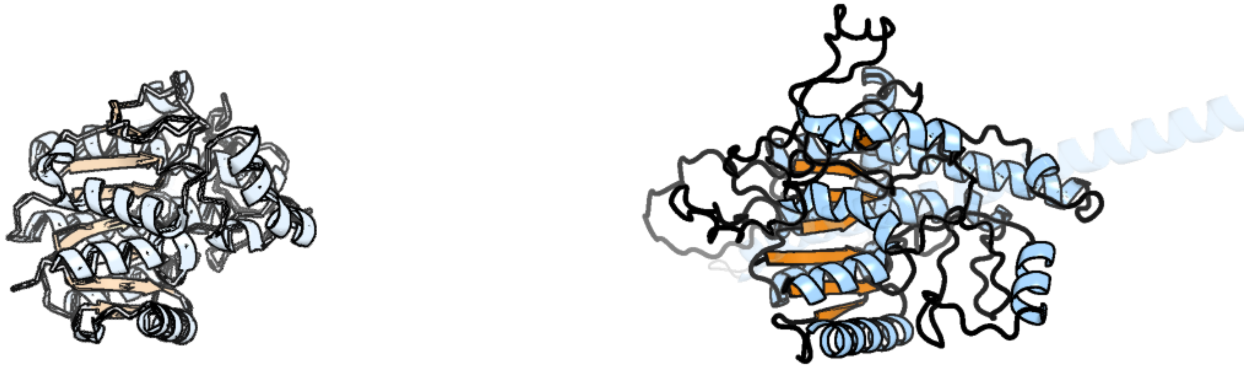

Figure 22: left: reference structure of 6l1h chain B. right: predicted structure of kaa0158005, unaligned sequences are shown as transparent

#### kaa0158301

- Sequence-based annotation for kaa0158301 (predicted in PE_PMME from hits to PF01066, UniRef50_P22140, UniRef50_Q9Y6K0) is hypothetical protein FNF28_06309 [Cafeteria roenbergensis]
- Best hit was 8gyw chain B: Choline/ethanolaminephosphotransferase 1

| target | prob | fident | alnlen | evalue | theader |
|---|---|---|---|---|---|
| 8gyw-assembly1.cif.gz_B | 1 | 0.197 | 446 | 1.425e-13 | Cryo-EM structure of human CEPT1 complexed with CDP-choline |
| 8ero-assembly1.cif.gz_A | 1 | 0.204 | 444 | 1.074e-12 | Structure of Xenopus cholinephosphotransferase1 in complex with CDP |
| 6tod-assembly2.cif.gz_B | 0.196 | 0.128 | 396 | 0.07233 | Crystal structure of the Orexin-1 receptor in complex with EMPA |

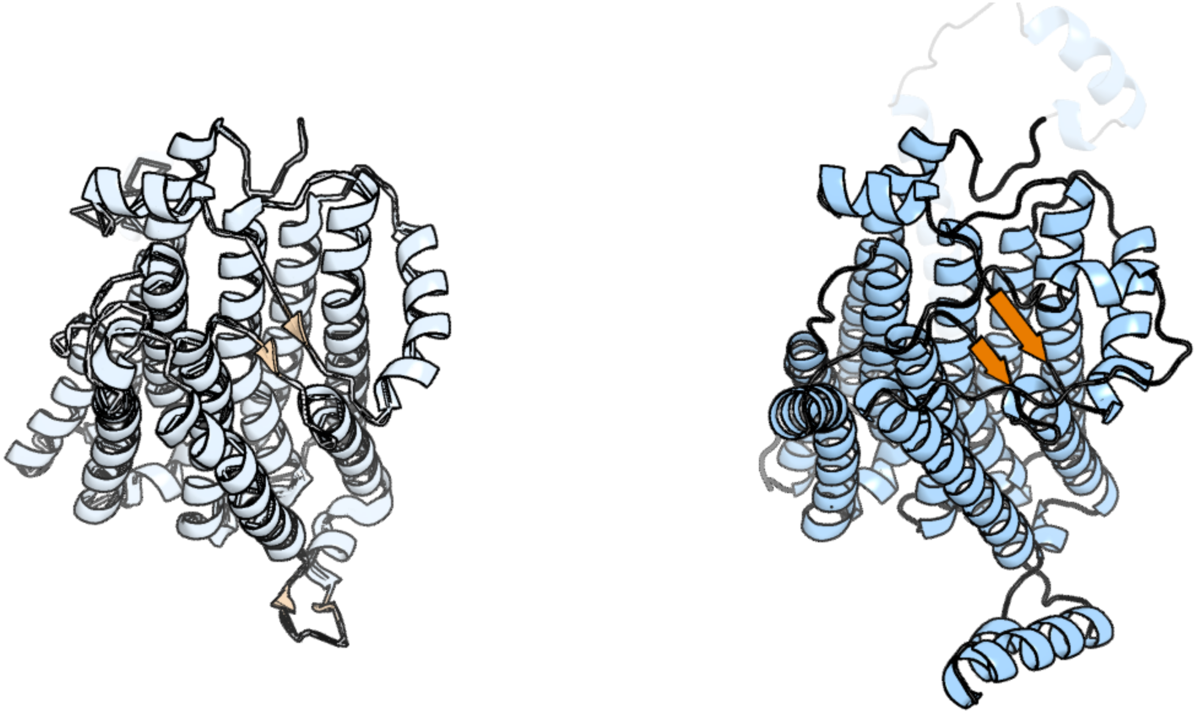

Figure 23: left: reference structure of 8gyw chain B. right: predicted structure of kaa0158301, unaligned sequences are shown as transparent

#### kaa0158689

- Sequence-based annotation for kaa0158689 (predicted in Sulfonolipids from hits to PF00106, UniRef50_A0A383U0M1) is hypothetical protein FNF28_06124 [Cafeteria roenbergensis]
- Best hit was 6d9y chain A: Short-chain dehydrogenase/reductase SDR

| target | prob | fident | alnlen | evalue | theader |
|---|---|---|---|---|---|
| 6d9y-assembly1.cif.gz_A | 1 | 0.449 | 247 | 1.801e-30 | Crystal structure of a short chain dehydrogenase/reductase SDR from Burkholderia phymatum with partially occupied NAD |
| 5x8h-assembly1.cif.gz_B | 1 | 0.321 | 246 | 1.112e-27 | Crystal structure of the ketone reductase ChKRED20 from the genome of Chryseobacterium sp. CA49 |
| 7tzp-assembly3.cif.gz_G | 1 | 0.362 | 248 | 2.521e-26 | Crystal Structure of Putataive Short-Chain Dehydrogenase/Reductase (FabG) from Klebsiella pneumoniae subsp. pneumoniae NTUH-K2044 in Complex with NADH |

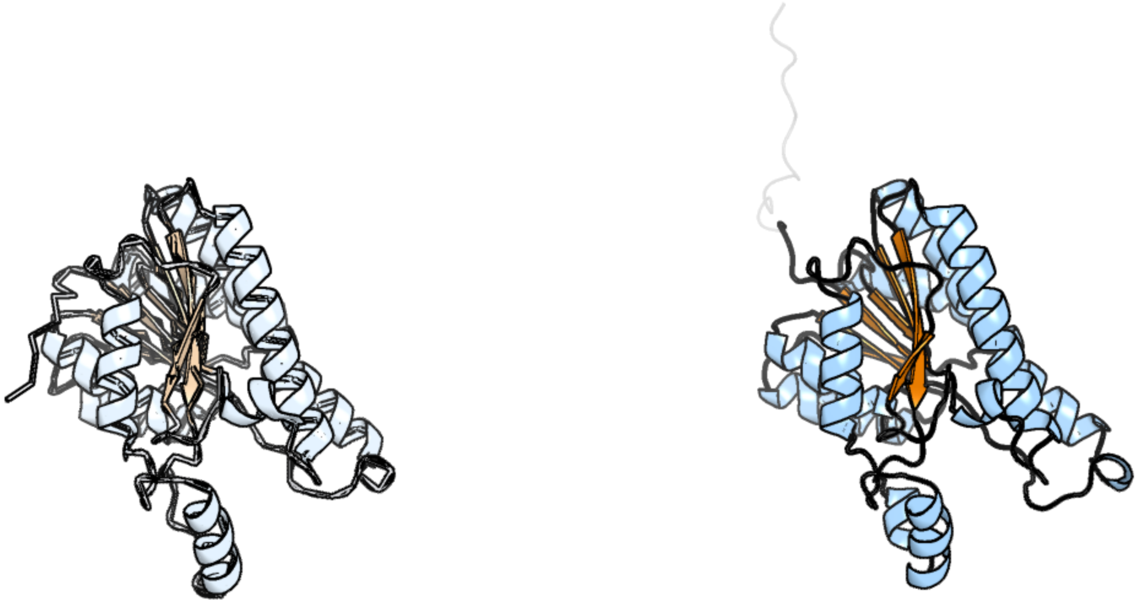

Figure 24: left: reference structure of 6d9y chain A. right: predicted structure of kaa0158689, unaligned sequences are shown as transparent

### kaa0158823

- Sequence-based annotation for kaa0158823 (predicted in Sulfonolipids from hits to PF00106) is hypothetical protein FNF28_06068 [Cafeteria roenbergensis]
- Best hit was 1yxm chain B: peroxisomal trans 2-enoyl CoA reductase

| target | prob | fident | alnlen | evalue | theader |
|---|---|---|---|---|---|
| 1yxm-assembly1.cif.gz_B | 1 | 0.41 | 278 | 1.259e-32 | Crystal structure of peroxisomal trans 2-enoyl CoA reductase |
| 1yxm-assembly1.cif.gz_A | 1 | 0.417 | 278 | 2.721e-32 | Crystal structure of peroxisomal trans 2-enoyl CoA reductase |
| 1yxm-assembly1.cif.gz_D | 1 | 0.418 | 277 | 8.47e-31 | Crystal structure of peroxisomal trans 2-enoyl CoA reductase |

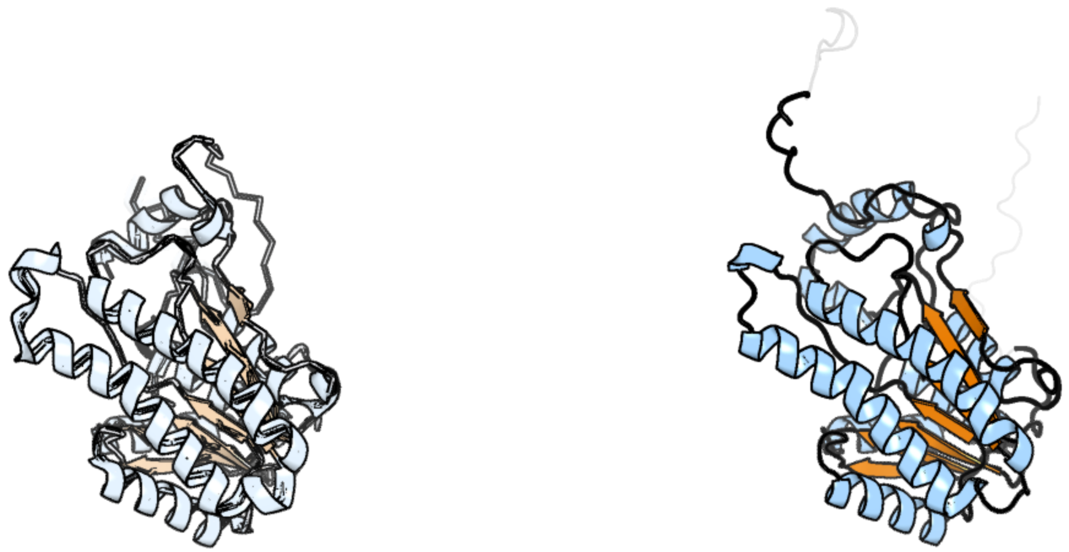

Figure 25: left: reference structure of 1yxm chain B. right: predicted structure of kaa0158823, unaligned sequences are shown as transparent

#### kaa0159043

- Sequence-based annotation for kaa0159043 (predicted in Sulfonolipids from hits to PF00106) is hypothetical protein FNF28_06017 [Cafeteria roenbergensis]
- Best hit was 3oml chain A: Peroxisomal Multifunctional Enzyme Type 2, CG3415

| target | prob | fident | alnlen | evalue | theader |
|---|---|---|---|---|---|
| 3oml-assembly1.cif.gz_A | 1 | 0.319 | 680 | 2.907e-51 | Structure of full-length peroxisomal multifunctional enzyme type 2 from Drosophila melanogaster |
| 1zbq-assembly3.cif.gz_F | 1 | 0.557 | 305 | 3.536e-39 | Crystal Structure Of Human 17-Beta-Hydroxysteroid Dehydrogenase Type 4 In Complex With NAD |
| 1gz6-assembly2.cif.gz_D | 1 | 0.544 | 305 | 5.405e-38 | (3R)-HYDROXYACYL-COA DEHYDROGENASE FRAGMENT OF RAT PEROXISOMAL MULTIFUNCTIONAL ENZYME TYPE 2 |

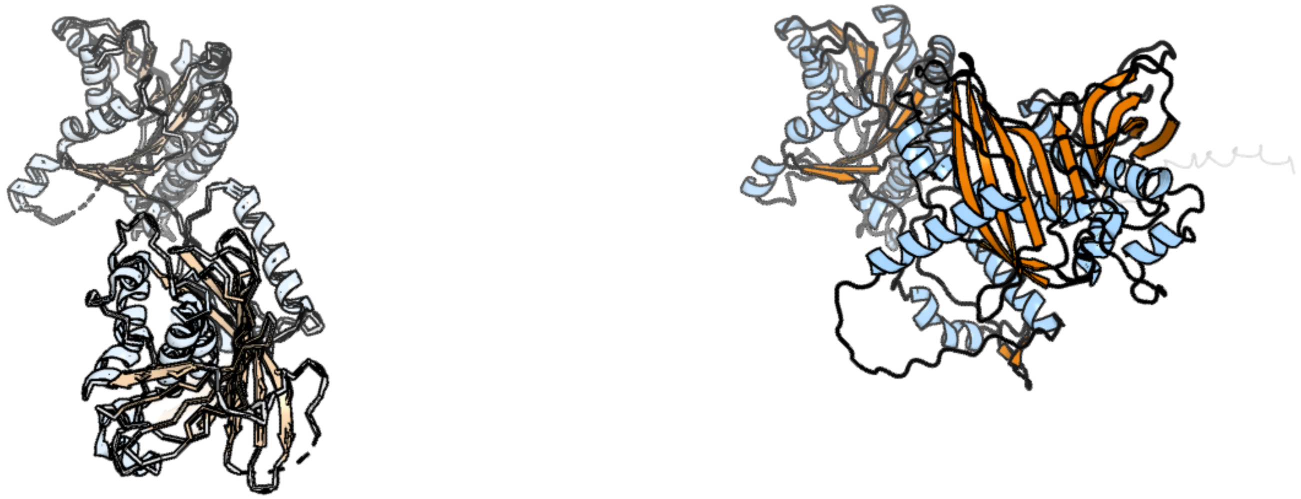

Figure 26: left: reference structure of 3oml chain A. right: predicted structure of kaa0159043, unaligned sequences are shown as transparent

#### kaa0159500

- Sequence-based annotation for kaa0159500 (predicted in Sulfonolipids from hits to PF00106, UniRef50_A0A383U0M1) is hypothetical protein FNF28_05857 [Cafeteria roenbergensis]
- Best hit was 5fyd chain A: OXIDOREDUCTASE, SHORT CHAIN DEHYDROGENASE/REDUCTASE FAMILY PROTEIN

| target | prob | fident | alnlen | evalue | theader |
|---|---|---|---|---|---|
| 5fyd-assembly1.cif.gz_A | 1 | 0.227 | 264 | 8.953e-18 | Structural and biochemical insights into 7beta-hydroxysteroid dehydrogenase stereoselectivity |
| 4yag-assembly1.cif.gz_B | 1 | 0.223 | 259 | 2.199e-16 | Crystal structure of LigL in complex with NADH from Sphingobium sp. strain SYK-6 |
| 3tjr-assembly1.cif.gz_A | 1 | 0.252 | 257 | 2.336e-16 | Crystal structure of a Rv0851c ortholog short chain dehydrogenase from Mycobacterium paratuberculosis |

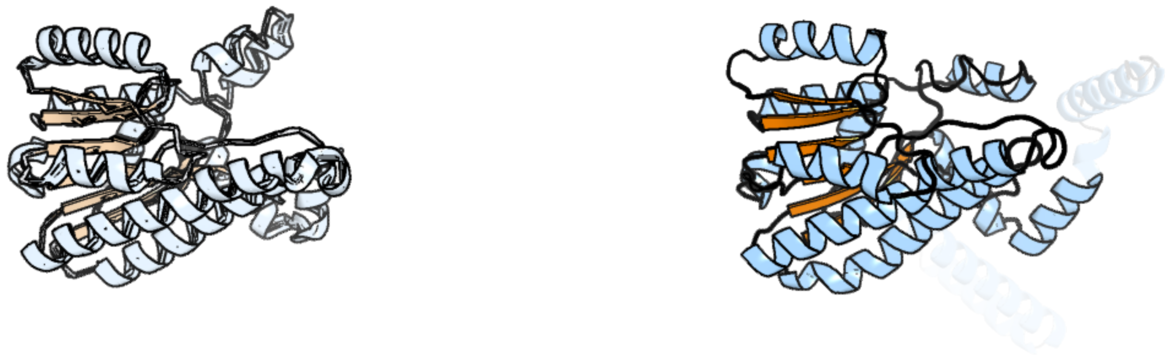

Figure 27: left: reference structure of 5fyd chain A. right: predicted structure of kaa0159500, unaligned sequences are shown as transparent

#### kaa0159680

- Sequence-based annotation for kaa0159680 (predicted in DGTS from hits to PF13649) is hypothetical protein FNF28_05785 [Cafeteria roenbergensis]
- Best hit was 6g4w chain q: Probable 18S rRNA (guanine-N(7))-methyltransferase

| target | prob | fident | alnlen | evalue | theader |
|---|---|---|---|---|---|
| 6g4w-assembly1.cif.gz_q | 1 | 0.591 | 191 | 2.284e-29 | Cryo-EM structure of a late human pre-40S ribosomal subunit - State A |
| 7wts-assembly1.cif.gz_q | 1 | 0.414 | 314 | 2.837e-28 | Cryo-EM structure of a human pre-40S ribosomal subunit - State UTP14 |
| 4qtu-assembly1.cif.gz_B | 1 | 0.526 | 192 | 5.698e-27 | Structure of S. cerevisiae Bud23-Trm112 complex involved in formation of m7G1575 on 18S rRNA (SAM bound form) |

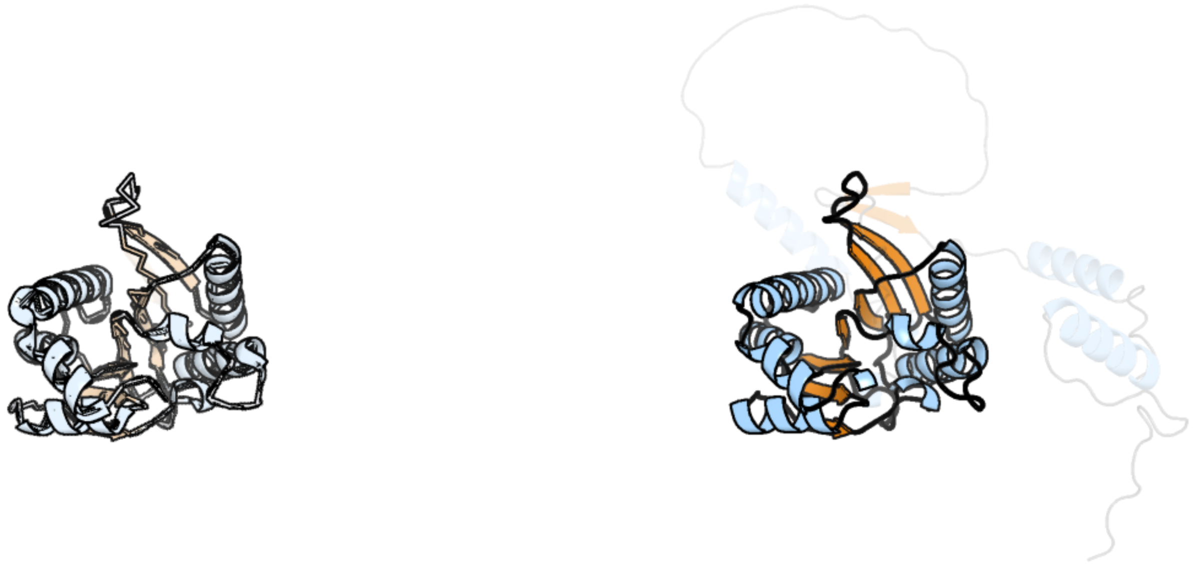

Figure 28: left: reference structure of 6g4w chain q. right: predicted structure of kaa0159680, unaligned sequences are shown as transparent

#### kaa0159710

- Sequence-based annotation for kaa0159710 (predicted in Sulfonolipids from hits to PF00106, UniRef50_A0A383U0M1) is hypothetical protein FNF28_05770 [Cafeteria roenbergensis]
- Best hit was 3rd5 chain A: MYPAA.01249.C

| target | prob | fident | alnlen | evalue | theader |
|---|---|---|---|---|---|
| 3rd5-assembly1.cif.gz_A | 1 | 0.348 | 330 | 9.568e-23 | Crystal structure of a putative uncharacterized protein from Mycobacterium Paratuberculosis |
| 7jk9-assembly1.cif.gz_A | 1 | 0.274 | 346 | 1.467e-19 | Helical filaments of plant light-dependent protochlorophyllide oxidoreductase (LPOR) bound to NADPH, Pchlide, and membrane |
| 6l1h-assembly2.cif.gz_B | 1 | 0.24 | 332 | 2.715e-19 | Crystal structure of light-dependent protochlorophyllide oxidoreductase from Thermosynechococcus elongatus |

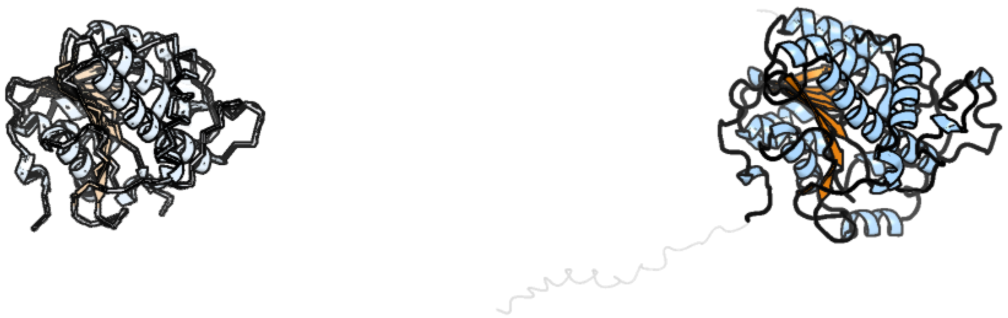

Figure 29: left: reference structure of 3rd5 chain A. right: predicted structure of kaa0159710, unaligned sequences are shown as transparent

#### kaa0159739

- Sequence-based annotation for kaa0159739 (predicted in Fu-FA from hits to PF01593) is hypothetical protein FNF28_05746 [Cafeteria roenbergensis]
- Best hit was 2hko chain A: Lysine-specific histone demethylase 1

| target | prob | fident | alnlen | evalue | theader |
|---|---|---|---|---|---|
| 2hko-assembly1.cif.gz_A | 1 | 0.192 | 317 | 4.639e-10 | Crystal structure of LSD1 |
| 4bay-assembly1.cif.gz_A | 1 | 0.191 | 323 | 1.028e-09 | Phosphomimetic mutant of LSD1-8a splicing variant in complex with CoREST |
| 7xw8-assembly1.cif.gz_A | 1 | 0.181 | 320 | 1.896e-09 | Crystal structure of Lysine Specific Demethylase 1 (LSD1) with TAK-418 distomer, FAD-adduct |

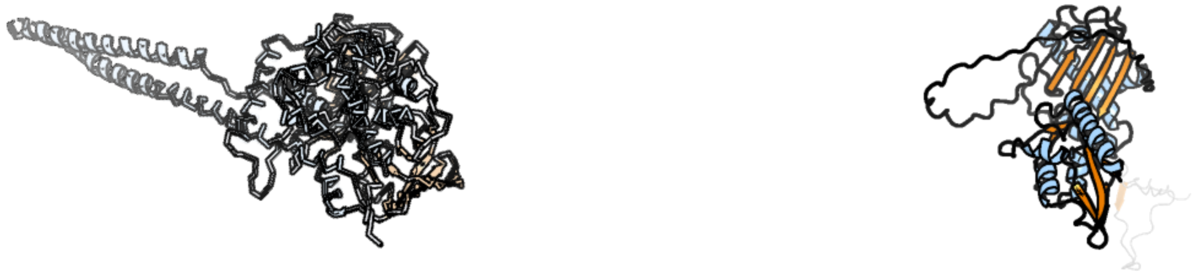

Figure 30: left: reference structure of 2hko chain A. right: predicted structure of kaa0159739, unaligned sequences are shown as transparent

### kaa0159740

- Sequence-based annotation for kaa0159740 (predicted in Fu-FA from hits to PF01593, UniRef50_Q5LTD2) is hypothetical protein FNF28_05747 [Cafeteria roenbergensis]
- No significant structural hit found

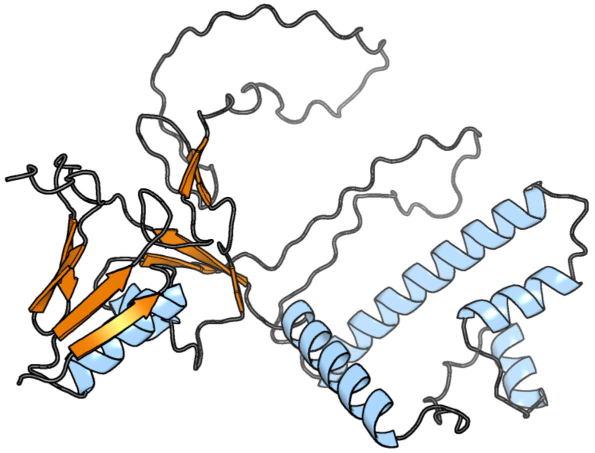

Figure 31: predicted structure of kaa0159740

#### kaa0159818

- Sequence-based annotation for kaa0159818 (predicted in DGTS from hits to PF13649) is hypothetical protein FNF28_05672 [Cafeteria roenbergensis]
- Best hit was 7mqa chain NL: Probable dimethyladenosine transferase

| target | prob | fident | alnlen | evalue | theader |
|---|---|---|---|---|---|
| 7mqa-assembly1.cif.gz_NL | 1 | 0.377 | 297 | 6.412e-29 | Cryo-EM structure of the human SSU processome, state post-A1 |
| 6zqg-assembly1.cif.gz_JL | 1 | 0.36 | 300 | 2.843e-28 | Cryo-EM structure of the 90S pre-ribosome from Saccharomyces cerevisiae, state Dis-C |
| 2h1r-assembly2.cif.gz_B | 1 | 0.331 | 296 | 1.803e-27 | Crystal structure of a dimethyladenosine transferase from Plasmodium falciparum |

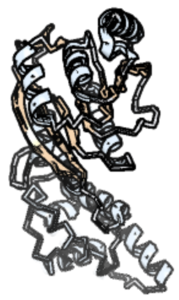

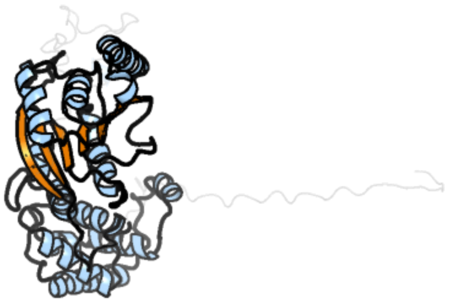

Figure 32: left: reference structure of 7mqa chain NL. right: predicted structure of kaa0159818, unaligned sequences are shown as transparent

#### kaa0159913

- Sequence-based annotation for kaa0159913 (predicted in Sulfonolipids from hits to PF00106) is hypothetical protein FNF28_05604 [Cafeteria roenbergensis]
- Best hit was 1nas chain A: SEPIAPTERIN REDUCTASE

| target | prob | fident | alnlen | evalue | theader |
|---|---|---|---|---|---|
| 1nas-assembly1.cif.gz_A | 1 | 0.297 | 292 | 3.644e-21 | SEPIAPTERIN REDUCTASE COMPLEXED WITH N-ACETYL SEROTONIN |
| 7dsf-assembly1.cif.gz_B | 1 | 0.301 | 289 | 1.184e-20 | The Crystal Structure of human SPR from Biortus. |
| 1z6z-assembly3.cif.gz_E | 1 | 0.302 | 288 | 1.898e-20 | Crystal Structure of Human Sepiapterin Reductase in complex with NADP+ |

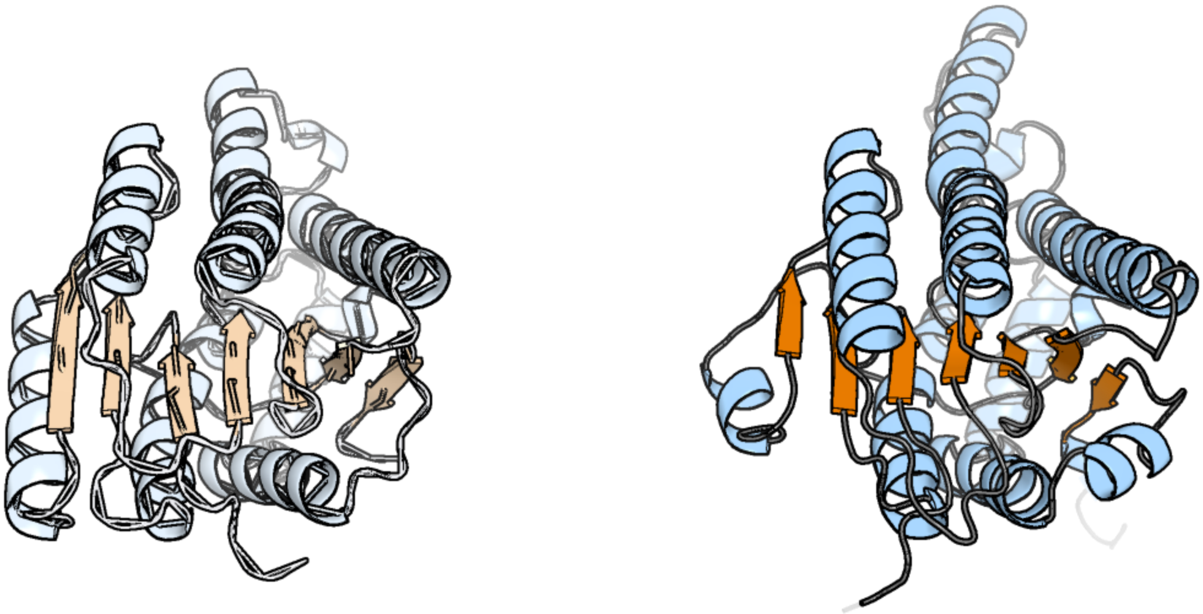

Figure 33: left: reference structure of 1nas chain A. right: predicted structure of kaa0159913, unaligned sequences are shown as transparent

#### kaa0160035

- Sequence-based annotation for kaa0160035 (predicted in Sulfonolipids from hits to PF00106) is hypothetical protein FNF28_05574 [Cafeteria roenbergensis]
- Best hit was 7xpq chain A: UDP-glucose 4-epimerase

| target | prob | fident | alnlen | evalue | theader |
|---|---|---|---|---|---|
| 7xpq-assembly1.cif.gz_A | 1 | 0.522 | 354 | 3.699e-46 | Crystal Structure of UDP-Glc/GlcNAc 4-Epimerase with NAD/UDP-GlcNAc |
| 3enk-assembly1.cif.gz_B | 1 | 0.507 | 351 | 1.41e-45 | 1.9A crystal structure of udp-glucose 4-epimerase from burkholderia pseudomallei |
| 7xpo-assembly1.cif.gz_B | 1 | 0.517 | 352 | 1.623e-44 | Crystal Structure of UDP-Glc/GlcNAc 4-Epimerase with NAD/UDP-Glc |

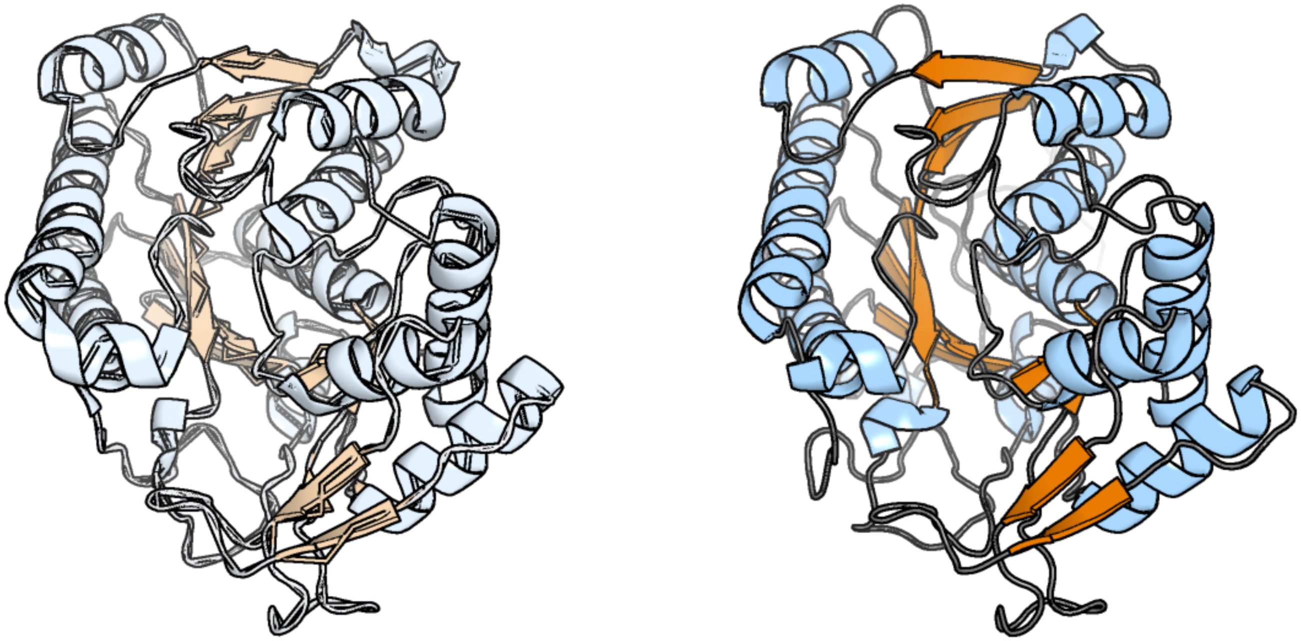

Figure 34: left: reference structure of 7xpq chain A. right: predicted structure of kaa0160035, unaligned sequences are shown as transparent

#### kaa0160201

- Sequence-based annotation for kaa0160201 (predicted in Sulfonolipids from hits to PF00155, UniRef90_A0A7Z8YEE0, UniRef90_D4IKV7) is hypothetical protein FNF28_05519 [Cafeteria roenbergensis]
- Best hit was 7k0m chain A: Serine palmitoyltransferase 1

| target | prob | fident | alnlen | evalue | theader |
|---|---|---|---|---|---|
| 7k0m-assembly1.cif.gz_A | 1 | 0.378 | 497 | 4.485e-47 | Human serine palmitoyltransferase complex SPTLC1/SPLTC2/ssSPTa/ORMDL3, class 1 |
| 7yj2-assembly1.cif.gz_A | 1 | 0.381 | 469 | 2.896e-46 | Cryo-EM structure of SPT-ORMDL3 (ORMDL3-N13A) complex |
| 7yjm-assembly1.cif.gz_A | 1 | 0.321 | 467 | 1.028e-41 | Cryo-EM structure of the monomeric atSPT-ORM1 complex |

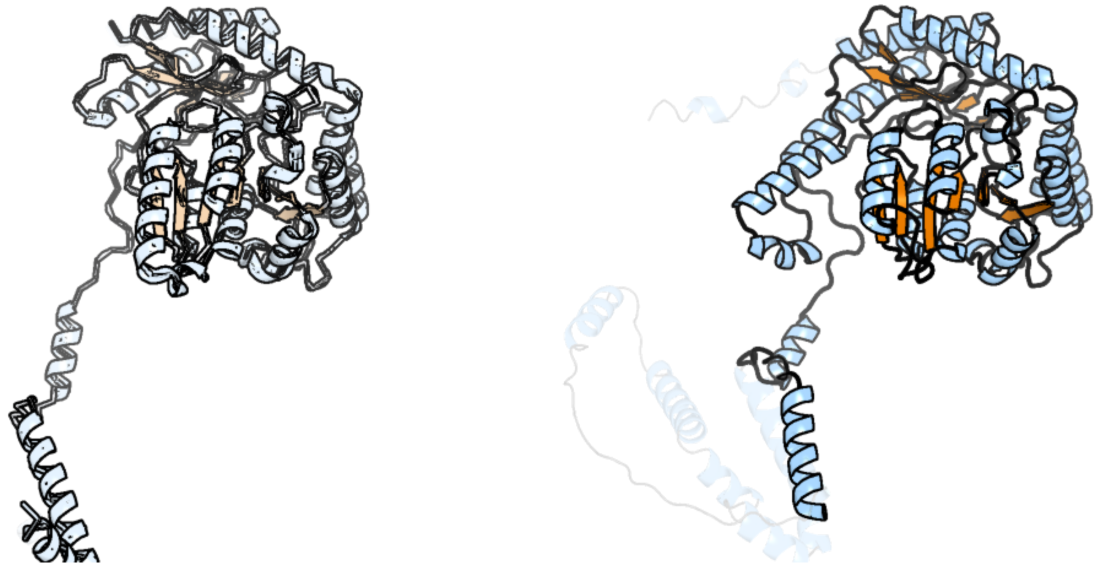

Figure 35: left: reference structure of 7k0m chain A. right: predicted structure of kaa0160201, unaligned sequences are shown as transparent

#### kaa0160314

- Sequence-based annotation for kaa0160314 (predicted in Sulfonolipids from hits to PF00155, UniRef90_A0A7Z8YEE0, UniRef90_D4IKV7) is hypothetical protein FNF28_05510 [Cafeteria roenbergensis]
- Best hit was 7v5i chain B: 2-amino-3-ketobutyrate coenzyme A ligase

| target | prob | fident | alnlen | evalue | theader |
|---|---|---|---|---|---|
| 7v5i-assembly1.cif.gz_B | 1 | 0.572 | 395 | 2.783e-58 | Structural insights into the substrate selectivity of acyl-CoA transferase |
| 1fc4-assembly1.cif.gz_B | 1 | 0.569 | 395 | 9.158e-58 | 2-AMINO-3-KETOBUTYRATE COA LIGASE |
| 7bxp-assembly1.cif.gz_B | 1 | 0.548 | 401 | 1.809e-57 | 2-amino-3-ketobutyrate CoA ligase from Cupriavidus necator |

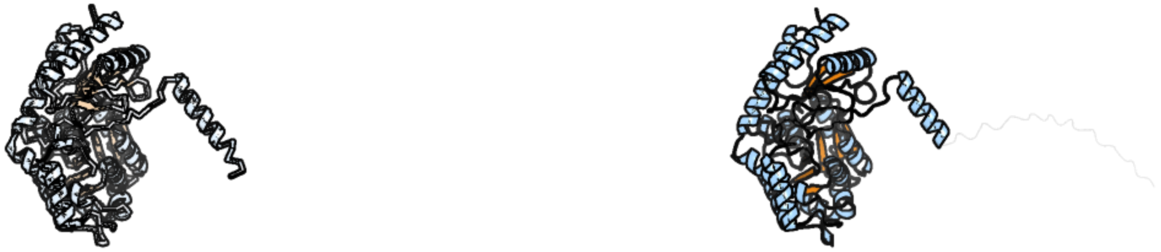

Figure 36: left: reference structure of 7v5i chain B. right: predicted structure of kaa0160314, unaligned sequences are shown as transparent

#### kaa0160603

- Sequence-based annotation for kaa0160603 (predicted in DGTS from hits to PF13649) is hypothetical protein FNF28_05401 [Cafeteria roenbergensis]
- Best hit was 4pip chain D: Histidine-specific methyltransferase EgtD

| target | prob | fident | alnlen | evalue | theader |
|---|---|---|---|---|---|
| 4pip-assembly4.cif.gz_D | 1 | 0.117 | 384 | 4.376e-08 | Engineered EgtD variant EgtD-M252V,E282A in complex with tryptophan and SAH |
| 1y8c-assembly1.cif.gz_A | 1 | 0.096 | 279 | 1.084e-07 | Crystal structure of a S-adenosylmethionine-dependent methyltransferase from Clostridium acetobutylicum ATCC 824 |
| 7sf5-assembly2.cif.gz_B | 1 | 0.135 | 400 | 4.229e-07 | M. tb EgtD in complex with HD3 |

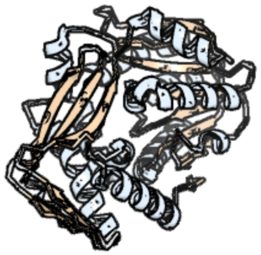

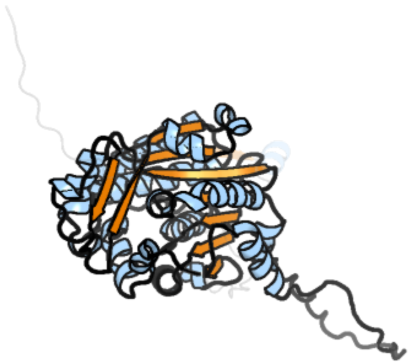

Figure 37: left: reference structure of 4pip chain D. right: predicted structure of kaa0160603, unaligned sequences are shown as transparent

#### kaa0160875

- Sequence-based annotation for kaa0160875 (predicted in Fu-FA from hits to PF01593, UniRef50_Q5LTD2) is hypothetical protein FNF28_05297 [Cafeteria roenbergensis]
- Best hit was 4ivm chain B: Protoporphyrinogen oxidase

| target | prob | fident | alnlen | evalue | theader |
|---|---|---|---|---|---|
| 4ivm-assembly1.cif.gz_B | 1 | 0.277 | 548 | 1.236e-40 | Structure of human protoporphyrinogen IX oxidase(R59G) |
| 3nks-assembly1.cif.gz_A | 1 | 0.278 | 546 | 9.346e-40 | Structure of human protoporphyrinogen IX oxidase |
| 4ivo-assembly1.cif.gz_B | 1 | 0.274 | 551 | 2.172e-39 | Structure of human protoporphyrinogen IX oxidase(R59Q) |

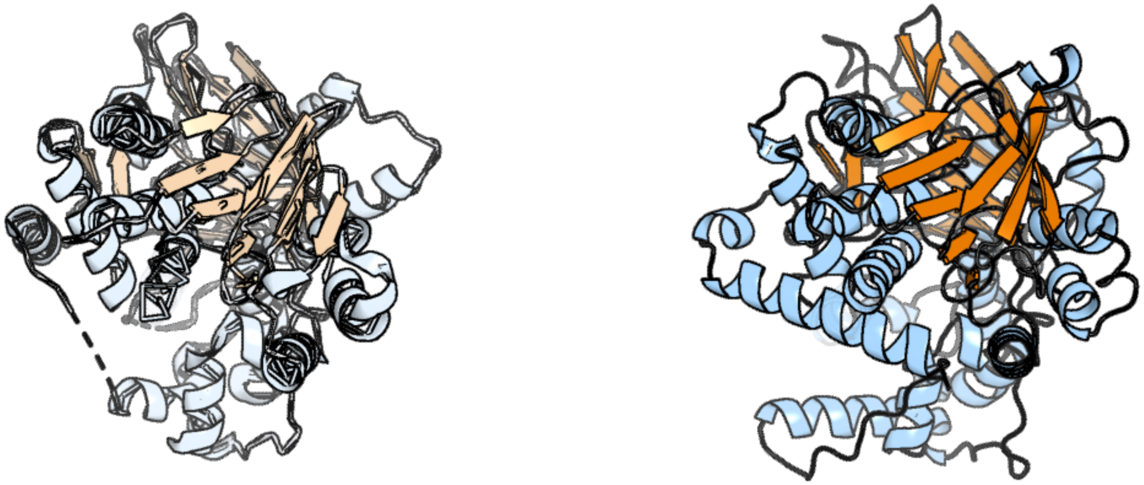

Figure 38: left: reference structure of 4ivm chain B. right: predicted structure of kaa0160875, unaligned sequences are shown as transparent

#### kaa0161056

- Sequence-based annotation for kaa0161056 (predicted in Sulfonolipids from hits to PF00155) is hypothetical protein FNF28_05204 [Cafeteria roenbergensis]
- Best hit was 1ibj chain A: CYSTATHIONINE BETA-LYASE

| target | prob | fident | alnlen | evalue | theader |
|---|---|---|---|---|---|
| 1ibj-assembly1.cif.gz_A | 1 | 0.392 | 403 | 3.358e-37 | Crystal structure of cystathionine beta-lyase from Arabidopsis thaliana |
| 6k1l-assembly1.cif.gz_B | 1 | 0.317 | 406 | 1.424e-33 | E53A mutant of a putative cystathionine gamma-lyase |
| 6k1m-assembly1.cif.gz_C | 1 | 0.322 | 403 | 2.162e-33 | Engineered form of a putative cystathionine gamma-lyase |

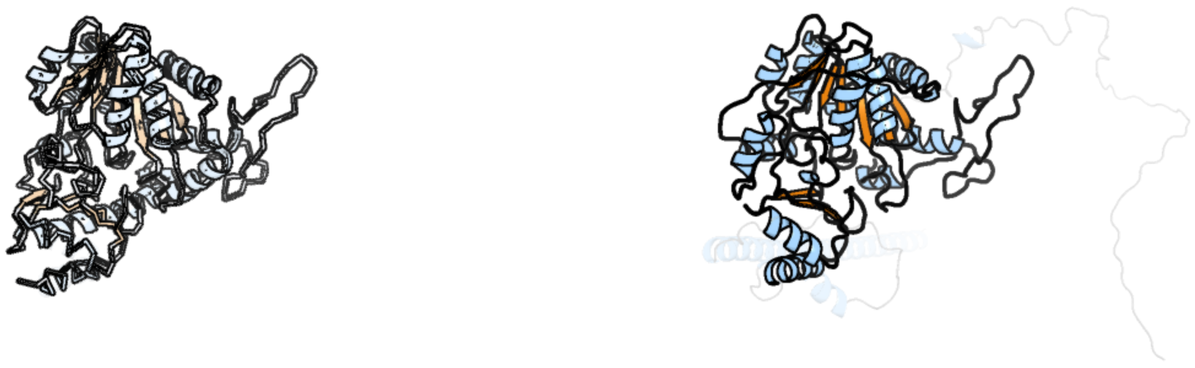

Figure 39: left: reference structure of 1ibj chain A. right: predicted structure of kaa0161056, unaligned sequences are shown as transparent

#### kaa0161448

- Sequence-based annotation for kaa0161448 (predicted in Sulfonolipids from hits to PF00291) is hypothetical protein FNF28_05040 [Cafeteria roenbergensis]
- Best hit was 2gn2 chain A: Threonine dehydratase catabolic

| target | prob | fident | alnlen | evalue | theader |
|---|---|---|---|---|---|
| 2gn2-assembly1.cif.gz_A | 1 | 0.396 | 318 | 6.634e-37 | Crystal structure of tetrameric biodegradative threonine deaminase (TdcB) from Salmonella typhimurium in complex with CMP at 2.5A resolution (Hexagonal form) |
| 1tdj-assembly1.cif.gz_A-2 | 1 | 0.317 | 410 | 8.236e-32 | THREONINE DEAMINASE (BIOSYNTHETIC) FROM E. COLI |
| 3iau-assembly2.cif.gz_B | 1 | 0.32 | 350 | 1.78e-31 | The structure of the processed form of threonine deaminase isoform 2 from Solanum lycopersicum |

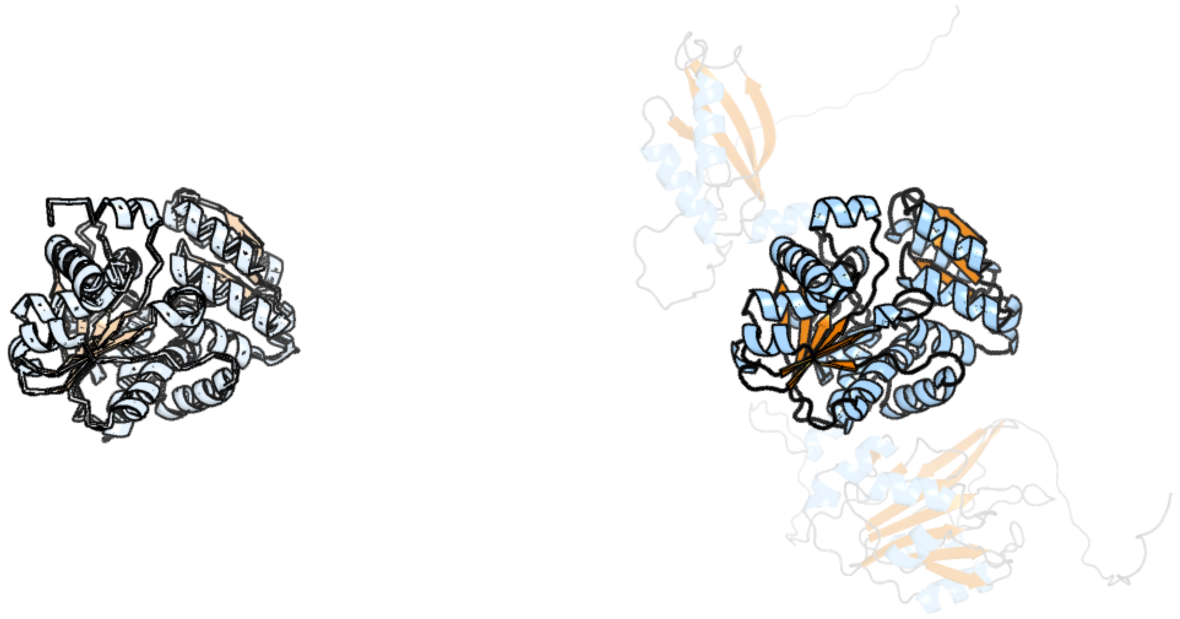

Figure 40: left: reference structure of 2gn2 chain A. right: predicted structure of kaa0161448, unaligned sequences are shown as transparent

#### kaa0161545

- Sequence-based annotation for kaa0161545 (predicted in Fu-FA from hits to PF06966) is hypothetical protein FNF28_05002 [Cafeteria roenbergensis]
- Best hit was 7c83 chain A: 3-oxo-5-alpha-steroid 4-dehydrogenase

| target | prob | fident | alnlen | evalue | theader |
|---|---|---|---|---|---|
| 7c83-assembly1.cif.gz_A | 1 | 0.408 | 262 | 1.456e-18 | Crystal structure of an integral membrane steroid 5-alpha-reductase PbSRD5A |
| 7bw1-assembly1.cif.gz_A | 1 | 0.4 | 257 | 3.93e-18 | Crystal structure of Steroid 5-alpha-reductase 2 in complex with Finasteride |
| 4quv-assembly3.cif.gz_B | 1 | 0.178 | 241 | 0.007361 | Structure of an integral membrane delta(14)-sterol reductase |

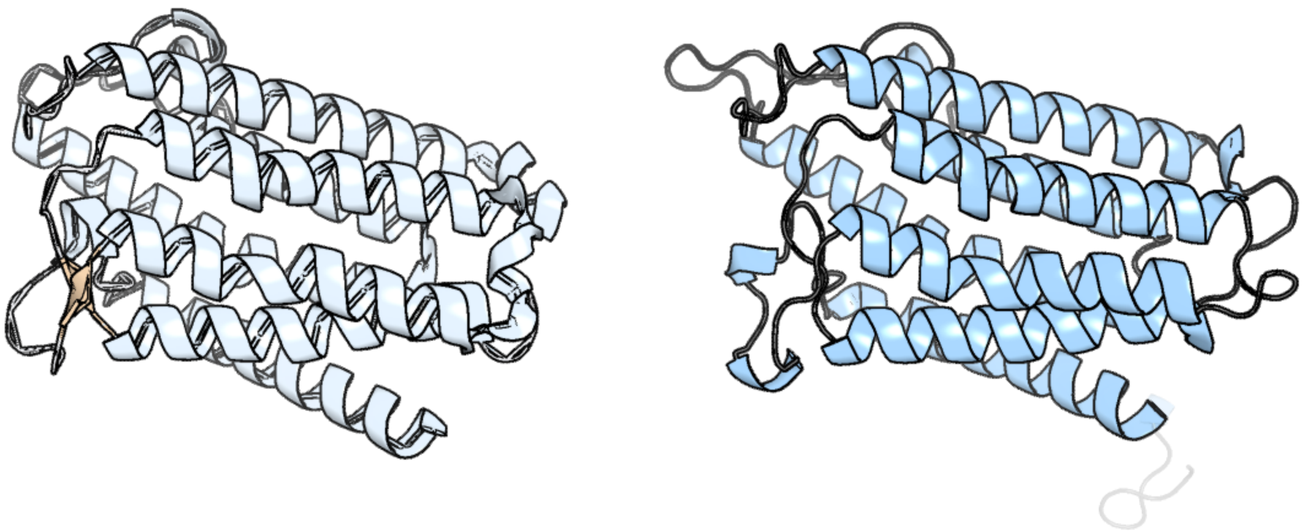

Figure 41: left: reference structure of 7c83 chain A. right: predicted structure of kaa0161545, unaligned sequences are shown as transparent

### kaa0161565

- Sequence-based annotation for kaa0161565 (predicted in DGTS from hits to PF13649) is hypothetical protein FNF28_05021 [Cafeteria roenbergensis]
- Best hit was 4hg2 chain A: Methyltransferase type 11

| target | prob | fident | alnlen | evalue | theader |
|---|---|---|---|---|---|
| 4hg2-assembly1.cif.gz_A | 1 | 0.3 | 300 | 1.475e-19 | The Structure of a Putative Type II Methyltransferase from Anaeromyxobacter dehalogenans. |
| 3g5t-assembly1.cif.gz_A | 1 | 0.195 | 342 | 2.713e-17 | Crystal structure of trans-aconitate 3-methyltransferase from yeast |
| 2yqz-assembly2.cif.gz_B | 1 | 0.195 | 343 | 3.746e-12 | Crystal Structure of Hypothetical Methyltransferase TTHA0223 from Thermus thermophilus HB8 complexed with S-adenosylmethionine |

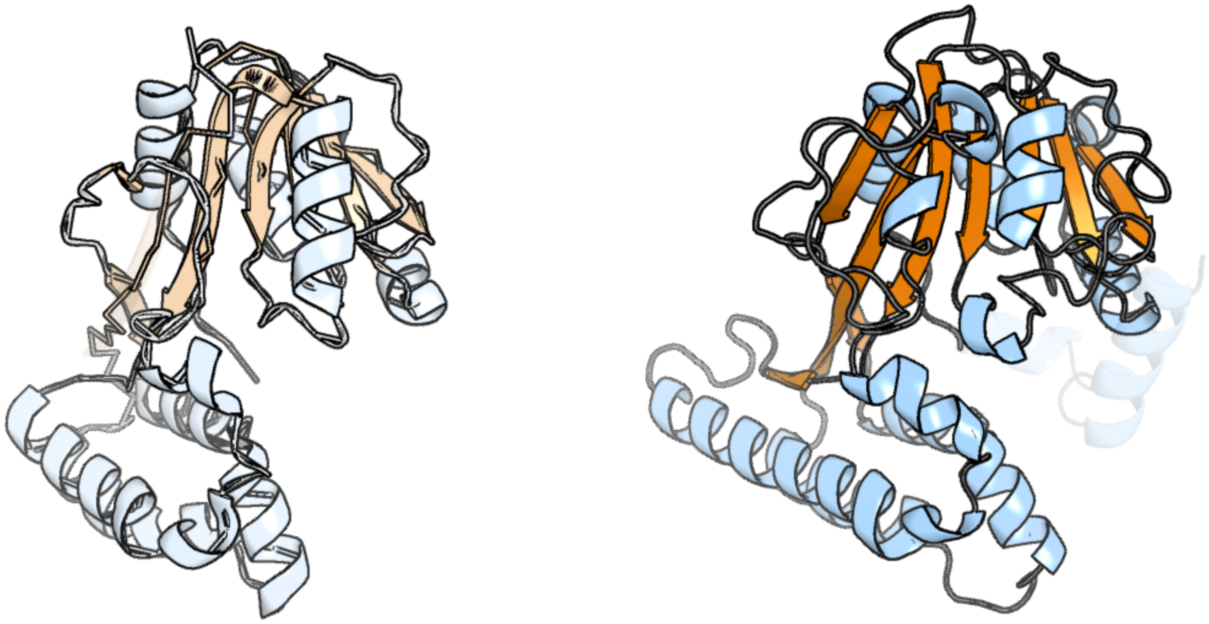

Figure 42: left: reference structure of 4hg2 chain A. right: predicted structure of kaa0161565, unaligned sequences are shown as transparent

### kaa0161568

- Sequence-based annotation for kaa0161568 (predicted in Fu-FA from hits to PF06966) is hypothetical protein FNF28_05024 [Cafeteria roenbergensis]
- Best hit was 4quv chain B: Delta(14)-sterol reductase

| target | prob | fident | alnlen | evalue | theader |
|---|---|---|---|---|---|
| 4quv-assembly3.cif.gz_B | 1 | 0.323 | 454 | 1.259e-23 | Structure of an integral membrane delta(14)-sterol reductase |
| 4quv-assembly3.cif.gz_A | 1 | 0.334 | 451 | 3.246e-23 | Structure of an integral membrane delta(14)-sterol reductase |
| 7bw1-assembly1.cif.gz_A | 0.984 | 0.168 | 321 | 0.06064 | Crystal structure of Steroid 5-alpha-reductase 2 in complex with Finasteride |

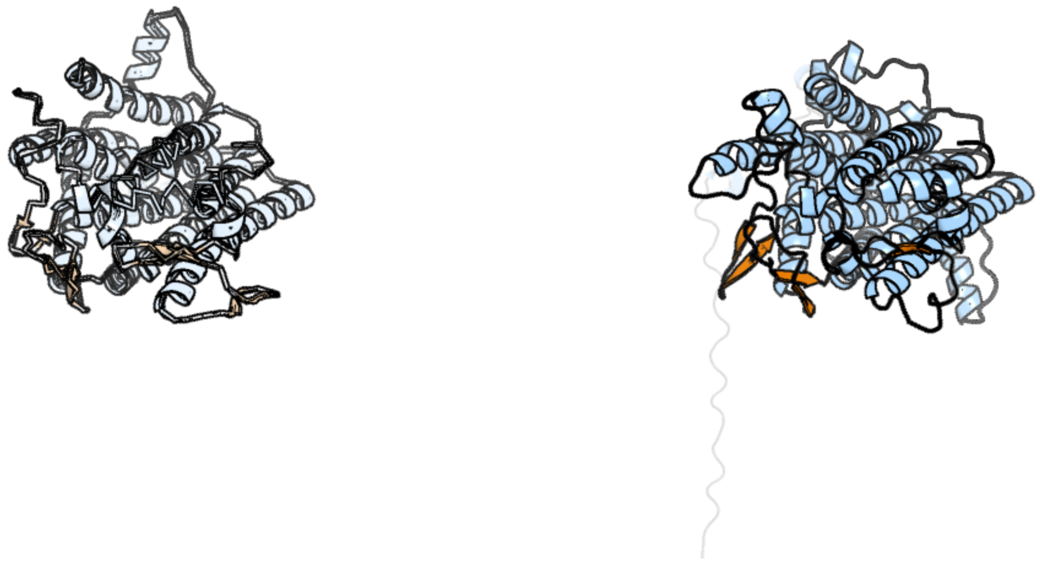

Figure 43: left: reference structure of 4quv chain B. right: predicted structure of kaa0161568, unaligned sequences are shown as transparent

#### kaa0161570

- Sequence-based annotation for kaa0161570 (predicted in Sulfonolipids from hits to PF00291, UniRef50_A0A2D9B818) is hypothetical protein FNF28_05026 [Cafeteria roenbergensis]
- Best hit was 4coo chain A: CYSTATHIONINE BETA-SYNTHASE

| target | prob | fident | alnlen | evalue | theader |
|---|---|---|---|---|---|
| 4coo-assembly1.cif.gz_A | 1 | 0.58 | 508 | 4.837e-73 | Crystal structure of human cystathionine beta-synthase (delta516-525) at 2.0 angstrom resolution |
| 4l28-assembly2.cif.gz_B | 1 | 0.581 | 507 | 2.159e-72 | Crystal structure of delta516-525 human cystathionine beta-synthase D444N mutant containing C-terminal 6xHis tag |
| 7qgt-assembly1.cif.gz_B | 1 | 0.58 | 508 | 4.699e-72 | Crystal structure of human cystathionine beta-synthase (delta516-525) in complex with AOAA. |

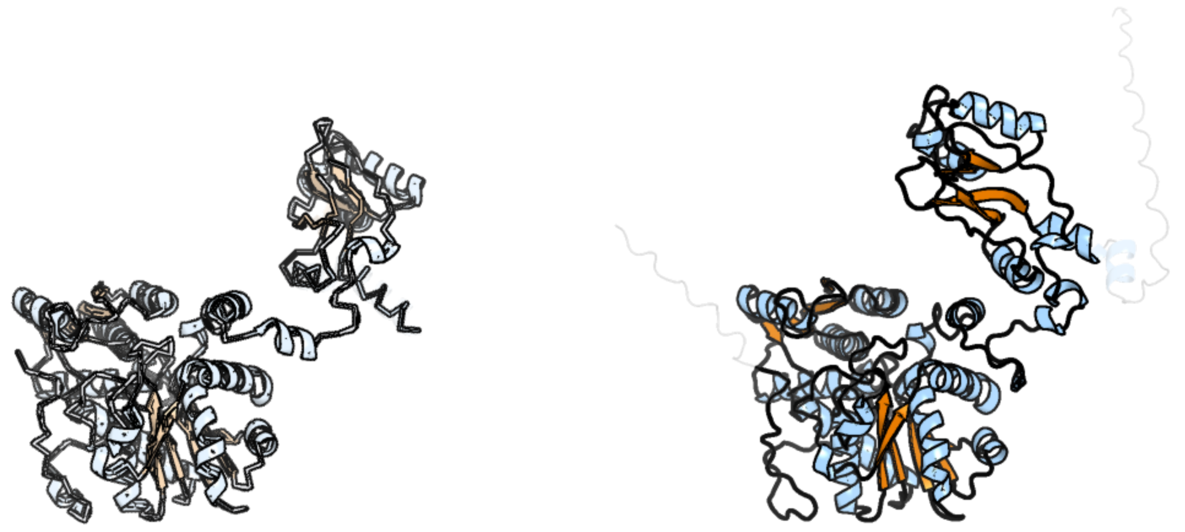

Figure 44: left: reference structure of 4coo chain A. right: predicted structure of kaa0161570, unaligned sequences are shown as transparent

### kaa0161571

- Sequence-based annotation for kaa0161571 (predicted in Fu-FA from hits to PF02353) is hypothetical protein FNF28_05027 [Cafeteria roenbergensis]
- Best hit was 6p3o chain A: Tetrahydroprotoberberine N-methyltransferase

| target | prob | fident | alnlen | evalue | theader |
|---|---|---|---|---|---|
| 6p3o-assembly1.cif.gz_A-2 | 1 | 0.326 | 153 | 7.048e-11 | Tetrahydroprotoberberine N-methyltransferase in complex with (S)-cis-N-methylstylopine and S-adenosylhomocysteine |
| 6gkz-assembly2.cif.gz_C | 1 | 0.379 | 153 | 9.583e-11 | Crystal structure of Coclaurine N-Methyltransferase (CNMT) bound to N-methylheliamine and SAH |
| 6gky-assembly1.cif.gz_A-2 | 1 | 0.389 | 154 | 9.583e-11 | Crystal structure of Coclaurine N-Methyltransferase (CNMT) bound to N-methylheliamine and SAH |

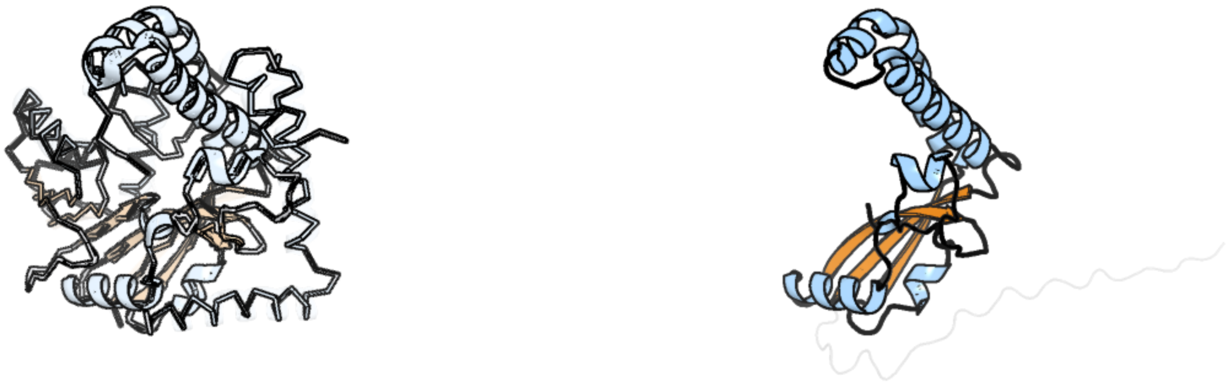

Figure 45: left: reference structure of 6p3o chain A. right: predicted structure of kaa0161571, unaligned sequences are shown as transparent

#### kaa0161746

- Sequence-based annotation for kaa0161746 (predicted in DGTS from hits to PF13649) is hypothetical protein FNF28_04933 [Cafeteria roenbergensis]
- Best hit was 3mer chain B: Slr1183 protein

| target | prob | fident | alnlen | evalue | theader |
|---|---|---|---|---|---|
| 3mer-assembly2.cif.gz_B | 1 | 0.255 | 196 | 2.498e-13 | Crystal Structure of the methyltransferase Slr1183 from Synechocystis sp. PCC 6803, Northeast Structural Genomics Consortium Target SgR145 |
| 6mro-assembly1.cif.gz_A | 1 | 0.202 | 222 | 2.258e-12 | Crystal structure of methyl transferase from Methanosarcina acetivorans at 1.6 Angstroms resolution, Northeast Structural Genomics Consortium (NESG) Target MvR53. |
| 4nec-assembly4.cif.gz_D | 1 | 0.187 | 235 | 5.383e-12 | Conversion of a Disulfide Bond into a Thioacetal Group during Echinomycin Biosynthesis |

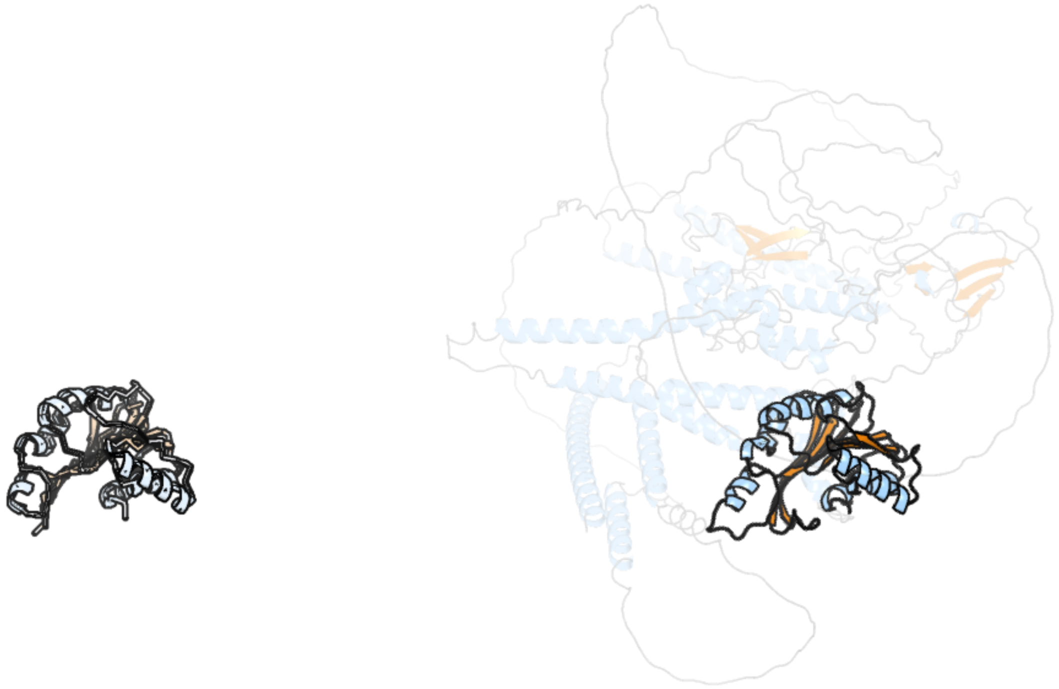

Figure 46: left: reference structure of 3mer chain B. right: predicted structure of kaa0161746, unaligned sequences are shown as transparent

### kaa0162082

- Sequence-based annotation for kaa0162082 (predicted in DGTS from hits to PF13649) is hypothetical protein FNF28_04828 [Cafeteria roenbergensis]
- Best hit was 5zq8 chain B: Uncharacterized RNA methyltransferase SP_1029

| target | prob | fident | alnlen | evalue | theader |
|---|---|---|---|---|---|
| 5zq8-assembly1.cif.gz_B | 1 | 0.165 | 556 | 7.059e-23 | Crystal structure of spRlmCD with U747 stemloop RNA |
| 5xj1-assembly1.cif.gz_A | 1 | 0.173 | 564 | 2.817e-22 | Crystal structure of spRlmCD |
| 2jjq-assembly1.cif.gz_A | 1 | 0.172 | 563 | 4.025e-16 | The crystal structure of Pyrococcus abyssi tRNA (uracil-54, C5)- methyltransferase in complex with S-adenosyl-L-homocysteine |

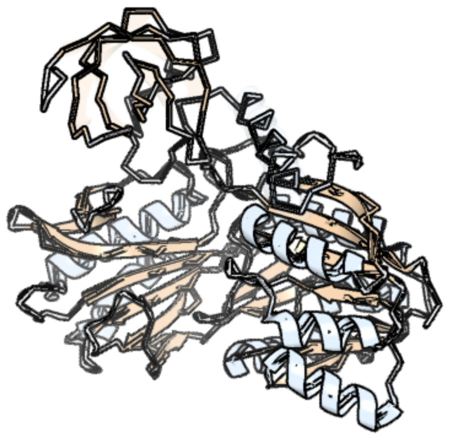

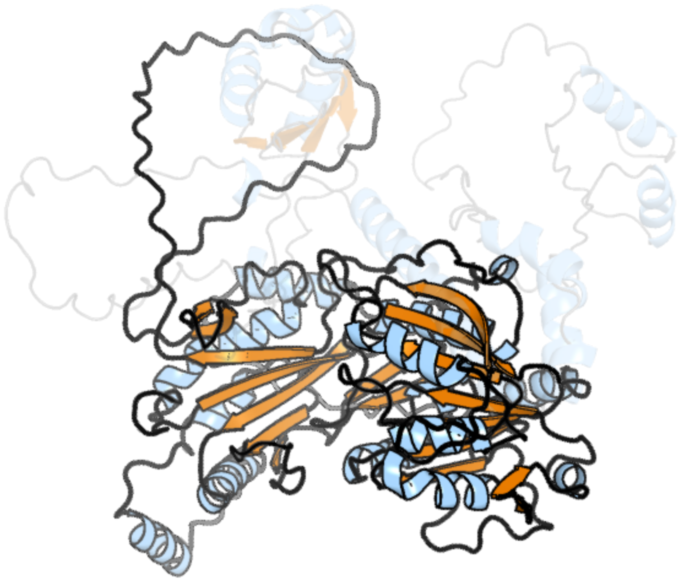

Figure 47: left: reference structure of 5zq8 chain B. right: predicted structure of kaa0162082, unaligned sequences are shown as transparent

#### kaa0162384

- Sequence-based annotation for kaa0162384 (predicted in Sulfonolipids from hits to PF00155) is hypothetical protein FNF28_04721 [Cafeteria roenbergensis]
- Best hit was 6ovg chain E: Cystathionine gamma-lyase

| target | prob | fident | alnlen | evalue | theader |
|---|---|---|---|---|---|
| 6ovg-assembly2.cif.gz_E | 1 | 0.576 | 406 | 1.454e-53 | L-Methionine Depletion with an Engineered Human Enzyme Disrupts Prostate Cancer Metabolism |
| 5tsu-assembly1.cif.gz_A | 1 | 0.562 | 407 | 3.975e-53 | Active conformation for Engineered human cystathionine gamma lyase (E59N, R119L, E339V) to depleting methionine |
| 5eig-assembly2.cif.gz_G | 1 | 0.581 | 394 | 8.218e-53 | Engineered human cystathionine gamma lyase (E59T, E339V) to deplet cysteine |

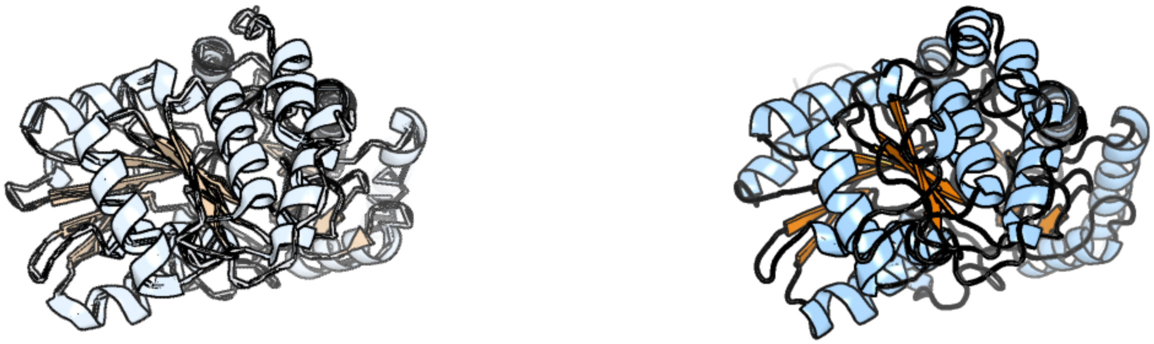

Figure 48: left: reference structure of 6ovg chain E. right: predicted structure of kaa0162384, unaligned sequences are shown as transparent

#### kaa0162392

- Sequence-based annotation for kaa0162392 (predicted in Sulfonolipids from hits to PF00106, UniRef50_A0A383U0M1) is hypothetical protein FNF28_04729 [Cafeteria roenbergensis]
- Best hit was 4fc7 chain D: Peroxisomal 2,4-dienoyl-CoA reductase

| target | prob | fident | alnlen | evalue | theader |
|---|---|---|---|---|---|
| 4fc7-assembly1.cif.gz_D | 1 | 0.541 | 266 | 6.941e-37 | Studies on DCR shed new light on peroxisomal beta-oxidation: Crystal structure of the ternary complex of pDCR |
| 1yxm-assembly1.cif.gz_A | 1 | 0.402 | 266 | 1.98e-30 | Crystal structure of peroxisomal trans 2-enoyl CoA reductase |
| 1yxm-assembly1.cif.gz_B | 1 | 0.4 | 265 | 2.368e-30 | Crystal structure of peroxisomal trans 2-enoyl CoA reductase |

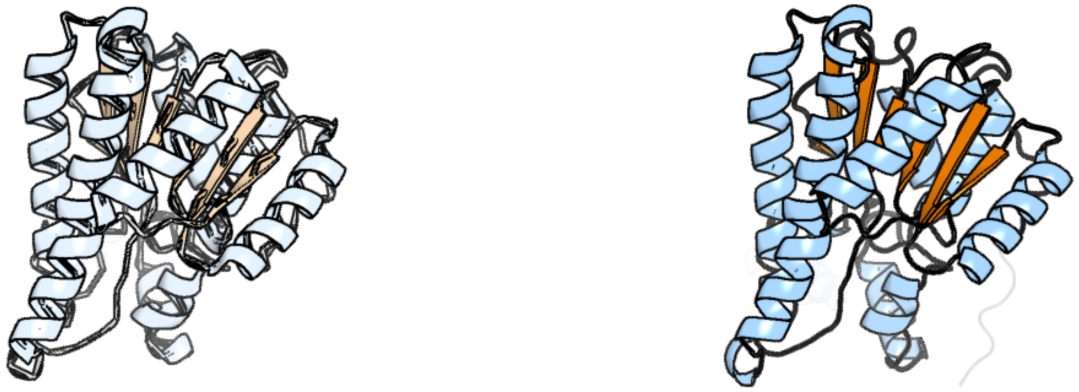

Figure 49: left: reference structure of 4fc7 chain D. right: predicted structure of kaa0162392, unaligned sequences are shown as transparent

#### kaa0162523

- Sequence-based annotation for kaa0162523 (predicted in PE_PMME from hits to PF01066) is hypothetical protein FNF28_04697 [Cafeteria roenbergensis]
- No significant structural hit found

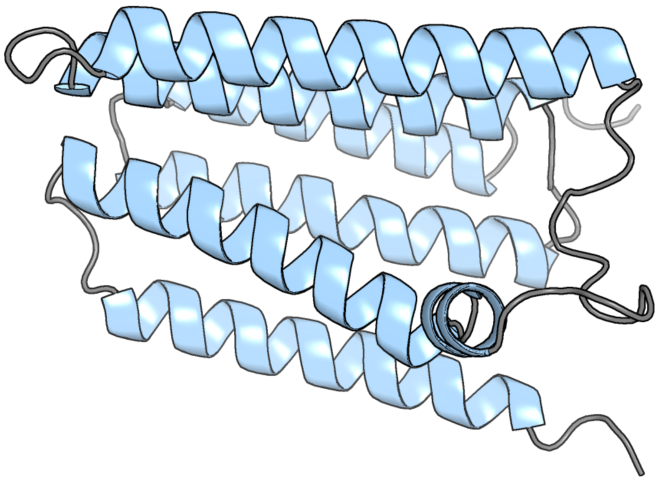

Figure 50: predicted structure of kaa0162523

#### kaa0162542

- Sequence-based annotation for kaa0162542 (predicted in Sulfonolipids from hits to PF00155, UniRef90_A0A7Z8YEE0, UniRef90_D4IKV7) is hypothetical protein FNF28_04636 [Cafeteria roenbergensis]
- Best hit was 7yjo chain B: Long chain base biosynthesis protein 2a

| target | prob | fident | alnlen | evalue | theader |
|---|---|---|---|---|---|
| 7yjo-assembly1.cif.gz_B | 1 | 0.556 | 469 | 3.142e-57 | Cryo-EM structure of the monomeric atSPT-ORM1 (LCB2a-deltaN5) complex |
| 7k0i-assembly1.cif.gz_B | 1 | 0.533 | 484 | 1.456e-56 | Human serine palmitoyltransferase complex SPTLC1/SPLTC2/ssSPTa |
| 7yjm-assembly1.cif.gz_B | 1 | 0.553 | 470 | 3.976e-56 | Cryo-EM structure of the monomeric atSPT-ORM1 complex |

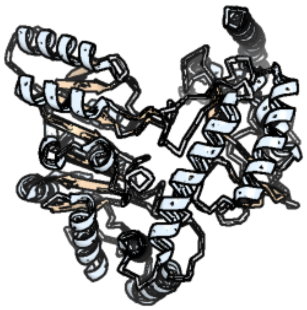

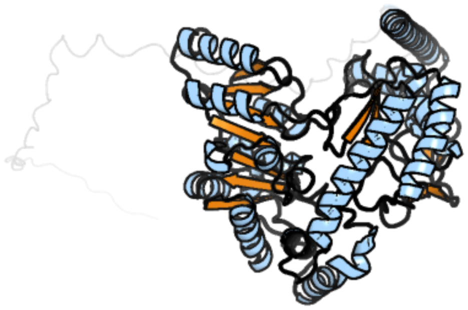

Figure 51: left: reference structure of 7yjo chain B. right: predicted structure of kaa0162542, unaligned sequences are shown as transparent

#### kaa0162677

- Sequence-based annotation for kaa0162677 (predicted in DGTS from hits to PF13649) is hypothetical protein FNF28_04620 [Cafeteria roenbergensis]
- Best hit was 6mro chain A: methyl transferase from Methanosarcina acetivorans

| target | prob | fident | alnlen | evalue | theader |
|---|---|---|---|---|---|
| 6mro-assembly1.cif.gz_A | 1 | 0.208 | 192 | 4.131e-13 | Crystal structure of methyl transferase from Methanosarcina acetivorans at 1.6 Angstroms resolution, Northeast Structural Genomics Consortium (NESG) Target MvR53. |
| 3sm3-assembly1.cif.gz_A-2 | 1 | 0.129 | 201 | 4.682e-13 | Crystal Structure of SAM-dependent methyltransferases Q8PUK2_METMA from Methanosarcina mazei. Northeast Structural Genomics Consortium Target MaR262. |
| 7f1e-assembly1.cif.gz_A | 1 | 0.148 | 242 | 4.682e-13 | Structure of METTL6 bound with SAM |

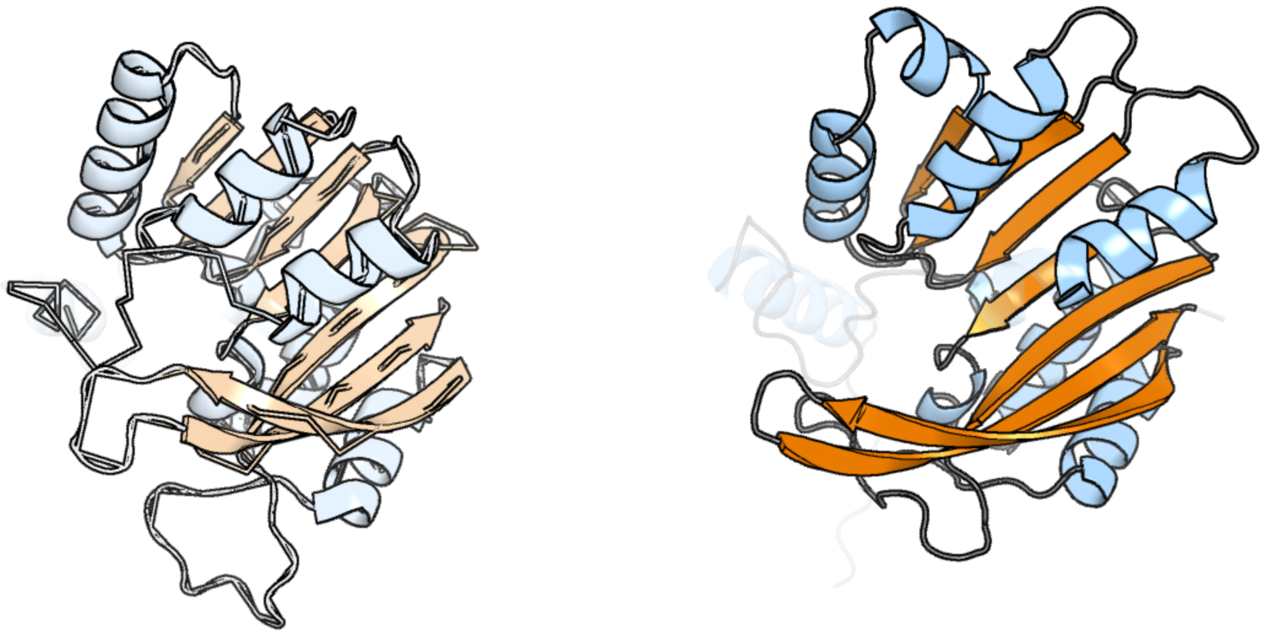

Figure 52: left: reference structure of 6mro chain A. right: predicted structure of kaa0162677, unaligned sequences are shown as transparent

#### kaa0162700

- Sequence-based annotation for kaa0162700 (predicted in Sulfonolipids from hits to PF00106, UniRef50_A0A383U0M1) is hypothetical protein FNF28_04581 [Cafeteria roenbergensis]
- Best hit was 8g9v chain C: 17-beta-hydroxysteroid dehydrogenase 13

| target | prob | fident | alnlen | evalue | theader |
|---|---|---|---|---|---|
| 8g9v-assembly2.cif.gz_C | 1 | 0.215 | 278 | 1.881e-20 | Crystal structures of 17-beta-hydroxysteroid dehydrogenase 13 |
| 3tjr-assembly1.cif.gz_A | 1 | 0.236 | 300 | 3.137e-20 | Crystal structure of a Rv0851c ortholog short chain dehydrogenase from Mycobacterium paratuberculosis |
| 8g93-assembly1.cif.gz_B | 1 | 0.221 | 293 | 3.938e-20 | Crystal structures of 17-beta-hydroxysteroid dehydrogenase 13 |

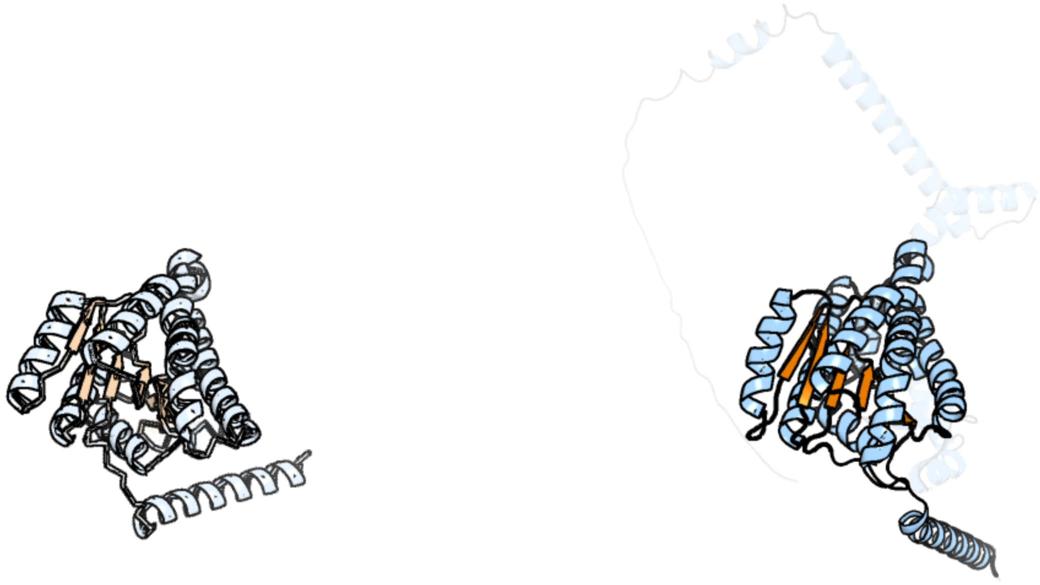

Figure 53: left: reference structure of 8g9v chain C. right: predicted structure of kaa0162700, unaligned sequences are shown as transparent

#### kaa0162783

- Sequence-based annotation for kaa0162783 (predicted in Sulfonolipids from hits to PF00106) is hypothetical protein FNF28_04540 [Cafeteria roenbergensis]
- Best hit was 4jro chain C: FabG protein

| target | prob | fident | alnlen | evalue | theader |
|---|---|---|---|---|---|
| 4jro-assembly1.cif.gz_C | 1 | 0.287 | 299 | 3.906e-19 | Crystal structure of 3-oxoacyl-[acyl-carrier protein]reductase (FabG)from Listeria monocytogenes in complex with NADP+ |
| 3ftp-assembly1.cif.gz_B | 1 | 0.296 | 300 | 5.723e-19 | Crystal structure of 3-Ketoacyl-(acyl-carrier-protein) reductase from Burkholderia pseudomallei at 2.05 A resolution |
| 2cdh-assembly1.cif.gz_G | 1 | 0.281 | 302 | 6.044e-19 | ARCHITECTURE OF THE THERMOMYCES LANUGINOSUS FUNGAL FATTY ACID SYNTHASE AT 5 ANGSTROM RESOLUTION. |

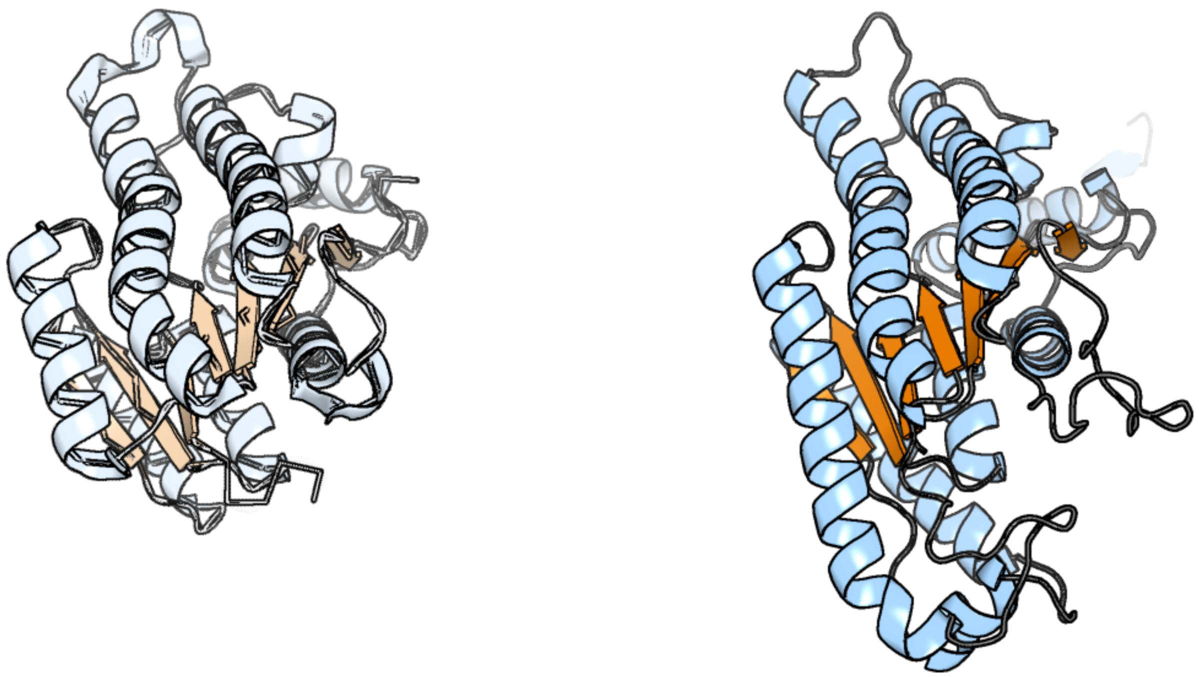

Figure 54: left: reference structure of 4jro chain C. right: predicted structure of kaa0162783, unaligned sequences are shown as transparent

#### kaa0162926

- Sequence-based annotation for kaa0162926 (predicted in DGTS from hits to PF13649) is hypothetical protein FNF28_04482 [Cafeteria roenbergensis]
- Best hit was 3ocj chain A: Putative exported protein

| target | prob | fident | alnlen | evalue | theader |
|---|---|---|---|---|---|
| 3ocj-assembly1.cif.gz_A | 1 | 0.326 | 297 | 1.422e-23 | The crystal structure of a possilbe exported protein from Bordetella parapertussis |
| 8ftv-assembly1.cif.gz_A-2 | 1 | 0.161 | 346 | 1.285e-09 | SgvM methyltransferase triple variant (M144V/F329V/T331A) with SAH and 2-oxo-4-phenylbutanoic acid |
| 8g5s-assembly1.cif.gz_B | 1 | 0.152 | 361 | 1.564e-09 | Crystal structure of apo TnmJ |

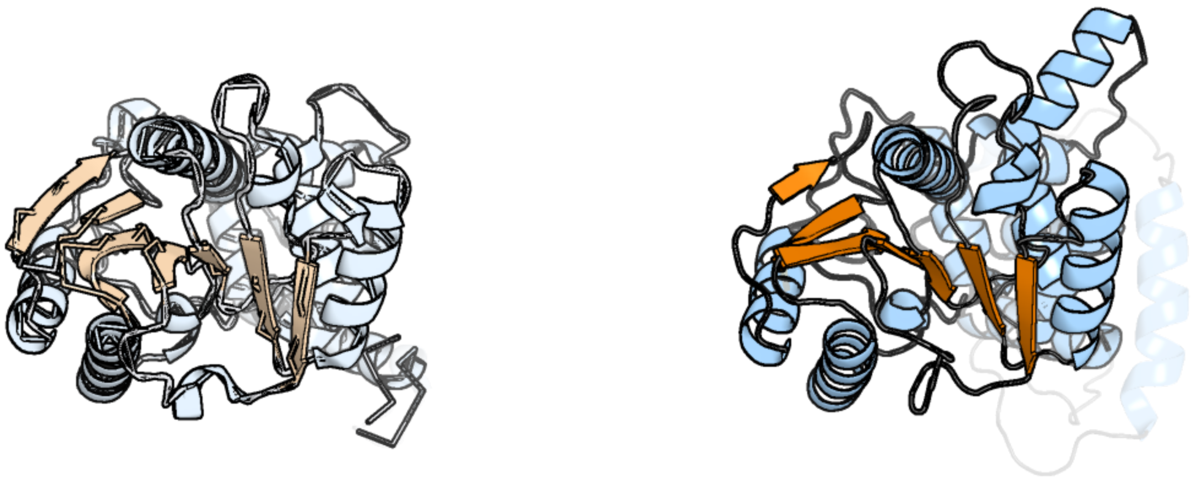

Figure 55: left: reference structure of 3ocj chain A. right: predicted structure of kaa0162926, unaligned sequences are shown as transparent

#### kaa0163236

- Sequence-based annotation for kaa0163236 (predicted in Sulfonolipids from hits to PF00155) is hypothetical protein FNF28_04386 [Cafeteria roenbergensis]
- Best hit was 7aat chain A: ASPARTATE AMINOTRANSFERASE

| target | prob | fident | alnlen | evalue | theader |
|---|---|---|---|---|---|
| 7aat-assembly1.cif.gz_A | 1 | 0.502 | 398 | 1.418e-52 | X-RAY STRUCTURE REFINEMENT AND COMPARISON OF THREE FORMS OF MITOCHONDRIAL ASPARTATE AMINOTRANSFERASE |
| 5ax8-assembly2.cif.gz_D | 1 | 0.513 | 399 | 3.38e-51 | Recombinant expression, purification and preliminary crystallographic studies of the mature form of human mitochondrial aspartate aminotransferase |
| 3pd6-assembly1.cif.gz_A | 1 | 0.511 | 399 | 4.666e-51 | Crystal structure of mouse mitochondrial aspartate aminotransferase, a newly identified kynurenine aminotransferase-IV |

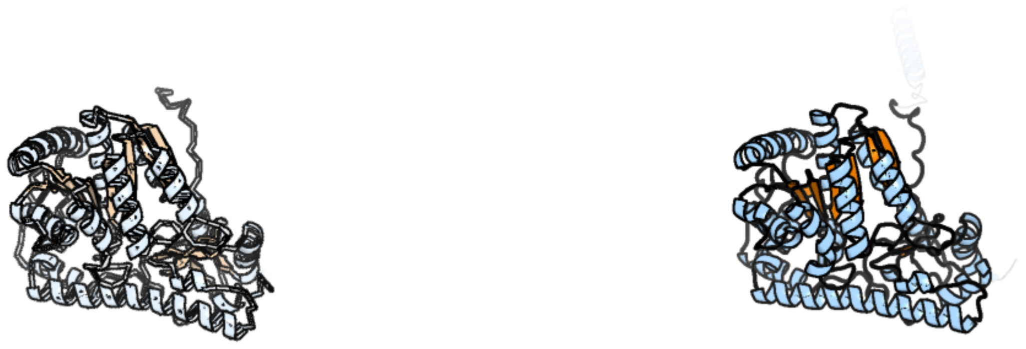

Figure 56: left: reference structure of 7aat chain A. right: predicted structure of kaa0163236, unaligned sequences are shown as transparent

#### kaa0163360

- Sequence-based annotation for kaa0163360 (predicted in Sulfonolipids from hits to PF00155) is hypothetical protein FNF28_04281 [Cafeteria roenbergensis]
- Best hit was 1jg8 chain A: L-allo-threonine aldolase

| target | prob | fident | alnlen | evalue | theader |
|---|---|---|---|---|---|
| 1jg8-assembly1.cif.gz_A | 1 | 0.421 | 346 | 1.15e-41 | Crystal Structure of Threonine Aldolase (Low-specificity) |
| 3wgc-assembly1.cif.gz_D | 1 | 0.457 | 341 | 2.077e-41 | Aeromonas jandaei L-allo-threonine aldolase H128Y/S292R double mutant |
| 3wgb-assembly1.cif.gz_C | 1 | 0.461 | 342 | 3.331e-41 | Crystal structure of aeromonas jandaei L-allo-threonine aldolase |

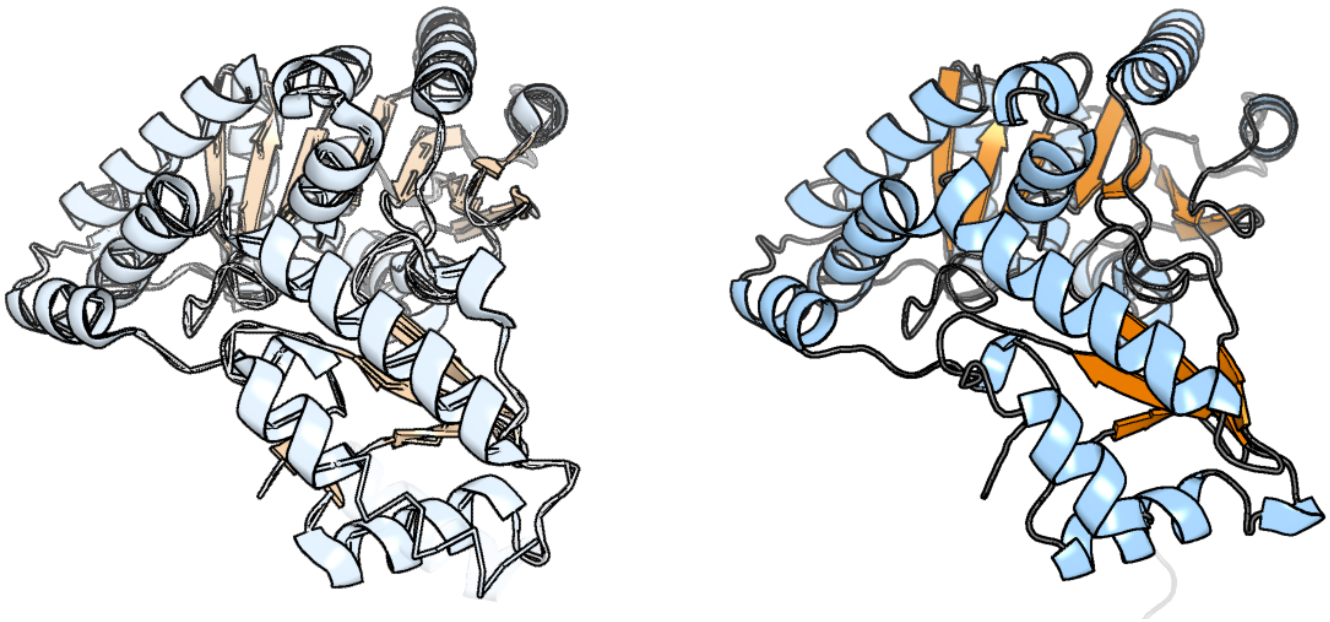

Figure 57: left: reference structure of 1jg8 chain A. right: predicted structure of kaa0163360, unaligned sequences are shown as transparent

### kaa0163518

- Sequence-based annotation for kaa0163518 (predicted in PE_PMME from hits to PF01066) is hypothetical protein FNF28_04175 [Cafeteria roenbergensis]
- Best hit was 7drk chain B: CDP-diacylglycerol–glycerol-3-phosphate 3-phosphatidyltransferase

| target | prob | fident | alnlen | evalue | theader |
|---|---|---|---|---|---|
| 7drk-assembly1.cif.gz_B | 1 | 0.211 | 203 | 9.24e-05 | Crystal structure of phosphatidylglycerol phosphate synthase in complex with cytidine diphosphate-diacylglycerol |
| 6wmv-assembly1.cif.gz_A | 0.997 | 0.168 | 208 | 0.01666 | Structure of a phosphatidylinositol-phosphate synthase (PIPS) from Mycobacterium kansasii with evidence of substrate binding |
| 6wm5-assembly1.cif.gz_A | 1 | 0.157 | 203 | 0.01922 | Structure of a phosphatidylinositol-phosphate synthase (PIPS) from Mycobacterium kansasii |

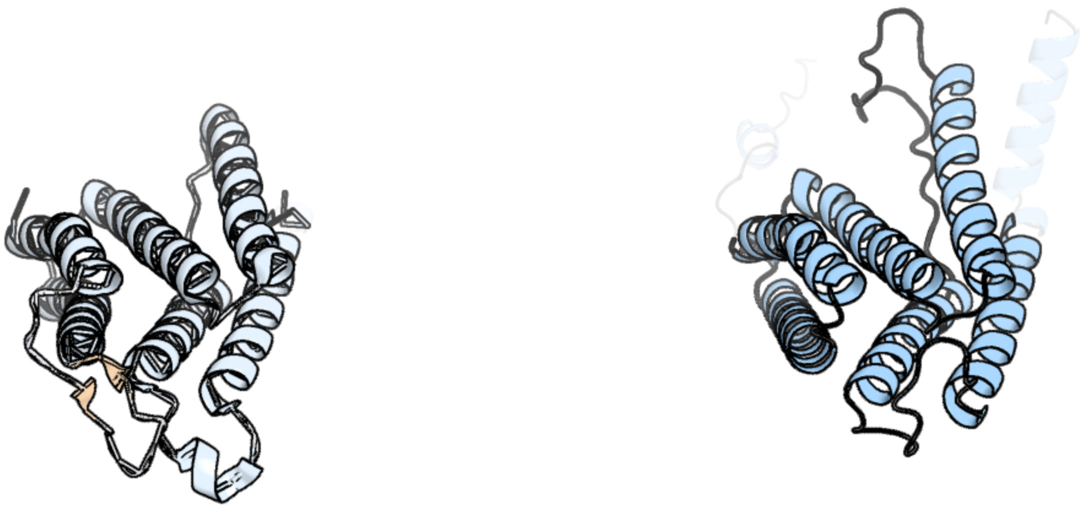

Figure 58: left: reference structure of 7drk chain B. right: predicted structure of kaa0163518, unaligned sequences are shown as transparent

#### kaa0163661

- Sequence-based annotation for kaa0163661 (predicted in Sulfonolipids from hits to PF00155) is hypothetical protein FNF28_04138 [Cafeteria roenbergensis]
- Best hit was 3tcm chain B: Alanine aminotransferase 2

| target | prob | fident | alnlen | evalue | theader |
|---|---|---|---|---|---|
| 3tcm-assembly1.cif.gz_B | 1 | 0.504 | 492 | 1.018e-58 | Crystal Structure of Alanine Aminotransferase from Hordeum vulgare |
| 3ihj-assembly1.cif.gz_A | 1 | 0.434 | 481 | 1.479e-47 | Human alanine aminotransferase 2 in complex with PLP |
| 4cvq-assembly1.cif.gz_B | 1 | 0.224 | 486 | 7.545e-31 | CRYSTAL STRUCTURE OF AN AMINOTRANSFERASE FROM ESCHERICHIA COLI AT 2. 11 ANGSTROEM RESOLUTION |

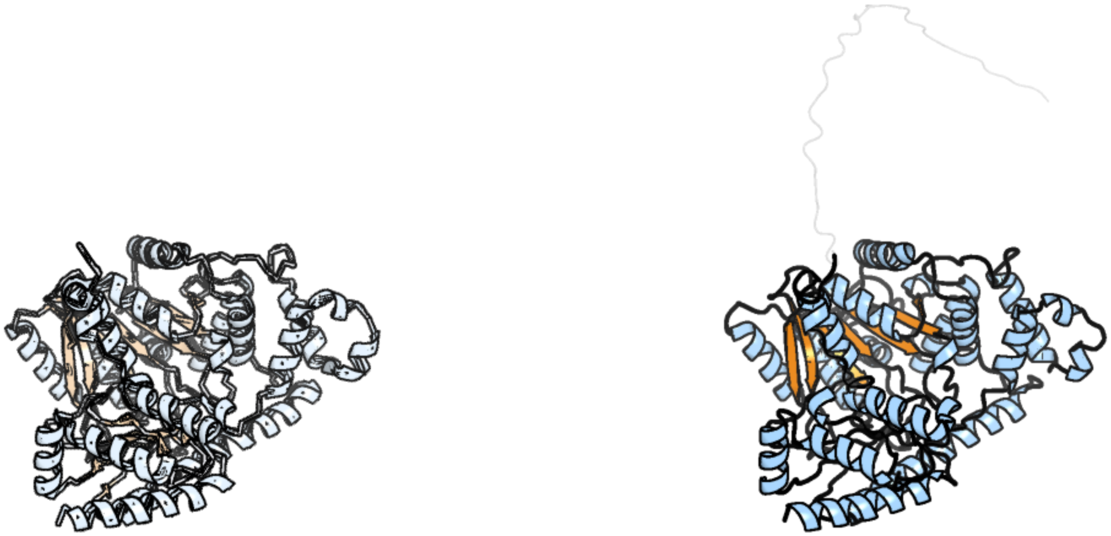

Figure 59: left: reference structure of 3tcm chain B. right: predicted structure of kaa0163661, unaligned sequences are shown as transparent

### kaa0163873

- Sequence-based annotation for kaa0163873 (predicted in DGTS from hits to PF13649) is hypothetical protein FNF28_04067 [Cafeteria roenbergensis]
- Best hit was 2pxx chain A: Uncharacterized protein MGC2408

| target | prob | fident | alnlen | evalue | theader |
|---|---|---|---|---|---|
| 2pxx-assembly1.cif.gz_A | 1 | 0.238 | 231 | 1.065e-15 | Human putative methyltransferase MGC2408 |
| 4nec-assembly3.cif.gz_C | 1 | 0.187 | 262 | 4.298e-09 | Conversion of a Disulfide Bond into a Thioacetal Group during Echinomycin Biosynthesis |
| 5m58-assembly1.cif.gz_B | 1 | 0.179 | 245 | 1.819e-08 | Crystal structure of CouO, a C-methyltransferase from Streptomyces rishiriensis |

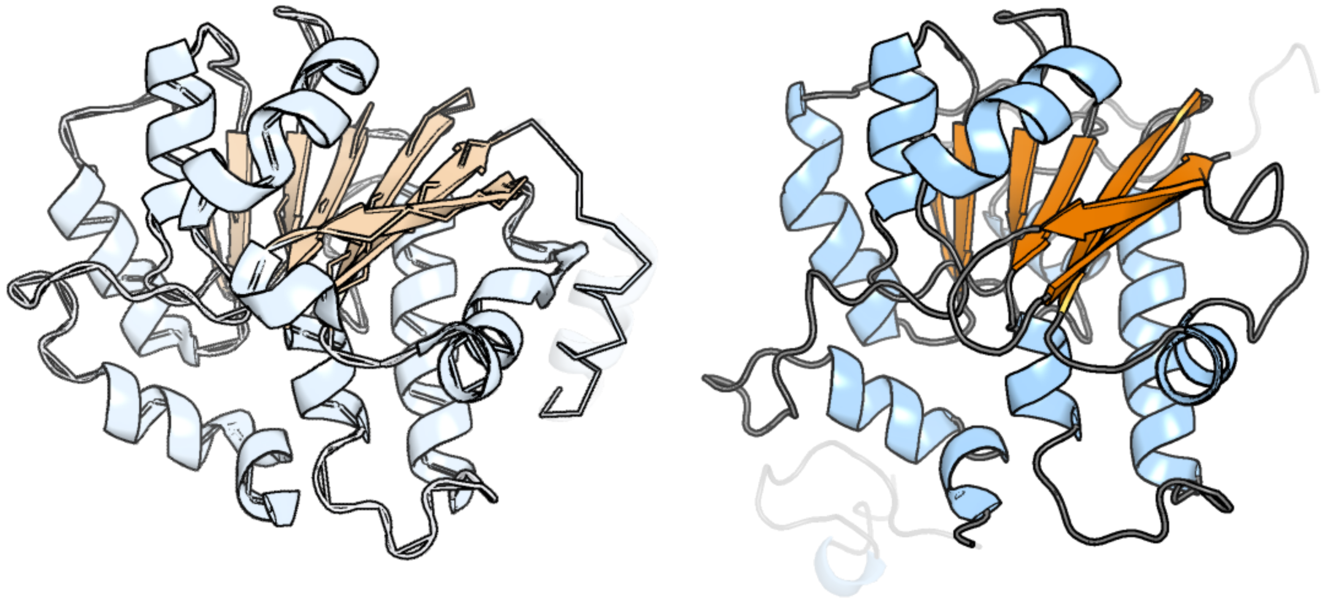

Figure 60: left: reference structure of 2pxx chain A. right: predicted structure of kaa0163873, unaligned sequences are shown as transparent

### kaa0163941

- Sequence-based annotation for kaa0163941 (predicted in Sulfonolipids from hits to PF00106, UniRef50_A0A383U0M1) is hypothetical protein FNF28_04047 [Cafeteria roenbergensis]
- Best hit was 3rd5 chain A: MYPAA.01249.C

| target | prob | fident | alnlen | evalue | theader |
|---|---|---|---|---|---|
| 3rd5-assembly1.cif.gz_A | 1 | 0.322 | 307 | 8.822e-19 | Crystal structure of a putative uncharacterized protein from Mycobacterium Paratuberculosis |
| 6l1h-assembly2.cif.gz_B | 1 | 0.241 | 307 | 2.405e-16 | Crystal structure of light-dependent protochlorophyllide oxidoreductase from Thermosynechococcus elongatus |
| 7jk9-assembly1.cif.gz_A | 1 | 0.243 | 316 | 5.532e-16 | Helical filaments of plant light-dependent protochlorophyllide oxidoreductase (LPOR) bound to NADPH, Pchlide, and membrane |

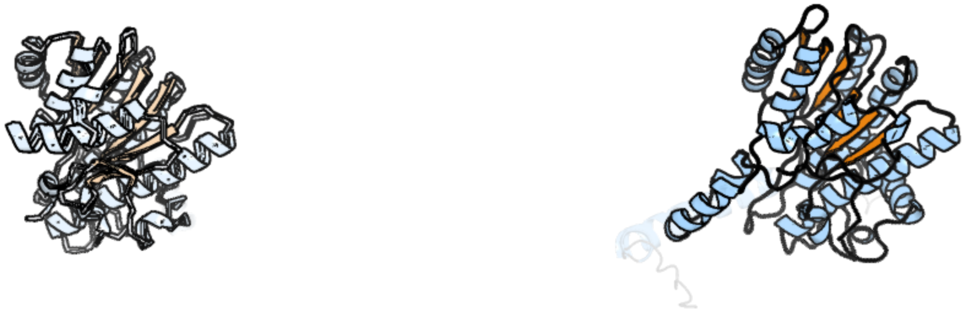

Figure 61: left: reference structure of 3rd5 chain A. right: predicted structure of kaa0163941, unaligned sequences are shown as transparent

#### kaa0163942

- Sequence-based annotation for kaa0163942 (predicted in Sulfonolipids from hits to PF00106, UniRef50_A0A383U0M1) is hypothetical protein FNF28_04048 [Cafeteria roenbergensis]
- Best hit was 7jk9 chain A: Protochlorophyllide reductase B, chloroplastic

| target | prob | fident | alnlen | evalue | theader |
|---|---|---|---|---|---|
| 7jk9-assembly1.cif.gz_A | 1 | 0.243 | 320 | 6.677e-20 | Helical filaments of plant light-dependent protochlorophyllide oxidoreductase (LPOR) bound to NADPH, Pchlide, and membrane |
| 6l1h-assembly2.cif.gz_B | 1 | 0.237 | 308 | 7.006e-19 | Crystal structure of light-dependent protochlorophyllide oxidoreductase from Thermosynechococcus elongatus |
| 3rd5-assembly1.cif.gz_A | 1 | 0.323 | 297 | 1.261e-18 | Crystal structure of a putative uncharacterized protein from Mycobacterium Paratuberculosis |

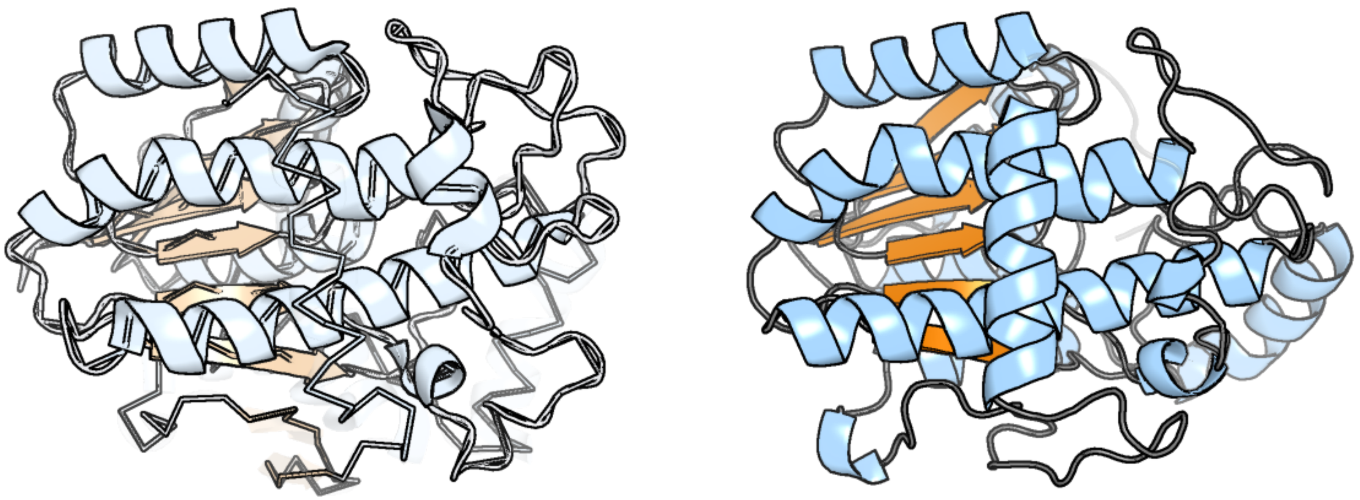

Figure 62: left: reference structure of 7jk9 chain A. right: predicted structure of kaa0163942, unaligned sequences are shown as transparent

#### kaa0164148

- Sequence-based annotation for kaa0164148 (predicted in DGTS from hits to PF00010) is hypothetical protein FNF28_03961 [Cafeteria roenbergensis]
- No significant structural hit found

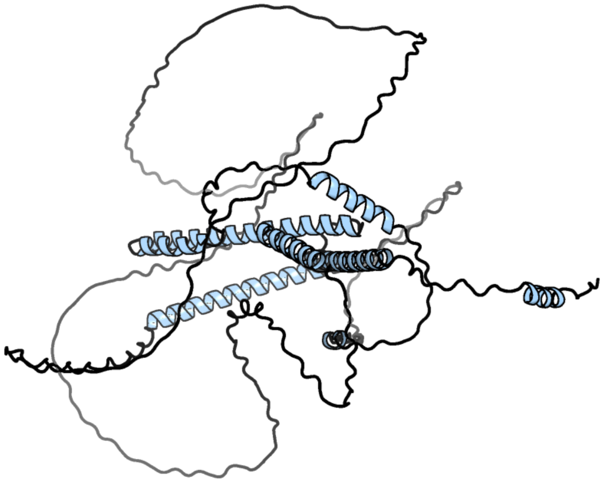

Figure 63: predicted structure of kaa0164148

### kaa0164503

- Sequence-based annotation for kaa0164503 (predicted in Sulfonolipids from hits to PF00106) is hypothetical protein FNF28_03844 [Cafeteria roenbergensis]
- Best hit was 3rd5 chain A: MYPAA.01249.C

| target | prob | fident | alnlen | evalue | theader |
|---|---|---|---|---|---|
| 3rd5-assembly1.cif.gz_A | 1 | 0.268 | 294 | 4.449e-16 | Crystal structure of a putative uncharacterized protein from Mycobacterium Paratuberculosis |
| 6l1g-assembly1.cif.gz_A | 1 | 0.203 | 315 | 1.734e-15 | Crystal structure of light-dependent protochlorophyllide oxidoreductase from Synechocystis sp. PCC 6803 |
| 6r48-assembly1.cif.gz_A | 1 | 0.216 | 309 | 1.831e-15 | Crystal structure of LPOR (Synechocystis) complexed with NADPH at 1.87A resolution. |

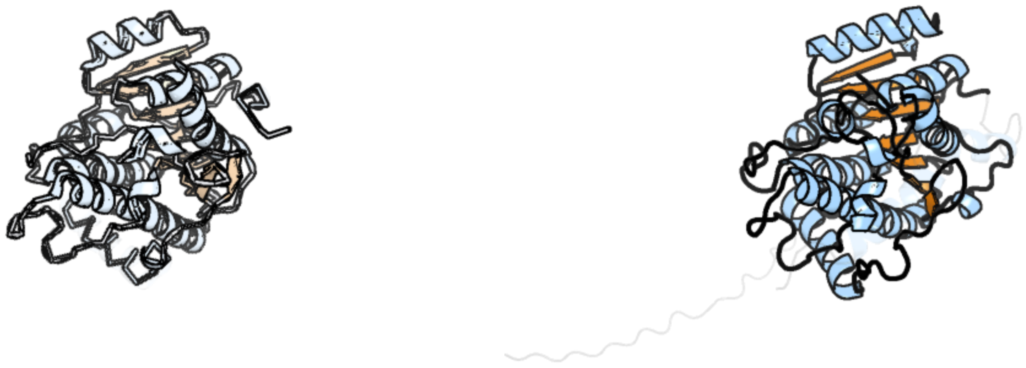

Figure 64: left: reference structure of 3rd5 chain A. right: predicted structure of kaa0164503, unaligned sequences are shown as transparent

### kaa0164550

- Sequence-based annotation for kaa0164550 (predicted in PE_PMME from hits to PF04191) is hypothetical protein FNF28_03792 [Cafeteria roenbergensis]
- No significant structural hit found

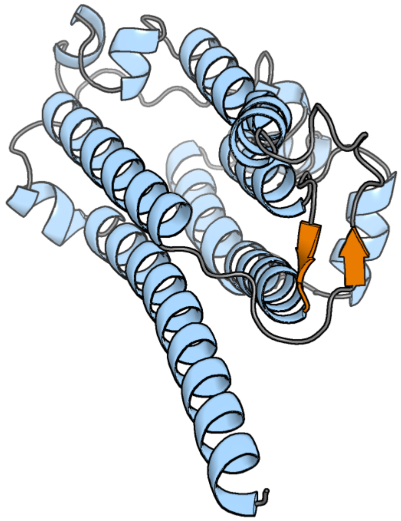

Figure 65: predicted structure of kaa0164550

### kaa0164596

- Sequence-based annotation for kaa0164596 (predicted in Fu-FA from hits to PF01593) is hypothetical protein FNF28_03755 [Cafeteria roenbergensis]
- Best hit was 3lxd chain A: FAD-dependent pyridine nucleotide-disulphide oxidoreductase

| target | prob | fident | alnlen | evalue | theader |
|---|---|---|---|---|---|
| 3lxd-assembly1.cif.gz_A | 1 | 0.312 | 439 | 8.675e-37 | Crystal Structure of Ferredoxin Reductase ArR from Novosphingobium aromaticivorans |
| 1q1r-assembly1.cif.gz_A | 1 | 0.272 | 451 | 1.293e-36 | Crystal Structure of Putidaredoxin Reductase from Pseudomonas putida |
| 3fg2-assembly1.cif.gz_P | 1 | 0.263 | 440 | 2.16e-36 | Crystal Structure of Ferredoxin Reductase for the CYP199A2 System from Rhodopseudomonas palustris |

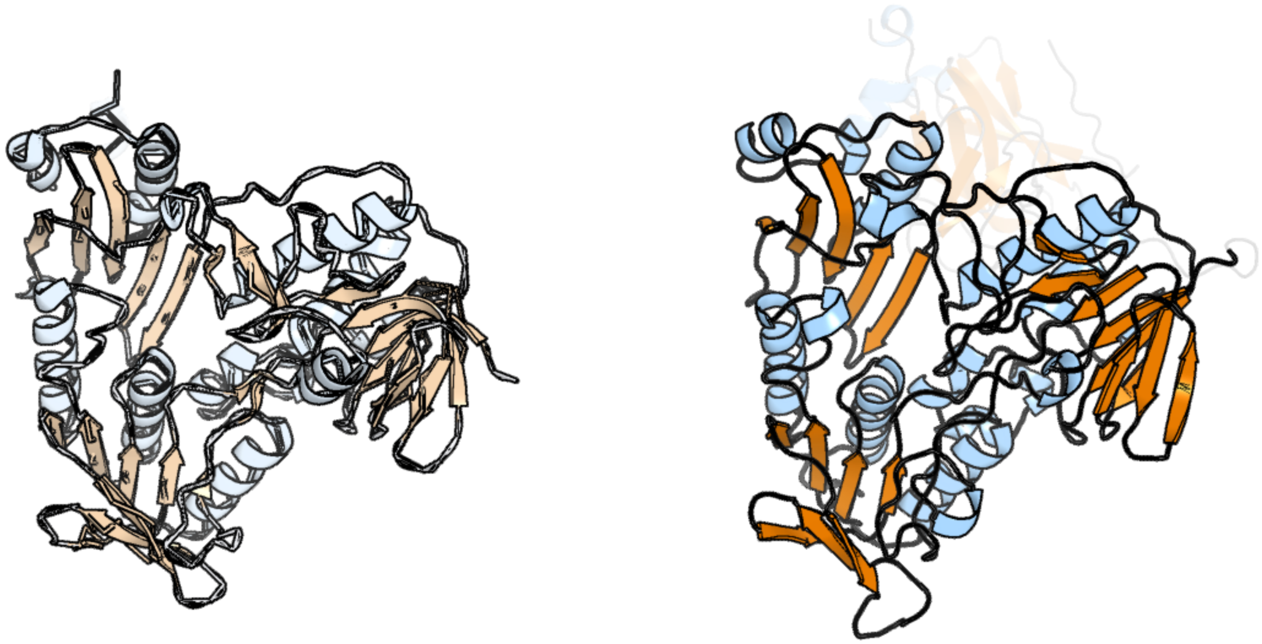

Figure 66: left: reference structure of 3lxd chain A. right: predicted structure of kaa0164596, unaligned sequences are shown as transparent

#### kaa0164643

- Sequence-based annotation for kaa0164643 (predicted in DGTS from hits to PF13649) is hypothetical protein FNF28_03704 [Cafeteria roenbergensis]
- Best hit was 2yqz chain B: Hypothetical protein TTHA0223

| target | prob | fident | alnlen | evalue | theader |
|---|---|---|---|---|---|
| 2yqz-assembly2.cif.gz_B | 1 | 0.175 | 381 | 1.897e-11 | Crystal Structure of Hypothetical Methyltransferase TTHA0223 from Thermus thermophilus HB8 complexed with S-adenosylmethionine |
| 5w7m-assembly1.cif.gz_A | 1 | 0.144 | 381 | 5.576e-11 | Crystal structure of RoqN |
| 1vl5-assembly4.cif.gz_D | 1 | 0.114 | 348 | 1.078e-10 | CRYSTAL STRUCTURE OF A PUTATIVE METHYLTRANSFERASE (BH2331) FROM BACILLUS HALODURANS C-125 AT 1.95 A RESOLUTION |

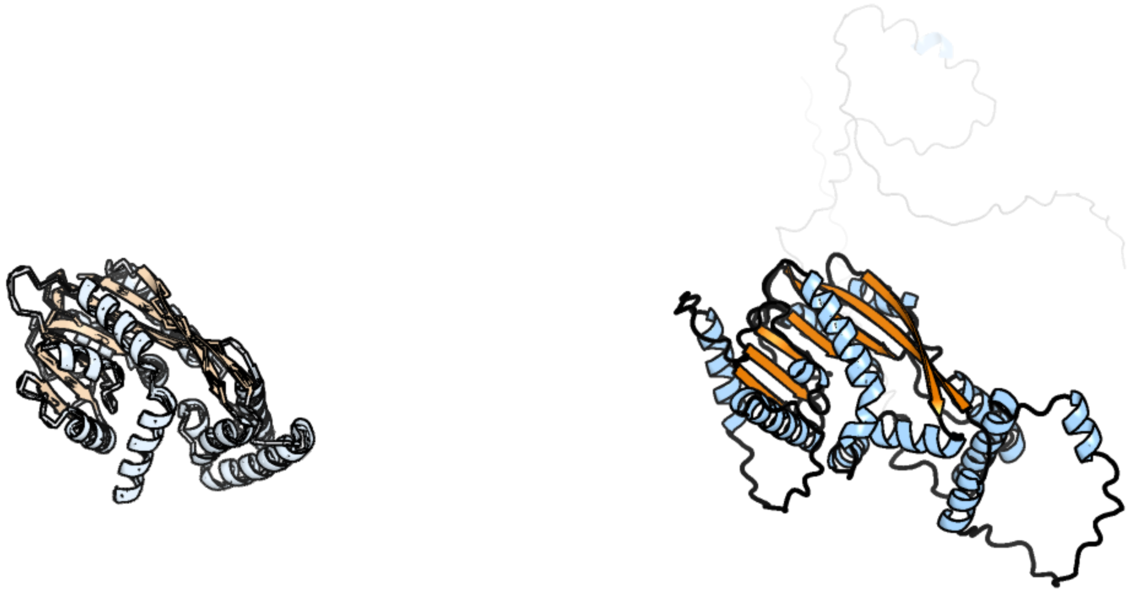

Figure 67: left: reference structure of 2yqz chain B. right: predicted structure of kaa0164643, unaligned sequences are shown as transparent

#### kaa0164792

- Sequence-based annotation for kaa0164792 (predicted in DGTS from hits to PF13649) is hypothetical protein FNF28_03679 [Cafeteria roenbergensis]
- Best hit was 8csr chain 7: Methyltransferase-like protein 17, mitochondrial

| target | prob | fident | alnlen | evalue | theader |
|---|---|---|---|---|---|
| 8csr-assembly1.cif.gz_7 | 1 | 0.241 | 265 | 1.437e-12 | Human mitochondrial small subunit assembly intermediate (State C) |
| 8csq-assembly1.cif.gz_7 | 1 | 0.237 | 265 | 1.677e-12 | Human mitochondrial small subunit assembly intermediate (State B) |
| 8cst-assembly1.cif.gz_7 | 1 | 0.237 | 261 | 2.533e-12 | Human mitochondrial small subunit assembly intermediate (State E) |

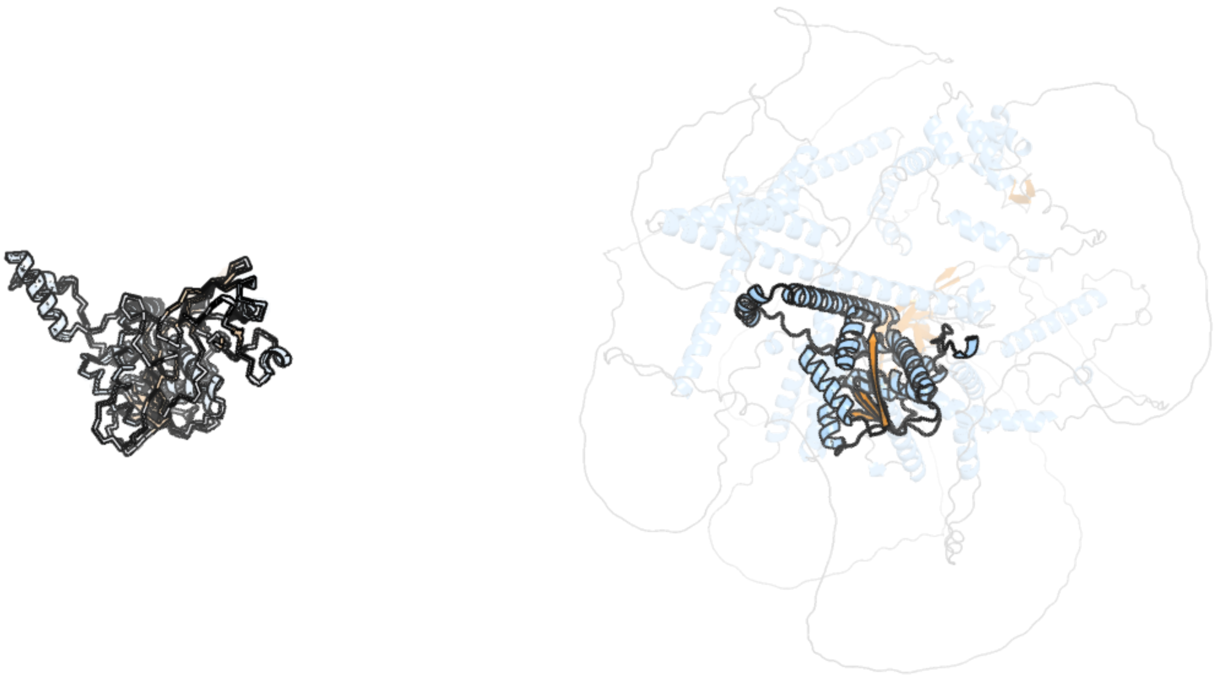

Figure 68: left: reference structure of 8csr chain 7. right: predicted structure of kaa0164792, unaligned sequences are shown as transparent

#### kaa0164796

- Sequence-based annotation for kaa0164796 (predicted in Sulfonolipids from hits to PF00106, UniRef50_A0A383U0M1) is hypothetical protein FNF28_03683 [Cafeteria roenbergensis]
- Best hit was 4o5o chain B: 3-hydroxyacyl-CoA dehydrogenase

| target | prob | fident | alnlen | evalue | theader |
|---|---|---|---|---|---|
| 4o5o-assembly1.cif.gz_B-2 | 1 | 0.515 | 260 | 3.675e-35 | X-ray Crystal Structure of a 3-hydroxyacyl-CoA dehydrogenase from Brucella suis |
| 4pn3-assembly1.cif.gz_A | 1 | 0.515 | 260 | 1.017e-34 | Crystal structure of 3-hydroxyacyl-CoA-dehydrogenase from Brucella melitensis |
| 4xgn-assembly2.cif.gz_E | 1 | 0.498 | 259 | 6.911e-34 | Crystal structure of 3-hydroxyacyl-CoA dehydrogenase in complex with NAD from Burkholderia thailandensis |

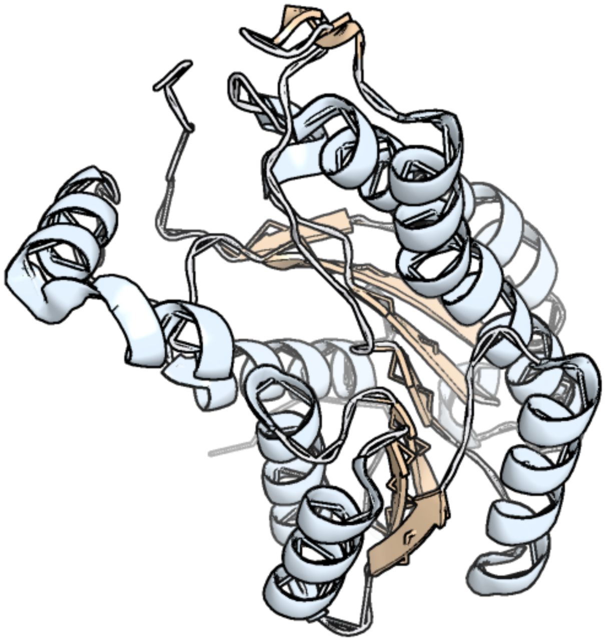

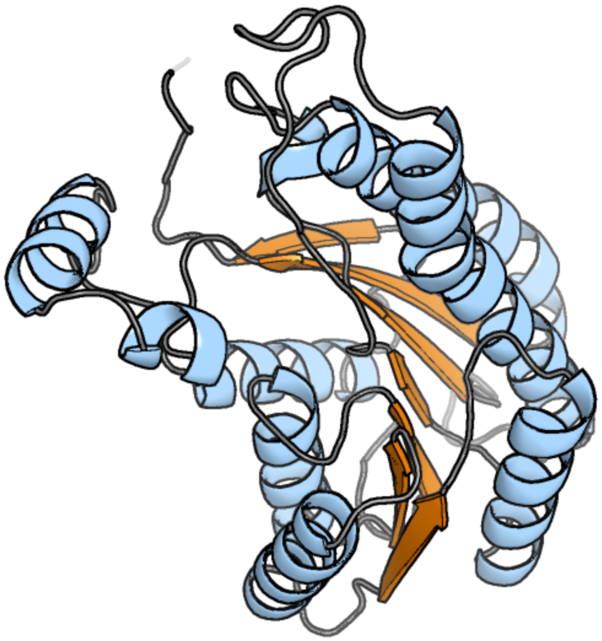

Figure 69: left: reference structure of 4o5o chain B. right: predicted structure of kaa0164796, unaligned sequences are shown as transparent

#### kaa0164992

- Sequence-based annotation for kaa0164992 (predicted in Sulfonolipids from hits to PF00106, UniRef50_A0A383U0M1) is hypothetical protein FNF28_03615 [Cafeteria roenbergensis]
- Best hit was 2b4q chain A: Rhamnolipids biosynthesis 3-oxoacyl-[acyl-carrier-protein] reductase

| target | prob | fident | alnlen | evalue | theader |
|---|---|---|---|---|---|
| 2b4q-assembly1.cif.gz_A | 1 | 0.424 | 257 | 3.367e-29 | Pseudomonas aeruginosa RhlG/NADP active-site complex |
| 2b4q-assembly1.cif.gz_B | 1 | 0.411 | 255 | 1.013e-26 | Pseudomonas aeruginosa RhlG/NADP active-site complex |
| 3r1i-assembly1.cif.gz_A-2 | 1 | 0.309 | 262 | 5.818e-24 | Crystal structure of a short-chain type dehydrogenase/reductase from Mycobacterium marinum |

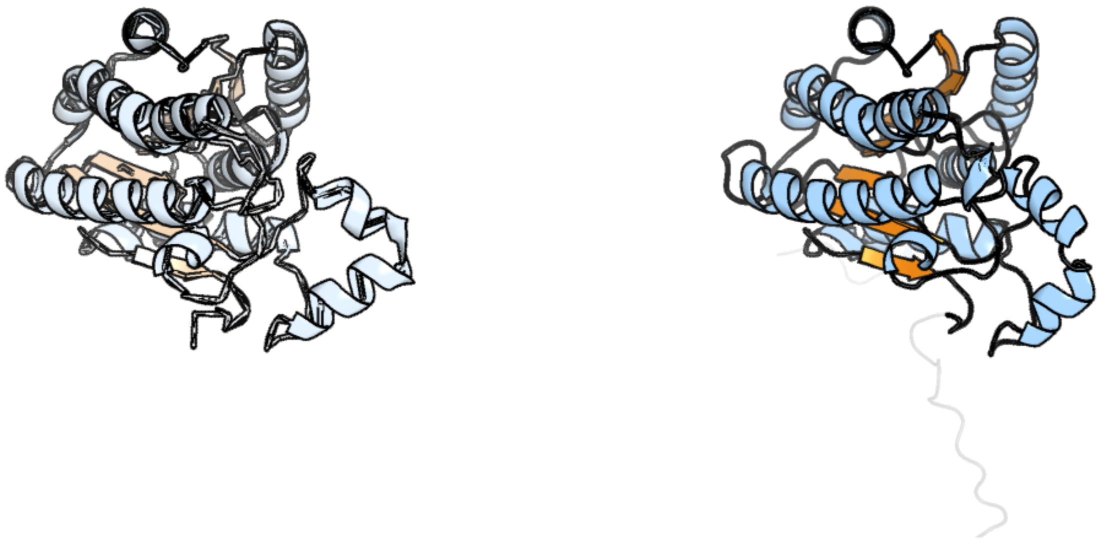

Figure 70: left: reference structure of 2b4q chain A. right: predicted structure of kaa0164992, unaligned sequences are shown as transparent

#### kaa0165176

- Sequence-based annotation for kaa0165176 (predicted in Sulfonolipids from hits to PF00155) is hypothetical protein FNF28_03575 [Cafeteria roenbergensis]
- Best hit was 6f77 chain F: Aspartate aminotransferase A

| target | prob | fident | alnlen | evalue | theader |
|---|---|---|---|---|---|
| 6f77-assembly3.cif.gz_F | 1 | 0.258 | 395 | 1.641e-25 | Crystal structure of the prephenate aminotransferase from Rhizobium meliloti |
| 8wkj-assembly1.cif.gz_A-2 | 1 | 0.243 | 386 | 6.58e-25 | The crystal structure of aspartate aminotransferases Lpg0070 from Legionella pneumophila |
| 5bj4-assembly1.cif.gz_A | 1 | 0.229 | 387 | 1.249e-24 | THERMUS THERMOPHILUS ASPARTATE AMINOTRANSFERASE TETRA MUTANT 2 |

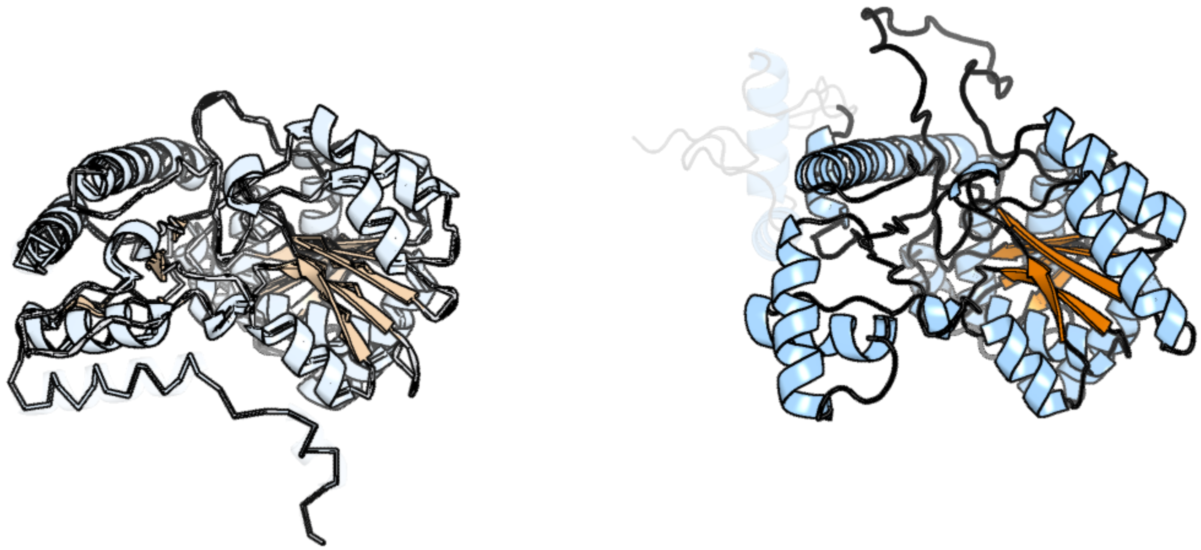

Figure 71: left: reference structure of 6f77 chain F. right: predicted structure of kaa0165176, unaligned sequences are shown as transparent

#### kaa0165429

- Sequence-based annotation for kaa0165429 (predicted in Sulfonolipids from hits to PF00291, UniRef50_A0A2D9B818) is hypothetical protein FNF28_03485 [Cafeteria roenbergensis]
- Best hit was 5jjc chain B: Cysteine synthase

| target | prob | fident | alnlen | evalue | theader |
|---|---|---|---|---|---|
| 5jjc-assembly2.cif.gz_B | 1 | 0.32 | 409 | 4.119e-31 | Crystal Structure of double mutant (Q96A-Y125A) O-Acetyl Serine Sulfhydralase from Brucella abortus |
| 5i7w-assembly2.cif.gz_B | 1 | 0.32 | 409 | 1.006e-30 | Crystal Structure of a Cysteine Synthase from Brucella suis |
| 5jis-assembly1.cif.gz_A | 1 | 0.317 | 412 | 1.257e-30 | The Crystal Structure of O-acetyl serine sulfhydralase from Brucella abortus |

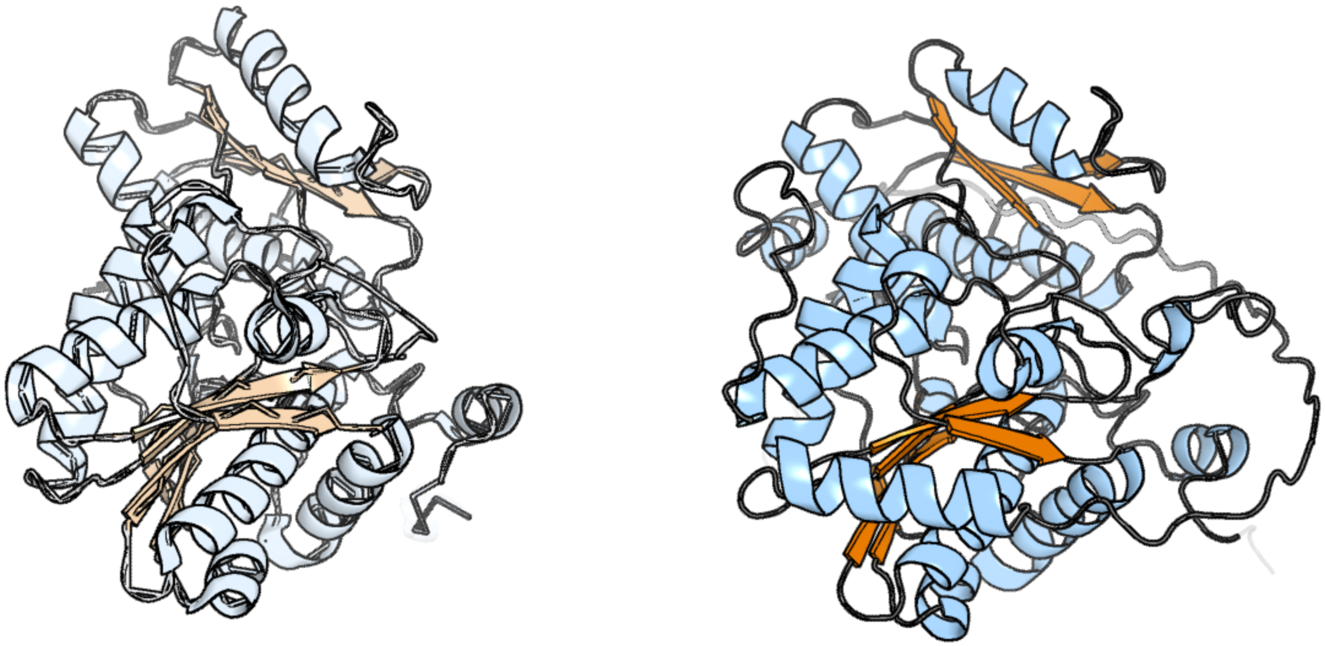

Figure 72: left: reference structure of 5jjc chain B. right: predicted structure of kaa0165429, unaligned sequences are shown as transparent

#### kaa0165655

- Sequence-based annotation for kaa0165655 (predicted in Sulfonolipids from hits to PF00106, UniRef50_A0A383U0M1) is hypothetical protein FNF28_03405 [Cafeteria roenbergensis]
- Best hit was 3l77 chain A: Short-chain alcohol dehydrogenase

| target | prob | fident | alnlen | evalue | theader |
|---|---|---|---|---|---|
| 3l77-assembly1.cif.gz_A | 1 | 0.286 | 241 | 1.773e-24 | X-ray structure alcohol dehydrogenase from archaeon Thermococcus sibiricus complexed with 5-hydroxy-NADP |
| 2bd0-assembly1.cif.gz_D | 1 | 0.283 | 243 | 2.567e-24 | Chlorobium tepidum Sepiapterin Reductase complexed with NADP and Sepiapterin |
| 3p19-assembly1.cif.gz_A | 1 | 0.266 | 255 | 2.187e-22 | Improved NADPH-dependent Blue Fluorescent Protein |

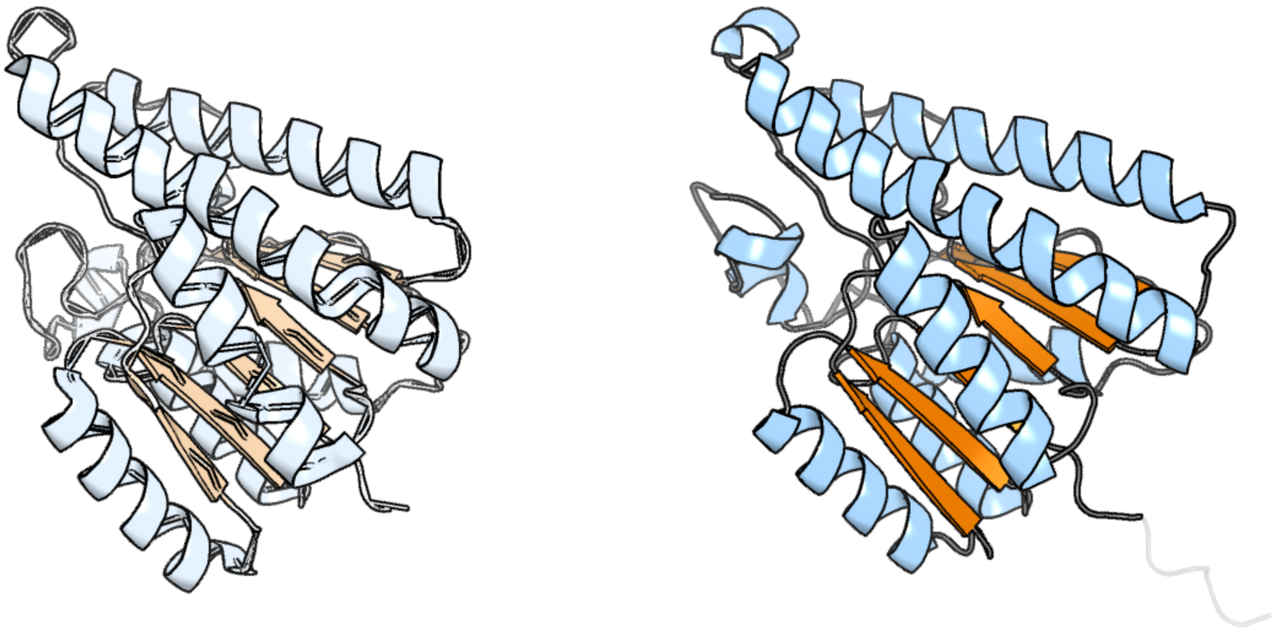

Figure 73: left: reference structure of 3l77 chain A. right: predicted structure of kaa0165655, unaligned sequences are shown as transparent

#### kaa0165836

- Sequence-based annotation for kaa0165836 (predicted in Sulfonolipids from hits to PF00106, UniRef50_A0A383U0M1) is hypothetical protein FNF28_03342 [Cafeteria roenbergensis]
- Best hit was 1wmb chain A: D(-)-3-hydroxybutyrate dehydrogenase

| target | prob | fident | alnlen | evalue | theader |
|---|---|---|---|---|---|
| 1wmb-assembly1.cif.gz_A | 1 | 0.555 | 261 | 4.069e-33 | Crystal structure of NAD dependent D-3-hydroxybutylate dehydrogenase |
| 2ztv-assembly1.cif.gz_C | 1 | 0.551 | 261 | 8.812e-33 | The binary complex of D-3-hydroxybutyrate dehydrogenase with NAD+ |
| 2ztv-assembly1.cif.gz_D | 1 | 0.544 | 261 | 3.912e-32 | The binary complex of D-3-hydroxybutyrate dehydrogenase with NAD+ |

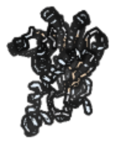

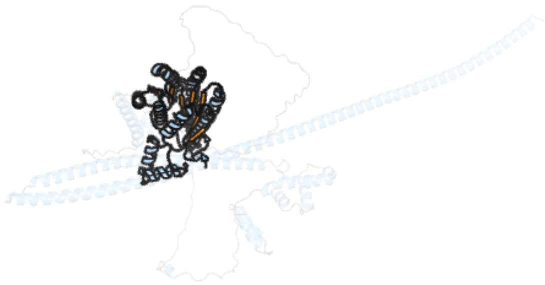

Figure 74: left: reference structure of 1wmb chain A. right: predicted structure of kaa0165836, unaligned sequences are shown as transparent

### kaa0165928

- Sequence-based annotation for kaa0165928 (predicted in Sulfonolipids from hits to PF00106) is hypothetical protein FNF28_03310 [Cafeteria roenbergensis]
- Best hit was 1zbq chain F: 17-beta-hydroxysteroid dehydrogenase 4

| target | prob | fident | alnlen | evalue | theader |
|---|---|---|---|---|---|
| 1zbq-assembly3.cif.gz_F | 1 | 0.521 | 305 | 3.507e-42 | Crystal Structure Of Human 17-Beta-Hydroxysteroid Dehydrogenase Type 4 In Complex With NAD |
| 1gz6-assembly2.cif.gz_D | 1 | 0.519 | 304 | 5.602e-41 | (3R)-HYDROXYACYL-COA DEHYDROGENASE FRAGMENT OF RAT PEROXISOMAL MULTIFUNCTIONAL ENZYME TYPE 2 |
| 1gz6-assembly2.cif.gz_C | 1 | 0.495 | 303 | 1.594e-39 | (3R)-HYDROXYACYL-COA DEHYDROGENASE FRAGMENT OF RAT PEROXISOMAL MULTIFUNCTIONAL ENZYME TYPE 2 |

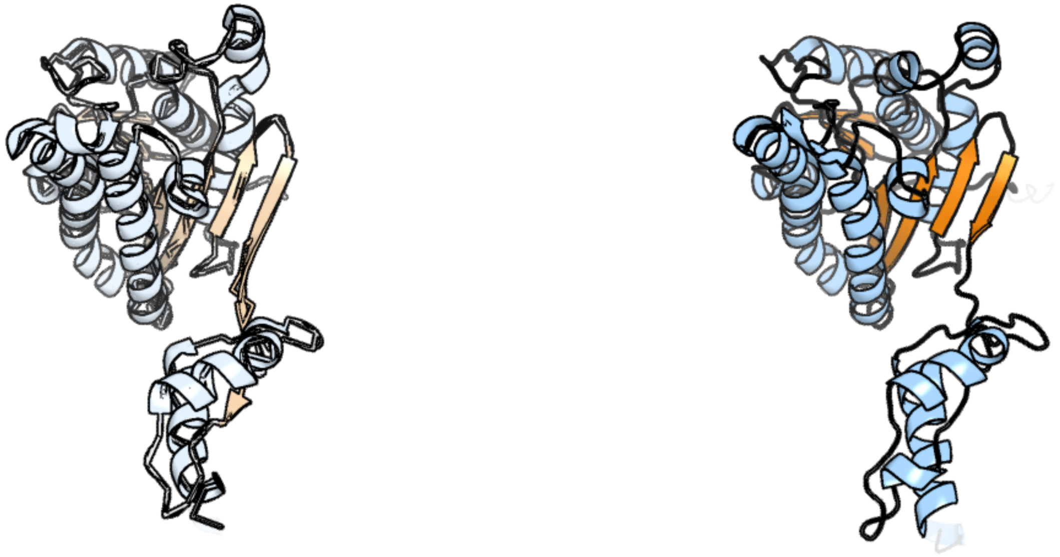

Figure 75: left: reference structure of 1zbq chain F. right: predicted structure of kaa0165928, unaligned sequences are shown as transparent

### kaa0166079

- Sequence-based annotation for kaa0166079 (predicted in DGTS from hits to PF13649) is hypothetical protein FNF28_03247 [Cafeteria roenbergensis]
- Best hit was 6nt2 chain A: Protein arginine N-methyltransferase 1

| target | prob | fident | alnlen | evalue | theader |
|---|---|---|---|---|---|
| 6nt2-assembly1.cif.gz_A | 1 | 0.277 | 379 | 9.522e-29 | type 1 PRMT in complex with the inhibitor GSK3368715 |
| 8g2g-assembly1.cif.gz_A | 1 | 0.295 | 382 | 3.501e-28 | Crystal structure of PRMT3 with compound YD1113 |
| 8bva-assembly1.cif.gz_A-2 | 1 | 0.28 | 389 | 8.663e-28 | Crystal Structure of Mus musculus Protein Arginine Methyltransferase 2 in complex with RSF1_114-126 |

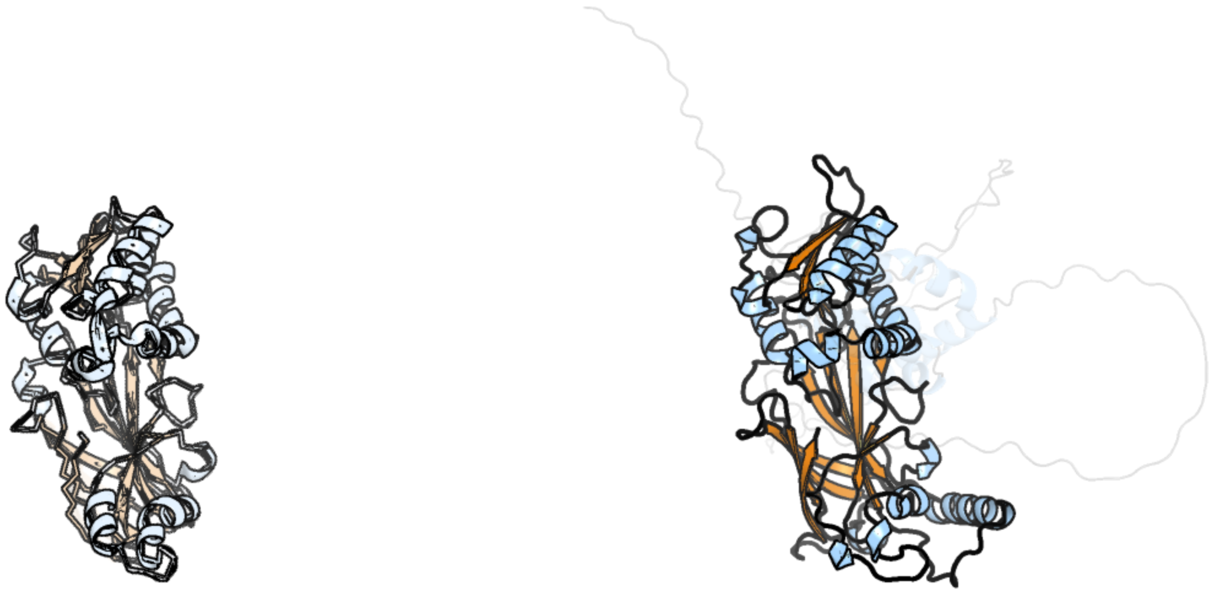

Figure 76: left: reference structure of 6nt2 chain A. right: predicted structure of kaa0166079, unaligned sequences are shown as transparent

#### kaa0166163

- Sequence-based annotation for kaa0166163 (predicted in Sulfonolipids from hits to PF00106, UniRef50_A0A383U0M1) is hypothetical protein FNF28_03211 [Cafeteria roenbergensis]
- Best hit was 8g9v chain B: 17-beta-hydroxysteroid dehydrogenase 13

| target | prob | fident | alnlen | evalue | theader |
|---|---|---|---|---|---|
| 8g9v-assembly1.cif.gz_B | 1 | 0.377 | 273 | 1.398e-28 | Crystal structures of 17-beta-hydroxysteroid dehydrogenase 13 |
| 8g9v-assembly4.cif.gz_H | 1 | 0.361 | 282 | 1.574e-28 | Crystal structures of 17-beta-hydroxysteroid dehydrogenase 13 |
| 8g9v-assembly1.cif.gz_A | 1 | 0.363 | 283 | 2.117e-28 | Crystal structures of 17-beta-hydroxysteroid dehydrogenase 13 |

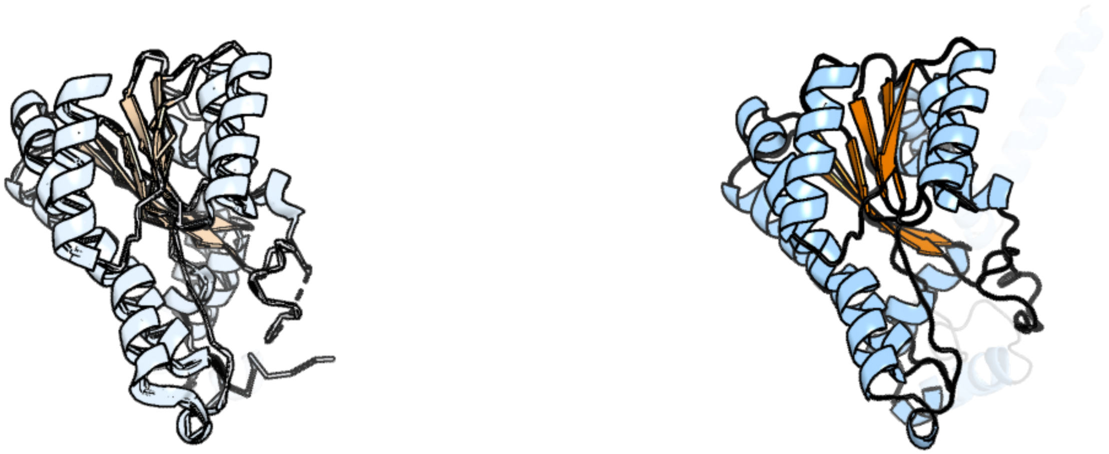

Figure 77: left: reference structure of 8g9v chain B. right: predicted structure of kaa0166163, unaligned sequences are shown as transparent

#### kaa0166380

- Sequence-based annotation for kaa0166380 (predicted in Fu-FA,DGTS from hits to PF01593, PF02353, PF13649, UniRef50_A0A1V0RIH1, UniRef50_Q5LTD2) is hypothetical protein FNF28_03149 [Cafeteria roenbergensis]
- Best hit was 6f7l chain A: Amine oxidase LkcE

| target | prob | fident | alnlen | evalue | theader |
|---|---|---|---|---|---|
| 6f7l-assembly1.cif.gz_A | 1 | 0.278 | 464 | 1.562e-27 | Crystal structure of LkcE R326Q mutant in complex with its substrate |
| 6f32-assembly1.cif.gz_A | 1 | 0.278 | 470 | 1.562e-27 | Crystal structure of a dual function amine oxidase/cyclase in complex with substrate analogues |
| 6f7v-assembly1.cif.gz_A | 1 | 0.284 | 467 | 3.761e-27 | Crystal structure of LkcE E64Q mutant in complex with LC-KA05 |

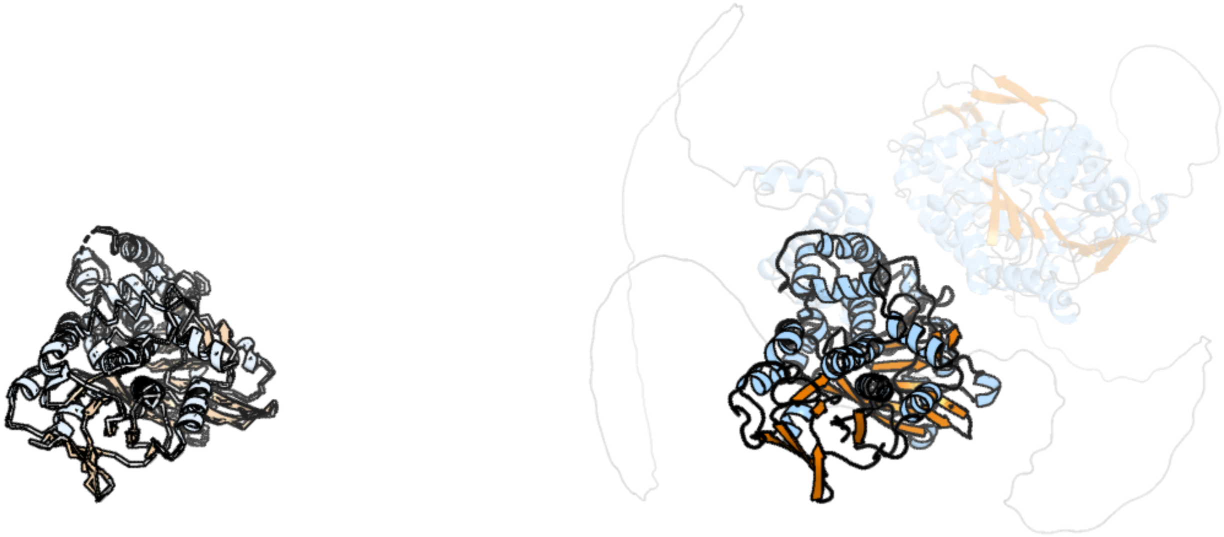

Figure 78: left: reference structure of 6f7l chain A. right: predicted structure of kaa0166380, unaligned sequences are shown as transparent

#### kaa0166382

- Sequence-based annotation for kaa0166382 (predicted in Fu-FA from hits to PF07103, UniRef50_A0A1W2C0P8) is hypothetical protein FNF28_03151 [Cafeteria roenbergensis]
- No significant structural hit found

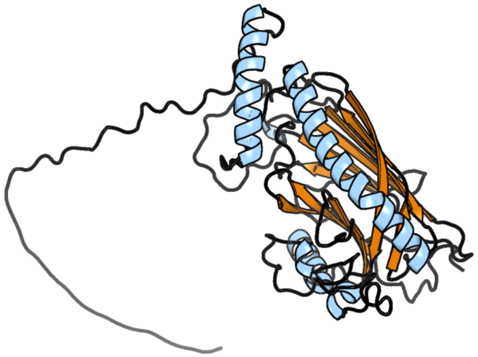

Figure 79: predicted structure of kaa0166382

#### kaa0166390

- Sequence-based annotation for kaa0166390 (predicted in DGTS from hits to PF00010) is hypothetical protein FNF28_03159 [Cafeteria roenbergensis]
- Best hit was 8osk chain M: Circadian locomoter output cycles protein kaput

| target | prob | fident | alnlen | evalue | theader |
|---|---|---|---|---|---|
| 8osk-assembly1.cif.gz_M | 1 | 0.188 | 186 | 2.866e-05 | Cryo-EM structure of CLOCK-BMAL1 bound to a nucleosomal E-box at position SHL+5.8 (composite map) |
| 8osl-assembly1.cif.gz_M | 1 | 0.208 | 168 | 3.448e-05 | Cryo-EM structure of CLOCK-BMAL1 bound to the native Por enhancer nucleosome (map 2, additional 3D classification and flexible refinement) |
| 8osk-assembly1.cif.gz_N | 1 | 0.168 | 178 | 9.239e-05 | Cryo-EM structure of CLOCK-BMAL1 bound to a nucleosomal E-box at position SHL+5.8 (composite map) |

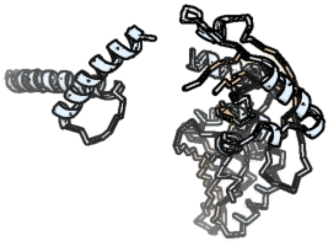

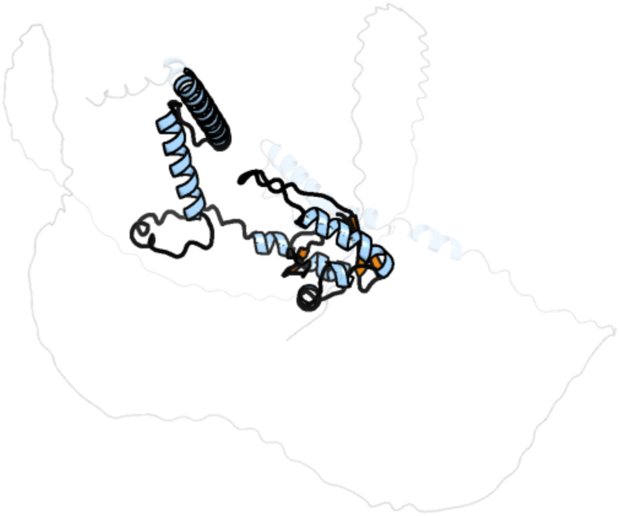

Figure 80: left: reference structure of 8osk chain M. right: predicted structure of kaa0166390, unaligned sequences are shown as transparent

### kaa0166943

- Sequence-based annotation for kaa0166943 (predicted in Sulfonolipids from hits to PF00155) is hypothetical protein FNF28_03014 [Cafeteria roenbergensis]
- Best hit was 3fvx chain B: Kynurenine–oxoglutarate transaminase 1

| target | prob | fident | alnlen | evalue | theader |
|---|---|---|---|---|---|
| 3fvx-assembly1.cif.gz_B | 1 | 0.323 | 433 | 1.044e-39 | Human kynurenine aminotransferase I in complex with tris |
| 4wp0-assembly1.cif.gz_B | 1 | 0.324 | 432 | 1.688e-39 | Crystal structure of human kynurenine aminotransferase-I with a C-terminal V5-hexahistidine tag |
| 4wp0-assembly2.cif.gz_C | 1 | 0.324 | 428 | 3.037e-39 | Crystal structure of human kynurenine aminotransferase-I with a C-terminal V5-hexahistidine tag |

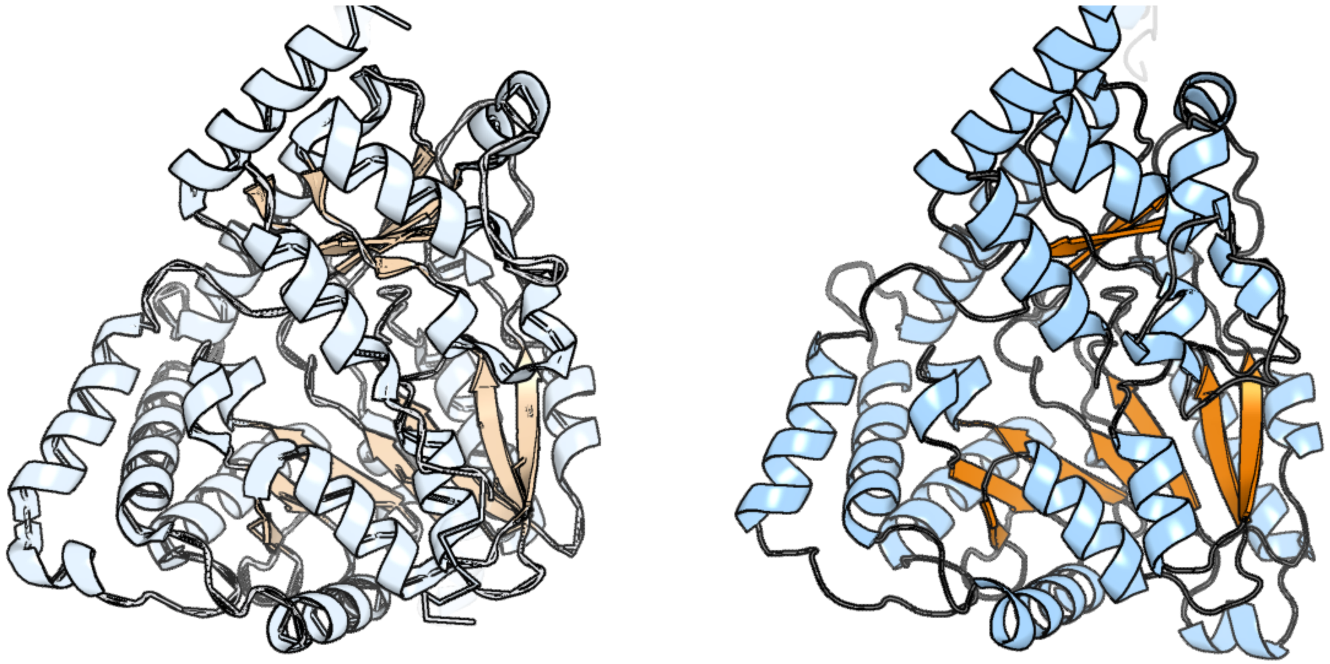

Figure 81: left: reference structure of 3fvx chain B. right: predicted structure of kaa0166943, unaligned sequences are shown as transparent

#### kaa0167551

- Sequence-based annotation for kaa0167551 (predicted in Sulfonolipids from hits to PF00106, UniRef50_A0A383U0M1) is hypothetical protein FNF28_02765 [Cafeteria roenbergensis]
- Best hit was 8g89 chain A: Hydroxysteroid 17-beta dehydrogenase 13

| target | prob | fident | alnlen | evalue | theader |
|---|---|---|---|---|---|
| 8g89-assembly1.cif.gz_A | 1 | 0.223 | 313 | 1.113e-16 | HSD17B13 in complex with cofactor and inhibitor |
| 8g9v-assembly1.cif.gz_B | 1 | 0.211 | 307 | 1.244e-16 | Crystal structures of 17-beta-hydroxysteroid dehydrogenase 13 |
| 8g93-assembly1.cif.gz_B | 1 | 0.212 | 296 | 2.171e-16 | Crystal structures of 17-beta-hydroxysteroid dehydrogenase 13 |

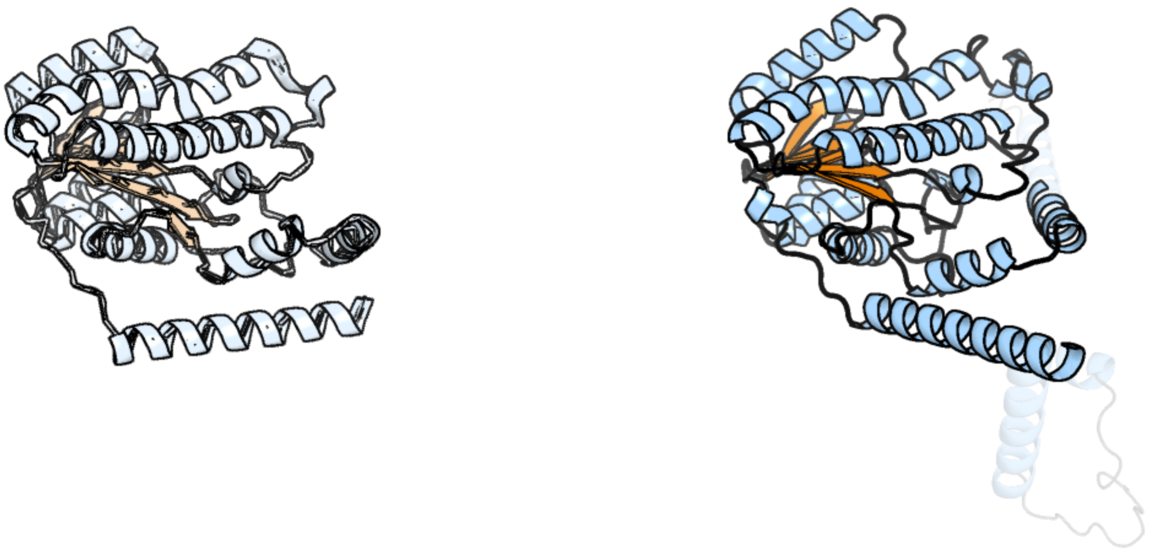

Figure 82: left: reference structure of 8g89 chain A. right: predicted structure of kaa0167551, unaligned sequences are shown as transparent

#### kaa0167555

- Sequence-based annotation for kaa0167555 (predicted in DGTS from hits to PF13649) is hypothetical protein FNF28_02769 [Cafeteria roenbergensis]
- Best hit was 8khq chain A: 5-histidylcysteine sulfoxide synthase/putative 4-mercaptohistidine N1-methyltranferase

| target | prob | fident | alnlen | evalue | theader |
|---|---|---|---|---|---|
| 8khq-assembly1.cif.gz_A | 1 | 0.344 | 258 | 4.956e-27 | Bifunctional sulfoxide synthase OvoA_Th2 in complex with histidine and cysteine |
| 8khq-assembly2.cif.gz_D | 1 | 0.351 | 262 | 8.857e-27 | Bifunctional sulfoxide synthase OvoA_Th2 in complex with histidine and cysteine |
| 8khq-assembly1.cif.gz_C | 1 | 0.347 | 273 | 7.593e-26 | Bifunctional sulfoxide synthase OvoA_Th2 in complex with histidine and cysteine |

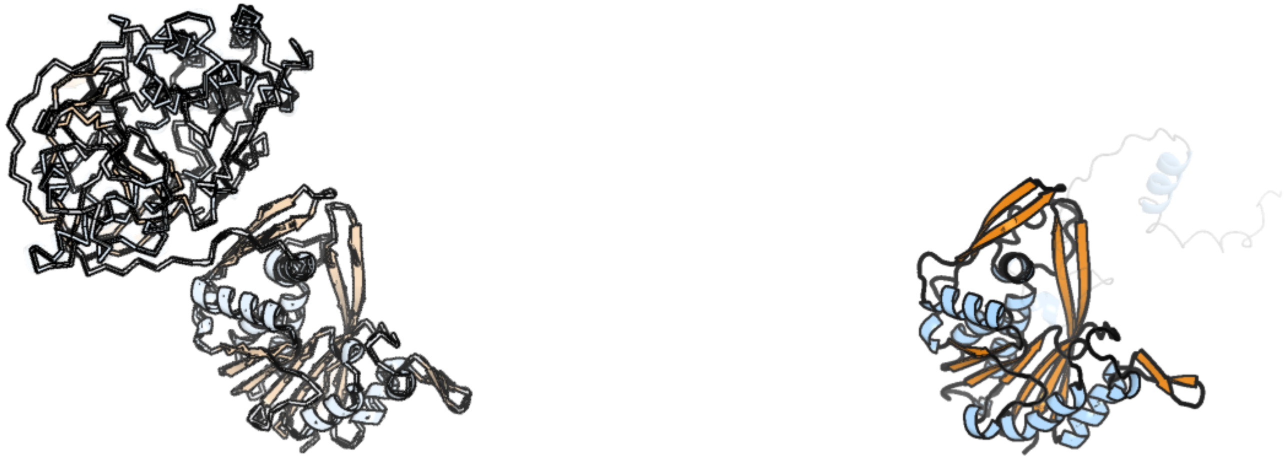

Figure 83: left: reference structure of 8khq chain A. right: predicted structure of kaa0167555, unaligned sequences are shown as transparent

#### kaa0167599

- Sequence-based annotation for kaa0167599 (predicted in DGTS from hits to PF13649) is hypothetical protein FNF28_02813 [Cafeteria roenbergensis]
- Best hit was 5w7k chain A: OxaG

| target | prob | fident | alnlen | evalue | theader |
|---|---|---|---|---|---|
| 5w7k-assembly1.cif.gz_A | 1 | 0.162 | 344 | 8.769e-13 | Crystal structure of OxaG |
| 5w7k-assembly2.cif.gz_B | 1 | 0.163 | 343 | 1.697e-12 | Crystal structure of OxaG |
| 3ccf-assembly1.cif.gz_A | 1 | 0.19 | 288 | 5.098e-12 | Crystal structure of putative methyltransferase (YP_321342.1) from Anabaena variabilis ATCC 29413 at 1.90 A resolution |

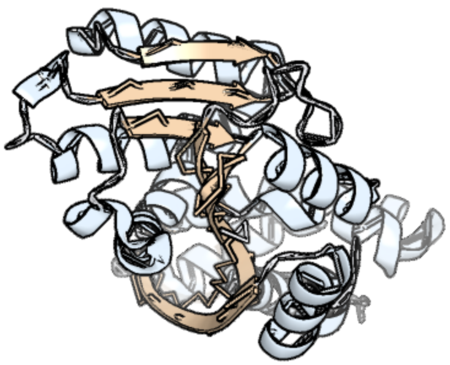

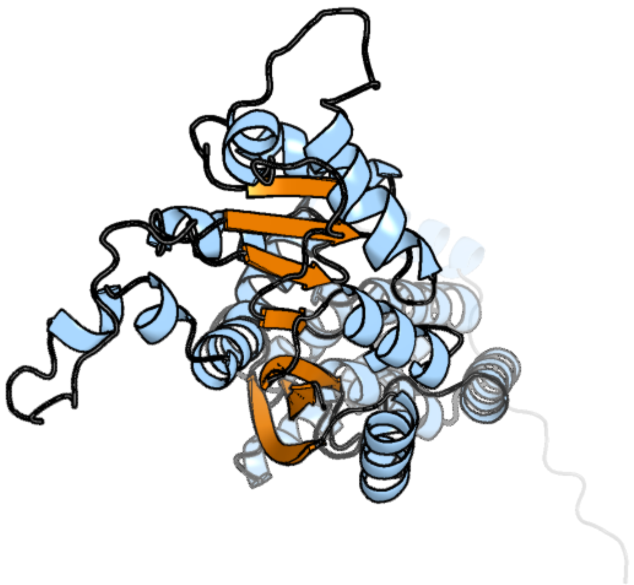

Figure 84: left: reference structure of 5w7k chain A. right: predicted structure of kaa0167599, unaligned sequences are shown as transparent

#### kaa0167909

- Sequence-based annotation for kaa0167909 (predicted in DGTS from hits to PF13649) is hypothetical protein FNF28_02655 [Cafeteria roenbergensis]
- Best hit was 4obx chain B: 2-methoxy-6-polyprenyl-1,4-benzoquinol methylase, mitochondrial

| target | prob | fident | alnlen | evalue | theader |
|---|---|---|---|---|---|
| 4obx-assembly1.cif.gz_B | 1 | 0.468 | 237 | 6.136e-27 | Crystal structure of yeast Coq5 in the apo form |
| 1xxl-assembly2.cif.gz_B | 1 | 0.166 | 264 | 5.278e-13 | The crystal structure of YcgJ protein from Bacillus subitilis at 2.1 A resolution |
| 5mgz-assembly1.cif.gz_B | 1 | 0.164 | 243 | 5.953e-13 | Streptomyces Spheroides NovO (8-demethylnovbiocic acid methyltransferase) with SAH |

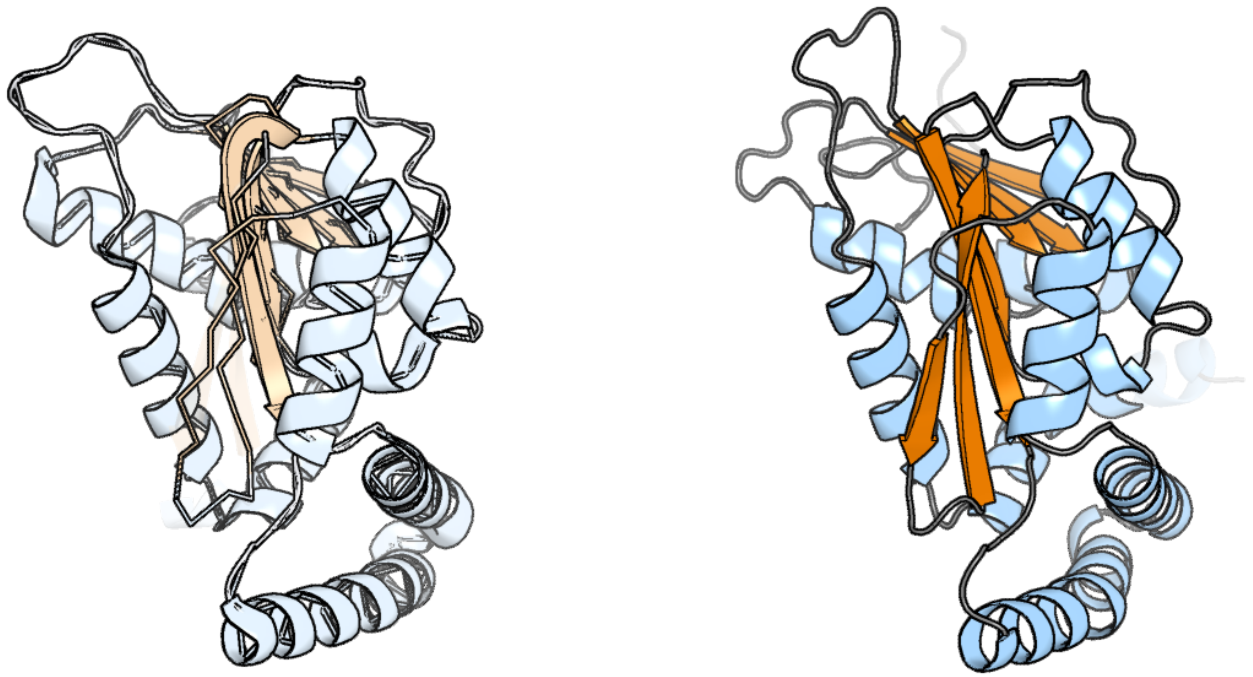

Figure 85: left: reference structure of 4obx chain B. right: predicted structure of kaa0167909, unaligned sequences are shown as transparent

#### kaa0167919

- Sequence-based annotation for kaa0167919 (predicted in Fu-FA,DGTS from hits to PF02353, PF13649) is hypothetical protein FNF28_02665 [Cafeteria roenbergensis]
- Best hit was 6nt2 chain A: Protein arginine N-methyltransferase 1

| target | prob | fident | alnlen | evalue | theader |
|---|---|---|---|---|---|
| 6nt2-assembly1.cif.gz_A | 1 | 0.583 | 329 | 1.087e-52 | type 1 PRMT in complex with the inhibitor GSK3368715 |
| 1ori-assembly1.cif.gz_A | 1 | 0.572 | 318 | 1.098e-50 | Structure of the predominant protein arginine methyltransferase PRMT1 |
| 5dst-assembly5.cif.gz_J | 1 | 0.547 | 318 | 1.67e-50 | Crystal structure of human PRMT8 in complex with SAH |

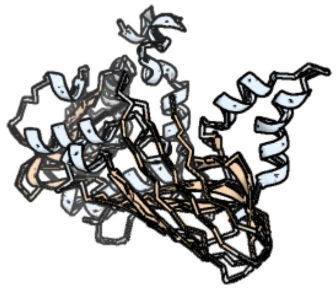

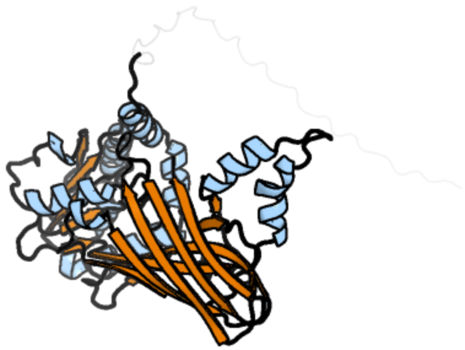

Figure 86: left: reference structure of 6nt2 chain A. right: predicted structure of kaa0167919, unaligned sequences are shown as transparent

### kaa0167921

- Sequence-based annotation for kaa0167921 (predicted in Sulfonolipids from hits to PF00155) is hypothetical protein FNF28_02667 [Cafeteria roenbergensis]
- Best hit was 3dyd chain B: Tyrosine aminotransferase

| target | prob | fident | alnlen | evalue | theader |
|---|---|---|---|---|---|
| 3dyd-assembly1.cif.gz_B | 1 | 0.497 | 388 | 1.172e-49 | Human Tyrosine Aminotransferase |
| 3dyd-assembly1.cif.gz_A | 1 | 0.498 | 389 | 7.262e-49 | Human Tyrosine Aminotransferase |
| 1bw0-assembly1.cif.gz_A | 1 | 0.387 | 423 | 8.588e-46 | CRYSTAL STRUCTURE OF TYROSINE AMINOTRANSFERASE FROM TRYPANOSOMA CRUZI |

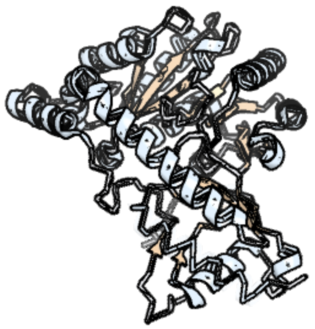

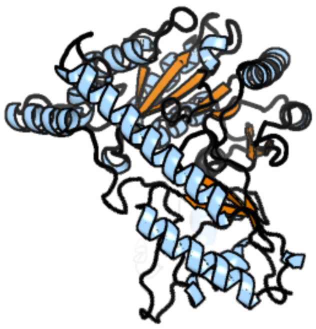

Figure 87: left: reference structure of 3dyd chain B. right: predicted structure of kaa0167921, unaligned sequences are shown as transparent

#### kaa0167946

- Sequence-based annotation for kaa0167946 (predicted in Sulfonolipids from hits to PF00155) is hypothetical protein FNF28_02692 [Cafeteria roenbergensis]
- Best hit was 1wst chain A: multiple substrate aminotransferase

| target | prob | fident | alnlen | evalue | theader |
|---|---|---|---|---|---|
| 1wst-assembly1.cif.gz_A-2 | 1 | 0.324 | 388 | 7.787e-38 | Crystal structure of multiple substrate aminotransferase (MsAT) from Thermococcus profundus |
| 8tn3-assembly1.cif.gz_B | 1 | 0.336 | 380 | 9.182e-37 | Structure of S. hygroscopicus aminotransferase MppQ complexed with pyridoxamine 5'-phosphate (PMP) |
| 3av7-assembly1.cif.gz_A | 1 | 0.307 | 377 | 1.211e-35 | Crystal structure of Pyrococcus horikoshii kynurenine aminotransferase in complex with PMP, KYN as substrates and KYA as products |

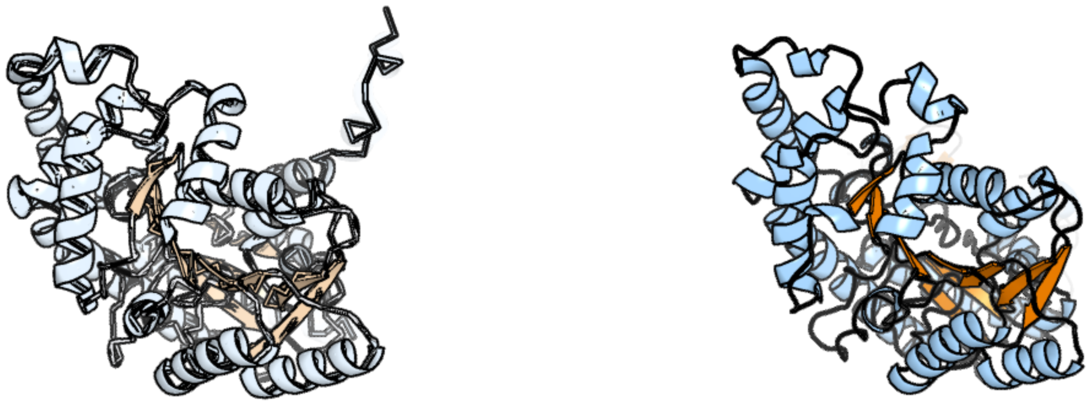

Figure 88: left: reference structure of 1wst chain A. right: predicted structure of kaa0167946, unaligned sequences are shown as transparent

#### kaa0168166

- Sequence-based annotation for kaa0168166 (predicted in Sulfonolipids from hits to PF00106, UniRef50_A0A383U0M1) is hypothetical protein FNF28_02585 [Cafeteria roenbergensis]
- Best hit was 3t4x chain A: Oxidoreductase, short chain dehydrogenase/reductase family

| target | prob | fident | alnlen | evalue | theader |
|---|---|---|---|---|---|
| 3t4x-assembly1.cif.gz_A | 1 | 0.429 | 268 | 7.014e-30 | Short chain dehydrogenase/reductase family oxidoreductase from Bacillus anthracis str. Ames Ancestor |
| 6jhb-assembly1.cif.gz_B-2 | 1 | 0.287 | 271 | 1.628e-22 | Crystal structure of NADPH and 4-hydroxyphenylpyruvic acid bound AerF from Microcystis aeruginosa |
| 4imr-assembly1.cif.gz_A | 1 | 0.344 | 267 | 2.599e-22 | Crystal structure of 3-oxoacyl (acyl-carrier-protein) reductase (target EFI-506442) from agrobacterium tumefaciens C58 with NADP bound |

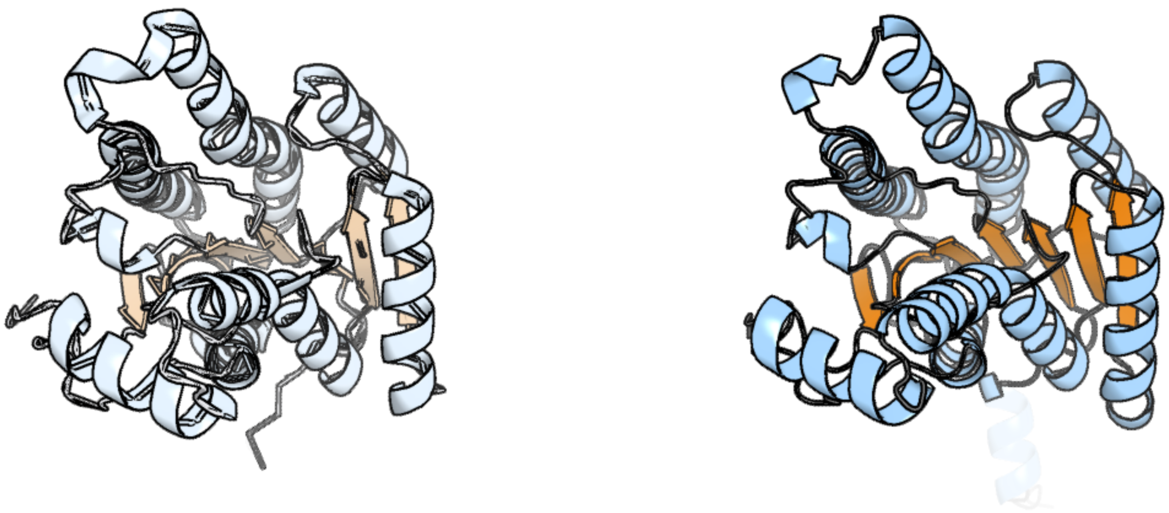

Figure 89: left: reference structure of 3t4x chain A. right: predicted structure of kaa0168166, unaligned sequences are shown as transparent

#### kaa0168203

- Sequence-based annotation for kaa0168203 (predicted in DGTS from hits to PF13649, UniRef50_A0A238JF33) is hypothetical protein FNF28_02621 [Cafeteria roenbergensis]
- Best hit was 1xxl chain B: YcgJ protein

| target | prob | fident | alnlen | evalue | theader |
|---|---|---|---|---|---|
| 1xxl-assembly2.cif.gz_B | 1 | 0.223 | 188 | 5.759e-13 | The crystal structure of YcgJ protein from Bacillus subitilis at 2.1 A resolution |
| 3l8d-assembly1.cif.gz_A-2 | 1 | 0.179 | 206 | 1.637e-12 | Crystal structure of methyltransferase from Bacillus Thuringiensis |
| 6f5z-assembly1.cif.gz_A | 1 | 0.146 | 245 | 1.851e-12 | Complex between the Haloferax volcanii Trm112 methyltransferase activator and the Hvo_0019 putative methyltransferase |

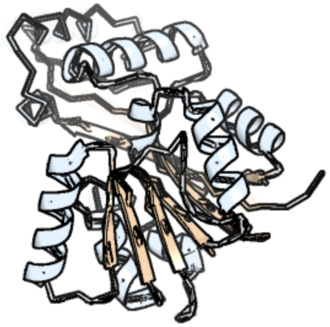

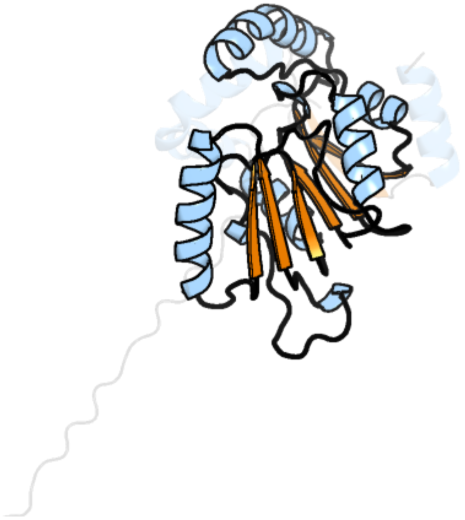

Figure 90: left: reference structure of 1xxl chain B. right: predicted structure of kaa0168203, unaligned sequences are shown as transparent

### kaa0168683

- Sequence-based annotation for kaa0168683 (predicted in Sulfonolipids from hits to PF00106, UniRef50_A0A383U0M1) is hypothetical protein FNF28_02422 [Cafeteria roenbergensis]
- Best hit was 1w8d chain A: 2,4-DIENOYL-COA REDUCTASE, MITOCHONDRIAL PRECURSOR

| target | prob | fident | alnlen | evalue | theader |
|---|---|---|---|---|---|
| 1w8d-assembly1.cif.gz_A | 1 | 0.553 | 278 | 2.585e-40 | Binary structure of human DECR. |
| 1w6u-assembly1.cif.gz_C | 1 | 0.539 | 278 | 3.988e-40 | Structure of human DECR ternary complex |
| 7ucw-assembly1.cif.gz_B | 1 | 0.543 | 276 | 4.514e-40 | Structure of mouse Decr1 in complex with 2'-5' oligoadenylate |

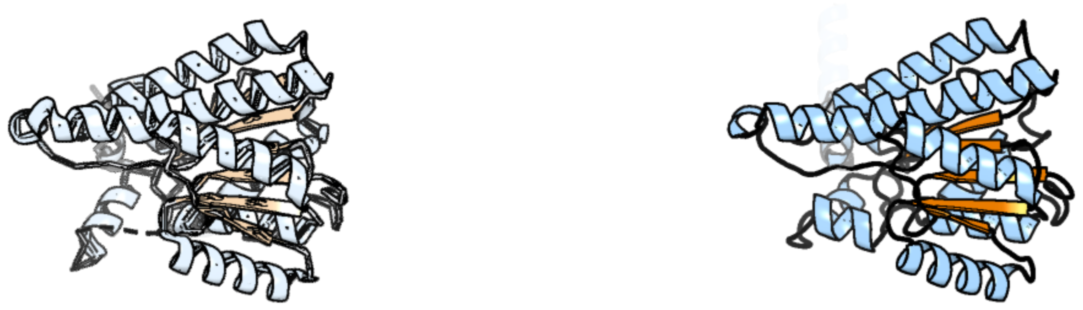

Figure 91: left: reference structure of 1w8d chain A. right: predicted structure of kaa0168683, unaligned sequences are shown as transparent

#### kaa0168703

- Sequence-based annotation for kaa0168703 (predicted in Sulfonolipids from hits to PF00155) is hypothetical protein FNF28_02442 [Cafeteria roenbergensis]
- Best hit was 4ge4 chain A: Kynurenine/alpha-aminoadipate aminotransferase, mitochondrial

| target | prob | fident | alnlen | evalue | theader |
|---|---|---|---|---|---|
| 4ge4-assembly1.cif.gz_A | 1 | 0.363 | 349 | 3.573e-30 | Kynurenine Aminotransferase II Inhibitors |
| 3ue8-assembly3.cif.gz_A | 1 | 0.365 | 350 | 2.912e-29 | Kynurenine Aminotransferase II Inhibitors |
| 6t8p-assembly1.cif.gz_B | 1 | 0.365 | 350 | 5.433e-29 | HKATII IN COMPLEX WITH LIGAND (2R)-N-benzyl-1-[6-methyl-5-(oxan-4-yl)-7-oxo-6H,7H-[1,3]thiazolo[5,4-d]pyrimidin-2-yl]pyrrolidine-2-carboxamide |

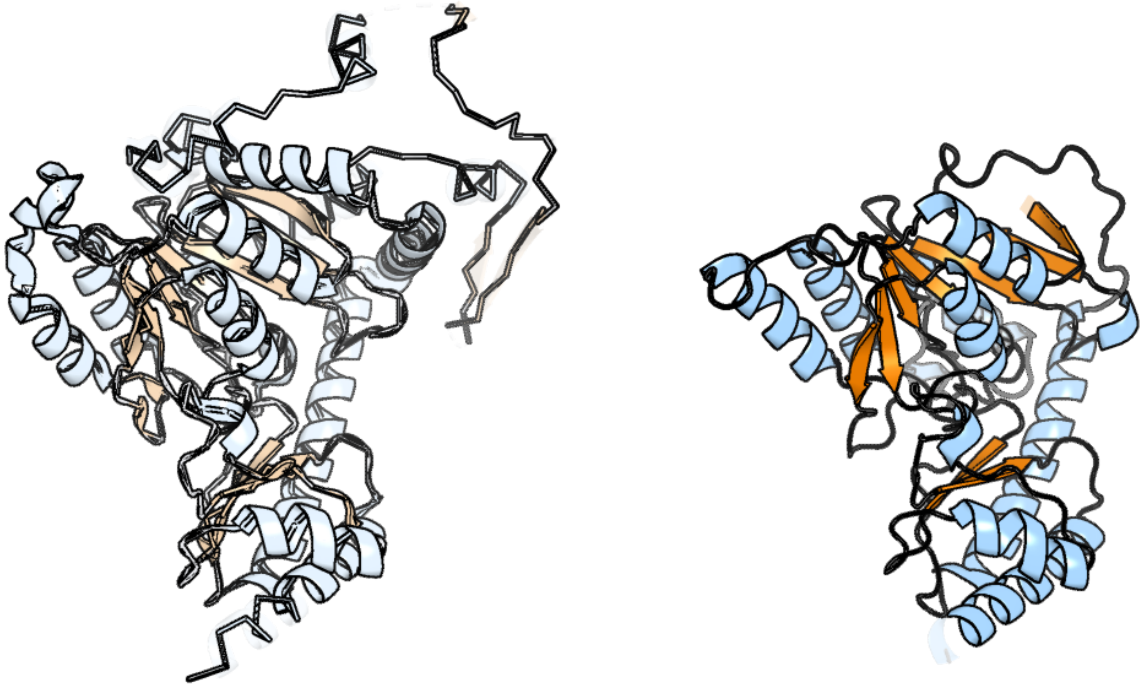

Figure 92: left: reference structure of 4ge4 chain A. right: predicted structure of kaa0168703, unaligned sequences are shown as transparent

#### kaa0168704

- Sequence-based annotation for kaa0168704 (predicted in Sulfonolipids from hits to PF00155) is hypothetical protein FNF28_02442 [Cafeteria roenbergensis]
- Best hit was 6t8q chain A: Kynurenine/alpha-aminoadipate aminotransferase, mitochondrial

| target | prob | fident | alnlen | evalue | theader |
|---|---|---|---|---|---|
| 6t8q-assembly1.cif.gz_A | 1 | 0.378 | 457 | 3.883e-41 | HKATII IN COMPLEX WITH LIGAND (2R)-N-benzyl-1-[6-methyl-5-(oxan-4-yl)-7-oxo-6H,7H-[1,3]thiazolo[5,4-d]pyrimidin-2-yl]pyrrolidine-2-carboxamide |
| 4ge4-assembly1.cif.gz_A | 1 | 0.368 | 459 | 2.886e-40 | Kynurenine Aminotransferase II Inhibitors |
| 6d0a-assembly1.cif.gz_A | 1 | 0.371 | 457 | 1.229e-39 | Crystal structure of Kynurenine Aminotransferase-II in apo-form, at 1.47 A resolution |

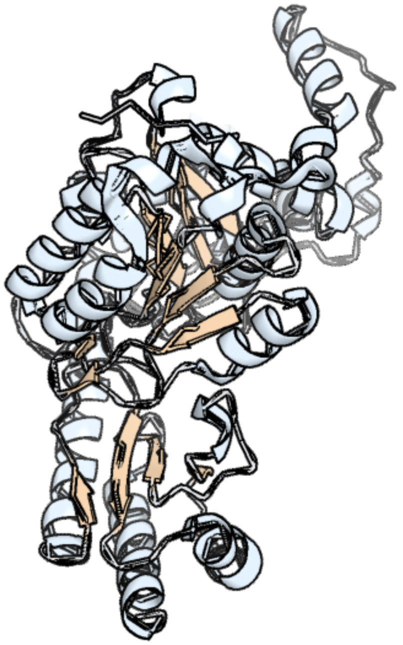

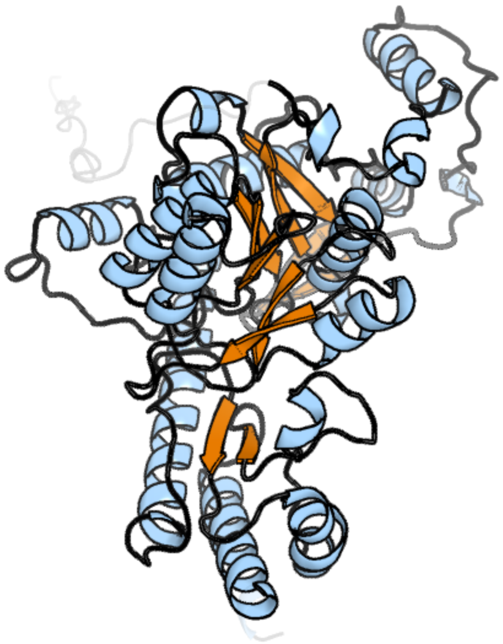

Figure 93: left: reference structure of 6t8q chain A. right: predicted structure of kaa0168704, unaligned sequences are shown as transparent

#### kaa0168801

- Sequence-based annotation for kaa0168801 (predicted in DGTS from hits to PF13649) is hypothetical protein FNF28_02348 [Cafeteria roenbergensis]
- Best hit was 5lv3 chain D: Histone-arginine methyltransferase CARM1

| target | prob | fident | alnlen | evalue | theader |
|---|---|---|---|---|---|
| 5lv3-assembly1.cif.gz_D | 1 | 0.439 | 316 | 3.032e-46 | Crystal structure of mouse CARM1 in complex with ligand LH1561Br |
| 6arj-assembly1.cif.gz_A | 1 | 0.444 | 315 | 5.187e-46 | Crystal structure of CARM1 with EPZ022302 and SAH |
| 7qrd-assembly1.cif.gz_A | 1 | 0.428 | 338 | 2.046e-45 | Crystal structure of mouse CARM1 in complex with histone H3_10-25 |

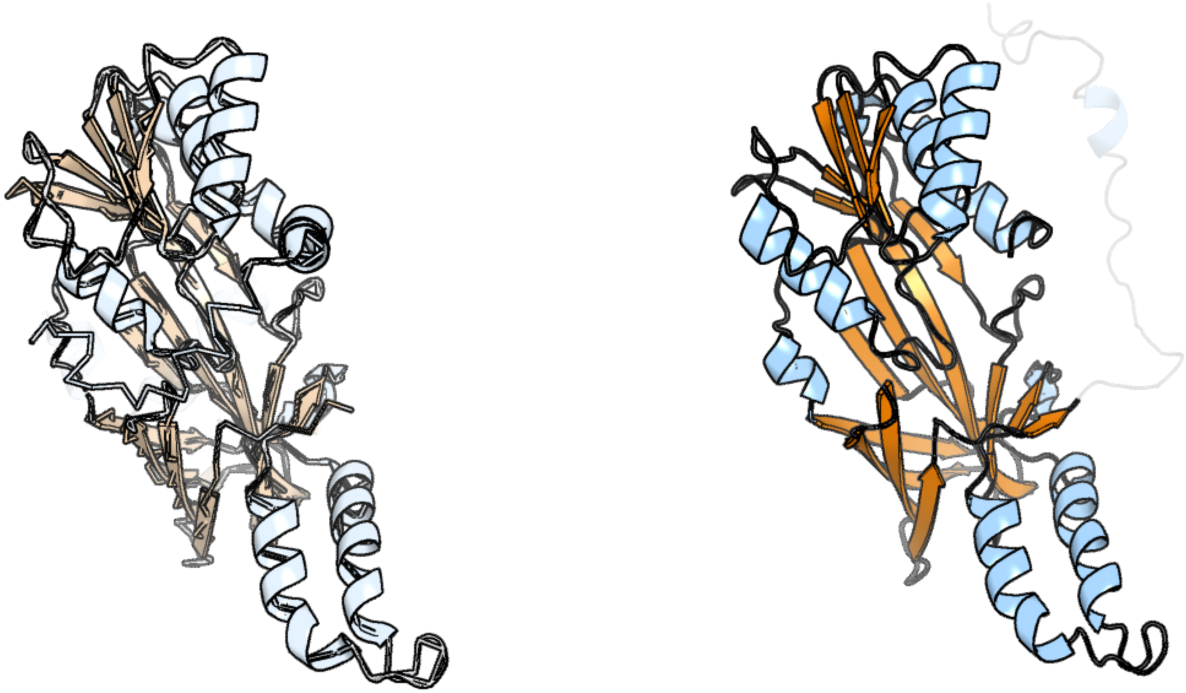

Figure 94: left: reference structure of 5lv3 chain D. right: predicted structure of kaa0168801, unaligned sequences are shown as transparent

#### kaa0169158

- Sequence-based annotation for kaa0169158 (predicted in DGTS from hits to PF13649) is hypothetical protein FNF28_02284 [Cafeteria roenbergensis]
- Best hit was 6h2v chain A: Methyltransferase-like protein 5

| target | prob | fident | alnlen | evalue | theader |
|---|---|---|---|---|---|
| 6h2v-assembly1.cif.gz_A | 1 | 0.364 | 214 | 3.121e-20 | Crystal structure of human METTL5-TRMT112 complex, the 18S rRNA m6A1832 methyltransferase at 2.5A resolution |
| 6h2u-assembly1.cif.gz_A | 1 | 0.352 | 213 | 1.816e-19 | Crystal structure of human METTL5-TRMT112 complex, the 18S rRNA m6A1832 methyltransferase at 1.6A resolution |
| 6h2v-assembly2.cif.gz_C | 1 | 0.328 | 216 | 1.689e-18 | Crystal structure of human METTL5-TRMT112 complex, the 18S rRNA m6A1832 methyltransferase at 2.5A resolution |

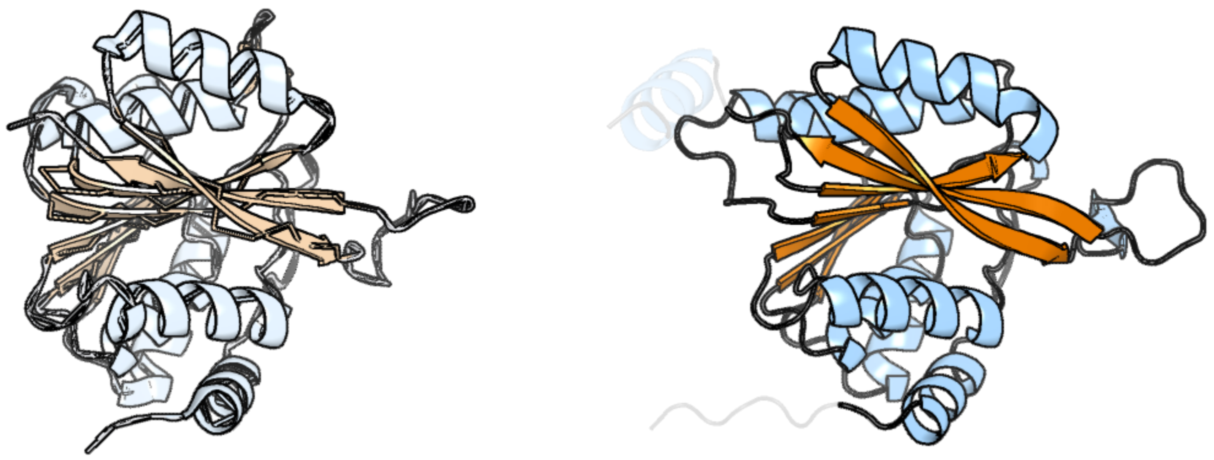

Figure 95: left: reference structure of 6h2v chain A. right: predicted structure of kaa0169158, unaligned sequences are shown as transparent

### kaa0169185

- Sequence-based annotation for kaa0169185 (predicted in Sulfonolipids from hits to PF00106, UniRef50_A0A383U0M1) is hypothetical protein FNF28_02310 [Cafeteria roenbergensis]
- Best hit was 6ihi chain A: Alclohol dehydrogenase

| target | prob | fident | alnlen | evalue | theader |
|---|---|---|---|---|---|
| 6ihi-assembly1.cif.gz_A | 1 | 0.296 | 260 | 1.094e-18 | Crystal structure of RasADH 3B3/I91V from Ralstonia.sp in complex with NADPH and A6O |
| 6ihi-assembly1.cif.gz_C | 1 | 0.286 | 265 | 1.094e-18 | Crystal structure of RasADH 3B3/I91V from Ralstonia.sp in complex with NADPH and A6O |
| 4bms-assembly2.cif.gz_E-2 | 1 | 0.3 | 260 | 1.31e-18 | Short chain alcohol dehydrogenase from Ralstonia sp. DSM 6428 in complex with NADPH |

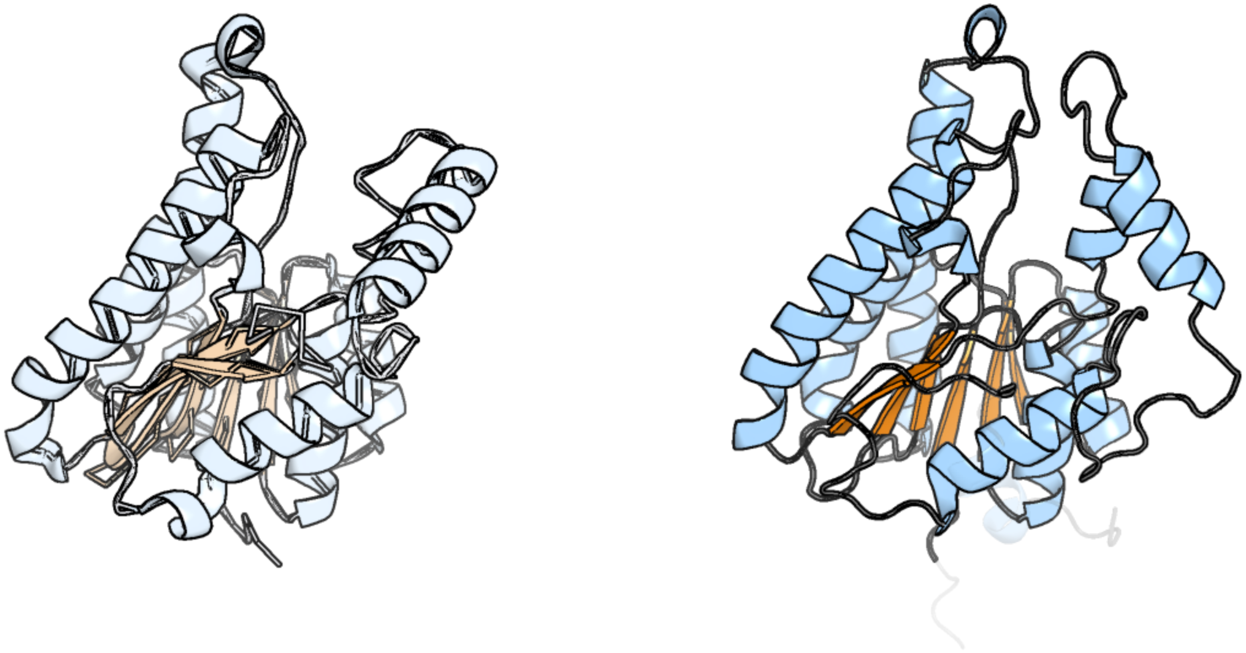

Figure 96: left: reference structure of 6ihi chain A. right: predicted structure of kaa0169185, unaligned sequences are shown as transparent

#### kaa0169376

- Sequence-based annotation for kaa0169376 (predicted in Sulfonolipids from hits to PF00155) is hypothetical protein FNF28_02157 [Cafeteria roenbergensis]
- Best hit was 5uts chain E: C-S Lyase Egt2

| target | prob | fident | alnlen | evalue | theader |
|---|---|---|---|---|---|
| 5uts-assembly3.cif.gz_E | 1 | 0.246 | 491 | 1.737e-27 | Carbon Sulfoxide lyase, Egt2 in the Ergothioneine biosynthesis pathway |
| 5uts-assembly1.cif.gz_G | 1 | 0.231 | 505 | 7.723e-27 | Carbon Sulfoxide lyase, Egt2 in the Ergothioneine biosynthesis pathway |
| 5v1x-assembly3.cif.gz_A | 1 | 0.238 | 494 | 8.56e-27 | Carbon Sulfoxide lyase, Egt2 Y134F in complex with its substrate |

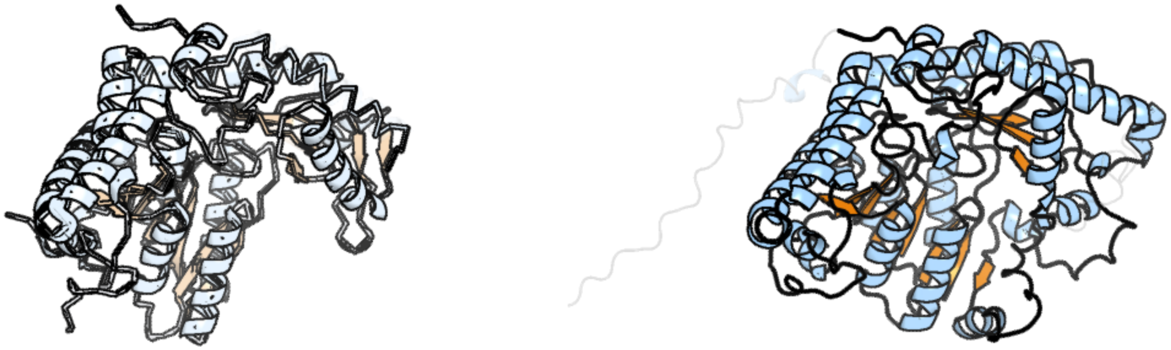

Figure 97: left: reference structure of 5uts chain E. right: predicted structure of kaa0169376, unaligned sequences are shown as transparent

#### kaa0169541

- Sequence-based annotation for kaa0169541 (predicted in Sulfonolipids from hits to PF00291) is hypothetical protein FNF28_01986 [Cafeteria roenbergensis]
- Best hit was 7ysk chain B: D-Cysteine desulfhydrase

| target | prob | fident | alnlen | evalue | theader |
|---|---|---|---|---|---|
| 7ysk-assembly1.cif.gz_B | 1 | 0.307 | 351 | 2.952e-24 | Crystal structure of D-Cysteine desulfhydrase from Pectobacterium atrosepticum |
| 7ysl-assembly1.cif.gz_B | 1 | 0.305 | 354 | 9.98e-24 | Crystal structure of D-Cysteine desulfhydrase with a trapped PLP-pyruvate geminal diamine |
| 1j0a-assembly1.cif.gz_A | 1 | 0.285 | 354 | 2.288e-22 | Crystal Structure Analysis of the ACC deaminase homologue |

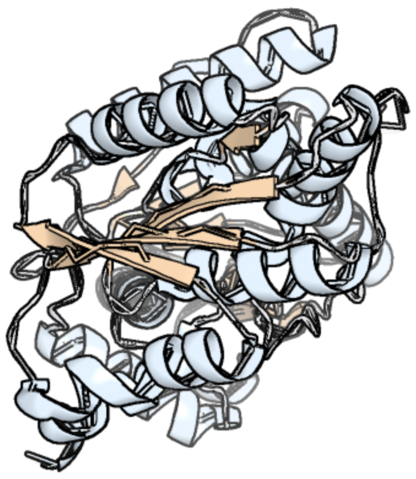
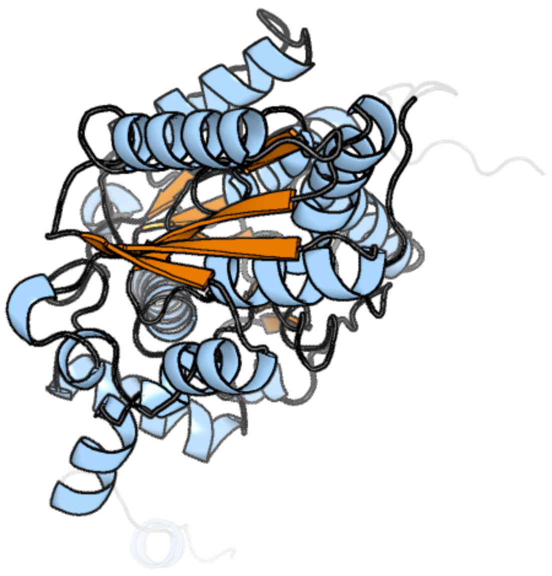

Figure 98: left: reference structure of 7ysk chain B. right: predicted structure of kaa0169541, unaligned sequences are shown as transparent

#### kaa0169592

- Sequence-based annotation for kaa0169592 (predicted in DGTS from hits to PF13649, UniRef50__A0A238JF33) is hypothetical protein FNF28__02036 [Cafeteria roenbergensis]
- Best hit was 3d2l chain A: SAM-dependent methyltransferase

| target | prob | fident | alnlen | evalue | theader |
|---|---|---|---|---|---|
| 3d2l-assembly1.cif.gz__A | 1 | 0.212 | 264 | 7.182e-19 | Crystal structure of SAM-dependent methyltransferase (ZP__00538691.1) from EXIGUOBACTERIUM SP. 255-15 at 1.90 A resolution |
| 7zkg-assembly1.cif.gz__A | 1 | 0.166 | 282 | 7.615e-19 | C-Methyltransferase PsmD from Streptomyces griseofuscus with bound cofactor (crystal form 2) |
| 1y8c-assembly1.cif.gz__A | 1 | 0.145 | 268 | 1.192e-17 | Crystal structure of a S-adenosylmethionine-dependent methyltransferase from Clostridium acetobutylicum ATCC 824 |

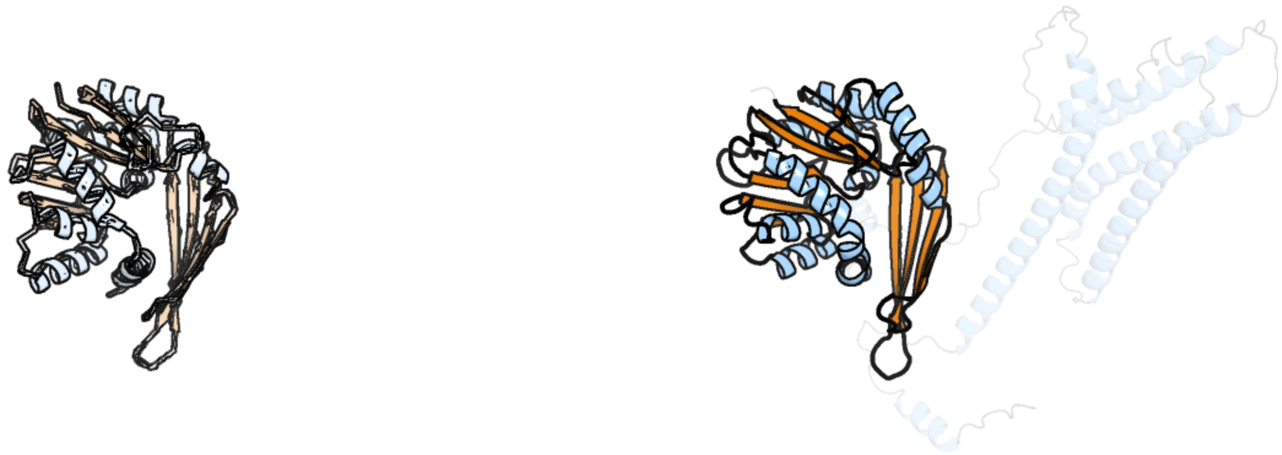

Figure 99: left: reference structure of 3d2l chain A. right: predicted structure of kaa0169592, unaligned sequences are shown as transparent

#### kaa0169679

- Sequence-based annotation for kaa0169679 (predicted in DGTS from hits to PF13649) is hypothetical protein FNF28_01957 [Cafeteria roenbergensis]
- Best hit was 7mqa chain NL: Probable dimethyladenosine transferase

| target | prob | fident | alnlen | evalue | theader |
|---|---|---|---|---|---|
| 7mqa-assembly1.cif.gz_NL | 1 | 0.64 | 278 | 5.149e-43 | Cryo-EM structure of the human SSU processome, state post-A1 |
| 6zqg-assembly1.cif.gz_JL | 1 | 0.537 | 283 | 3.346e-38 | Cryo-EM structure of the 90S pre-ribosome from Saccharomyces cerevisiae, state Dis-C |
| 2h1r-assembly2.cif.gz_B | 1 | 0.478 | 278 | 1.485e-36 | Crystal structure of a dimethyladenosine transferase from Plasmodium falciparum |

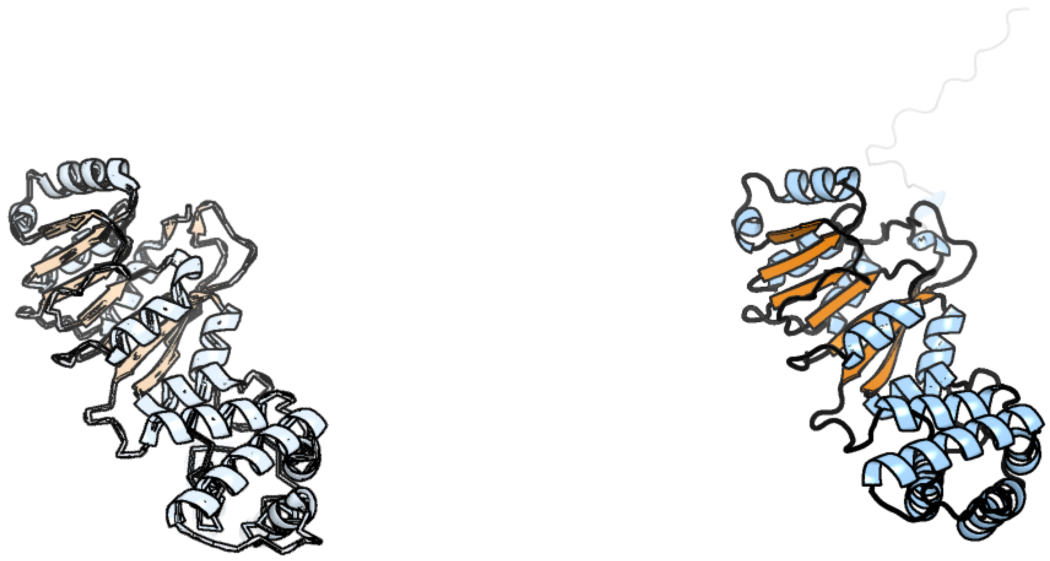

Figure 100: left: reference structure of 7mqa chain NL. right: predicted structure of kaa0169679, unaligned sequences are shown as transparent

#### kaa0169689

- Sequence-based annotation for kaa0169689 (predicted in DGTS from hits to PF13649) is hypothetical protein FNF28_01967 [Cafeteria roenbergensis]
- Best hit was 5una chain E: 7SK snRNA methylphosphate capping enzyme

| target | prob | fident | alnlen | evalue | theader |
|---|---|---|---|---|---|
| 5una-assembly5.cif.gz_E | 1 | 0.19 | 352 | 7.388e-14 | Fragment of 7SK snRNA methylphosphate capping enzyme |
| 5una-assembly6.cif.gz_F | 1 | 0.184 | 347 | 7.388e-14 | Fragment of 7SK snRNA methylphosphate capping enzyme |
| 6dcc-assembly1.cif.gz_A | 1 | 0.177 | 349 | 8.335e-14 | Structure of methylphosphate capping enzyme methyltransferase domain in complex with 5' end of 7SK RNA |

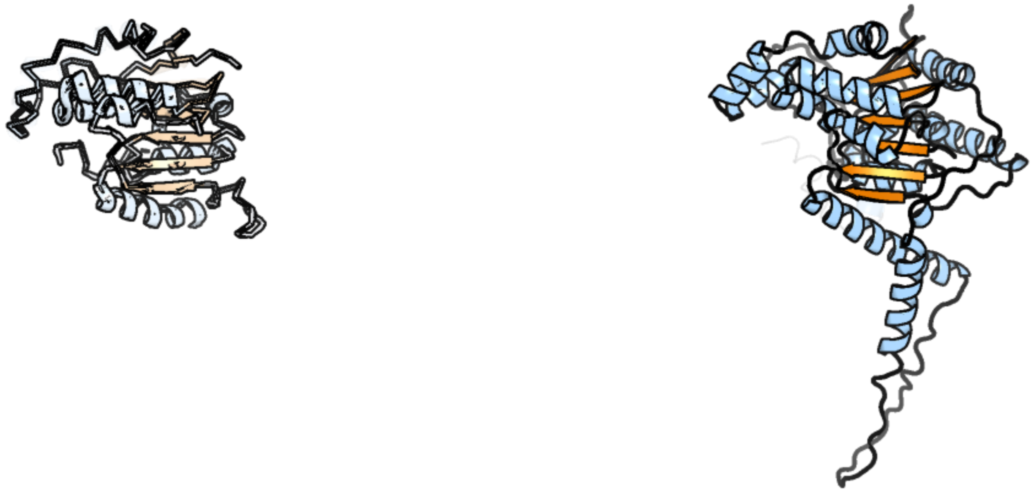

Figure 101: left: reference structure of 5una chain E. right: predicted structure of kaa0169689, unaligned sequences are shown as transparent

#### kaa0169966

- Sequence-based annotation for kaa0169966 (predicted in Sulfonolipids from hits to PF00155) is hypothetical protein FNF28_01756 [Cafeteria roenbergensis]
- Best hit was 5veh chain B: Kynurenine aminotransferase

| target | prob | fident | alnlen | evalue | theader |
|---|---|---|---|---|---|
| 5veh-assembly1.cif.gz_B | 1 | 0.354 | 412 | 1.446e-35 | Re-refinement OF THE PDB STRUCTURE 1yiz of Aedes aegypti kynurenine aminotransferase |
| 5veh-assembly1.cif.gz_A | 1 | 0.346 | 419 | 1.691e-35 | Re-refinement OF THE PDB STRUCTURE 1yiz of Aedes aegypti kynurenine aminotransferase |
| 5veq-assembly1.cif.gz_A | 1 | 0.363 | 421 | 8.536e-35 | MOUSE KYNURENINE AMINOTRANSFERASE III, RE-REFINEMENT OF THE PDB STRUCTURE 3E2Y |

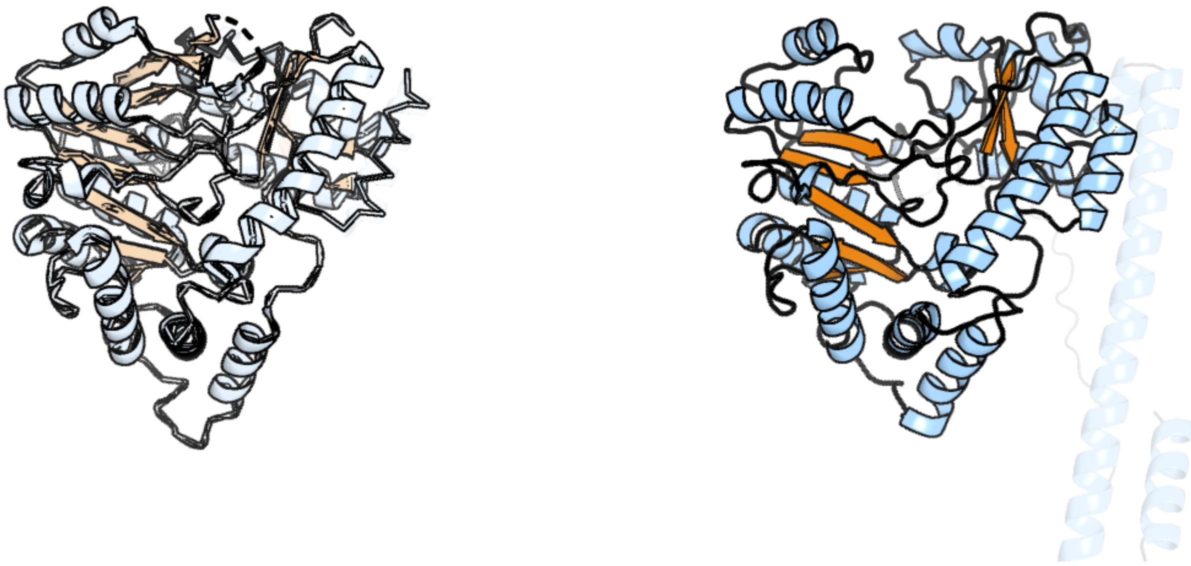

Figure 102: left: reference structure of 5veh chain B. right: predicted structure of kaa0169966, unaligned sequences are shown as transparent

### kaa0170054

- Sequence-based annotation for kaa0170054 (predicted in Fu-FA,DGTS from hits to PF02353, PF13649, UniRef50_A0A1V0RIH1) is hypothetical protein FNF28_01663 [Cafeteria roenbergensis]
- Best hit was 5z9o chain A: Cyclopropane-fatty-acyl-phospholipid synthase

| target | prob | fident | alnlen | evalue | theader |
|---|---|---|---|---|---|
| 5z9o-assembly1.cif.gz_A | 1 | 0.253 | 398 | 3.32e-21 | The crystal structure of Cyclopropane-fatty-acyl-phospholipid synthase from Lactobacillus acidophilus |
| 5z9o-assembly2.cif.gz_B | 1 | 0.254 | 393 | 6.693e-21 | The crystal structure of Cyclopropane-fatty-acyl-phospholipid synthase from Lactobacillus acidophilus |
| 7qos-assembly2.cif.gz_B | 1 | 0.242 | 409 | 4.961e-20 | Cyclopropane fatty acid synthase from Aquifex aeolicous with bound ligands |

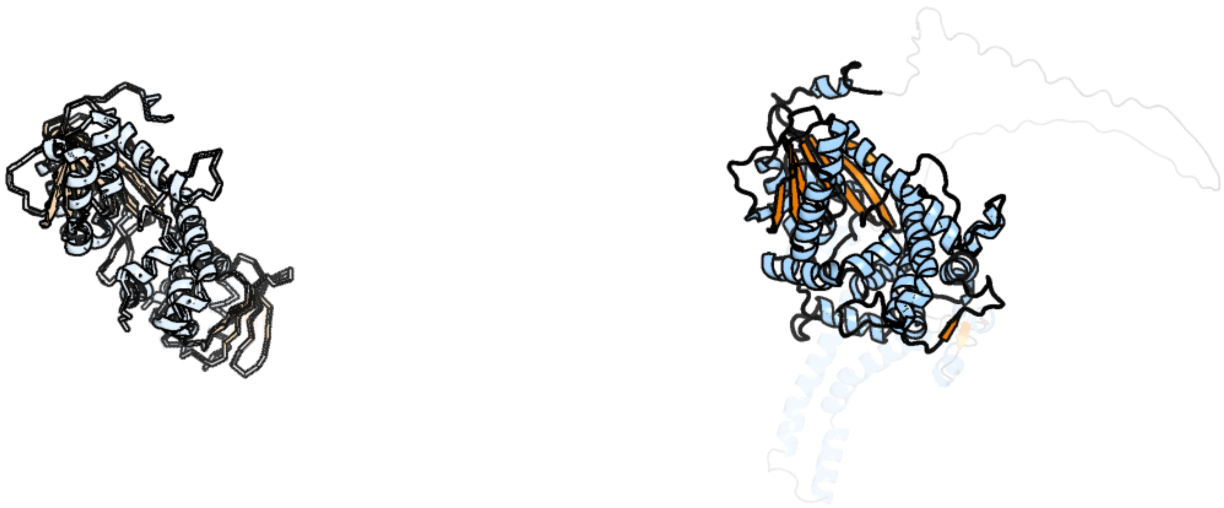

Figure 103: left: reference structure of 5z9o chain A. right: predicted structure of kaa0170054, unaligned sequences are shown as transparent

### kaa0170162

- Sequence-based annotation for kaa0170162 (predicted in Fu-FA from hits to PF01593) is hypothetical protein FNF28_01583 [Cafeteria roenbergensis]
- Best hit was 4rep chain A: Gamma-carotene desaturase

| target | prob | fident | alnlen | evalue | theader |
|---|---|---|---|---|---|
| 4rep-assembly1.cif.gz_A | 1 | 0.136 | 643 | 2.157e-19 | Crystal Structure of gamma-carotenoid desaturase |
| 3ka7-assembly1.cif.gz_A | 1 | 0.144 | 631 | 6.639e-16 | Crystal Structure of an oxidoreductase from Methanosarcina mazei. Northeast Structural Genomics Consortium target id MaR208 |
| 4dgk-assembly1.cif.gz_A | 1 | 0.153 | 630 | 9.078e-16 | Crystal structure of Phytoene desaturase CRTI from Pantoea ananatis |

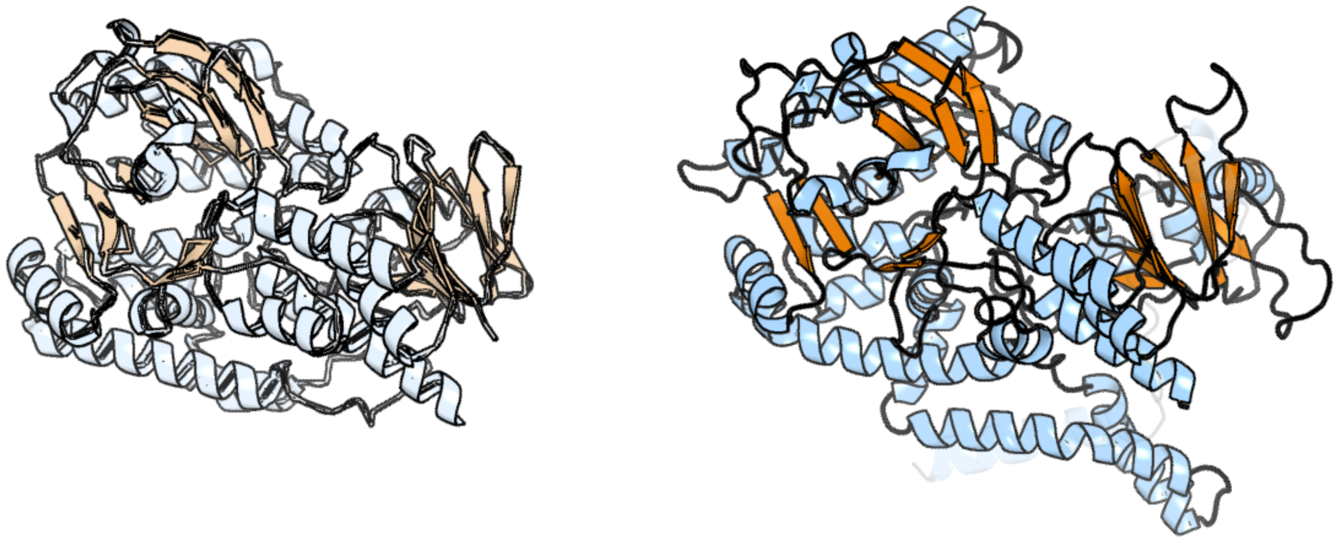

Figure 104: left: reference structure of 4rep chain A. right: predicted structure of kaa0170162, unaligned sequences are shown as transparent

### kaa0170308

- Sequence-based annotation for kaa0170308 (predicted in Sulfonolipids from hits to PF00155) is hypothetical protein FNF28_01536 [Cafeteria roenbergensis]
- Best hit was 1b8g chain B: PROTEIN (1-AMINOCYCLOPROPANE-1-CARBOXYLATE SYNTHASE)

| target | prob | fident | alnlen | evalue | theader |
|---|---|---|---|---|---|
| 1b8g-assembly1.cif.gz_B | 1 | 0.297 | 461 | 3.915e-38 | 1-AMINOCYCLOPROPANE-1-CARBOXYLATE SYNTHASE |
| 7dlw-assembly1.cif.gz_B | 1 | 0.284 | 460 | 5.722e-38 | Crystal structure of Arabidopsis ACS7 in complex with PPG |
| 7dlw-assembly2.cif.gz_D | 1 | 0.28 | 456 | 8.827e-38 | Crystal structure of Arabidopsis ACS7 in complex with PPG |

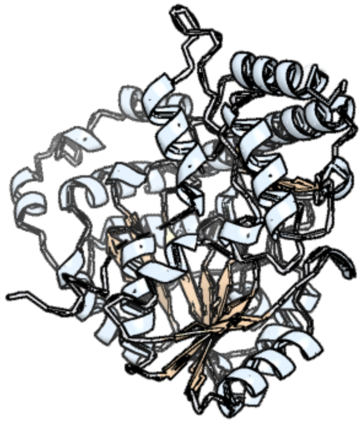

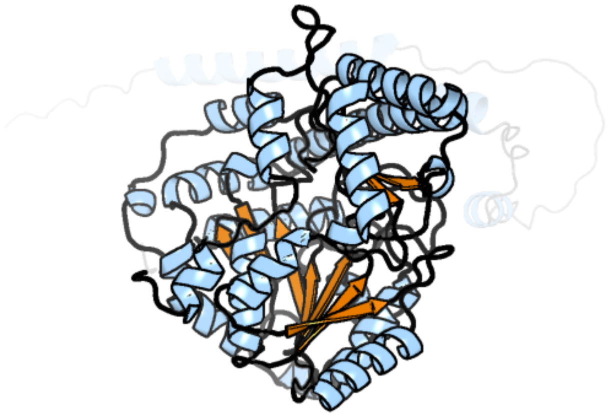

Figure 105: left: reference structure of 1b8g chain B. right: predicted structure of kaa0170308, unaligned sequences are shown as transparent

### kaa0170430

- Sequence-based annotation for kaa0170430 (predicted in Sulfonolipids from hits to PF00106) is hypothetical protein FNF28_01424 [Cafeteria roenbergensis]
- Best hit was 3wj7 chain C: Putative oxidoreductase

| target | prob | fident | alnlen | evalue | theader |
|---|---|---|---|---|---|
| 3wj7-assembly1.cif.gz_C | 1 | 0.234 | 367 | 3.099e-19 | Crystal structure of gox2253 |
| 6kv9-assembly1.cif.gz_A-2 | 1 | 0.214 | 369 | 7.669e-19 | MoeE5 in complex with UDP-glucuronic acid and NAD |
| 7ys8-assembly1.cif.gz_B | 1 | 0.196 | 371 | 3.538e-18 | Crystal Structure of UDP-glucose 4-epimerase (Rv3634c) from Mycobacterium tuberculosis |

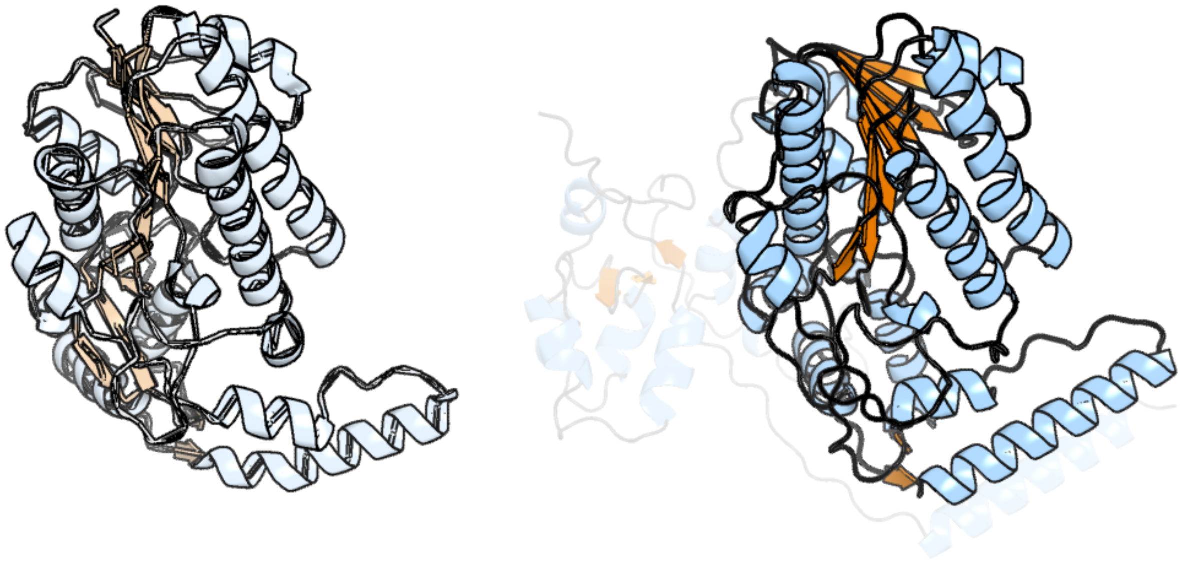

Figure 106: left: reference structure of 3wj7 chain C. right: predicted structure of kaa0170430, unaligned sequences are shown as transparent

### kaa0170472

- Sequence-based annotation for kaa0170472 (predicted in DGTS from hits to PF13649) is hypothetical protein FNF28_01466 [Cafeteria roenbergensis]
- Best hit was 2o05 chain B: Spermidine synthase

| target | prob | fident | alnlen | evalue | theader |
|---|---|---|---|---|---|
| 2o05-assembly1.cif.gz_B | 1 | 0.469 | 298 | 2.255e-36 | Human spermidine synthase |
| 6o64-assembly2.cif.gz_C | 1 | 0.421 | 301 | 3.003e-36 | Crystal Structure of Arabidopsis thaliana Spermidine Synthase isoform 2 (AtSPDS2) |
| 6o64-assembly1.cif.gz_B | 1 | 0.411 | 304 | 7.09e-36 | Crystal Structure of Arabidopsis thaliana Spermidine Synthase isoform 2 (AtSPDS2) |

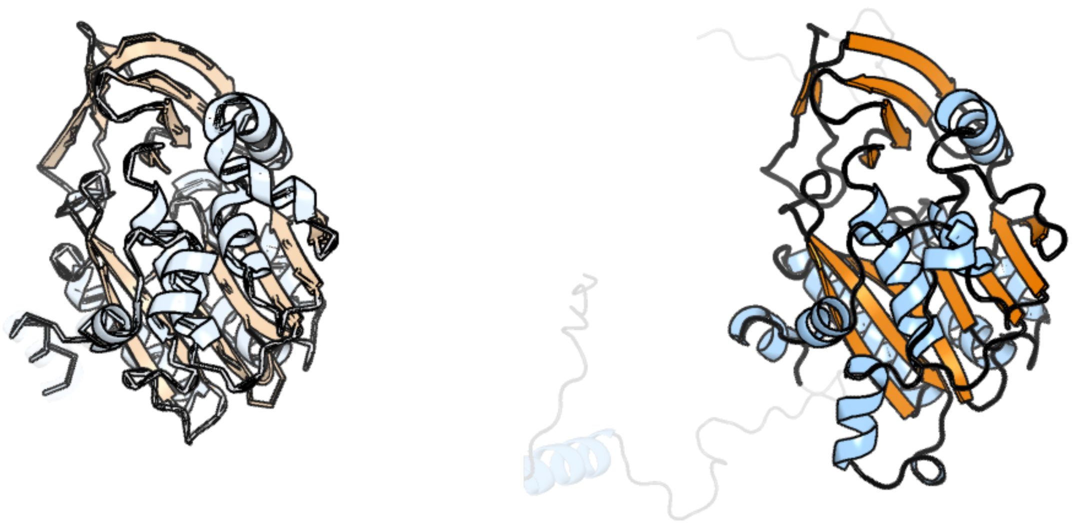

Figure 107: left: reference structure of 2o05 chain B. right: predicted structure of kaa0170472, unaligned sequences are shown as transparent

#### kaa0170620

- Sequence-based annotation for kaa0170620 (predicted in Fu-FA from hits to PF06966) is hypothetical protein FNF28_01382 [Cafeteria roenbergensis]
- No significant structural hit found

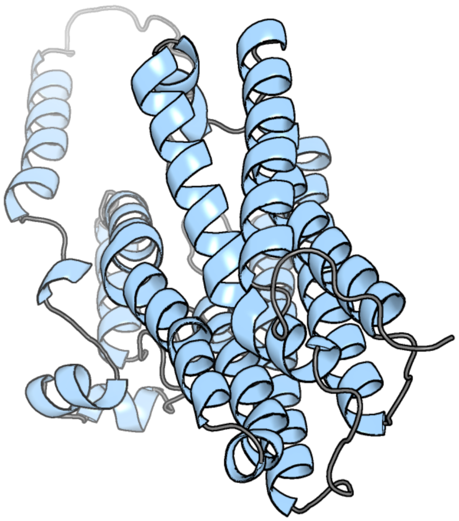

Figure 108: predicted structure of kaa0170620

### kaa0170685

- Sequence-based annotation for kaa0170685 (predicted in Sulfonolipids from hits to PF00155) is hypothetical protein FNF28_01230 [Cafeteria roenbergensis]
- Best hit was 8bj4 chain D: histidinol-phosphate aminotransferase

| target | prob | fident | alnlen | evalue | theader |
|---|---|---|---|---|---|
| 8bj4-assembly2.cif.gz_D | 1 | 0.358 | 385 | 2.887e-32 | Crystal structure of Medicago truncatula histidinol-phosphate aminotransferase (HISN6) in apo form |
| 6fwh-assembly1.cif.gz_A | 1 | 0.6 | 195 | 1.188e-24 | Acanthamoeba IGPD in complex with R-C348 to 1.7A resolution |
| 4r8d-assembly1.cif.gz_A | 1 | 0.262 | 408 | 1.28e-23 | Crystal structure of Rv1600 encoded aminotransferase in complex with PLP-MES from Mycobacterium tuberculosis |

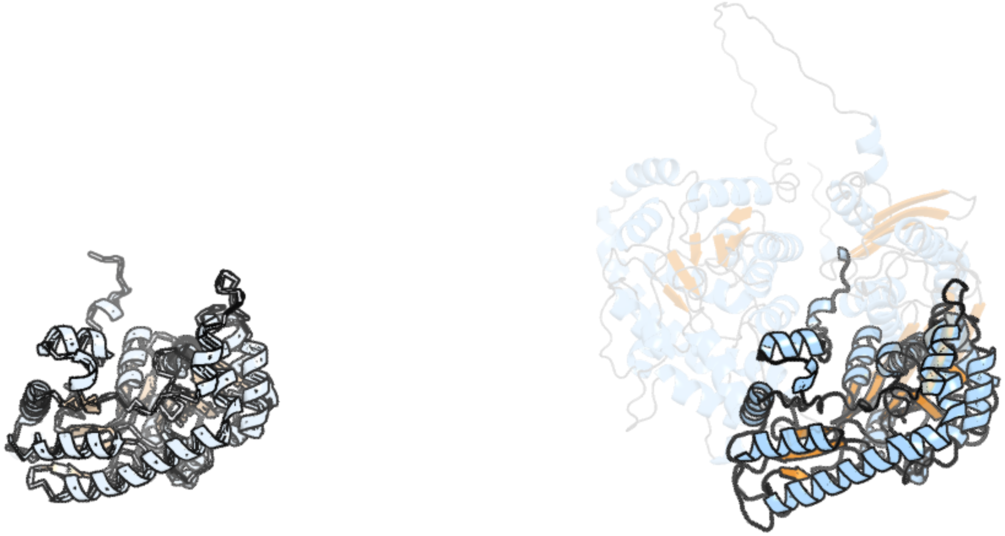

Figure 109: left: reference structure of 8bj4 chain D. right: predicted structure of kaa0170685, unaligned sequences are shown as transparent

### kaa0170833

- Sequence-based annotation for kaa0170833 (predicted in Sulfonolipids from hits to PF00106, UniRef50_A0A383U0M1) is hypothetical protein FNF28_01106 [Cafeteria roenbergensis]
- Best hit was 3wxb chain B: Uncharacterized protein

| target | prob | fident | alnlen | evalue | theader |
|---|---|---|---|---|---|
| 3wxb-assembly1.cif.gz_B | 1 | 0.264 | 242 | 1.087e-14 | Crystal structure of NADPH bound carbonyl reductase from chicken fatty liver |
| 1sny-assembly1.cif.gz_A-2 | 1 | 0.227 | 242 | 1.551e-14 | Carbonyl reductase Sniffer of D. melanogaster |
| 5idw-assembly1.cif.gz_B | 1 | 0.24 | 237 | 1.032e-13 | Crystal structure of an oxidoreductase from Burkholderia vietnamiensis in complex with NADP |

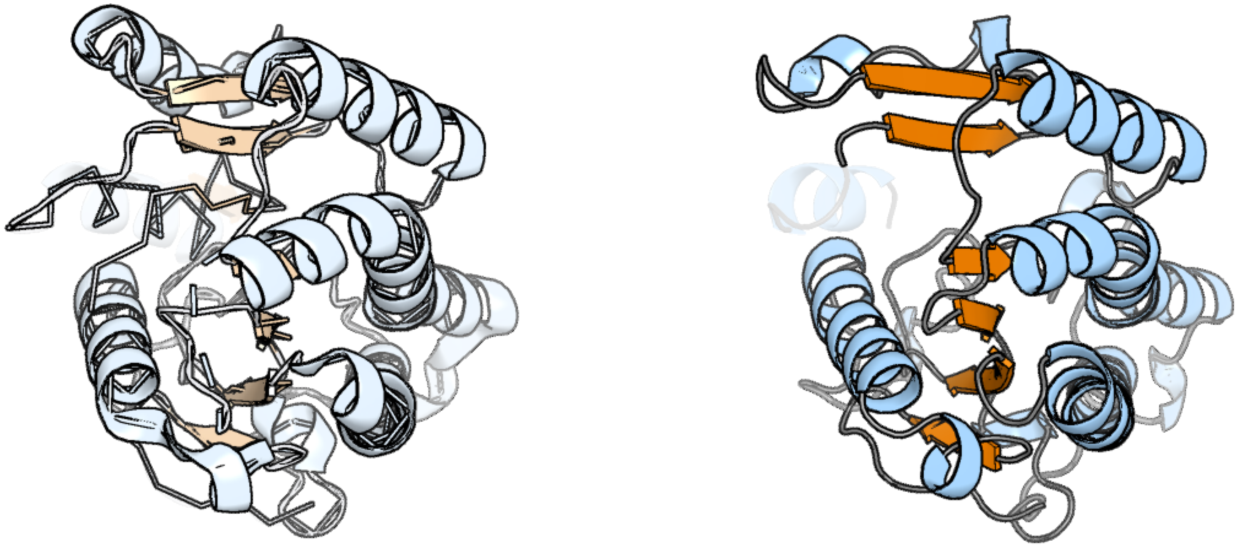

Figure 110: left: reference structure of 3wxb chain B. right: predicted structure of kaa0170833, unaligned sequences are shown as transparent

### kaa0170835

- Sequence-based annotation for kaa0170835 (predicted in DGTS from hits to PF13649) is hypothetical protein FNF28_01108 [Cafeteria roenbergensis]
- Best hit was 7y9c chain A: Protein-L-histidine N-pros-methyltransferase

| target | prob | fident | alnlen | evalue | theader |
|---|---|---|---|---|---|
| 7y9c-assembly1.cif.gz_A | 1 | 0.356 | 202 | 9.178e-17 | Crystal structure of METTL9 in complex with SLC39A5 peptide and SAH |
| 8gzf-assembly1.cif.gz_A | 1 | 0.361 | 199 | 1.402e-16 | Crystal Structure of METTL9-SAH |
| 8bvi-assembly1.cif.gz_A | 1 | 0.314 | 216 | 5.977e-15 | Crystal structure of the METTL9-like histidine methyltransferase from Ostreococcus tauri |

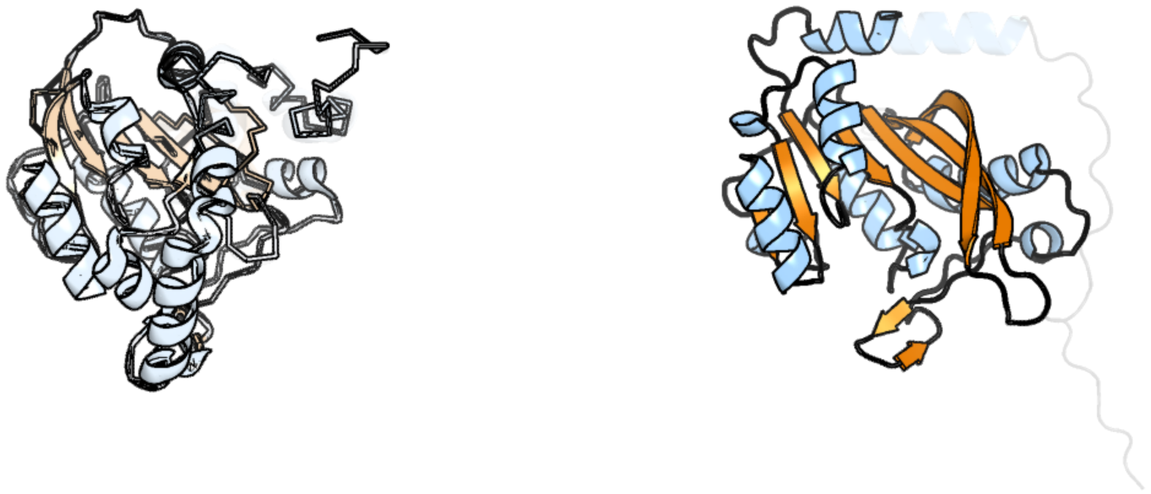

Figure 111: left: reference structure of 7y9c chain A. right: predicted structure of kaa0170835, unaligned sequences are shown as transparent

### kaa0170843

- Sequence-based annotation for kaa0170843 (predicted in DGTS from hits to PF13649) is hypothetical protein FNF28_01116 [Cafeteria roenbergensis]
- Best hit was 2pxx chain A: Uncharacterized protein MGC2408

| target | prob | fident | alnlen | evalue | theader |
|---|---|---|---|---|---|
| 2pxx-assembly1.cif.gz_A | 1 | 0.262 | 221 | 1.28e-14 | Human putative methyltransferase MGC2408 |
| 5wcj-assembly1.cif.gz_A | 1 | 0.218 | 311 | 1.775e-11 | Crystal Structure of Human Methyltransferase-like protein 13 in complex with SAH |
| 4nec-assembly3.cif.gz_C | 1 | 0.178 | 230 | 2.099e-07 | Conversion of a Disulfide Bond into a Thioacetal Group during Echinomycin Biosynthesis |

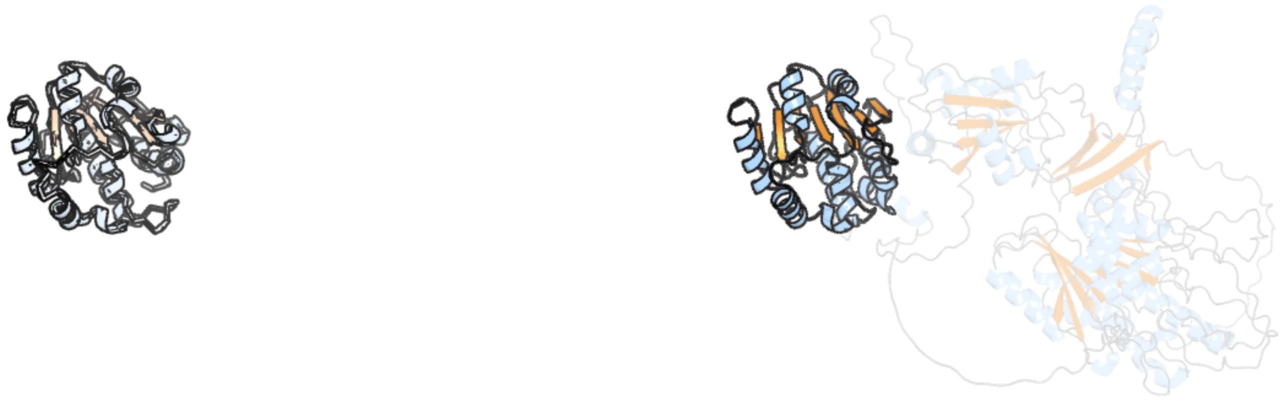

Figure 112: left: reference structure of 2pxx chain A. right: predicted structure of kaa0170843, unaligned sequences are shown as transparent

#### kaa0170858

- Sequence-based annotation for kaa0170858 (predicted in Sulfonolipids from hits to PF00106, UniRef50_A0A383U0M1) is hypothetical protein FNF28_01131 [Cafeteria roenbergensis]
- Best hit was 3pk0 chain C: Short-chain dehydrogenase/reductase SDR

| target | prob | fident | alnlen | evalue | theader |
|---|---|---|---|---|---|
| 3pk0-assembly1.cif.gz_C | 1 | 0.208 | 259 | 7.139e-18 | Crystal structure of Short-chain dehydrogenase/reductase SDR from Mycobacterium smegmatis |
| 6ihi-assembly1.cif.gz_C | 1 | 0.275 | 232 | 9.699e-18 | Crystal structure of RasADH 3B3/I91V from Ralstonia.sp in complex with NADPH and A6O |
| 4dry-assembly1.cif.gz_A | 1 | 0.199 | 256 | 1.031e-17 | The crystal structure of 3-oxoacyl-[acyl-carrier-protein] reductase from Rhizobium meliloti |

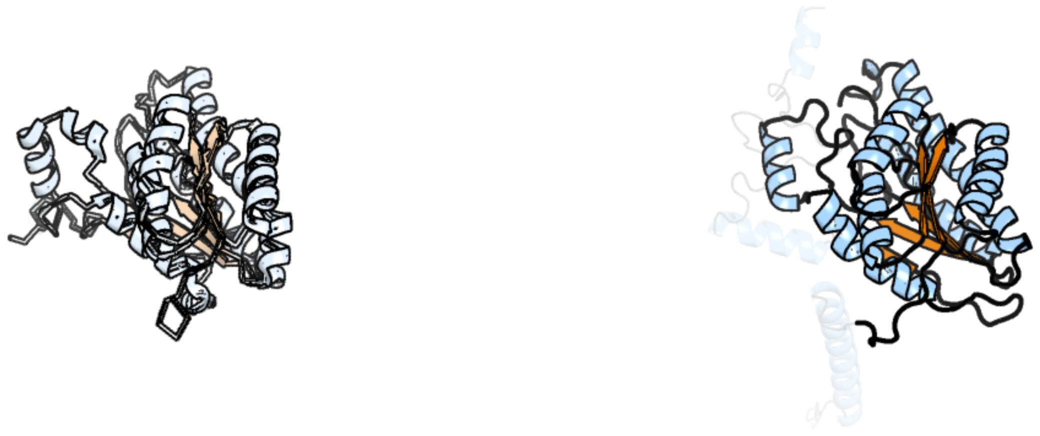

Figure 113: left: reference structure of 3pk0 chain C. right: predicted structure of kaa0170858, unaligned sequences are shown as transparent

#### kaa0171011

- Sequence-based annotation for kaa0171011 (predicted in Sulfonolipids from hits to PF00106, UniRef50_A0A383U0M1) is hypothetical protein FNF28_01016 [Cafeteria roenbergensis]
- Best hit was 5itv chain A: Dihydroanticapsin 7-dehydrogenase

| target | prob | fident | alnlen | evalue | theader |
|---|---|---|---|---|---|
| 5itv-assembly1.cif.gz_A | 1 | 0.338 | 254 | 5.56e-27 | Crystal structure of Bacillus subtilis BacC Dihydroanticapsin 7-dehydrogenase in complex with NADH |
| 2d1y-assembly1.cif.gz_C | 1 | 0.394 | 256 | 5.363e-26 | Crystal structure of TT0321 from Thermus thermophilus HB8 |
| 2d1y-assembly1.cif.gz_B | 1 | 0.386 | 256 | 1.526e-25 | Crystal structure of TT0321 from Thermus thermophilus HB8 |

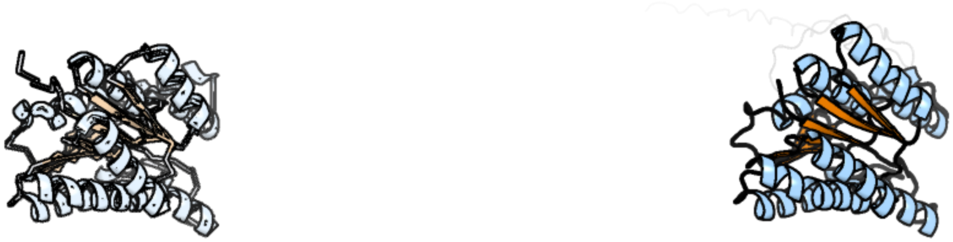

Figure 114: left: reference structure of 5itv chain A. right: predicted structure of kaa0171011, unaligned sequences are shown as transparent

### kaa0171081

- Sequence-based annotation for kaa0171081 (predicted in DGTS from hits to PF13649) is hypothetical protein FNF28_01086 [Cafeteria roenbergensis]
- Best hit was 6v0p chain A: Protein arginine N-methyltransferase 5

| target | prob | fident | alnlen | evalue | theader |
| --- | --- | --- | --- | --- | --- |
| 6v0p-assembly1.cif.gz_A | 1 | 0.396 | 771 | 2.607e-81 | PRMT5 complex bound to covalent PBM inhibitor BRD6711 |
| 8g1u-assembly1.cif.gz_I | 1 | 0.398 | 767 | 6.505e-81 | Structure of the methylosome-Lsm10/11 complex |
| 7uy1-assembly1.cif.gz_A | 1 | 0.395 | 771 | 8.175e-81 | HUMAN PRMT5:MEP50 COMPLEX WITH MTA and Fragment 5 Bound |

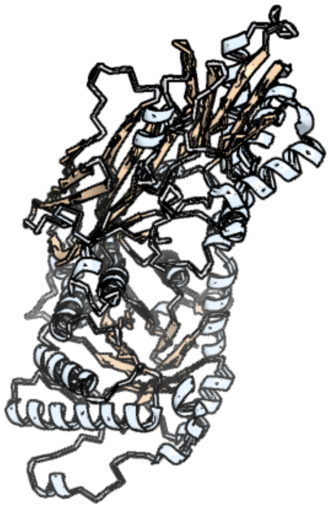

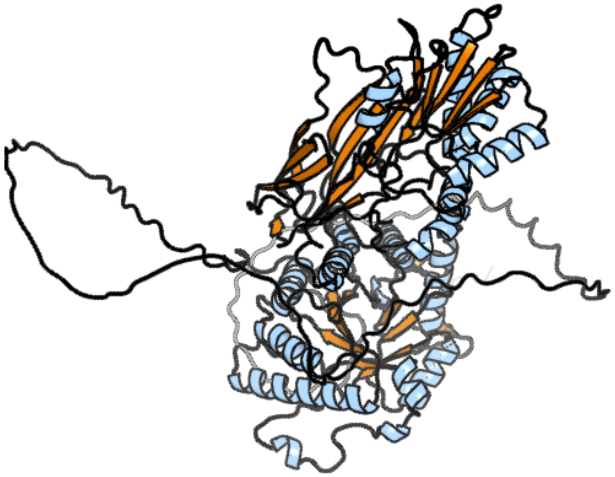

Figure 115: left: reference structure of 6v0p chain A. right: predicted structure of kaa0171081, unaligned sequences are shown as transparent

### kaa0171141

- Sequence-based annotation for kaa0171141 (predicted in Sulfonolipids from hits to PF00106) is hypothetical protein FNF28_00908 [Cafeteria roenbergensis]
- Best hit was 7ys8 chain B: UDP-glucose 4-epimerase

| target | prob | fident | alnlen | evalue | theader |
|---|---|---|---|---|---|
| 7ys8-assembly1.cif.gz_B | 1 | 0.207 | 405 | 1.695e-17 | Crystal Structure of UDP-glucose 4-epimerase (Rv3634c) from Mycobacterium tuberculosis |
| 3wj7-assembly1.cif.gz_C | 1 | 0.201 | 406 | 2.785e-16 | Crystal structure of gox2253 |
| 4lw8-assembly1.cif.gz_B | 1 | 0.27 | 277 | 3.645e-16 | Crystal structure of a putative epimerase from Burkholderia cenocepacia J2315 |

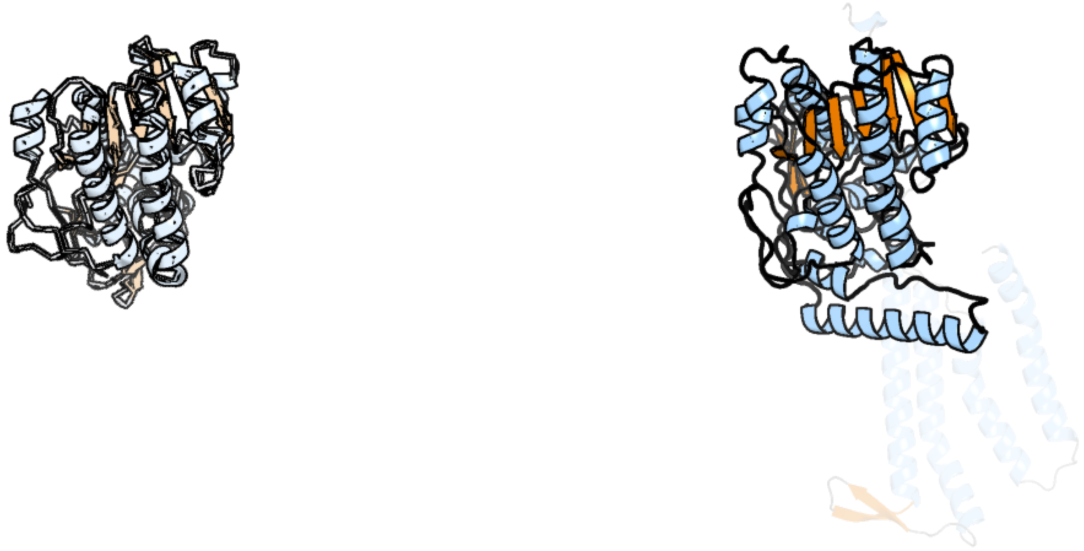

Figure 116: left: reference structure of 7ys8 chain B. right: predicted structure of kaa0171141, unaligned sequences are shown as transparent

### kaa0171150

- Sequence-based annotation for kaa0171150 (predicted in DGTS from hits to PF13649) is hypothetical protein FNF28_00917 [Cafeteria roenbergensis]
- Best hit was 7f1e chain A: tRNA N(3)-methylcytidine methyltransferase METTL6

| target | prob | fident | alnlen | evalue | theader |
|---|---|---|---|---|---|
| 7f1e-assembly1.cif.gz_A | 1 | 0.184 | 363 | 2.164e-16 | Structure of METTL6 bound with SAM |
| 7ezg-assembly1.cif.gz_A | 1 | 0.198 | 362 | 2.157e-15 | The structure of the human METTL6 enzyme in complex with SAH |
| 7f1e-assembly2.cif.gz_B | 1 | 0.184 | 363 | 4.198e-15 | Structure of METTL6 bound with SAM |

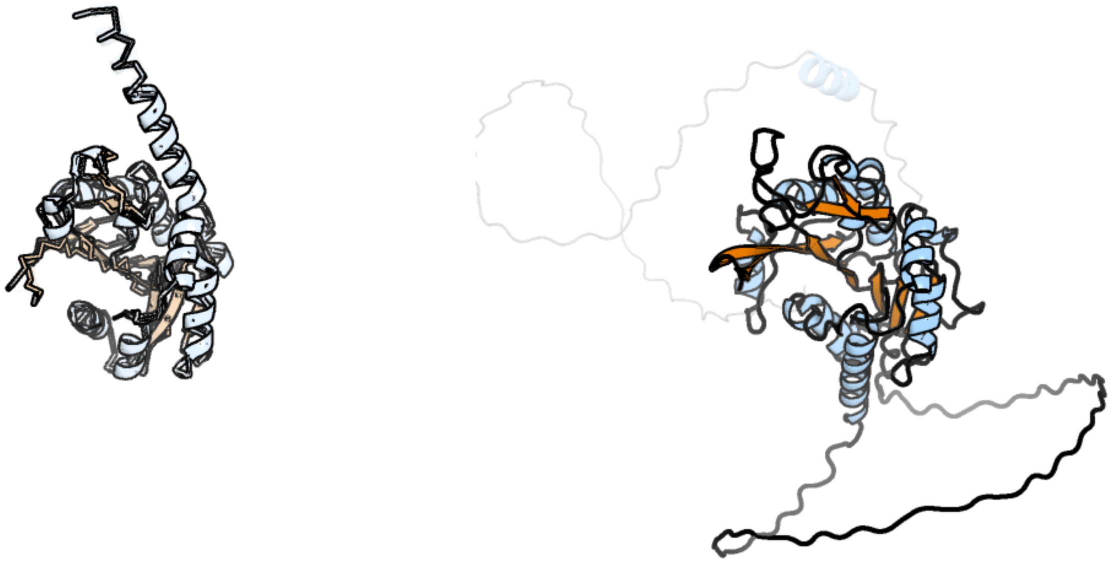

Figure 117: left: reference structure of 7f1e chain A. right: predicted structure of kaa0171150, unaligned sequences are shown as transparent

### kaa0171167

- Sequence-based annotation for kaa0171167 (predicted in Fu-FA from hits to PF06966) is hypothetical protein FNF28_00933 [Cafeteria roenbergensis]
- Best hit was 7w3y chain A: Isoform 2 of Potassium voltage-gated channel subfamily D member 3

| target | prob | fident | alnlen | evalue | theader |
|---|---|---|---|---|---|
| 7w3y-assembly1.cif.gz_A | 1 | 0.29 | 224 | 7.317e-10 | CryoEM structure of human Kv4.3 |
| 7f0j-assembly1.cif.gz_B | 1 | 0.323 | 198 | 8.011e-10 | CryoEM structure of human Kv4.2 |
| 5wie-assembly2.cif.gz_H | 1 | 0.341 | 196 | 9.605e-10 | Crystal structure of a Kv1.2-2.1 chimera K+ channel V406W mutant in an inactivated state |

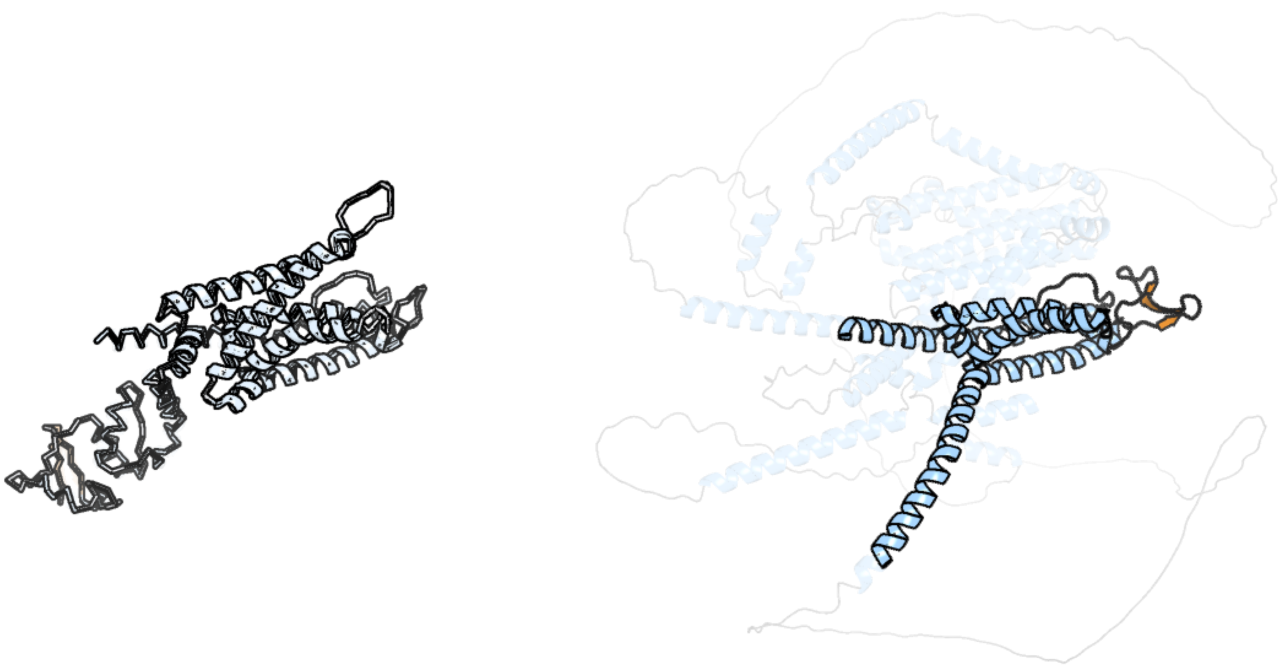

Figure 118: left: reference structure of 7w3y chain A. right: predicted structure of kaa0171167, unaligned sequences are shown as transparent

#### kaa0171345

- Sequence-based annotation for kaa0171345 (predicted in DGTS from hits to PF13649) is hypothetical protein FNF28_00836 [Cafeteria roenbergensis]
- Best hit was 5e9w chain C: mRNA cap guanine-N7 methyltransferase

| target | prob | fident | alnlen | evalue | theader |
|---|---|---|---|---|---|
| 5e9w-assembly3.cif.gz_C | 1 | 0.188 | 473 | 9.481e-18 | Crystal structure of mRNA cap guanine-N7 methyltransferase obtained by limited proteolysis |
| 3bgv-assembly3.cif.gz_C | 1 | 0.203 | 466 | 2.216e-17 | Crystal structure of mRNA cap guanine-N7 methyltransferase in complex with SAH |
| 5e9w-assembly2.cif.gz_B | 1 | 0.199 | 467 | 3.112e-17 | Crystal structure of mRNA cap guanine-N7 methyltransferase obtained by limited proteolysis |

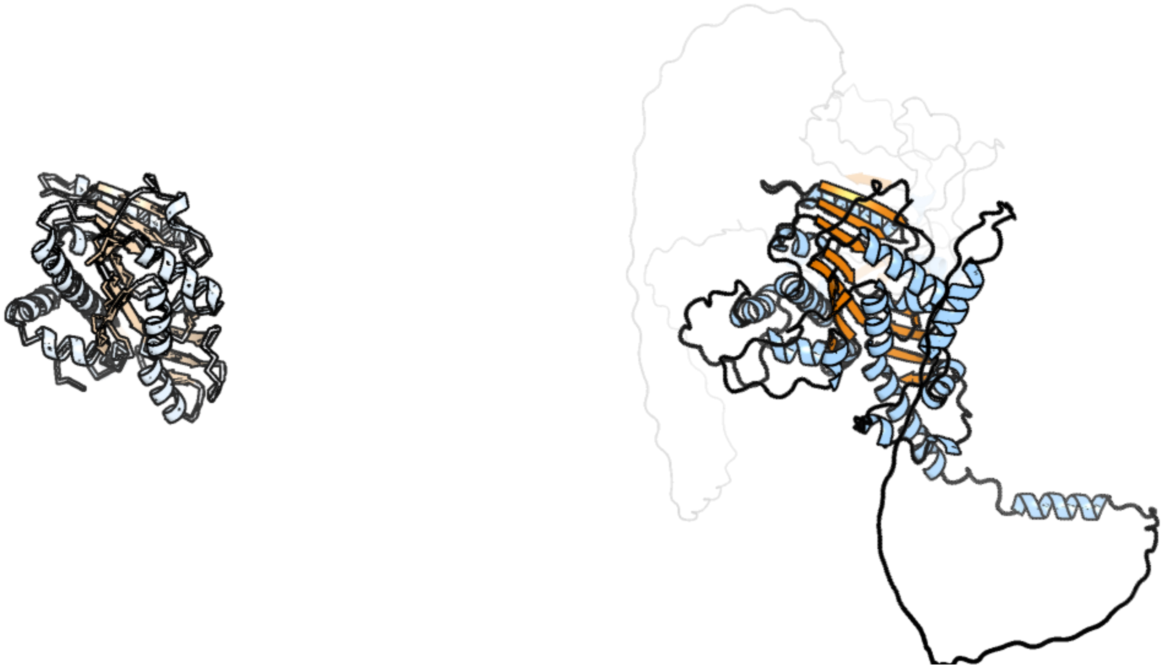

Figure 119: left: reference structure of 5e9w chain C. right: predicted structure of kaa0171345, unaligned sequences are shown as transparent

#### kaa0171444

- Sequence-based annotation for kaa0171444 (predicted in DGTS from hits to PF13649) is hypothetical protein FNF28_00656 [Cafeteria roenbergensis]
- Best hit was 2pxx chain A: Uncharacterized protein MGC2408

| target | prob | fident | alnlen | evalue | theader |
|---|---|---|---|---|---|
| 2pxx-assembly1.cif.gz_A | 1 | 0.276 | 217 | 1.116e-19 | Human putative methyltransferase MGC2408 |
| 3sm3-assembly1.cif.gz_A-2 | 1 | 0.13 | 223 | 5.895e-11 | Crystal Structure of SAM-dependent methyltransferases Q8PUK2_METMA from Methanosarcina mazei. Northeast Structural Genomics Consortium Target MaR262. |
| 7ndm-assembly1.cif.gz_A | 1 | 0.174 | 229 | 3.292e-10 | Crystal structure of the heterocyclic toxin methyltransferase from Mycobacterium tuberculosis with bound substrate 4-hydroxyisoquinolin-1(2H)-one |

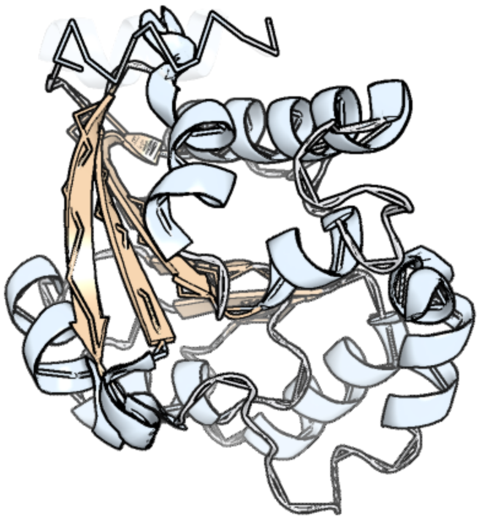

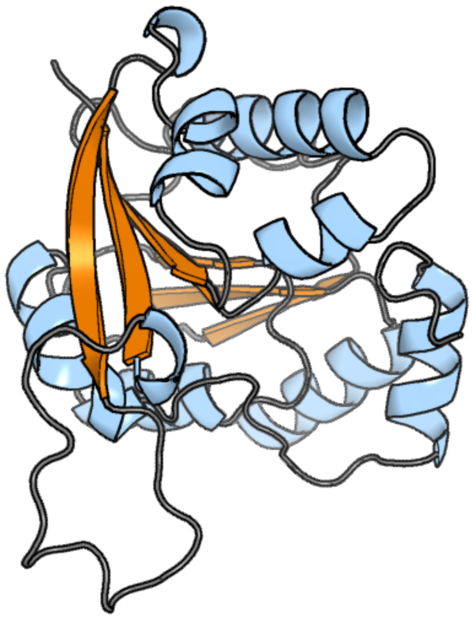

Figure 120: left: reference structure of 2pxx chain A. right: predicted structure of kaa0171444, unaligned sequences are shown as transparent

### kaa0171457

- Sequence-based annotation for kaa0171457 (predicted in PE_PMME from hits to PF01066) is hypothetical protein FNF28_00669 [Cafeteria roenbergensis]
- Best hit was 7b1l chain A: CDP-diacylglycerol–serine O-phosphatidyltransferase

| target | prob | fident | alnlen | evalue | theader |
|---|---|---|---|---|---|
| 7b1l-assembly1.cif.gz_A | 1 | 0.215 | 172 | 3.45e-06 | Crystal structure of phosphatidyl serine synthase (PSS) in the closed conformation with bound citrate. |
| 6h59-assembly1.cif.gz_A | 0.999 | 0.192 | 171 | 0.02036 | Crystal structure of Mycobacterium tuberculosis phosphatidylinositol phosphate synthase (PgsA1) with CDP-DAG bound |
| 4o6m-assembly1.cif.gz_A | 0.984 | 0.155 | 174 | 0.03335 | Structure of AF2299, a CDP-alcohol phosphotransferase (CMP-bound) |

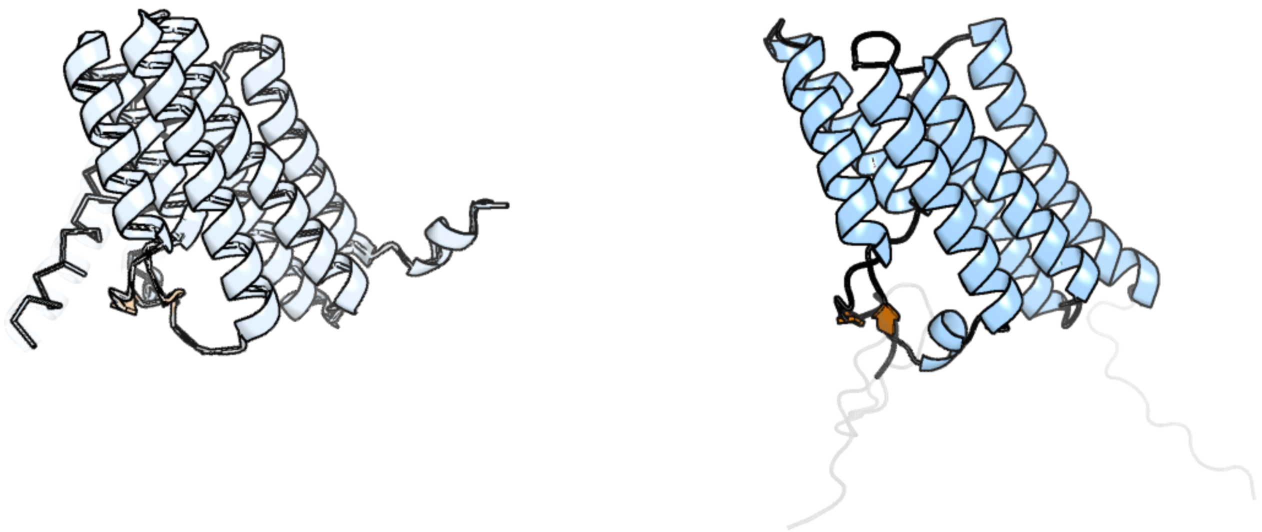

Figure 121: left: reference structure of 7b1l chain A. right: predicted structure of kaa0171457, unaligned sequences are shown as transparent

#### kaa0171475

- Sequence-based annotation for kaa0171475 (predicted in Sulfonolipids from hits to PF00106, UniRef50_A0A383U0M1) is hypothetical protein FNF28_00687 [Cafeteria roenbergensis]
- Best hit was 8g9v chain B: 17-beta-hydroxysteroid dehydrogenase 13

| target | prob | fident | alnlen | evalue | theader |
|---|---|---|---|---|---|
| 8g9v-assembly1.cif.gz_B | 1 | 0.24 | 283 | 8.022e-20 | Crystal structures of 17-beta-hydroxysteroid dehydrogenase 13 |
| 8g9v-assembly1.cif.gz_A | 1 | 0.231 | 289 | 1.353e-19 | Crystal structures of 17-beta-hydroxysteroid dehydrogenase 13 |
| 8g89-assembly1.cif.gz_A | 1 | 0.257 | 284 | 2.282e-19 | HSD17B13 in complex with cofactor and inhibitor |

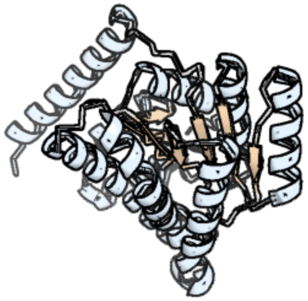

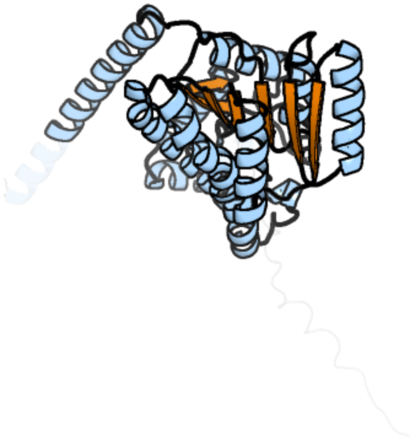

Figure 122: left: reference structure of 8g9v chain B. right: predicted structure of kaa0171475, unaligned sequences are shown as transparent

### kaa0171498

- Sequence-based annotation for kaa0171498 (predicted in Fu-FA,DGTS from hits to PF02353, PF13649) is hypothetical protein FNF28_00708 [Cafeteria roenbergensis]
- Best hit was 5bxy chain B: RNA methyltransferase

| target | prob | fident | alnlen | evalue | theader |
|---|---|---|---|---|---|
| 5bxy-assembly2.cif.gz_B | 1 | 0.313 | 150 | 2.224e-13 | Crystal structure of RNA methyltransferase from Salinibacter ruber in complex with S-Adenosyl-L-homocysteine |
| 5fa8-assembly1.cif.gz_A | 1 | 0.253 | 154 | 1.211e-11 | SAM complex with aKMT from the hyperthermophilic archaeon Sulfolobus islandicu |
| 5jwj-assembly1.cif.gz_A | 1 | 0.253 | 154 | 1.341e-09 | NMR solution structure of a thermophilic lysine methyl transferase from Sulfolobus islandicus |

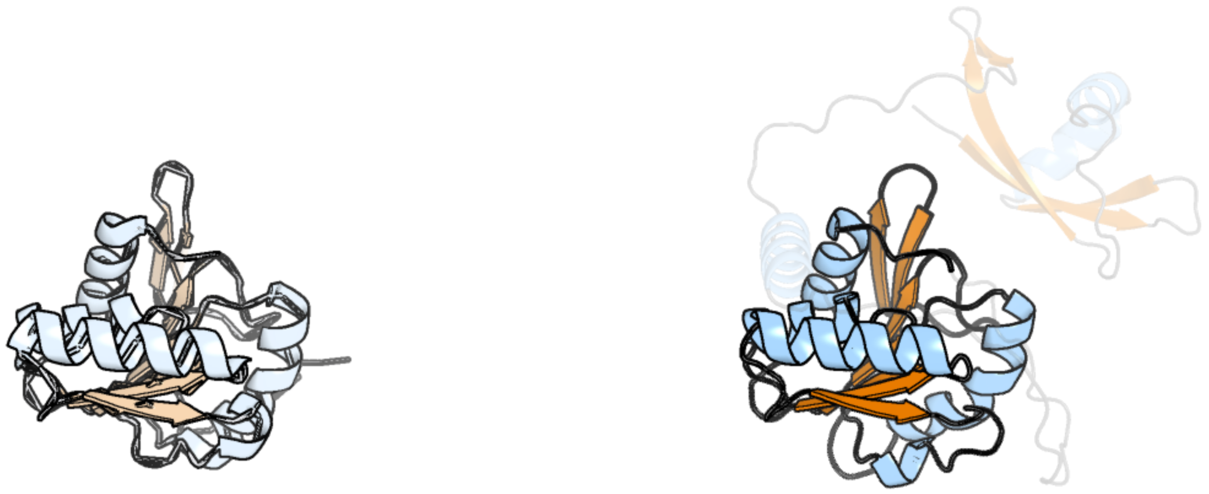

Figure 123: left: reference structure of 5bxy chain B. right: predicted structure of kaa0171498, unaligned sequences are shown as transparent

#### kaa0172037

- Sequence-based annotation for kaa0172037 (predicted in Sulfonolipids from hits to PF00291) is hypothetical protein FNF28_00354 [Cafeteria roenbergensis]
- Best hit was 5ey5 chain D: LBCA-b

| target | prob | fident | alnlen | evalue | theader |
|---|---|---|---|---|---|
| 5ey5-assembly1.cif.gz_D | 1 | 0.665 | 401 | 2.211e-58 | LBCATS |
| 6cut-assembly1.cif.gz_A | 1 | 0.572 | 402 | 5.581e-56 | Engineered Holo TrpB from Pyrococcus furiosus, PfTrpB7E6 with (2S,3S)-isopropylserine bound as the external aldimine |
| 6ami-assembly1.cif.gz_B | 1 | 0.567 | 407 | 2.017e-55 | Engineered tryptophan synthase b-subunit from Pyrococcus furiosus, PfTrpB4D11 with Trp non-covalently bound |

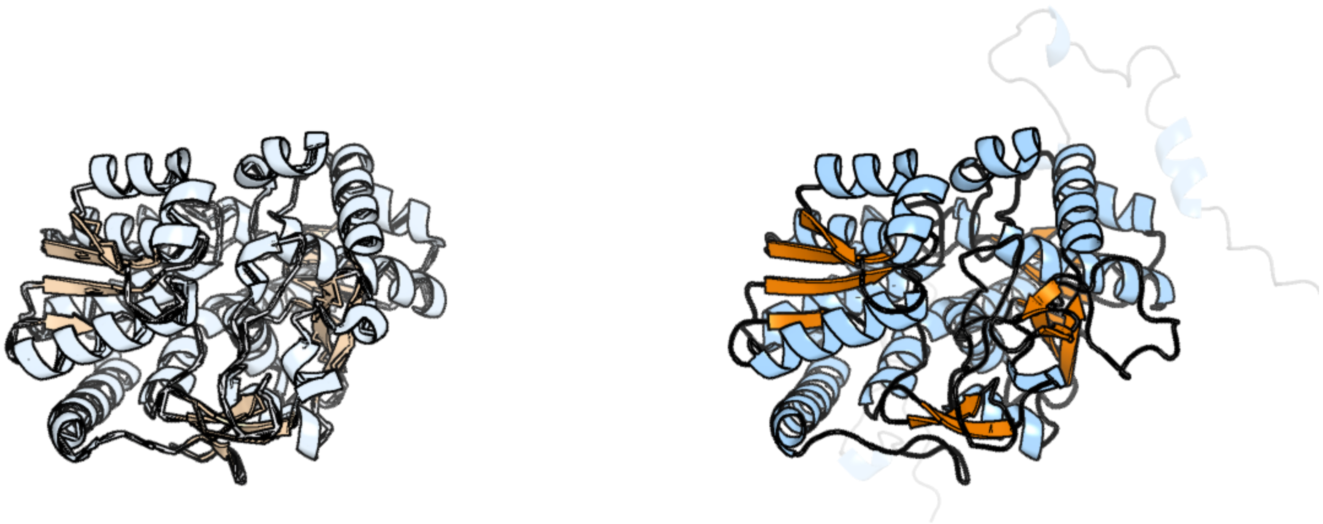

Figure 124: left: reference structure of 5ey5 chain D. right: predicted structure of kaa0172037, unaligned sequences are shown as transparent

### kaa0172061

- Sequence-based annotation for kaa0172061 (predicted in Sulfonolipids from hits to PF00106, UniRef50_A0A383U0M1) is hypothetical protein FNF28_00378 [Cafeteria roenbergensis]
- Best hit was 6g4l chain A: 17-beta-hydroxysteroid dehydrogenase 14

| target | prob | fident | alnlen | evalue | theader |
|---|---|---|---|---|---|
| 6g4l-assembly1.cif.gz_A | 1 | 0.28 | 250 | 2.641e-22 | 17beta-hydroxysteroid Dehydrogenase Type 14 Mutant Y253A in Complex With a Non-steroidal Inhibitor |
| 6qck-assembly1.cif.gz_A | 1 | 0.277 | 252 | 4.457e-22 | 17beta-hydroxysteroid dehydrogenase 14 variant T205 in complex with FB262 |
| 5l7w-assembly1.cif.gz_A | 1 | 0.276 | 253 | 5.624e-22 | 17beta-hydroxysteroid dehydrogenase 14 variant T205 in complex with a non-steroidal inhibitor. |

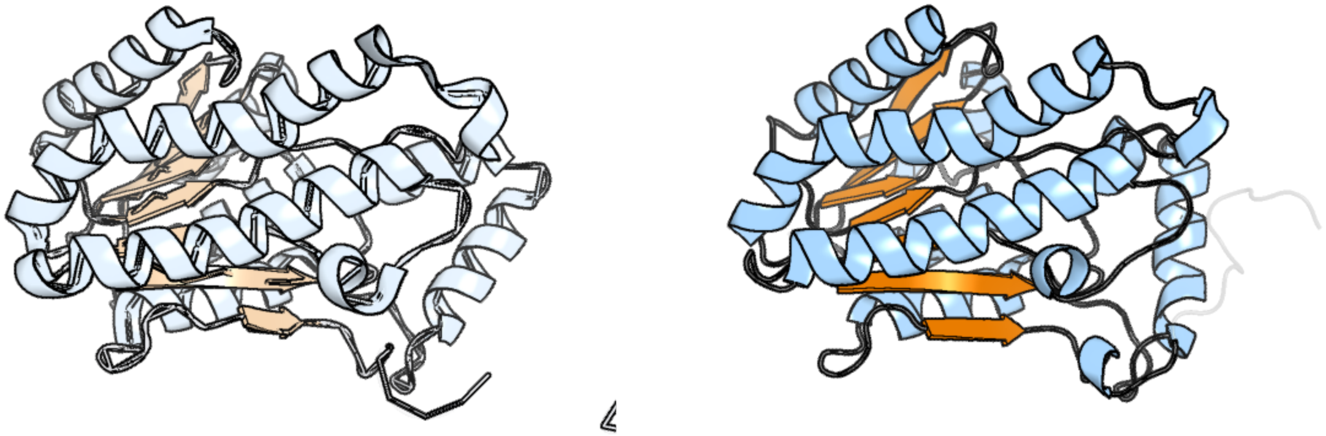

Figure 125: left: reference structure of 6g4l chain A. right: predicted structure of kaa0172061, unaligned sequences are shown as transparent

#### kaa0172071

- Sequence-based annotation for kaa0172071 (predicted in Sulfonolipids from hits to PF00106, UniRef50_A0A383U0M1) is hypothetical protein FNF28_00388 [Cafeteria roenbergensis]
- Best hit was 3lls chain A: 3-ketoacyl-(Acyl-carrier-protein) reductase

| target | prob | fident | alnlen | evalue | theader |
|---|---|---|---|---|---|
| 3lls-assembly1.cif.gz_A | 1 | 0.278 | 212 | 3.171e-14 | Crystal structure of 3-ketoacyl-(acyl-carrier-protein) reductase from Mycobacterium tuberculosis |
| 6zzs-assembly2.cif.gz_E-2 | 1 | 0.232 | 211 | 5.642e-13 | Crystal structure of (R)-3-hydroxybutyrate dehydrogenase from Acinetobacter baumannii complexed with NAD+ and 3-oxovalerate |
| 7e6o-assembly1.cif.gz_C | 1 | 0.241 | 207 | 6.755e-13 | Crystal structure of polyol dehydrogenase from Paracoccus denitrificans |

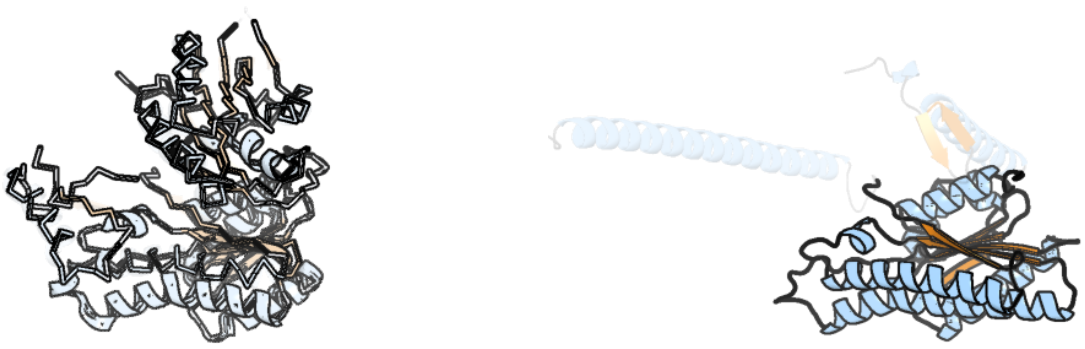

Figure 126: left: reference structure of 3lls chain A. right: predicted structure of kaa0172071, unaligned sequences are shown as transparent

### kaa0172187

- Sequence-based annotation for kaa0172187 (predicted in Sulfonolipids from hits to PF00155, UniRef90_A0A7Z8YEE0, UniRef90_D4IKV7) is hypothetical protein FNF28_00190 [Cafeteria roenbergensis]
- Best hit was 7yjo chain B: Long chain base biosynthesis protein 2a

| target | prob | fident | alnlen | evalue | theader |
|---|---|---|---|---|---|
| 7yjo-assembly1.cif.gz_B | 1 | 0.541 | 469 | 8.938e-57 | Cryo-EM structure of the monomeric atSPT-ORM1 (LCB2a-deltaN5) complex |
| 7yjm-assembly1.cif.gz_B | 1 | 0.541 | 469 | 1.575e-55 | Cryo-EM structure of the monomeric atSPT-ORM1 complex |
| 7k0i-assembly1.cif.gz_B | 1 | 0.51 | 476 | 2.409e-55 | Human serine palmitoyltransferase complex SPTLC1/SPLTC2/ssSPTa |

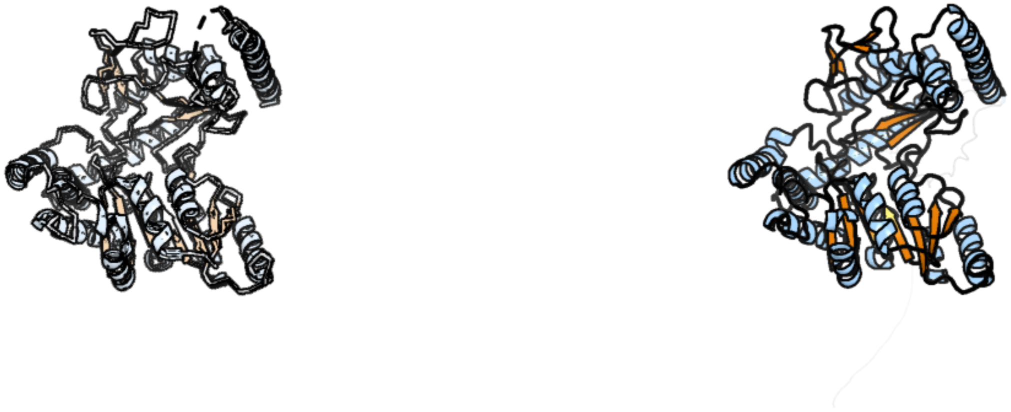

Figure 127: left: reference structure of 7yjo chain B. right: predicted structure of kaa0172187, unaligned sequences are shown as transparent

### kaa0172196

- Sequence-based annotation for kaa0172196 (predicted in PE_PMME from hits to PF04191) is hypothetical protein FNF28_00199 [Cafeteria roenbergensis]
- Best hit was 5v7p chain A: Protein-S-isoprenylcysteine O-methyltransferase

| target | prob | fident | alnlen | evalue | theader |
|---|---|---|---|---|---|
| 5v7p-assembly1.cif.gz_A | 1 | 0.316 | 221 | 3.251e-12 | Atomic structure of the eukaryotic intramembrane Ras methyltransferase ICMT (isoprenylcysteine carboxyl methyltransferase), in complex with a monobody |
| 5vg9-assembly1.cif.gz_A | 1 | 0.321 | 221 | 5.644e-11 | Structure of the eukaryotic intramembrane Ras methyltransferase ICMT (isoprenylcysteine carboxyl methyltransferase) without a monobody |
| 4a2n-assembly1.cif.gz_B | 1 | 0.172 | 215 | 0.003496 | Crystal Structure of Ma-ICMT |

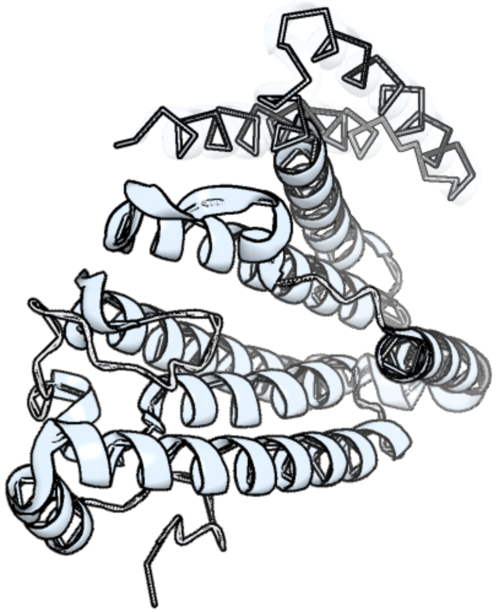

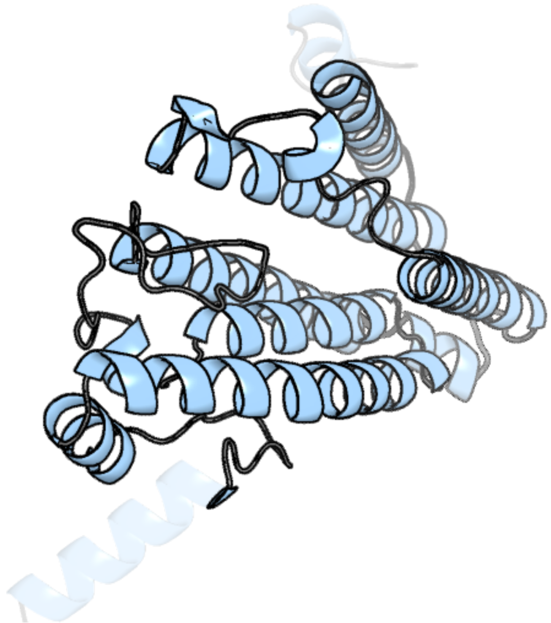

Figure 128: left: reference structure of 5v7p chain A. right: predicted structure of kaa0172196, unaligned sequences are shown as transparent

#### kaa0172207

- Sequence-based annotation for kaa0172207 (predicted in Sulfonolipids from hits to PF00291, UniRef50_A0A2D9B818) is hypothetical protein FNF28_00210 [Cafeteria roenbergensis]
- Best hit was 2gn2 chain A: Threonine dehydratase catabolic

| target | prob | fident | alnlen | evalue | theader |
|---|---|---|---|---|---|
| 2gn2-assembly1.cif.gz_A | 1 | 0.306 | 329 | 1.543e-25 | Crystal structure of tetrameric biodegradative threonine deaminase (TdcB) from Salmonella typhimurium in complex with CMP at 2.5A resolution (Hexagonal form) |
| 7nbf-assembly1.cif.gz_AAA | 1 | 0.285 | 333 | 1.443e-24 | Crystal structure of human serine racemase in complex with DSiP fragment Z126932614, XChem fragment screen. |
| 5cvc-assembly2.cif.gz_B-2 | 1 | 0.291 | 329 | 1.443e-24 | Structure of maize serine racemase |

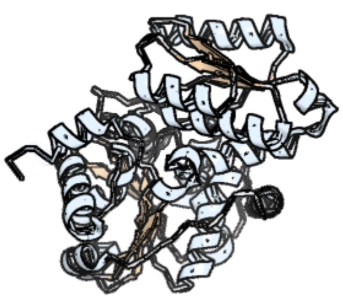

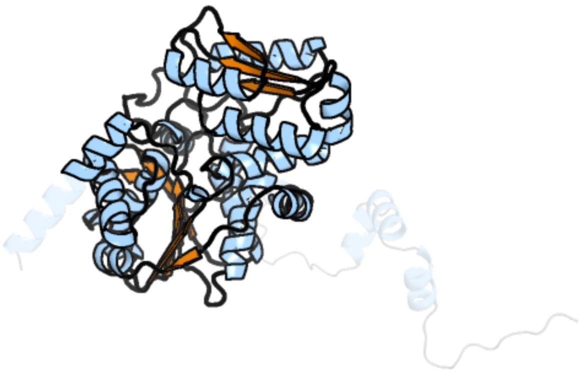

Figure 129: left: reference structure of 2gn2 chain A. right: predicted structure of kaa0172207, unaligned sequences are shown as transparent

### kaa0172390

- Sequence-based annotation for kaa0172390 (predicted in DGTS from hits to PF13649) is hypothetical protein FNF28_00073 [Cafeteria roenbergensis]
- Best hit was 6uv6 chain B: D-glucose O-methyltransferase

| target | prob | fident | alnlen | evalue | theader |
|---|---|---|---|---|---|
| 6uv6-assembly2.cif.gz_B | 1 | 0.191 | 256 | 5.98e-11 | AtmM with bound rebeccamycin analogue |
| 6f5z-assembly1.cif.gz_A | 1 | 0.156 | 250 | 6.754e-11 | Complex between the Haloferax volcanii Trm112 methyltransferase activator and the Hvo_0019 putative methyltransferase |
| 2ip2-assembly1.cif.gz_B | 1 | 0.152 | 197 | 1.584e-10 | Structure of the Pyocyanin Biosynthetic Protein PhzM |

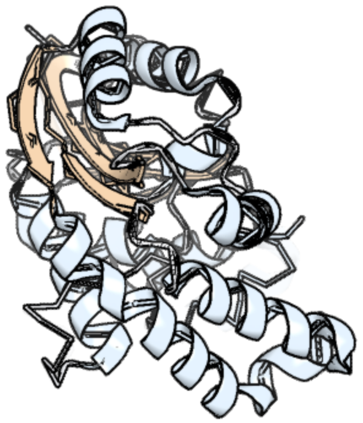

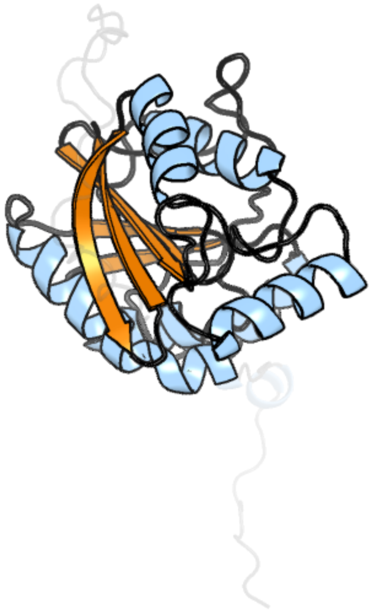

Figure 130: left: reference structure of 6uv6 chain B. right: predicted structure of kaa0172390, unaligned sequences are shown as transparent
